# ELLA: Modeling Subcellular Spatial Variation of Gene Expression within Cells in High-Resolution Spatial Transcriptomics

**DOI:** 10.1101/2024.09.23.614515

**Authors:** Jade Xiaoqing Wang, Xiang Zhou

## Abstract

Spatial transcriptomic technologies are becoming increasingly high-resolution, enabling precise measurement of gene expression at the subcellular level. Here, we introduce a computational method called subcellular expression localization analysis (ELLA), for modeling the subcellular localization of mRNAs and detecting genes that display spatial variation within cells in high-resolution spatial transcriptomics. ELLA creates a unified cellular coordinate system to anchor diverse cell shapes and morphologies, utilizes a nonhomogeneous Poisson process to model spatial count data, leverages an expression gradient function to characterize subcellular expression patterns, and produces effective control of type I error and high statistical power. We illustrate the benefits of ELLA through comprehensive simulations and applications to four spatial transcriptomics datasets from distinct technologies, where ELLA not only identifies genes with distinct subcellular localization patterns but also associates these patterns with unique mRNA characteristics. Specifically, ELLA shows that genes enriched in the nucleus exhibit an abundance of long noncoding RNAs or protein-coding mRNAs, often characterized by longer gene lengths. Conversely, genes containing signal recognition peptides, encoding ribosomal proteins, or involved in membrane related activities tend to enrich in the cytoplasm or near the cellular membrane. Furthermore, ELLA reveals dynamic subcellular localization patterns during the cell cycle, with certain genes showing decreased nuclear enrichment in the G1 phase while others maintain their enrichment patterns throughout the cell cycle. Overall, ELLA represents a calibrated, powerful, robust, scalable, and versatile tool for modeling subcellular spatial expression variation across diverse high-resolution spatial transcriptomic platforms.

## Introduction

Spatial transcriptomics is a collection of new genomics technologies designed to measure gene expression within tissues while preserving spatial localization information. Recent technological advancements have substantially improved the spatial resolution of spatial transcriptomics, facilitating expression measurements at cellular and subcellular levels. Specifically, *in situ* RNA-sequencing techniques, such as ISS [1], FISSEQ [2], STARmap [3], and Ex-seq [4], achieve a spatial resolution under 1 μm, which is much smaller than the size of a typical cell. Recent high-throughput sequencing-based techniques, such as Seq-Scope [5], VisiumHD [6], Open-ST [7], and Stereo-seq [8], offer spatial resolutions in the range of 0.5-2 μm. *In situ* imaging techniques, such as MERFISH [9], SeqFISH+ [10], MERSCOPE [11], CosMx [12], and 10X Xenium [13], provide spatial resolutions as fine as 0.1-0.2 μ m (Fig. S1). Together, these high-resolution spatial transcriptomics technologies have enabled expression measurement at subcellular resolution, providing unprecedented opportunities to interrogate the intracellular localization and distribution of mRNAs within cells.

The intracellular localization and distribution of mRNAs are vital for cellular functions. They ensure the targeted delivery of mRNAs and facilitate localized protein synthesis, enabling precise regulation of gene expression within specific subcellular compartments. The spatial localization of mRNAs empowers cells to respond rapidly to local cues and signals, adapting effectively to changing environments and supporting specialized cellular functions [14]. For example, the localization of mRNAs encoding for β-actin at the leading edges of fibroblasts or the lamellipodia of myoblasts ensures localized protein synthesis of actin, supporting proper cell polarity and motility [15]. In addition, the spatial localization of mRNA contributes to cellular organization and differentiation, aiding in the establishment and maintenance of distinct cellular identities and functions, influencing asymmetric cell division and cell fate determination across various organisms. For example, the spatially localized expression of *Oskar* at the posterior end of the embryo is essential for the development and assembly of the germ plasm in *Drosophila*, facilitating germ cell formation [16]. As another example, *Ash* mRNA localizes to the bud tip in *S. cerevisiae* to establish asymmetry of HO endonuclease gene expression, which is important for mating type switching [17]. Given the importance of proper mRNA spatial localization, their misplacement often leads to detrimental effects and has been associated with multiple diseases [16]. For example, disruptions in axonal mRNA transport and localization contribute to neurodegeneration in Huntington’s disease [18]. Therefore, understanding the spatial localization and distribution of mRNA within cells is crucial for unraveling the complexity of cellular structure and function, as well as for elucidating the cellular mechanisms underlying disease etiology.

Despite its importance, however, characterizing the subcellular spatial organization of mRNAs in high-resolution spatial transcriptomics turns out to be a computationally challenging task. Only two methods have been developed for this purpose, each with its own limitations. Specifically, Bento [19] employs pre-trained random forest classifiers to categorize each gene into five pre-defined subcellular RNA localization patterns, while SPRAWL [20] relies on four metrics to identify four pre-specified subcellular patterns. However, both methods are limited to imaging-based spatial transcriptomics data, failing to leverage the vast amount of high-resolution spatial transcriptomics obtained from recent sequencing-based technologies. Additionally, they are constrained to detect genes with pre-defined localization patterns, thus limiting the discovery of any new spatial localization patterns and suffering from low statistical power. Besides these major limitations, Bento requires nuclear boundary information, which may not be readily available in some spatial transcriptomics datasets. In addition, Bento is only applicable to analyzing a single cell and lacks the ability to borrow the spatial localization pattern shared across multiple cells. Conversely, SPRAWL is only applicable to analyzing multiple cells, not a single cell, and is unable to distinguish between enrichment and depletion in the pre-specified localization patterns due to the nature of its two-sided tests.

To address the above limitations, we present subcellular Expression LocaLization Analysis (ELLA), a computational method for modeling the subcellular localization of mRNAs and detecting genes that display spatial variation within cells across a range of high-resolution spatial transcriptomics technologies. ELLA comes with three unique features. First, it creates a unified cellular coordinate system, which allows for anchoring diverse cell shapes and morphologies regardless of the spatial transcriptomics technology, thus enabling the joint modeling of cells with distinct shapes. Second, it constructs a novel nonhomogeneous Poisson process model to directly and explicitly model the spatial occurrence of expression measurements within cells, thus facilitating powerful and effective modeling of both spatial count data from sequencing-based techniques and spatial binary data from imaging-based techniques. Finally, it devises an expression intensity function to model the subcellular spatial distribution of mRNAs along the cellular radius, which, when paired with a range of kernel functions, is capable of capturing a wide variety of expression distribution patterns within cells without the need to restrict to pre-defined localization patterns. As a result, ELLA can be applied to an arbitrary number of cells and detect a wide variety of subcellular localization patterns across diverse spatial transcriptomic techniques, all with effective type I error control and high statistical power. With a computationally efficient algorithm, ELLA is also scalable to tens of thousands of genes across tens of thousands of cells.

We illustrate the benefits of ELLA through comprehensive simulations and applications to four spatial transcriptomics datasets. In the real data applications, ELLA not only identifies genes with distinct subcellular localization patterns but also reveals that these patterns are associated with unique mRNA characteristics. Specifically, genes enriched in the nucleus show an abundance of long noncoding RNAs (lncRNAs) and protein-coding mRNAs, often characterized by longer gene lengths. Conversely, genes containing signal recognition peptides, encoding ribosomal proteins, or involved in membrane related activities such as synaptic transmission and G protein coupled receptor activities, tend to enrich in the cytoplasm or near the cellular membrane. Moreover, genes exhibit dynamic subcellular localization during the cell cycle, with some showing decreased nuclear enrichment in the G1 phase, while others maintain their patterns of enrichment regardless of cell cycle phases.

## Results

### Method overview

ELLA is described in Methods, with its technical details provided in Supplementary Notes and method schematic displayed in Fig. 1a. Briefly, ELLA is a statistical method for modeling the subcellular localization of mRNAs and detecting spatially variable genes with subcellular spatial expression patterns in high-resolution spatial transcriptomics (Fig. S1). ELLA examines one gene at a time, creates a unified cellular coordinate system through defining a cellular radius in each cell that points from the center of the nucleus towards the cellular boundary, relies on a nonhomogeneous Poisson process (NHPP) to capture the spatial distribution of expression measurements within cells, devises an expression intensity function and computes a P value to capture any subcellular expression patterns observed along the cellular radius. ELLA is capable of borrowing information across cells through a joint likelihood framework to substantially improve detection power, while taking advantage of multiple intensity kernel functions to capture the distinct subcellular expression patterns that may be encountered in various biological settings to ensure robust performance. In addition, ELLA relies on a fast binning algorithm for approximate position computation and utilizes Adam optimization for scalable inference. As a result, ELLA is computationally efficient and is easily scalable to tens of thousands of genes measured in tens of thousands of cells. ELLA is implemented in Python, freely accessible from https://xiangzhou.github.io/software/.

**Fig. 1.**
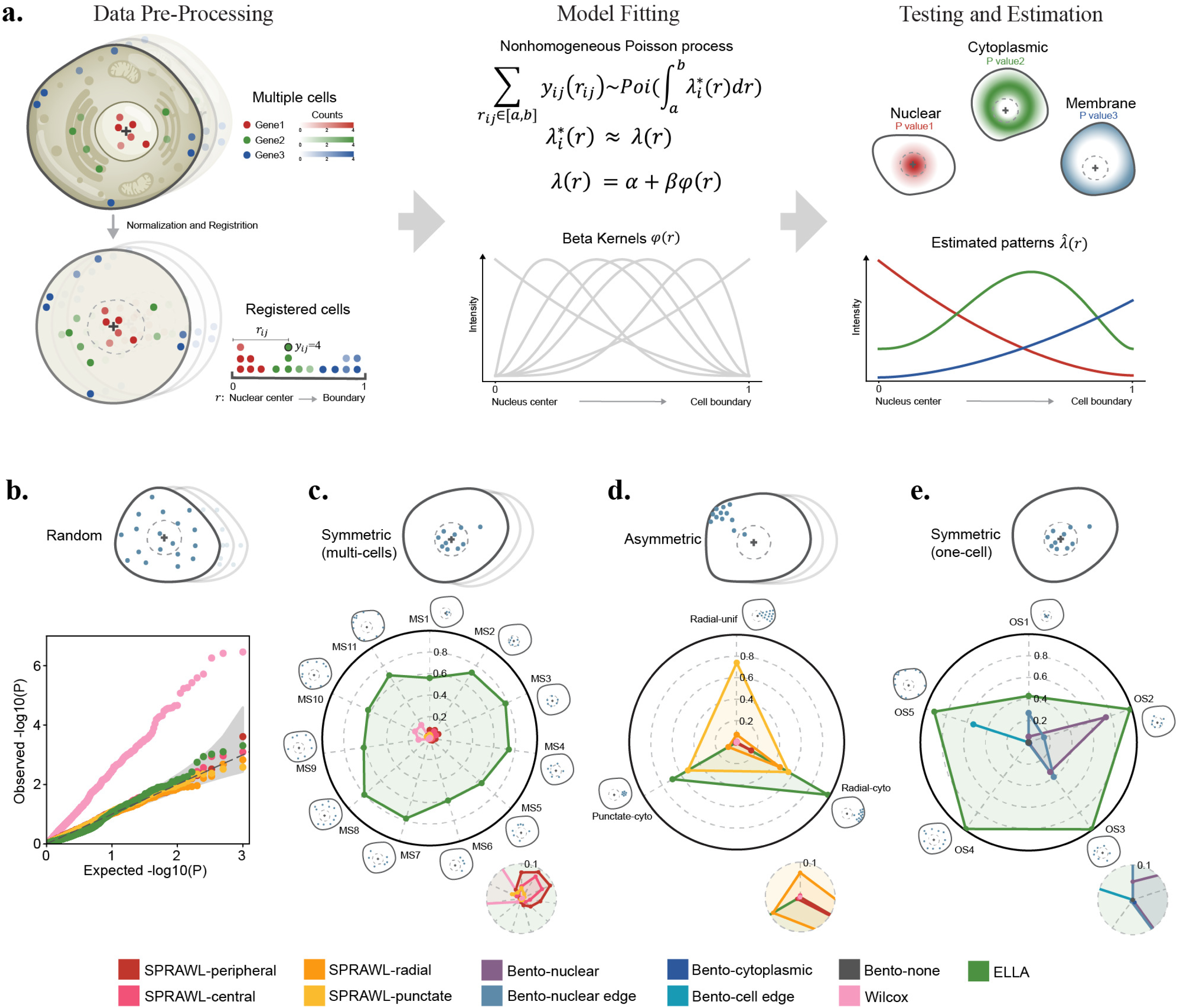
Schematic of ELLA and simulation results. **a.** ELLA is a method designed for modeling the subcellular localization of mRNAs and detecting genes that display spatial variation within cells in high-resolution spatial transcriptomics. ELLA takes as inputs the high-resolution spatial gene expression data along with nuclear center and cell segmentation information. It first performs data pre-processing including normalization and registration to create a unified cellular coordinate system to anchor diverse cell shapes and morphologies through defining a cellular radius in each cell that points from the center of the nucleus towards the cellular boundary. It then fits a nonhomogeneous Poisson process model for each gene to capture its spatial distribution within cells, computes a P value to capture any subcellular expression pattern observed along the cellular radius, and estimates such pattern in the form of estimated pattern expression intensity and pattern score. ELLA is capable of borrowing information across cells through a joint likelihood framework to substantially improve detection power, while taking advantage of multiple intensity kernel functions to capture the distinct subcellular expression patterns that may be encountered in various biological settings to ensure robust performance. **b.** Quantile-quantile plots of the expected and observed -log10 P values for different methods in the baseline null simulation, where gene expression is randomly distributed spatially within cells. ELLA was compared to SPRAWL, Bento, and Wilcox. **c.** Radar plots show the power of different methods in the alternative simulations with multiple cells across eleven symmetric subcellular expression patterns, where gene expression is enriched in specific subcellular regions within cells. ELLA was compared to SPRAWL and Wilcox and power was evaluated based on 5% FDR. **d.** Radar plots of the power of different methods in the alternative simulations with multiple cells across three asymmetric subcellular expression patterns, where gene expressions exhibit distinct asymmetric patterns. ELLA was compared to SPRAWL and Wilcox and power was evaluated based on 5% FDR. **e.** Radar plots show the power of different methods in the additional alternative simulations with one cell across five symmetric subcellular expression patterns, where gene expression is enriched in specific subcellular regions within cells. ELLA was compared to Bento and power was evaluated based on 5% FDR.

### Simulations

We performed comprehensive simulations on imaging-based spatial transcriptomics to evaluate the performance of ELLA and compared it with three methods. The three methods include SPRAWL [20], Bento [19], and Wilcox, where Wilcox denotes a modified Wilcoxon rank sum test [21] that uses expression measurements normalized by the area of subcellular regions to examine the difference in expression between nuclear and cytoplasmic areas. All methods examine one gene at a time and all methods except Bento produce a P value for each gene; Bento outputs five prediction probabilities for five pre-specified cellular localization patterns, which cannot be converted to a P value. Among these methods, ELLA can analyze either one or multiple cells; SPRAWL and Wilcox can only analyze multiple cells; and Bento can only analyze one cell. Therefore, we compared ELLA with SPRAWL and Wilcox in all our main simulations on multiple cells while compared ELLA with Bento in additional simulations on only one cell. Unlike ELLA and SPRAWL, both Bento and Wilcox require nuclear boundary information in addition to cell boundary information (Tab. S1). We provide the actual nuclear boundary information to Bento and Wilcox, although this information may not be readily available in certain sequencing-based techniques such as Seq-Scope [5] and Stereo-seq [8] and may not be accurately inferred in other techniques.

Simulation details are provided in Methods. Briefly, we sampled *n* different embryonic fibroblast cells from seqFISH+ data (Fig. S2) and simulated expression counts for 1,000 genes to be spatially distributed within these cells. We examined type I error control of different methods in null simulations, where the simulated gene expression counts are randomly distributed spatially within each cell without any specific subcellular spatial expression patterns (Fig. 1b, S3). We also examined the power of different methods in alternative simulations, where the simulated gene expression counts are enriched in specific subcellular regions within the cells, exhibiting either symmetric (consisting of eleven distinct symmetric patterns; Fig. 1c, S4-5) or asymmetric patterns (three distinct asymmetric patterns, Fig. 1d, S6). In the simulations, we first created a baseline setting and then varied the number of cells (*n*), the gene expression level (*m*), and in the alternative settings, the strength of the subcellular expression patterns (*S*; Methods), one at a time on top of the baseline setting, to create additional settings. In total, we examined 13 null and 40 alternative settings, with 1,000 replicates per setting.

In the null simulations, the P values from ELLA are well calibrated across settings, and so are the P values from SPRAWL, although SPRAWL failed to produce P values for the radial and punctate metrics in *m*=1 (one count per cell) settings (Fig 1b, S7-8). The inability of SPRAWL to produce P values in these settings arises from its inability to make use of cells with less than two counts of the gene, which constitutes a large fraction of gene-cell combinations in the real data (e.g. 57.8% in the MERFISH data) [22]. Wilcox yielded inflated P values, especially in settings where the gene expression level is low, or where the number of cells is large (Fig 1b, S7-8). The P value inflation observed in Wilcox suggests that the simple normalization procedure and the non-parametric Wilcoxon test are not sufficient to control for variance heterogeneity and subsequently type I error (Fig. S9, Tab. S2).

In the alternative simulations, because some methods failed to control for type I error, we evaluated power based on a fixed false-discovery rate (FDR) to ensure a fair comparison across methods (Methods). We first examined the eleven subcellular expression patterns in the symmetric pattern category, including two patterns with nucleus enrichment, two patterns with nuclear edge enrichment, five patterns with cytoplasmic enrichment, and two patterns with membrane enrichment. Based on an FDR threshold of 0.05, ELLA achieves consistently higher power (average=0.63, range=0.41-0.80) than the other methods (SPRAWL: average=0.04, range=0.00-0.09; Wilcox: average=0.04, range=0.00-0.15) in detecting each of the eleven patterns (Fig. 1c). For SPRAWL, its radial and punctate metrics tend to exhibit very low power in detecting any of the patterns (average=0.01, range=0.00-0.03), presumably because these metrics are not well suited for detecting symmetric patterns. The peripheral and central metrics of SPRAWL have low power for detecting the cytoplasmic enrichment patterns (average=0.01, range=0.00-0.04) but have slightly higher powers for detecting the membrane and nuclear enrichment patterns (average=0.05, range=0.01-0.09), as one might expect. Also as expected, the power of ELLA, SPRAWL, and Wilcox all improves with increasing number of cells, increasing expression level, and increasing pattern strength across all eleven patterns, although the power of ELLA improves much faster compared to the other two methods (Fig. S10). For example, at an FDR of 0.05, the power of ELLA in detecting the first nucleus pattern is 0.01 with 10 cells but increases to 1.00 with 300 cells, while the power SPRAWL’s central metric only increases from 0.01 to 0.42 and the power of SPRAWL’s peripheral metric only increases from 0.00 to 0.66. The exceptions are Wilcox and SPRAWL’s radial metric, whose power for detecting nucleus patterns remains below 0.05 and barely improves as the number of cells increases.

ELLA is also more powerful than the other methods in detecting two of the three asymmetric subcellular expression patterns. These include the radial-cyto and punctate-cyto patterns, where gene expression is enriched in either a circular sector or a small subcellular disc in the cytoplasm (Fig. 1d). Specifically, for the radial-cyto pattern, ELLA achieved a power of 0.55 while Wilcox achieved a power of 0.00. For SPRAWL, its peripheral, central, radial, and punctate metrics achieved a power of 0.10, 0.00, 0.16, and 0.23, respectively. For the punctate-cyto pattern, ELLA achieved a power of 0.65 while Wilcox had zero power. For SPRAWL, its peripheral, central, radial, and punctate metrics achieved a power of 0.17, 0.00, 0.16, and 0.39, respectively (Fig. 1d). Certainly, because ELLA models expression patterns along the cellular radius, it is not powered to detect radial-unif asymmetric pattern, where gene expression is enriched in a circular sector of the cell completely uniformly (Fig. 1d), a scenario unlikely in practical biological applications.

Importantly, ELLA not only achieves high power in detecting genes with various subcellular expression patterns but also accurately estimates these patterns (Fig. S11-12). Specifically, the average KL-divergences achieved by ELLA for estimating the two pattern categories are 0.12 and 0.29, respectively (Tab. S3). To further summarize the observed subcellular pattern, ELLA computes a subcellular pattern score for each gene. This score represents the relative position of subcellular expression enrichment, with zero indicating enrichment in the cell nucleus and one indicating enrichment on the cell membrane; Methods). The majority of the pattern scores (77%) are within 0.1 of the truth across the three pattern categories, underscoring the accuracy of ELLA (Fig. S13-14, Tab. S4).

We performed additional simulations with only one cell in order to compare ELLA with Bento (Fig. 1e). Bento is capable of detecting five pre-specified patterns including enrichment in nucleus, nuclear edge, cytoplasm, cell boundary, and none. To favor the comparison towards Bento, we focused on comparing ELLA with Bento under five symmetric patterns that Bento specifically models, where gene expression is enriched in nucleus (including 2 patterns), nuclear edge (1), cytoplasm (1), or cellular boundary (1) under a relatively high expression level (*m*=30) and a high pattern strength (*S*=9) (Fig. S15). Because Bento cannot produce P values, we used the prediction probabilities output from Bento to rank genes, with which we measured powers based on FDR (Methods). We are able to compute FDR for Bento in simulations only because we know the truth, which is certainly unknown for any real data applications. In the simulations, ELLA achieves high power (Fig. 1e, average=0.86, range=0.43-0.99) and accuracy (Fig. S16-17, Tab. S5) across all five patterns, consistently outperforming Bento (average=0.10, range=0.00-0.75).

### Seq-Scope mouse liver data

We applied ELLA to analyze four published datasets obtained using different high-resolution spatial transcriptomics technologies (Methods). The four datasets include a liver data by Seq-Scope [5], an embryo data by Stereo-seq [8], an NIH/3T3 embryonic fibroblast cell line data by seqFISH+ [10], and a brain data by MERFISH [22].

We first analyzed the Seq-Scope mouse liver data (Fig. 2a, S18-30), which contains 497 to 1,349 genes measured on 870 cells from four cell types, with 82 to 276 cells per cell type (Fig. S31-32). The four cell types include periportal hepatocyte (PP; *n*=276) and pericentral hepatocyte (PC; *n*=276) in normal mice, and PP (*n*=236) and PC (*n*=82) cells in early-onset liver failure mice (TD, [23]; Fig. S33). We were only able to apply ELLA to the data as SPRAWL and Bento are not applicable to sequencing-based data and the nuclear boundary information required for Wilcox and Bento was not available.

**Fig. 2.**
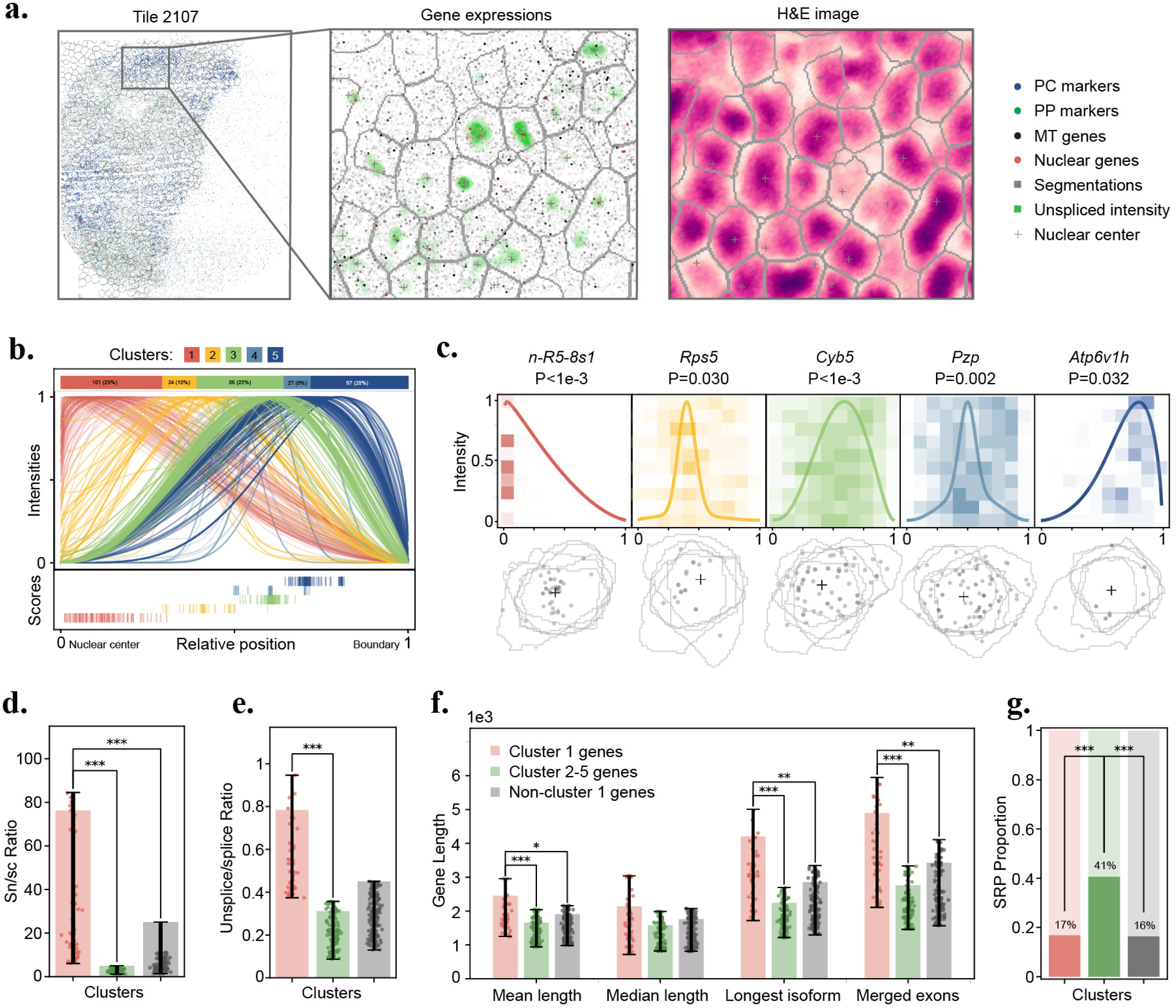
Seq-Scope mouse liver data analysis. **a.** Data snapshot for tissue tile 2107. Left panel displays the full tile with expression from four gene sets (PC marker genes, blue dots; PP marker genes, green dots; Tab. S10; mitochondria genes, black dots; nuclear genes, red dots; Tab. S13) along with cell segmentation boundaries. Middle panel zooms into a subregion and displays unspliced expression densities (colored green), nuclear centers (crosses), gene expression from the four lists, along with cell segmentation boundaries. Right panel displays the H&E staining image of the subregion. **b.** Estimated spatial expression pattern for genes in each of the five gene pattern clusters identified by ELLA. Upper panel shows the number and proportion of genes across five pattern clusters. Middle panel displays the estimated expression intensities for genes across clusters. Lower panel displays the estimated pattern score for genes across clusters. **c.** Example genes and cells for the five pattern clusters. One example gene is shown for each pattern cluster. Upper panel shows the gene name, ELLA P value, and the estimated expression intensities overlayed on the density heat map. Each row of the density heat map visualizes the number of counts standardized by area in 10 randomly selected cells across relative positions with intervals of 0.1. Lower panel displays the expressions of the corresponding genes within five selected cells, overlayed with cell boundaries and aligned nuclear centers (crosses). **d.** Bar plot shows the average sn/sc RNA ratio in the form of snRNA expression level normalized by scRNA expression level across genes in pattern clusters 1 (red), 2-5 (green), and non-cluster 1 genes (i.e. clusters 2-5 plus the nonsignificant genes; grey). Nuclear enriched genes (cluster 1) tend to exhibit higher relative snRNA expression levels. **e.** Bar plot shows the average unspliced/spliced expression ratio across genes in pattern clusters 1, 2-5, and non-cluster 1 genes. Nuclear enriched genes (cluster 1) tend to exhibit higher unspliced/spliced expression ratios. **f.** Bar plot displays the average gene length, measured by four metrics (x-axis), across genes in pattern clusters 1, 2-5, and non-cluster 1 genes. Nuclear enriched genes (cluster 1) tend to exhibit longer gene lengths. **g.** Bar plot displays the proportions of SRP-coded genes for genes in pattern clusters 1, 2-5, and non-cluster 1 genes. Cytoplasmic enriched genes (clusters 2-5) frequently encode SRP. Statistical significance for pair-wise comparisons (*: <0.05; **: <0.01; ***: <0.001) is based on Mann-Whitney U test (**d**-**f**) or Fisher’s exact test (**g**). The error bars represent the 25th and 75th percentiles, and data points beyond this range are not included (**d**-**f**).

At an FDR of 5%, ELLA identified 84, 123, 98, and 40 genes that display subcellular expression patterns in normal PP, PC cells and TD PP, PC cells, respectively. 77 of these genes, including one transcription factor (*Mlxipl*), were detected in two or more cell types. Based on their subcellular spatial expression patterns, we clustered the detected genes into five distinct pattern clusters (Fig. 2b, Method): 101 genes (29%) display a nuclear expression pattern (clusters 1), 34 (10%) genes display a nuclear edge expression pattern (cluster 2), and 210 genes (61%) display one of the three cytoplasmic expression patterns near the cellular membrane (cluster 3-5). Example cells from the five clusters are shown in Fig. 2c.

The detected genes from ELLA allow us to comprehensively investigate the properties of genes that display distinct enrichment patterns within cells. For nuclear genes (clusters 1) with subcellular enrichment near the nuclear center, we found them to have significantly higher snRNA expression in a similar cell type from a separate study [24] (clusters 1 vs clusters 2-5 fold enrichment=15.31, Mann-Whitney U test P value = 5e-22; cluster 1 vs all the remaining genes - consisting of cluster 2-5 genes and nonsignificant genes, fold enrichment=4.80, P value=1e-11; Fig. 2d) with significantly higher unsplice/splice ratio supporting their nuclear enrichment (cluster 1 vs clusters 2-5 fold enrichment=2.52, Mann-Whitney U test P value=8e-21; cluster 1 vs all the other genes fold enrichment=1.08, P value=0.58; Fig. 2e). For genes with subcellular enrichment in the cytoplasm (cluster 4-5), we found them to frequently encode a signal recognition peptide (SRPs; proportion=40.57%) as compared to the genes in the nuclear cluster 1 (proportion=16.83%; Fisher’s exact test P value=2e-5) or the remaining genes (proportion=14.95%; P value=9e-21; Fig. 2g). SRPs are short sequence segments located at the N-termini of newly synthesized proteins that are translated at the endoplasmic reticulum (ER) and sorted towards the secretory pathway [25]. Importantly, we found that nuclear genes (cluster 1) have significantly longer gene lengths compared to genes in the other clusters or the remaining genes, both in terms of the average isoform length (Mann-Whitney U test P value=6e-4 and 0.02), the longest isoform length (P value=1e-6 and 4e-3), and the total length across exons (P value=2e-7 and 6e-3; Fig. 2f). These new findings suggest that long genes may require additional time to be transcribed and exported [26] and their enrichment in the nucleus may serve as a reservoir so that they can be quickly exported to the cytoplasm for translation in response to stimuli [27]. Notably, this novel discovery is consistently observed across the other three datasets we analyzed, as presented in later sections.

We explored additional biological insights by focusing on the normal PC cell type, which has the largest number of genes with subcellular spatial expression patterns, to carefully examine the 123 genes detected by ELLA (Fig. S19). Among the 34 nuclear (cluster 1) genes (Fig. S34, Tab. S6), three of them (*Malat1*, *Neat1* and *Gm13775*) are long non-coding RNAs that are previously known to be localized to the nucleus [28]. 30 of them are protein encoding genes including two previously known nuclear-enriched mRNAs *Chd9* and *Ppara*, dovetailing recent findings that retention of mRNAs in the nucleus may help buffer noise in the stochastic mRNA production process [21]. Seven of them (*Malat1*, *Neat*, *n-R5-8s1*, *Gm24601*, *Mlxipl*, *Mafb*, and *Echdc2*) were also found among the top 10 nuclear-enriched genes identified in the original Seq-Scope study, which explicitly searched for genes enriched within 10μm from the nuclear center [5]. Among the seven genes, four encode transcription factors or proteins with transcription factor activity. For example, *Mlxipl*, one of these genes, is a transcription factor retained in the nuclear speckles in the liver [29]. Finally, all nine significant mitochondrial genes were detected as cytoplasmic localized (cluster 3; Fig. S35a, Tab. S7) and all three significant PC cell type marker genes were detected as cytoplasmic or membrane localized (clusters 3 and 5; Fig. S35b, Tab. S7).

### Stereo-seq mouse embryo data

Next, we analyzed the Stereo-seq mouse embryo data, focusing on two major cell types localized in the cardiothoracic region on slice E1S3 (Fig. 3a, S36-37): precursor muscle cells, or myoblasts (596 cells with 2,008 genes); and mature muscle cells, or cardiomyocytes (553 cells with 1,743 genes; Fig. S38). We were only able to apply ELLA to the data as SPRAWL and Bento are not applicable to sequencing-based data, and the nuclear boundary information required for Wilcox and Bento was not available in this data.

**Fig. 3.**
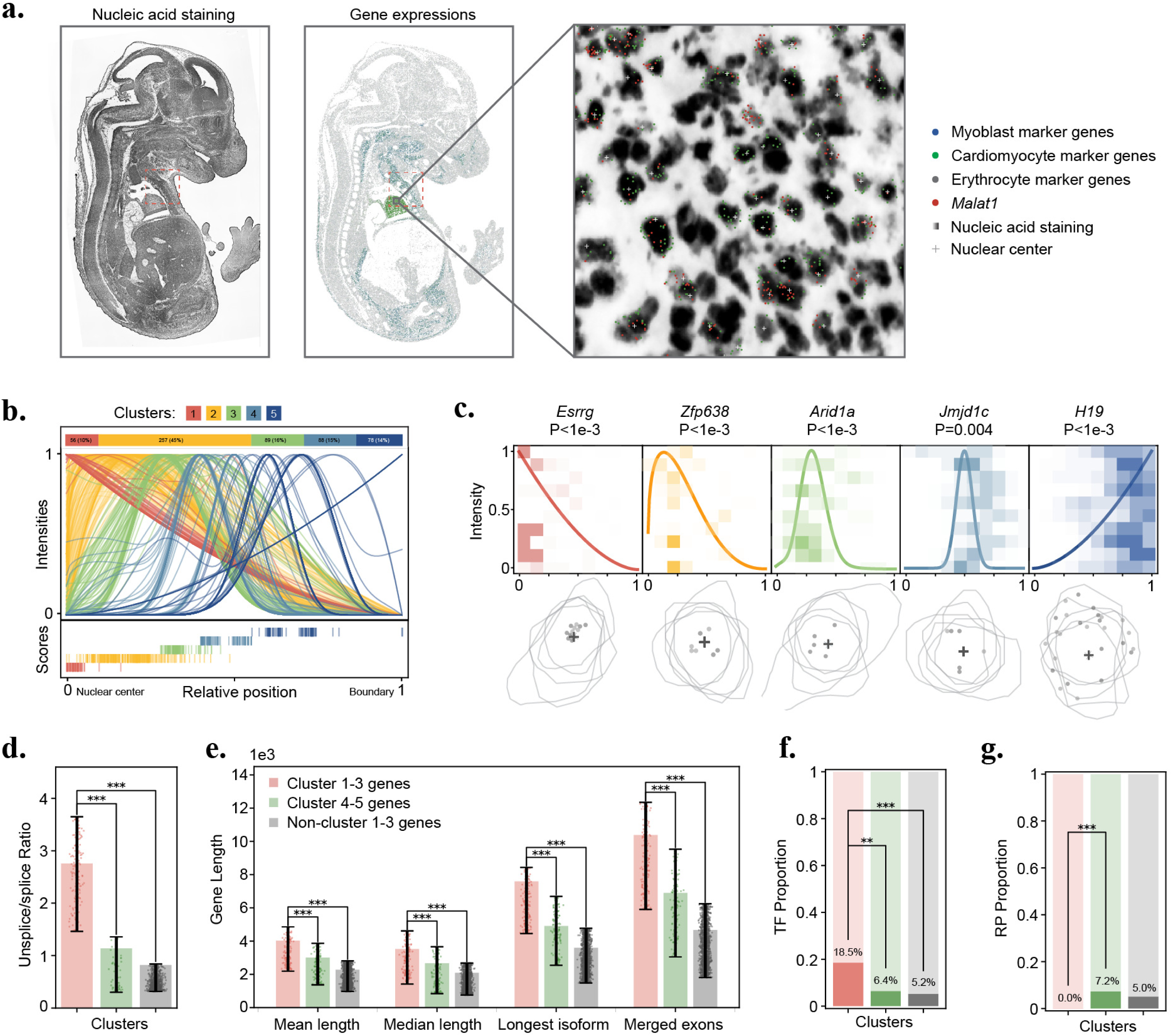
Stereo-seq mouse embryo data analysis. **a.** Data snapshot for tissue slice E1S3. Left panel displays the nucleic acid staining image of the slice. Middle panel displays expression from three gene sets (Erythrocyte marker genes, grey dots; Myoblast marker genes, blue dots; Cardiomyocyte marker genes, green dots; Tab. S11). Right panel zooms into a subregion and displays expression from three gene sets (Myoblast marker genes, blue dots; Cardiomyocyte marker genes, green dots; *Malat1*, red dots), nuclear centers (crosses) overlayed on the nucleic acid staining image. **b.** Estimated spatial expression pattern for genes in each of the five gene pattern clusters identified by ELLA. Upper panel shows the number and proportion of genes across five pattern clusters. Middle panel displays the estimated expression intensity for genes across clusters. Lower panel displays the estimated pattern score for genes across clusters. **c.** Example genes and cells for the five pattern clusters. One example gene is shown for each pattern cluster. Upper panel shows the gene name, ELLA P value, and the estimated expression intensities overlayed on the density heat map. Each row of the density heat map visualizes the number of counts standardized by area in 10 randomly selected cells across relative positions with intervals of 0.1. Lower panel displays the expressions of the corresponding genes within five selected cells, overlayed with cell boundaries and aligned nuclear centers (crosses). **d.** Bar plot shows the average unspliced/spliced expression ratio across genes in pattern clusters 1-3 (red), 4-5 (green), and non-cluster 1-3 genes (i.e. clusters 4-5 plus the nonsignificant genes; grey). Genes enriched close to nuclear center (cluster 1-3) tend to exhibit higher unspliced/spliced expression ratios. **e.** Bar plot displays average gene length, measured by four metrics (x-axis), across genes in pattern clusters 1-3, 4-5, and non-cluster 1-3 genes. Genes enriched close to nuclear center (cluster 1-3) tend to exhibit longer gene lengths. **f.** Bar plot displays the proportions of transcription factors (TFs) for genes in pattern clusters 1-3, 4-5, and non-cluster 1-3 genes. Genes enriched close to nuclear center (cluster 1-3) contain a higher proportion of TFs. **g.** Bar plot displays the proportions of ribosomal protein (RP) genes for genes in pattern clusters 1-3, 4-5, and non-cluster 1-3 genes. Cytoplasmic enriched genes (clusters 4-5) contain a higher proportion of RP genes. Statistical significance for pair-wise comparisons (*: <0.05; **: <0.01; ***: <0.001) is based on Mann-Whitney U test (**d**-**e**) or Fisher’s exact test (**f**-**g**). The error bars represent the 25th and 75th percentiles, and data points beyond this range are not included (**d**-**e**).

At an FDR of 5%, ELLA identified 264 and 304 genes to be spatially variable within myoblasts and cardiomyocytes, respectively (Fig. S39). 89 genes were detected in both cell types including 12 transcription factors. Based on their subcellular spatial expression patterns, we clustered the detected genes into five distinct clusters (Methods, Fig. 3b): 56 genes (10%) display a nuclear expression pattern (clusters 1), 346 genes (61%) display one of the two nuclear edge expression patterns (cluster 2-3), and 166 genes (29%) display one of the two cytoplasmic expression patterns (cluster 4-5). Example cells from the five clusters are shown in Fig. 3c.

The genes detected by ELLA again allow us to comprehensively investigate the properties of genes that display distinct enrichment patterns within cells. For nuclear genes (clusters 1-3) with subcellular enrichment near the nuclear center, we found them to have significantly higher unsplice/splice ratio (clusters 1-3 vs clusters 4-5, fold enrichment=2.42, P value=1e-24; clusters 1-3 vs all the remaining genes, fold enrichment=3.36, P value=5e-96; Fig. 3d), which is also negatively correlated with the expression pattern score (Pearson correlation=-0.464, P value=1e-152). In addition, genes in clusters 1-3 are enriched with transcription factors (proportion=14.43%) as compared to the other clusters (clusters 4-5, proportion=5.42%, Fisher’s exact test P value=2e-3) or the remaining genes (proportion=5.20%, P value=2e-10; Fig. 3f). For the genes with subcellular enrichment in the cytoplasm (clusters 4-5), we found them to contain a significantly higher proportion of ribosomal protein (RP) genes (clusters 4-5, 7.23% vs clusters 1-3, 0%, Fisher’s exact test P value=3e-7; cluster 3-4 vs all the remaining genes, 4.38%, P value=0.086; Fig. 3g, Methods), supporting their localized synthesis. Importantly, nuclear genes (clusters 1-3) also tend to have longer gene lengths compared to genes in the other clusters or the remaining genes, in terms of the average isoform length (P value=2e-12 and 2e-60), the median isoform length (P value=2e-8 and 4e-34), the longest isoform length (P value=1e-16 and 5e-84), and the total length across exons (P value=1e-14 and 1e-87; Fig. 3e). Finally, in terms of 3’UTR length (Supplementary Notes 1, Fig. S40), 19 genes display significant variation across five expression pattern clusters (Fig. S41), 21 genes display significant correlation with expression pattern strength (Fig. S42), and 18 genes display significant correlation with expression pattern score (Fig. S43).

We further investigated the shared and distinct features of the genes detected by ELLA in both myoblasts and cardiomyocytes to reveal additional biological insights (Fig. S44). Both cell types exhibit a similar proportion of genes across the five expression pattern clusters, with common genes displaying similar estimated expression intensities (Fig. S45-46). Among the detected genes, 12 transcription factors are detected in both cell types (16 unique in myoblasts and 27 unique in cardiomyocytes; Fig. S47a). These transcription factors are enriched in GO gene sets related to regulation of transcription, development, and various regulatory categories (Fig. S47b-d). In addition, among the detected genes, 7 long noncoding genes are detected in both cell types (5 unique in myoblasts and 0 unique in cardiomyocytes; Fig. S48), including five (*Xist*, *Meg3*, *Kcnq1ot1*, *Malat1*, and *Airn*) localized near the nuclear center (cluster 1-3). One mitochondrial gene (*mt-Nd1*) is detected in both cell types (0 unique in myoblasts and one unique in cardiomyocytes; Fig. S49a) and it displays a cytoplasmic localized pattern (cluster 4; Fig. S49b). Two cardiomyocyte cell type marker genes (*Acta1* and *Myh3*) are detected as localized within cytoplasmic region (clusters 4) in the cardiomyocyte cell type (Fig. S50).

### SeqFish+ mouse embryonic fibroblast data

Next, we analyzed the NIH/3T3 mouse embryonic fibroblast cell line data generated by seqFISH+ [10], which contains 2,747 genes measured on 171 embryonic fibroblast cells (Fig. 4a, S51). We were unable to apply SPRAWL due to its heavy computational burden but were able to apply Bento as this data contains nucleus segmentation information.

**Fig. 4.**
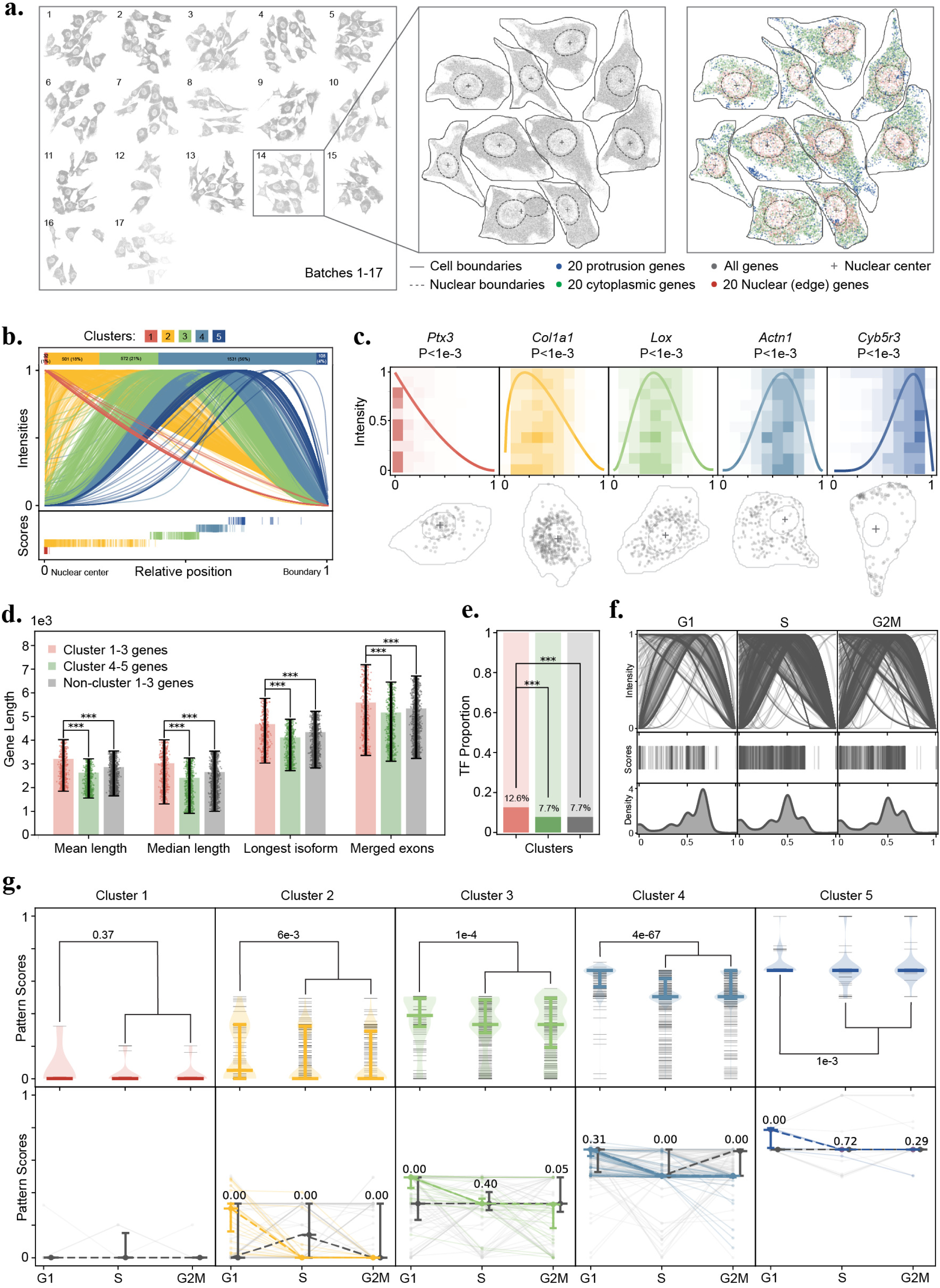
SeqFISH+ mouse embryonic fibroblast data analysis. **a.** Data snapshot for the embryonic fibroblast cell line. Left panel displays all transcripts (grey dots) measured across 17 batches. Middle panel zooms into batch 14 and displays all transcripts (grey dots), nuclear centers (crosses), along with nuclear segmentation boundaries (grey dashed line) and cell segmentation boundaries (grey solid line). Right panel displays batch 14 with expression from three gene sets (nuclear or nuclear edge genes, red dots; cytoplasmic genes, green dots; protrusion genes, blue dots; Tab. S14) and nuclear centers (crosses) overlayed with nuclear segmentation boundaries (grey dashed line) and cell segmentation boundaries (grey solid line). **b.** Estimated spatial expression pattern for genes in each of the five gene pattern clusters identified by ELLA. Upper panel shows the number and proportion of genes across five pattern clusters. Middle panel displays the estimated expression intensities for genes across clusters. Lower panel displays the estimated pattern score for genes across clusters. **c.** Example genes and cells for the five pattern clusters. One example gene is shown for each pattern cluster. Upper panel shows the gene name, ELLA P value, and the estimated expression intensity overlayed on the density heat map. Each row of the density heat map visualizes the number of counts standardized by area in 10 randomly selected cells across relative positions with intervals of 0.1. Lower panel displays the expressions of the corresponding genes within one selected cell, overlayed with cell boundary and nuclear center (cross). **d.** Bar plot displays average gene length, measured by four metrics (x-axis), across genes in pattern clusters 1-3 (red), 4-5 (green), and non-cluster 1-3 genes (i.e. clusters 4-5 plus the nonsignificant genes; grey). Genes enriched close to nuclear center (clusters 1-3) tend to exhibit longer gene lengths. **e.** Bar plot displays the proportions of transcription factors (TFs) for genes in pattern clusters 1-3, 4-5, and non-cluster 1-3 genes. Genes enriched close to nuclear center (clusters 1-3) contain a higher proportion of TFs. **f.** Estimated spatial expression pattern of genes across cell cycle phases (from left to right: G1, S, and G2M). Upper panels display the estimated expression intensities of genes across cell cycle phases. Middle panels display the estimated pattern score of genes across cell cycle phases. Lower Panel display the distribution of the estimated pattern score of genes across cell cycle phases. **g.** Upper panel displays Violin plots of the estimated pattern scores across cell cycle phases for genes in different pattern clusters. Genes significant in the G1 phase are less likely to be enriched close to nuclear center and display larger pattern scores compared to the genes in the S and G2M phases based on one side Mann-Whitney U test. Lower panel displays line plots of patterns score trajectories with respect to cell cycle phases for genes commonly detected in all cell cycle phases across pattern clusters. A subset of genes in clusters 2-5 display decreasing pattern scores through G1, S, and G2M phases (colored lines) while all six cluster 1 genes retain the nuclear pattern across all cell cycle phases (grey lines). Statistical significance for pair-wise comparisons (*: <0.05; **: <0.01; ***: <0.001) is based on Mann-Whitney U test (**d**) or Fisher’s exact test (**e**). The error bars represent the 25th and 75th percentiles, and data points beyond this range are not included (**d**).

At an FDR of 5%, ELLA identified 2,744 genes to display subcellular spatial expression patterns, with 244 being transcription factors. The subcellular expression patterns of the detected genes can be clustered into five distinct clusters (Fig. 4b, Methods): 32 genes (1%) display a nuclear expression pattern (cluster 1), 1,073 genes (39%) display one of the two nuclear edge expression patterns (clusters 2-3), and 1,639 genes (60%) display one of the two cytoplasmic expression patterns (clusters 4-5). The identified genes included 57 out of 60 genes with subcellular localization patterns detected through an *ad hoc* procedure in the seqFISH+ original study. The localization categorization of the 57 genes closely aligns with the pattern reported in the original study but with finer details: for example, 20 genes detected as enriched generally in the nuclear and perinuclear regions in the original study were clustered here as either cluster 2 (8 genes) or cluster 3 (12 genes) genes (Fig. S52). Example cells from the five clusters are shown in Fig. 4c.

Because Bento is only applicable to individual cells, we randomly selected 20 cells (Fig. S53) and applied both ELLA and Bento to analyze one cell at a time on 356-1,213 (mean=808) genes with more than 10 counts. Across cells, Bento classified 38.2% genes to one of the four compartmental patterns, 21.5% genes to a pattern called “none”, and the remaining 40.39% genes to either none of these five patterns or multiple patterns (Fig. S54a). Certainly, Bento is unable to produce P values nor quantifications of statistical significance for any of the genes. ELLA was able to allocate all genes to five identified patterns, with 13.4% genes achieving statistical significance (5% FDR; Fig. S54b-c). For genes detected by ELLA and classified by Bento to patterns other than none, their expression pattern classifications are largely consistent with each other, although ELLA offers more detailed results (Fig. S55). For example, 92.6% of the “nuclear” patterned genes detected by Bento were also identified as nuclear genes by ELLA, and these genes were classified by ELLA into two separate clusters (66.0% genes in cluster 1 with nuclear pattern and 26.5% genes in cluster 2 with nuclear edge pattern).

The genes detected by ELLA allow us to comprehensively investigate the properties of genes that display distinct enrichment patterns within cells. For genes with subcellular enrichment near the nuclear center (clusters 1-3), we found them to have significantly longer gene lengths compared to genes in the other clusters (clusters 4-5) or the remaining genes, in terms of the average isoform length (P value=8e-19 and 1e-18), the median isoform length (P value=6e-14 and 7e-14), the longest isoform length (P value=6e-11 and 7e-11), and the total length across exons (P value=1e-4 and 1e-4; Fig. 4d). These four types of gene lengths are also significantly negatively correlated with the ELLA pattern scores (Pearson correlation ranges from −0.17 to −0.07; P values range from 1e-19 to 2e-4). Genes with enrichment near the nuclear center (clusters 1-3) are also enriched with transcription factors (proportion=12.56%) as compared to the other clusters (clusters 1 and 4-5, proportion=7.73%, P value=3e-4) or the remaining genes (proportion=7.72%, P value=2e-4; Fig. 4e).

ELLA also provides a unique opportunity for us to explore whether cell cycle may influence the subcellular spatial localization of gene expression, as the data is collected from cultured cells that undergo continuous cell division. To do so, we first clustered fibroblast cells into three distinct cell-cycle phases, including G1 (n=36, 21%), S (n=83, 49%), and G2M (n=52, 30%). We then applied ELLA to analyze each cell phase separately and detected 728, 2,368, and 1,726 genes with subcellular spatial expression patterns, respectively (Fig. 4f). We found that genes significant in the G1 phase are less likely to be enriched close to the nuclear center and display larger pattern scores compared to the genes in the S and G2M phases, regardless which cluster the genes belong to (pattern score fold enrichment in G1 vs S and G2M=2.33, 1.31, 1.12, 1.16, and 1.02, for the five clusters, respectively; one side Mann-Whitney U test P value=0.37, 0.01, 1e-4, 4e-67, 1e-3; Fig. 4g), suggesting that DNA replication during the S phase enhances nuclear enrichment in S and G2M phases. Among the detected genes, 723 are shared across three cell cycles, including 6 (1%), 105 (15%), 123 (17%), 442 (61%), and 47 (7%) genes for each of the five clusters, respectively. Among the shared genes, a subset of genes in clusters 2-5 display decreasing pattern scores through G1, S, and G2M phases, corresponding to increasing enrichment towards the nucleus. In contrast, all six cluster 1 genes retain the nuclear pattern across all cell cycle phases (Fig. 4g, Methods), suggesting that some genes are capable of retaining their nuclear enrichment throughout the cell cycle phases [21].

### MERFISH mouse brain data

Lastly, we analyzed the adult mouse brain data generated by MERFISH [22] (Fig. 5a, S56). We focused on four major cell types residing in midbrain: excitatory neurons (EX, *n*=577), inhibitory neurons (IN, *n*=525), astrocytes (Astr, *n*=480), and oligodendrocytes (Olig, *n*=948) with 557-878 genes per cell type (Fig. S57). Besides ELLA, we were able to also apply SPRAWL to the data, but unable to apply Wilcox and Bento as the nuclear boundary information required for these two methods were not available in this data.

**Fig. 5.**
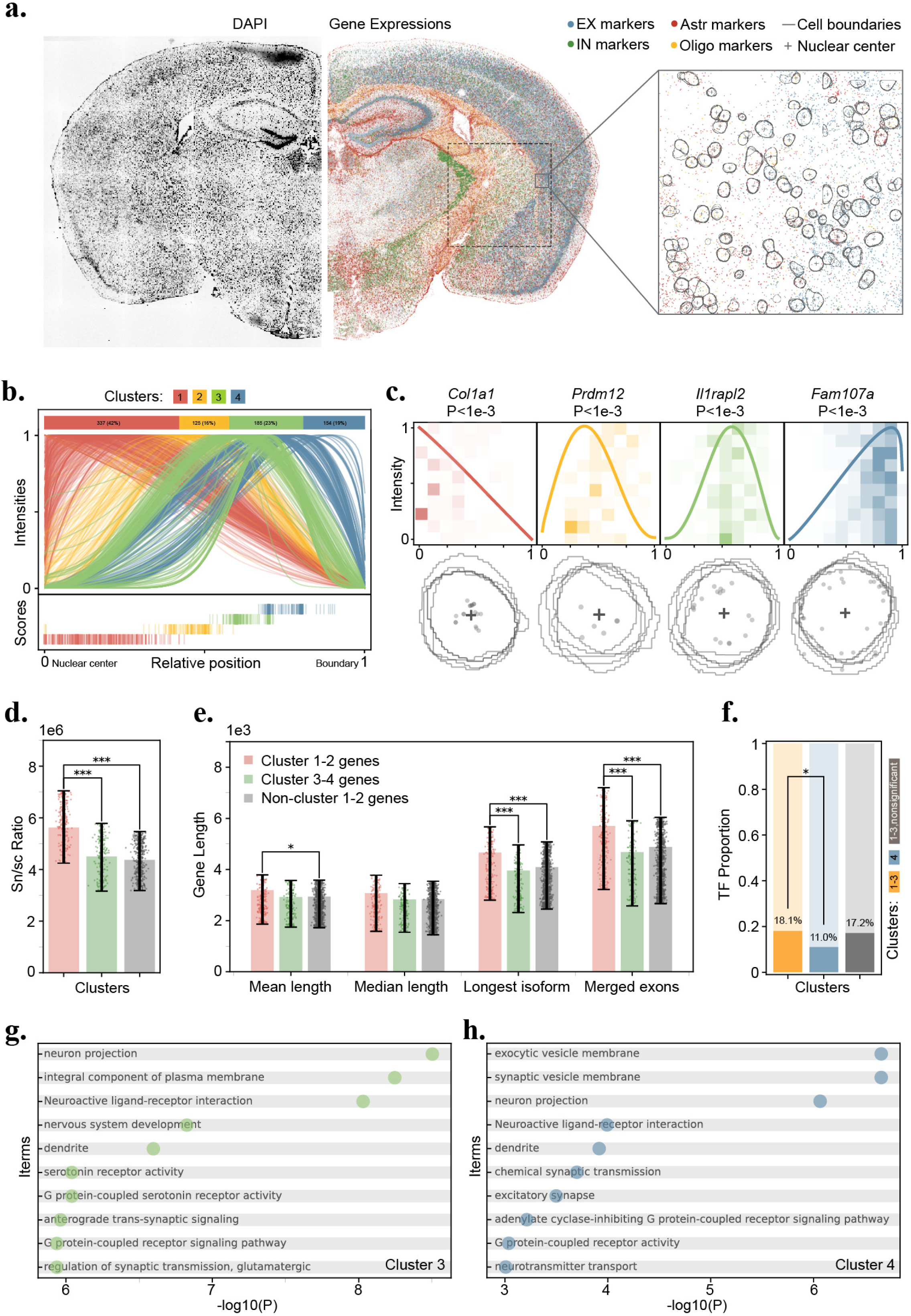
MERFISH mouse brain data analysis. **a.** Data snapshot for tissue slice hemi-brain region. Left panel shows the DAPI staining image. Middle panel displays expression for four gene sets (EX marker genes, blue dots; IN marker genes, green dots; Astr marker genes, red dots; Olig marker genes, orange dots; Tab. S12). Right panel zooms into a subregion and displays expression for the four gene sets along with cell centroids (crosses) overlayed with cell segmentation boundaries across five z stacks. **b.** Estimated spatial expression pattern for genes in each of the four gene pattern clusters identified by ELLA. Upper panel shows the number and proportion of genes across four pattern clusters. Middle panel displays the estimated expression intensity for genes across clusters. Lower panel displays the estimated pattern score for genes across clusters. **c.** Example genes and cells for the four pattern clusters. One example gene is shown for each pattern cluster. Upper panel shows the gene name, ELLA P value, and the estimated expression intensities overlayed on the density heat map. Each row of the density heat map visualizes the number of counts standardized by area in 10 randomly selected cells across relative positions with intervals of 0.1. Lower panel displays the expression of the corresponding genes within five selected cells, overlayed with cell boundaries and aligned nuclear centers (crosses). **d.** Bar plot shows the average sn/sc RNA ratio in the form of snRNA expression level normalized by scRNA expression level across genes in pattern clusters 1-2 (red), 3-4 (green), and non-cluster 1-2 genes (i.e. clusters 3-4 plus the nonsignificant genes; grey). Genes enriched close to nuclear center (clusters 1-2) tend to exhibit higher relative snRNA expression levels. **e.** Bar plot displays average gene length, measured by four metrics (x-axis), across genes in pattern clusters 1-2, 3-4, and non-cluster 1-2 genes. Genes enriched close to nuclear center (clusters 1-2) tend to exhibit longer gene lengths. **f.** Bar plot displays the proportions of transcription factors (TFs) for genes in pattern clusters 1-3 (orange), 4 (blue), and non-cluster 4 genes (i.e. clusters 1-3 plus the nonsignificant genes; grey). Genes enriched close to cell boundary (cluster 4) contain a lower proportion of TFs. **g** and **h**. Stem plots show the -log10 P values of the top 10 enriched gene sets in GSEA analysis for genes in pattern cluster 3 and 4 respectively. Gene sets enriched with cluster 3 or 4 genes are related to dendrites and synaptic transmission and signaling. Statistical significance for pair-wise comparisons (*: <0.05; **: <0.01; ***: <0.001) is based on Mann-Whitney U test (**d**-**e**) or Fisher’s exact test (**f**). The error bars represent the 25th and 75th percentiles, and data points beyond this range are not included (**d**-**e**).

At an FDR of 5%, ELLA identified 235, 250, 169, and 147 (total=801, total distinct=502) genes that display subcellular spatial expression patterns in EX, IN, Astr, and Olig cells, respectively (Fig. S57). 227 of these genes, including 36 transcription factors, were detected in two or more cell types. The subcellular spatial expression patterns of the detected genes can be clustered into four distinct pattern clusters (Fig. 5b, Method): 337 genes (42%) display a nuclear expression pattern (cluster 1), 125 (16%) genes display a nuclear edge expression pattern (cluster 2), and 339 genes (42%) display one of the two cytoplasmic expression patterns (clusters 3-4). Example cells from the four clusters are shown in Fig. 5c. Compared to the number of genes (801) detected by ELLA, the peripheral, central, radial, and punctate metrics of SPRAWL detected 572, 305, 138, and 238 genes, respectively, with 434 distinct genes in total, the majority of which (345; 79.49%) are overlapped with ELLA (Fig. S58). Note that SPRAWL radial and punctate metrics excluded 57.8% of the unqualified gene-cell pairs that have less than two counts of a gene in a cell, which likely leads to their lower power as well as their failure in producing P values for a small percentage of genes across cell types (2.3%, 272 genes).

The genes detected by ELLA allow us to comprehensively investigate the properties of genes that display distinct enrichment patterns within cells. For genes with subcellular enrichment near the nuclear center (clusters 1-2), we found them to have significantly higher snRNA expression in the same cell types from a separate study (clusters 1-2 vs clusters 3-4, fold enrichment=1.25, P value = 1e-15; cluster 1-2 vs all remaining genes fold enrichment=1.29, P value=2e-31; Fig. 5d; [30]). We also found them to have significantly longer gene lengths compared to genes in the other clusters or the remaining genes, in terms of the average isoform length (P value=0.083 and 0.021), the longest isoform length (P value=6e-5 and 1e-6), and the total length across exons (P value=2e-6 and 2e-9; Fig. 5e). In addition, the cluster 4 genes contain a lower proportion of transcription factors (proportion=11.04%) as compared to the other clusters (cluster 1-3, proportion=18.08%, P value=0.041) or the remaining genes (proportion=17.16%, P value=0.598; Fig. 5f). Gene sets enriched with the cluster 1-2 genes are related to various functions including transcription regulation (Fig. S59), while gene sets enriched with cluster 3-4 genes are particularly related to dendrites and synaptic transmission and signaling (Fig. 5g-h).

We investigated the shared and distinct features of the genes detected by ELLA in the two neuronal cell types, excitatory and inhibitory neurons to reveal additional biological insights. Excitatory neurons contain a slightly higher proportion of nuclear localized genes (cluster 1) and a lower proportion of cell membrane localized genes (cluster 4) compared to inhibitory neurons (Fig. S60). A fraction of the detected genes (Jaccard index=31.8%) are shared between the two neuronal types, with 38, 3, 8, and 11 shared genes detected across clusters 1-4 and with similar estimated expression patterns (Fig. S61-62). In addition, the majority of the detected transcription factors (126) are shared between the two neuronal types, while 20 are uniquely detected in excitatory neurons and 9 uniquely detected in inhibitory neurons (Fig. S63). The 126 shared transcription factors are enriched in 112 gene sets related to various transcription regulations and neuron differentiation (Fig. S64-65). Four out of eight long noncoding genes are detected in both cell types (Fig. S66a). Three out of four of the common long noncoding genes are localized in the nucleus in both cell types (cluster 1: *A830036E02Rik*, *B020031H02Rik*, and *Dlx6os1*), and one gene (*Rmst*) is localized in nuclear edge in the excitatory neurons (cluster 2; Fig. S66b). Most cell type marker genes detected by ELLA belong to clusters 3-4 with cytoplasmic or membrane localization patterns except for one gene (*Cux2* being detected as cluster 2, nuclear edge localized, in excitatory neurons; Fig. S67).

## Discussion

We have presented ELLA, a statistical method for modeling and detecting spatially variable genes within cells that display various subcellular spatial expression patterns in high-resolution spatial transcriptomic studies. ELLA models the spatial distribution of gene expression measurements along the cellular radius using a nonhomogeneous Poisson process, leverages multiple kernel functions to detect a variety of subcellular spatial expression patterns, and is capable of analyzing a large number of genes and cells. We have illustrated the benefits of ELLA through simulations and real data applications.

We have primarily focused on utilizing ELLA to capture the spatial variation of gene expression along the cellular radius within cells, which is inherently one-dimensional and rotation invariant. Detecting rotation-invariant and radial symmetric patterns enables information sharing across multiple cells, thereby enhancing statistical power. In addition, rotation-invariant patterns facilitate results interpretation, as the detected genes can be naturally categorized into cellular compartments, including nucleus, nucleus membrane, and cellular membrane. The framework of ELLA, however, is general and can be extended to two- or three-dimensional cellular space, enabling modeling of 2D cellular space with kernels defined on a unit circle or 3D cellular space with kernels defined on a unit ball. Use of different kernels on higher dimensional space may further enhance the power of ELLA. For example, radial kernel functions may be particularly effective in detecting genes with radial patterns in 2D cellular space -- a pattern that, although unlikely to be biological, the one-dimensional version of ELLA is ill equipped to detect, as shown in the simulations. Such extensions, however, necessitate careful consideration, as additional modeling features, such as rotation invariance, may need to be incorporated into the kernel structure to effectively utilize information from multiple cells.

ELLA leverages nuclear center and cellular boundary information extracted from the spatial transcriptomics data or its accompanying histology image data to register and segment cells through multiple pre-processing steps. These pre-processing steps can vary substantially across different spatial transcriptomics technologies. For example, the accompanying H&E and nucleic acid staining images in Seq-Scope and Stereo-seq need to be registered with the spatial transcriptomics data to obtain the cellular boundary information, while the DAPI images in imaging-based datasets have already aligned with the spatial transcriptomics data without the need for further registration. Similarly, the nucleus center in imaging-based datasets is determined as the geometric center of the nuclear segmentation, while in sequencing-based datasets is determined based on the enrichment of unspliced sequencing read counts. Importantly, ELLA provides accompanying scripts tailored to distinct spatial transcriptomics platforms to streamline these pre-processing steps. In addition to the nuclear center and cellular boundary information, additional data such as nuclear boundary information can also be integrated into ELLA as needed. In such cases, the registration step of ELLA can be extended to register cells based on the nuclear center, nuclear boundary, as well as cellular boundary. Furthermore, the modeling framework of ELLA can be extended to accommodate this additional information, as well as other subcellular compartmentalization information, such as annotations on nucleolus or ER membrane, as technologies continue to improve in the future. Investigating the effectiveness of ELLA in the context of additional feature information represents an important avenue for future research.

## Methods

### ELLA overview

#### Subcellular resolution spatial transcriptomics and data preprocessing

We consider a high-resolution spatial transcriptomics study that collects gene expression measurements at subcellular level. For sequencing-based techniques such as Seq-Scope [5] and Stereo-seq [8], the expression of a gene is measured on a set of pre-specified spatial locations and is represented as the number of read counts mapped to the mRNA transcripts of the gene. For imaging-based techniques such as seqFISH+ [10] and MERFISH [9], the expression of a gene is measured as the presence of a hybridization signal of its mRNA across spatial locations. Regardless of the techniques, we assume that *G* genes are measured on *S* spatial locations, where these locations have known two-dimensional *x* and *y* spatial coordinates that are recorded during the experiment. For a gene *g*, its raw expression measurement at each location is represented either as a count (for sequencing-based techniques) or as a binary label (for imaging-based techniques) depending on the spatial transcriptomic technique.

To facilitate joint modeling across cells, we create a unified cellular coordinate system to anchor diverse cell shapes and morphologies. To do so, for the high-resolution spatial transcriptomics data, we first follow standard data preprocessing procedures to segment the tissue into cells. We cluster these cells into different cell types based on marker gene expression. For each cell in turn, we obtain the center of its nucleus and assign the spatial coordinates to all expression measurement locations within the cell. For each measured location inside the cell, we calculate two distances: its distance to the nuclear center *d*_1_, and its distance to the cell boundary *d*_2_ in the opposite direction from the nuclear center (Fig. S68a). With these two distances, we further calculate the relative position of the measured location inside the cell as the ratio between the nuclear distance and the summation of the two distances *d*^’^_1_ = *d*_1_/(*d*_1_ + *d*_2_). The relative position ranges between 0 and 1 and allows us to create a unified coordinate system across cells, enabling the joint modeling of multiple cells regardless of their sizes and shapes (Fig. S68b). Importantly, we compute the cellular distances for each measured location efficiently using a binning-based numerical approximation approach. Specifically, we first divided each cell from the center of nucleus into 100 circular sectors of equal angle measure. In each sector *v*, we denote *r*_*v*_ as the maximum distance between the center of the nucleus and the cellular boundary in the sector using cell segmentation boundary or mask. For each expression measurement location within the sector, we obtain its distance from the center of the nucleus and normalize it by *r*_*v*_ to obtain its relative position. This binning-based approximation approach speeds up computation through eliminating the requirement of computing the distance of each measurement location to the cell boundary, facilitating parallel computation across cells and sectors.

#### ELLA model for detecting genes with subcellular spatial expression patterns

With the expression measurements and their relative positions within each cell, we aim to identify spatially variable genes that display subcellular spatial expression patterns along the cellular radius that points from the center of the nucleus towards the cellular boundary. The genes with subcellular spatial expression patterns are often localized in certain cellular compartments such as nucleus, cytoplasm, Golgi apparatus, or cell membrane and may display distinct enrichment associated with such compartmentalization. To identify those genes, we examine one gene at a time and jointly model its expression measurements within *n* cells that belong to a given cell type. For the *i*th cell (*i* = 1, …, *n*), we assume that the gene is measured on *m*_*i*_ spatial locations. For the *j*th measured location (*j* = 1, …, *m*_*i*_), we denote the measured gene expression value as *y*_*ij*_, which is either a count or a binary value. We denote the relative position of *j*th measured location as *r*_*ij*_ ∈ [0,1], where 0 corresponds to the center of the nucleus and 1 corresponds to the cellular boundary.

We model the subcellular spatial localization of gene expression within each cell using a one-dimensional nonhomogeneous Poisson process (NHPP) model. Specifically, we assume that the gene expression counts summed across all relative positions within a given interval [*a*, *b*] ⊂ [0, 1] on the cellular radius follow a Poisson distribution, with the rate parameter being the integration of an underlying NHPP density function in the interval [*a*, *b*], where the NHPP density function may vary with respect to the relative position within the cell. Mathematically, the model is expressed as:

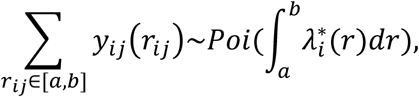

where *Poi* denotes a Poisson distribution and λ^∗^(*r*) is the unknown NHPP density function depending on the relative position *r*. We assume that the NHPP density function λ^∗^(*r*) is expressed as a product of three terms

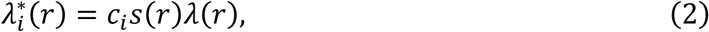

where *c*_*i*_, the total read depth for the *i*th cell, is used for normalization purpose and is calculated as the summation of the total read counts across all genes within the cell; *S*(*r*) = 2π*r* is another normalization term to capture the fact that the cellular area corresponds to a particular radius *r* is proportional to the size of the radius (that is, the area between *r* and *r* + Δ*r* is π(*r* + Δ*r*)^2^ − π*r*^2^ = 2π*r* + π(Δ*r*)^2^ → 2π*r* as Δ*r* → 0); and λ(*r*) is the key term of interest, the subcellular spatial expression intensity function that captures the subcellular spatial expression pattern along the cellular radius.

With the above NHPP model, we can write down the joint likelihood of the subcellular gene expression across *n* cells as:

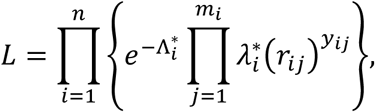

with *Λ^*^_i_ = ∫^1^_0_λ^*^_i_(r)dr*. Note that we have assumed that the subcellular spatial expression intensity function λ(*r*) is shared across cells, allowing us to borrow information across cells to enhance the detection of subcellular spatial expression patterns.

The intensity function λ(*r*) is key for modeling the subcellular spatial expression pattern of the given gene. In particular, if a gene does not display subcellular spatial expression pattern and is instead uniformly distributed within the cells, then λ(*r*) is expected to be a constant that is invariant to the relative position *r*. In contrast, if a gene displays subcellular spatial expression pattern, then λ(*r*) is expected to vary as a function of the relative position *r*.

Therefore, in the above NHPP model, identifying genes that display subcellular spatial expression pattern within cells is equivalent to testing whether λ(*r*) is a constant or not. The statistical power of such hypothesis test will inevitably vary depending on how the specified expression intensity function λ(*r*) matches the true underlying subcellular spatial expression pattern displayed by the gene of focus. For example, an intensity function enriched near zero will be particularly useful for detecting subcellular expression patterns that are also enriched in the nuclear, while an intensity function enriched near one will be particularly useful for detecting subcellular expression patterns that are also enriched near the cellular membrane. However, the true underlying subcellular spatial pattern for any gene is unfortunately unknown and may vary across genes. To ensure robust identification of subcellular spatial expression genes across various spatial patterns, we consider using a total of *k*=22 different kernel functions φ_1_(*r*), …, φ_*k*_(*r*) inside the intensity function λ(*r*) to capture a wide variety of possible subcellular spatial expression patterns (Fig. S68c). In particular, each function is a Beta probability density function defined on the interval [0,1], characterized by one of the 22 sets of shape parameters (Tab. S8) with a mode centering on 0, 0.1, 0.2, …, or 1. Note that, while we use these 22 kernel functions as default kernels in the present study, our method and software implementation can easily incorporate various numbers or types of intensity kernels as desired by the user.

For each kernel *l* = 1, …, *k* in turn, we model the intensity function in the form of λ(*r*) = α_*l*_ + β_*l*_φ_*l*_(*r*), where α_*l*_ is the nonnegative intercept parameter and β_*l*_ is the nonnegative scaling parameter for the *l* th kernel function. With the functional form of λ(*r*), we can test the null hypothesis *H*_0_: β_*l*_ = 0, that λ(*r*) is a constant. Rejecting the null hypothesis allows us to detect genes that display subcellular spatial expression patterns captured by the particular kernel. We perform inference and hypothesis test for each kernel in turn using a likelihood ratio test. In particular, we first maximize the log likelihood both under the null and under the alternative using the Adam optimizer in PyTorch [31]. Afterwards, we obtain the corresponding P value asymptotically based on an equal mixture of two chi-square distributions with degrees of freedom being zero and one [32]. Afterwards, we combine the *k* different P values calculated using different kernels into a single P value using the Cauchy combination rule [33, 34]. Specifically, we convert each of the *k* P values into a Cauchy statistic, aggregate the *k* Cauchy statistics through summation, and convert the summation back to a single P value based on the standard Cauchy distribution. The Cauchy rule takes advantage of the fact that a combination of Cauchy random variables also follows a Cauchy distribution regardless of whether these random variables are correlated or not. Therefore, the Cauchy combination rule allows us to effectively combine multiple potentially correlated P values into a single P value for every gene. Finally, we control FDR across genes using the Benjamini-Yekutieli procedure, which is effective for arbitrary dependency among test statistics. We used an FDR cutoff of 0.05 for declaring significance.

#### Estimation of the subcellular spatial expression pattern with ELLA

While the primary focus of ELLA is on hypothesis testing, it can also be used to estimate the subcellular spatial expression pattern for the detected genes. Specifically, for gene *g*, we can first obtain the *k* estimated intensity functions for each of the *k* kernel functions as

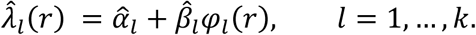

where α^_*l*_ and β^_*l*_ are the estimates for the corresponding parameters. Because each of the *k* estimated intensity functions captures a particular aspect of the overall subcellular spatial expression intensity function λ(*r*), we estimate λ(*r*) with a weighted combination of the estimated intensity functions in the form of

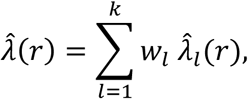

where *W*_*l*_ is the weight for the *l*th intensity function with ∑^*k*^ *W*_*l*_ = 1. The weights can be derived based on Bayesian model averaging [35]. In particular, we denote the model with *l* th kernel function as *M*_*l*_ and denote the data as *D*. The posterior distribution for λ(*r*) is in the form of: *P*(λ(*r*)|*D*) = ∑^*k*^ *P*(λ(*r*)|*M*_*l*_, *D*)*P*(*M*_*l*_|*D*), with the posterior mean estimate being λ^(*r*) = *E*[*P*(λ(*r*)|*D*)] = ∑^*k*^ *E*[*P*(λ(*r*)|*M*_*l*_, *D*)]*P*(*M*_*l*_|*D*) = ∑^*k*^ λ^_*l*_(*r*)*P*(*M*_*l*_|*D*). Therefore, the weights are in the form

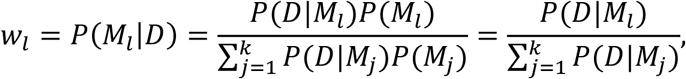

where the last equation holds due to the equal prior assumption on each model, with *P*(*M*_*j*_) = 1/*k* (*j* = 1, …, *k*). We approximate *P*(*D*|*M*_*l*_) with the maximized likelihood estimates to obtain the weights and subsequently λ^(*r*) (Supplementary Notes 2).

ELLA is implemented in python, with an underlying PyTorch Adam for efficient CPU or GPU computation. The software ELLA, together with all analysis code used in the present study, are freely available at https://xiangzhou.github.io/software/.

### Compared methods

We compared ELLA with three methods: (1) SPRAWL [20], (2) Bento [19], and (3) Wilcox. For both SPRAWL and Bento, we followed the tutorial on their corresponding GitHub pages and used the recommended default parameter settings.

SPRAWL takes RNA location information from subcellular multiplexed imaging datasets as inputs and does not explicitly require nuclear boundary or nuclear center information. SPRAWL examines one gene at a time and uses four localization metrics to capture four different types of subcellular spatial enrichment patterns that include peripheral, central, radial, and punctate. Specifically, the peripheral metric is used to identify peripheral/anti-peripheral patterns where the expression enrichment is either proximal or distal from the cell membrane. The central metric is used to identify central/anti-central patterns where the expression enrichment is either proximal or distal from the cell centroid. The radial metric is used to identify radial/anti-radial patterns where a gene is either aggregated or depleted in a sector of the cell. The punctate metric is used to identify punctate/anti-punctate patterns where a gene displays either self-colocalizing/self-aggregating or self-repulsion inside the cell. Because the radial and punctate metrics can only be computed for cells with no less than two expression counts, we had to filter out cells with less than two counts when analyzing a given gene for these two metrics. For each gene and each metric in turn, SPRAWL computes a score for every cell and averages them across cells in a particular cell type to obtain the per-cell-type score. SPRAWL then converted the per-cell-type score to a P value based on a standard normal distribution and used the Benjamini Hochberg (BH) procedure for FDR control. We used an FDR threshold of 0.05 to obtain significant genes.

Bento takes RNA location information from subcellular multiplexed imaging datasets as inputs and requires nuclear and cell boundaries as additional information. For each gene-cell pair in turn, Bento computes 13 spatial summary statistics and uses its RNAforest function, which consists of five independent pretrained binary random forest classifiers, to produce five binary labels that classify gene expression pattern into one of the five patterns including nuclear, nuclear edge, cytoplasmic, cell edge, and none. For each gene in the cell, we obtained the classification probability *p*_*c*_ for each pattern *c* and used 1 − *p*_*c*_ to rank genes for the pattern, which allowed us to measure powers based on FDR in the simulations. However, due to its use of classification probability, it is not feasible to obtain FDR control in any real datasets with Bento.

Wilcox, a Wilcoxon rank sum test-based approach developed in the present study, detects genes that are differentially expressed between two subcellular regions: the nucleus and the cytoplasm. We focus on these two subcellular regions because we can extract the nuclear boundary and cell boundary in many spatial transcriptomics studies. To detect those genes, for each cell in turn, we first extracted the gene expression counts within the nucleus as well as the gene expression counts in the cytoplasm. We then normalized the two counts by the corresponding cellular areas for the two subcellular regions. Afterwards, we performed Wilcoxon rank sum test across cells to detect genes that are differentially expressed between the nucleus and the cytoplasm.

### Simulations

We performed comprehensive simulations based on imaging data to evaluate the performance of ELLA and compare it with other methods. We did not perform simulations based on sequencing data as neither SPRAWL nor Bento can be applied to analyze these data. For simulations, we first extracted the cell boundaries of the embryonic fibroblast cells from the seqFISH+ data, calculated the minimal and maximal radius of each cell, obtained a list of 90 reasonably shaped cells with the ratio of minimal and maximal radius ≥0.3, and extracted their nuclear centers and boundaries. We then sampled with replacement *n* cells from these cells. For each cell in turn, we applied the same binning strategy used in ELLA preprocess to divide the cell from the center of nucleus into 100 circular sectors of equal angle measure. In each sector *v*, we denote *r*_*v*_ as the maximum distance between the center of nucleus and the cellular boundary in the sector. We calculated the approximate area of the sector *v* as π*r*^2^/100. We also denote θ_*v,min*_ and θ_*v,max*_ as the minimal and maximum angle measurement of the sector, respectively. For the alternative simulations, we further divided each circular sector into 25 annulus sectors with equal distances.

With the above preparations, we simulated gene expression for 1,000 genes, where each gene is expressed as a binary count on *m* subcellular localizations in each cell as imaging data. In the null simulations, none of these genes display cellular spatial expression patterns. In the alternative simulations, 800 genes are null while 200 genes display different types of subcellular expression patterns. Specifically, in the null simulations, we first randomly sampled the number of measured locations inside each sector (*m*_*v*_). We set *m*_*v*_ to be proportional to the area of the sector using the function “np.random.choice” with the constraint ∑_*v*_ *m*_*v*_ = *m*. For each of the *m*_*v*_ locations in sector *v*, we obtained two independent random variables, *u*_1_ and *u*_2_, from a uniform distribution *U*(0,1), and converted them into the radius (*r*) and angle (θ) coordinates for the location, where *r* = *r*_*v*√_*u*_1_ and θ = θ_*v,min*_ + *u*_2_(θ_*v,max*_ − θ_*v,min*_). The radius and angle coordinates are further converted to the *x* and *y* coordinates in the form of *x* = *r* cos(θ) and *y* = *r* sin(θ).

In the alternative simulations, we simulated gene expression to exhibit subcellular expression patterns from three pattern categories: symmetric, radial, and punctate. For the symmetric pattern category, we considered eleven different expression patterns, including two patterns with nucleus enrichment, two patterns with nuclear edge enrichment, five patterns with cytoplasmic enrichment, and two patterns with membrane enrichment. For each pattern, we first randomly sampled the number of measured locations inside each sector (*m*_*v*_). We set *m*_*v*_ to be proportional to the area of the sector using the function “np.random.choice” with the constraint ∑_*v*_ *m*_*v*_ = *m*. We then constructed the expression intensity function λ^true^(*r*) in the form of λ^true^(*r*) = α + βφ(*r*), where φ(*r*) is set to be one of the eleven beta probability density functions described earlier (upper panel in Fig. S68c). Each beta probability density function is characterized by one of the eleven sets of shape parameters (Set 1 in Tab. S8), with a mode centering on 0, 0.1, 0.2, …, or 1. With λ^true^(*r*), we define the patten strength *S* as (maxλ^true^(*r*)-minλ^true^(*r*))/minλ^true^(*r*). We also compute λ^∗true^(*r*) = 2π*r*λ^true^(*r*) and further *p* = ∫^*rq,max*^_rq,min_ λ^∗true^(*r*)*dr* [36], which represents the probability of observing an expression measurement in the *q*-th annulus sector. Afterwards, we simulated the number of expression measurement locations in each annulus sector, *m*_*v*1_, …, *m*_*v*20_∼ Multinomial (*m*_*v*_, *p*_1_, …, *p*_20_), with the total number of measured locations in the sector being *m*_*v*_ = ∑_*q*_ *m*_*vq*_. We then applied the same strategy described in the above paragraph to simulate the *x* and *y* coordinates for each of the *m*_*vq*_ locations within each annular sector *q*.

In the symmetric pattern, we created different simulation settings by varying the number of cells (*n*), expression level (*m*), the subcellular expression patterns, and pattern strength (*S*). To do so, for each pattern, we first create a baseline simulation setting where we set the number of cells to be *n*=100, the expression level to be *m*=5, and in the case of alternative simulations, the pattern strength to be moderate (*S*=0.6). We then varied the cell number (*n*=10, 20, 50, 100, 200, 300 or 500), expression level (*m*=1, 2, 10, 20, 50, 100), and pattern strength (*S* ranges from 0.1 to 1.0 with increments of 0.1), one parameter at a time on top of the baseline settings for each of the 11 symmetric patterns to create 22simulations settings. The detailed parameters for each simulation setting are listed in Tab. S9. We performed 10 simulation replicates in each setting.

For the radial patterns, we first consider a radial-unif setting where gene expression is enriched in one sector of the cell with the expression counts within the sector being randomly distributed. For each cell and each gene in turn, we randomly selected a sector with a central angle π/2. We sampled the number of measuring locations in the sector, *m*_1_, from a binomial distribution Bin(*m*, 0.5). We also sampled the number of measuring locations in the complementary sector with a central angle of 3/2π, *m*_2_, to be *m* − *m*_1_. Afterwards, we randomly sampled the *x* and *y* coordinates for each measurement location in the same way as described in the null simulations. Therefore, the gene expression is enriched in one sector of the cell with a fold enrichment of 3.0. Next, we consider a radial-cyto setting where the gene expression is not only enriched within the sector but are also further enriched in the cytoplasm. To do so, on top of the radial-uniform setting, we used the intensity function described in the symmetric pattern #7 to simulate the x and y coordinates for the measurement locations in the selected circular sector that has a central angle of π/2. In addition, we randomly sampled the *x* and *y* coordinates in the complementary circular sector with a central angle of 3/2π for the measurement locations in the same way as described in the null simulations. Therefore, the average gene expression inside the sector is also 3.0 times higher than that in the remaining parts of the cell, while the expression within the sector is enriched in the cytoplasmic region due to symmetric pattern #7 with a fold enrichment of approximately 5.1.

For the punctate pattern, we consider a punctate-cyto setting where gene expression is enriched in a small subcellular disc in the cytoplasm. To do so, we set the radius coordinate for the center of the punctate disc to be 0.8 and randomly sampled the corresponding angle coordinate θ from a uniform distribution *U*(0, 2π). We then converted the radius and angle coordinates to the location coordinates (*x*_*c*_, *y*_*c*_). Afterwards, we set the radius of the punctate disc to be 1/10 of the average cell diameter, which consists of 30 pixels for seqFISH+ cells. We sampled the number of measurement locations within the punctate disc, *m*_1_, from a binomial distribution Bin(*m*, 0.2). We randomly sampled the *x* and *y* coordinates for the *m*_1_ locations inside the punctate disc as well as those for the remaining *m* − *m*_1_ locations in the entire cell including the punctate disc using the same strategy in the null simulations. The expression in the punctate disc is on average 5.03 times higher than that in the remaining parts of the cell. For radial and punctate patterns, we also performed 10 simulations replicates for each of the three settings.

In summary, we used the simulations introduced above to systematically evaluate and compare the performance of different methods in terms of type I error control and statistical power. Specifically, type I error control was assessed by generating QQ plots of the log-transformed p-values (log10) under the null hypothesis, which allowed us to examine how closely the observed distribution of P values matched the expected uniform distribution under the null. Power was assessed by calculating the proportion of genes detected exhibiting any subcellular expression pattern, calculated as the fraction of true positive genes detected at an FDR threshold of 0.05, across various simulation scenarios, including different pattern types (symmetric, radial, and punctate) and varying parameters such as the number of cells, expression levels, and pattern strength.

### Analyzed datasets

We examined four public high-resolution spatial transcriptomics datasets described below.

### Seq-Scope mouse liver data

Seq-Scope is a spatial barcoding technology with a spatial resolution comparable to an optical microscope. It is based on a solid-phase amplification of randomly barcoded single-molecule oligonucleotides using an Illumine sequencing platform. These RNA-capturing barcoded clusters represent the pixels of Seq-Scope and are ∼0.5-0.8 μm apart from each other with an average distance of 0.6 μm, capturing 848 UMI on average per 10 μm diameter bin.

We downloaded the mouse liver data from the Seq-Scope resources website [37]. The data contains 5.88± 4.22 (mean ±sd) number of genes per pixel, with a total of 32,976 genes measured across ∼2 × 10^7^ locations. The Seq-Scope mouse liver data contains 10 tiles sequenced on one MiSeq flow cell with each tile being a 1mm-wide circular imaging area. Among these 10 tiles, six of them are from a normal mouse fragmented frozen liver section and four of them are from an early-onset liver failure mouse model section (TD; [23]). The tiles cover liver portal-central tissue zonation and contain two main cell types: hepatocytes and non-parenchymal cells (NPC) such as macrophages, hepatic stellate cells, endothelial cells, and red blood cells. Our analyses focus on the hepatocytes which can be further divided into periportal (PP) and pericentral (PC) cells. The two Seq-Scope tissue sections (normal and TD) each comes with multiple H&E staining images, including high-resolution images (10X) covering a portion of the normal and TD tile areas and low-resolution images (4X) covering nearly all the normal and TD tile areas. We used the low resolution (4X) images to ensure high coverage of the tiles.

The Seq-Scope mouse liver data consists of two data modalities, namely the spatial transcriptomics data and the accompanying H&E staining images. For the spatial transcriptomics data, we obtained the unspliced and spliced gene expression counts on each measured location using STARsolo from the raw fastq files. For the H&E staining images, we concatenated all the images from the normal tissue section or the TD tissue section, segmented individual cells on the concatenated image using Cellpose ([38]; Fig. S18-23), and obtained cells that overlapped with the tile areas. On each tile, we plotted the unspliced expression reads to visualize cell nucleus and plotted the total UMI counts to visualize the tissue boundaries (Fig. S24-26). These nucleus and tissue boundary information were used to manually align each spatial transcriptomics tile to the concatenated normal or TD H&E images (Fig. S27-28). After modality alignment, we assigned each spatial location to a cell based on the aligned cell segmentation results (Fig. S29-30). For each cell in turn, we used numpy.argmax function in python to declare its nuclear center, which is defined to be the location within 200 units (∼2μm) from the cell boundary where the maximum of unspliced read counts density is observed. In each tile, we filtered out cells with a low-quality nuclear center where the unspliced read count density values at the nuclear center is below the 95% quantile value across locations or where the spliced read count density at the nuclear center is above the 95% quantile value across locations. In addition, we obtained cell type marker genes for each of the three cell types (PP, PC, and NPC; Tab. S10; [5]) and obtained the total counts of cell type marker genes for each cell. Note that the NPC cells, such as macrophages, hepatic stellate cells, endothelial, and red blood cells, are relatively rare across the tiles and are hard to segment due to their small sizes on the H&E-based images. Therefore, following the original Seq-Scope study, we removed NPC cells that are characterized by NPC marker gene counts above the 95% quantile across all cells. Afterwards, we normalized the PP and PC marker gene counts for the remaining cells first across genes to have zero mean and unit standard deviation and then across cells to have zero mean and unit standard deviation. We then summed the normalized PP and PC marker genes separately in each cell to obtain a PP score and a PC score per cell. We annotated a cell as a PP cell if its PP score is greater than the PC score and annotated a cell as a PC cell otherwise. Such annotations largely align with Seq-Scope’s original cell type annotations (Fig. S32). We removed cells with extreme sizes, including extremely large cells with x or y coordinate range (max-min) exceeding the 95% quantile value across cells within the cell type or extremely small cells with x or y coordinate range below the 5% quantile value. After quality control, we obtained 276 normal PP cells, 276 normal PC cells, 236 TD PP cells, and 82 TD PC cells. Genes expressed in more than 50 cells and with more than 3 counts in at least 5 cells were retained, leading to 497 to 1,349 genes per cell type.

### Stereo-seq mouse embryo data

Stereo-seq combined DNA nanoball (DNB)-patterned arrays and *in situ* RNA capture to enhance the spatial resolution of omics-sequencing. Standard DNB chips have spots with approximately 0.22 μm diameter and a center-to-center distance of 0.5 or 0.715 μm, providing up to 400 spots per 100 μ*m*^2^ for tissue RNA capture. Stereo-seq captured UMI counts range on average from 69 per 2 μm diameter bin (for bin3, 3×3 DNB) to 1,450 per 10 μm diameter bin (for bin 14, 14×14 DNB, equivalent to ∼one medium size cell).

We downloaded the raw sequencing data on slice E1S3 of the Stereo-seq mouse embryo data from CNGB Nucleotide Sequence Archive [39]. We downloaded the processed gene expression (bin1) data and the accompanying nucleic acid staining image from MOSTA [40]. Slice E1S3 is a profiled sagittal frozen tissue section with 10μm thickness from a C57BL/6 mouse embryo on day E16.5. It covers all major tissues and organs including Epidermis, Meninges, Cartilage, Jaw and tooth, Choroid plexus, Kidney, GI tract, Spinal cord, Muscle, Heart, Bone, Cartilage primordium, Brain, Adrenal gland, Connective tissue, Thymus, Blood vessel, Liver, Olfactory epithelium, Lung, Pancreas, and Mucosal epithelium. The nucleic acid staining image of the slice was stained using BM purple and was imaged using a Ti-7 Nikon Eclipse microscope. We considered 25 cell types along with cell type marker genes from the Stereo-seq study (Tab. S11). The 25 cell types include Cardiomyocyte, Chondrocyte, Choroid plexus, Dorsal midbrain neuron, Ganglion, Endothelial cell, Keratinocyte, Epithelial cell, Erythrocyte, Facial fibroblast, Fibroblast, Forebrain neuron, Forebrain radial glia, Hepatocyte, Immune cell, Limb fibroblast, Macrophage, Meninges cell, Mid-/hindbrain and spinal cord neuron, Myoblast, Olfactory epithelial cell, Radial glia, Smooth muscle cell, Spinal cord neuron, and Diencephalon neuron. We processed the Stereo-seq data in the same way as we did for the Seq-Scope data except for the modality alignment step which is omitted here as the Stereo-seq slice was accompanied by nucleic acid staining that has already been aligned with the slices (Fig. S36). The processed data contains cell label of each location, cell center, cell boundary, cell type, and read depth of each cell (Fig. S37). We annotated cell types based on 75 cell type marker genes provided by the original study, resulting in an average of 3,689 cells (median=3,968, min=782, max=5,314) per cell type (Fig. S38). We focused on a cardiothoracic region on slice E1S3 (Fig. 3a) and two major cell types: precursor muscle cells, or myoblasts, and mature muscle cells, or cardiomyocytes. Similar quality control steps were conducted as described in the Seq-Scope data preprocessing. We retained genes expressed in more than 30 cells.

### SeqFISH+ mouse fibroblast data

SeqFish+ performs super-resolution imaging and multiplexing of 10,000 genes in a single cell using sequential hybridizations and imaging with a standard confocal microscope. We obtained the seqFISH+ NIH/3T3 fibroblast data preprocessed by Bento from [41]. The raw seqFISH+ data consists of two modalities: the spatial transcriptomics measurements and an accompanying DAPI staining image. The spatial transcriptomics modality of the data contains 3,726 genes with at least 10 counts expressed in at least one cell and 179 cells with nuclear segmentation results, with a resolution of 103nm. The downloaded seqFISH+ data comes with cell segmentation boundaries and nuclear segmentation boundaries, each represented by a set of points densely scattered along the boundaries. With the nucleus segmentation information, we computed the nuclear center of each cell as the k-means center of all nucleus boundary points. We computed the average nuclear radius of each cell by averaging the distance of all nuclear boundary points to the nuclear center. We computed the average cell radius of each cell by averaging the distance of all cell boundary points to the nuclear center. Afterwards, we computed the nucleus-cell ratio of each cell by dividing the average nuclear radius with the average cell radius. We excluded eight cells that have a nuclear-cell ratio beyond two standard deviations from the mean (Fig. S51a). We focused on the remaining 171 cells for analysis. These cells have an average nuclear-cell ratio of 0.46 (Fig. S51b). We retained genes expressed in more than 50 cells and with more than 3 counts in at least 5 cells, resulting in 2,747 genes for analysis.

### MERFISH adult mouse brain data

The mouse brain MERFISH dataset contains over 200 adult mouse brain slices from 4 mice and covers a panel of ∼1,100 selected genes with around 8 million cells. The dataset consists of two data modalities, namely the spatial transcriptomics data and the accompanying DAPI and polyA staining images. We focused on one coronal slice of mouse 2 from the 220501_wb3_co2_15_5z18R_merfish5 experiment and obtained the preprocessed data from [42]. The obtained data was measured on a coronal tissue slice with 10 μm thickness and contains five 1.5-μm-thick optical z-stacks, with 1,147 genes measured on ∼100,000 cells. The data also includes cell segmentation information in the form of sets of points densely scattered along the boundaries for each z-stack (0-4), along with cell centroid information shared across z-stacks (Fig. S56). For each measured transcript, we calculated its relative position to nuclear center based on the cell segmentation on the z-stack that it belongs to as well as the shared cell centroid. We exclude cells whose centroid is outside or too close to (< 0.5 μm) its segmentation boundaries on the baseline stack (z=0). In addition, we measured the variability of cell segmentation boundaries on each non-baseline stack (z>0) versus that on the baseline stack (z=0) by KL divergence. We excluded cells whose cell segmentation boundaries are highly variable across z-stacks based on a KL divergence threshold of 0.5. We obtained cell type marker genes (Tab. S12) from the Stereo-seq study for four cell types that include excitatory neurons (EX), inhibitory neurons (IN), astrocytes (Astr), and oligodendrites (Olig). We then carried out the same cell typing procedure as described in the Seq-Scope and Stereo-seq datasets above. We focused on four major cell types residing in the midbrain: excitatory neurons (EX, *n* =577), inhibitory neurons (IN, *n* =525), astrocytes (Astr, *n*=480), and oligodendrocytes (Olig, *n*=948), with 557-878 genes per cell type. Similar quality control steps were conducted as described in the Seq-Scope data preprocessing. After quality control, we retained 480-948 cells per cell type. We retained genes expressed in more than 50 cells, resulting in 557-878 genes per cell type for analysis.

### Real data analysis details

#### Subcellular expression pattern score

After obtaining the estimated subcellular expression intensity function λ^(*r*), we computed a subcellular expression pattern score *r*^∗^, defined as the relative position corresponding to the mode/peak of the estimated expression intensity function: *r*^∗^ = argmax λ^(*r*). Therefore, *r*^∗^ *r*∈[0,1] ranges from zero to one, with a value close to zero indicating expression enrichment in the center of the cell nucleus and a value close to one indicating expression enrichment on the cell boundary.

#### Gene clustering based on the estimated expression pattern

We clustered genes into different spatial pattern categories based on their estimated intensity functions. To do so, for each detected gene, we evaluated its estimated intensity function λ^(*r*) at 21 equidistant points, ranging from *r* =0 to *r* =1 with increments of 0.05. Additionally, we calculated the difference between consecutive functional values to obtain 20 differences. We then pooled the 21 functional values and 20 differences for each gene and used them as input for k-means clustering. We determined the optimal number of gene clusters using the Elbow method [43].

#### Transcription factor analysis

To examine the subcellular localization of transcription factors, we obtained a list of 1,358 mouse transcription factors from FANTOM5 SSTAR [26]. For all datasets, we examined the proportions of transcription factors that are measured in the datasets between pairs of gene clusters with Fisher’s exact tests.

#### Computing the unspliced-spliced ratio

In the sequencing-based datasets (Seq-Scope and Stereo-seq), for each gene in turn, we calculated the unspliced-spliced ratio for each cell by dividing the total unspliced counts (plus a pseudo count of one) by the total spliced counts (plus a pseudo count of one). We then computed the average value of this ratio across cells. We applied Mann-Whitney U tests to test the unspliced-spliced ratios between pairs of gene clusters across cell types.

#### snRNA-seq analysis

We examined the genes detected in the Seq-Scope dataset using a matched single-nucleus RNA sequencing (snRNA-seq) dataset. The snRNA-seq data was collected on mouse hepatocytes and was downloaded from the BRAIN Initiated Cell Census Network (BICCN) consortium 2021 [24]. For each gene in turn, we defined its sn-sc ratio as the average gene counts per nucleus in the snRNA-seq data divided by the average gene counts per cell in the Seq-Scope data. We applied Mann-Whitney U tests to test the sn-sc ratios between pairs of clusters across cell types. We also examined the genes detected in the MERFISH adult mouse brain dataset using a matched snRNA-seq dataset [30] that was collected on adult brain sections and calculated sn-sc ratios for the corresponding four cell types (EX, IN, Astr, and Olig) in the same way. We filtered out lowly expressed nonsignificant genes (average sc counts <1.5) due to the sparsity of the data.

#### Gene length analysis

We performed gene length analysis in the four datasets. To do so, we excluded mitochondrial genes and genes that have not been mapped to a chromosome as their gene length information is unavailable. We extracted four types of gene length measurements using GTF tools [44] from the same reference genome (mm10.gtf) that were used for alignment. The four measurements include (i) mean, (ii) median, (iii) longest single isoform, and (iv) total length across exons, all in the unit of base pairs. We then applied Mann-Whitney U tests to test the difference between pairs of gene clusters for each measurement.

#### SRP and RP analysis

In the Seq-Scope data, for each gene in turn, we used DeepSig [45] with Gencode [46] to predict whether the corresponding protein contains SRP. To do so, we downloaded protein sequences in the form of protein coding transcripts fasta files from Gencode release M28, used DeepSig to analyze the protein sequence, and referred to the genes corresponding to proteins with SRPs as SRP-coded genes. For genes with multiple protein isoforms, we used the longest isoform for SRP prediction. We examined the proportions of SRP-coded genes between pairs of gene clusters with Fisher’s exact tests. In the Stereo-seq data, we identified a list of ribosomal protein (RP) genes whose gene ID starts with RPS or RPL. These are genes of the nuclear genome that encode the protein subunits of the ribosome. These genes are expected to be enriched in the cytoplasm as ribosomal subunits are exported from the nucleus to the cytoplasm after their assembly in the nucleolus. We examined the proportions of RP genes between pairs of gene clusters with Fisher’s exact tests.

#### Cell by cell analysis in the seqFISH+ data

We randomly picked 20 cells from the seqFISH+ data. For each cell in turn, we kept genes with more than 10 counts in this analysis. We applied Bento following its instruction to classify each gene in each cell into five binary labels, corresponding to “nuclear”, “nuclear edge”, “cytoplasmic”, “cell edge”, and “none” patterns. We also applied ELLA to analyze each cell separately. We collected the estimated expression intensities λ^(*r*) of all genes and carried out k-means clustering (Methods) to obtain their pattern cluster labels.

#### Cell cycle-based analysis in the seqFISH+ data

In the seqFISH+ data, we computed the single cell gene counts and used Seurat to classify the fibroblasts into three cell subclusters corresponding to G1 (n=36, 21%), S (n=83, 49%), and G2M (n=52, 30%) cell cycle phases. We kept genes that are expressed in at least 30 cells and that have more than 3 counts in at least 5 cells, resulting in 756, 2475, and 1776 genes for G1, S, and G2M cells respectively. We then applied ELLA to analyze one gene at a time for each cell cycle subcluster. Afterwards, we retrieved ELLA genes pattern cluster labels (1-5) obtained using all fibroblasts, calculated pattern scores for genes obtained in the cell cycle specific ELLA analysis, examined these patterns cores across gene clusters, and carried out one side Mann-Whitney U test to compare the pattern scores between G1 and S/G2M subclusters (Fig. 4g, upper panel). We also focused on 723 genes commonly detected across the three cell-cycle subclusters and identified a set of genes in each gene pattern cluster with decreasing pattern scores from G1 to S and from S to G2M. Specifically, in each pattern cluster, we computed the increasement in pattern scores for each gene from G1 to S (denoted as *S*1) and another increase in pattern score from S to G2M (denoted as *S*2). We identified a set of genes with increasing pattern scores from G1 to S to G2M, characterized by *S*1 ≤ 0, *S*2 ≤ 0, and *S*1 + *S*2 < 0. We applied Mann-Whitney U tests to compare the pattern scores between the identified genes and the remaining genes at G1, S, and G2M phases across gene pattern clusters (Fig. 4g, lower panel).

## Data availability

The original public data used in this study can be accessed through the following links: (normal and diseased) mouse liver data by Seq-Scope available at https://lee.lab.medicine.umich.edu/seq-scope; mouse embryo E1S3 data by Stereo-seq available at https://db.cngb.org/search/project/CNP0001543/; mouse embryonic fibroblast data by seqFISH+ at https://figshare.com/articles/dataset/Bento_spatial_AnnData_formatted_datasets/15109236/2; and adult mouse brain data by MERFISH available at https://download.brainimagelibrary.org/29/3c/293cc39ceea87f6d/. Details about the data we used in this study are provided in the Analyzed datasets.

## Code availability

The ELLA software program and source code have been deposited at https://xiangzhou.github.io/software/ and https://github.com/jadexq/ELLA. All scripts used to reproduce all the analyses are also available on the websites.

## Acknowledgements

This study was supported by the National Institutes of Health (NIH) grants R01GM126553, R01HG011883 and R01GM144960, all to X.Z.

