## Supplementary tables, figures, and text. for "ELLA: Modeling Subcellular Spatial Variation of Gene Expression within Cells in High-Resolution Spatial Transcriptomics"

#### Supplementary Figures

**Supplementary Figure 1. An overview of high-resolution spatial transcriptomic techniques.** Selected high-resolution spatial transcriptomic techniques are listed with the original year of publication, broad categorization of the technique (colored triangles), achieved spatial resolutions (red boxes), and the number of genes capable of being accessed (blue circles). These techniques achieve expression measurement resolutions at cellular or subcellular levels.

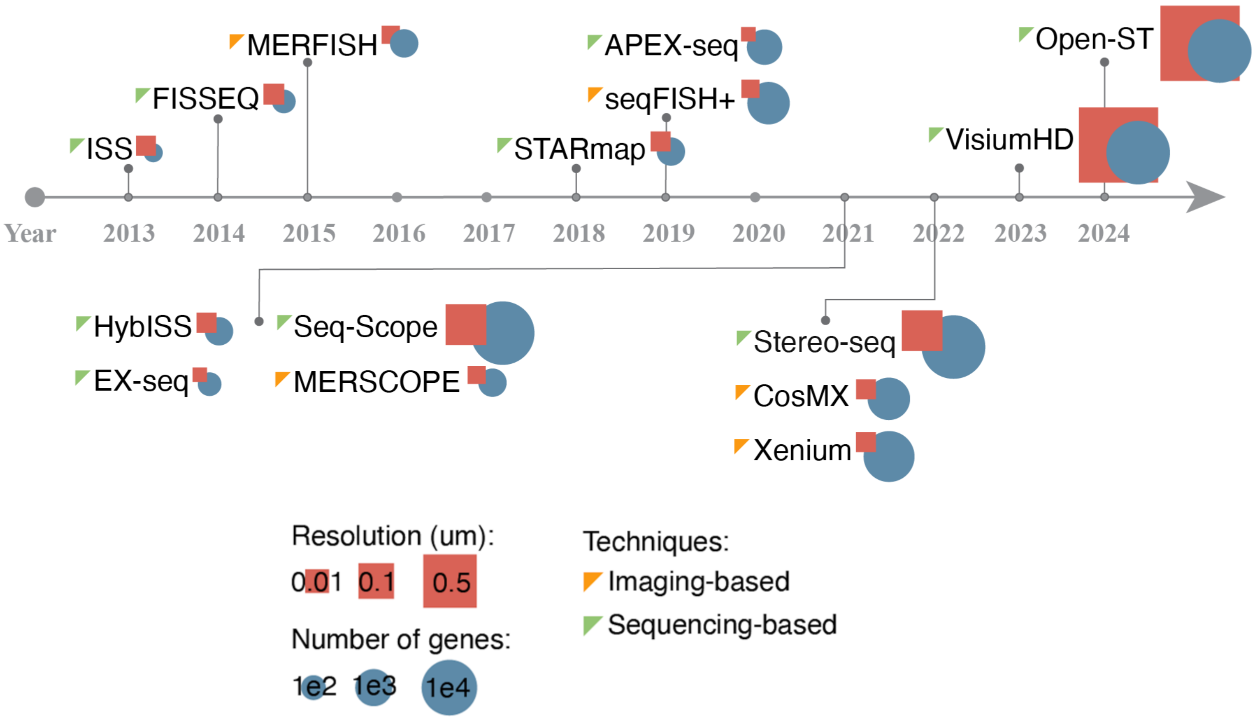

* For untargeted methods such as FISSEQ, APEX-seq, Seq-Scope, we considered the number of genes presented in their original studies.
* The sizes of square and circle markers are created to show roughly the relative resolution or number of genes between technologies. Please refer to the original studies or online information for their precise resolution and gene panel information.

**Supplementary Figure 2. Real cells used in the simulations.** Cell boundary and nuclear boundary are displayed for randomly selected embryonic fibroblast cells from the seqFISH+ data that are used in simulations.

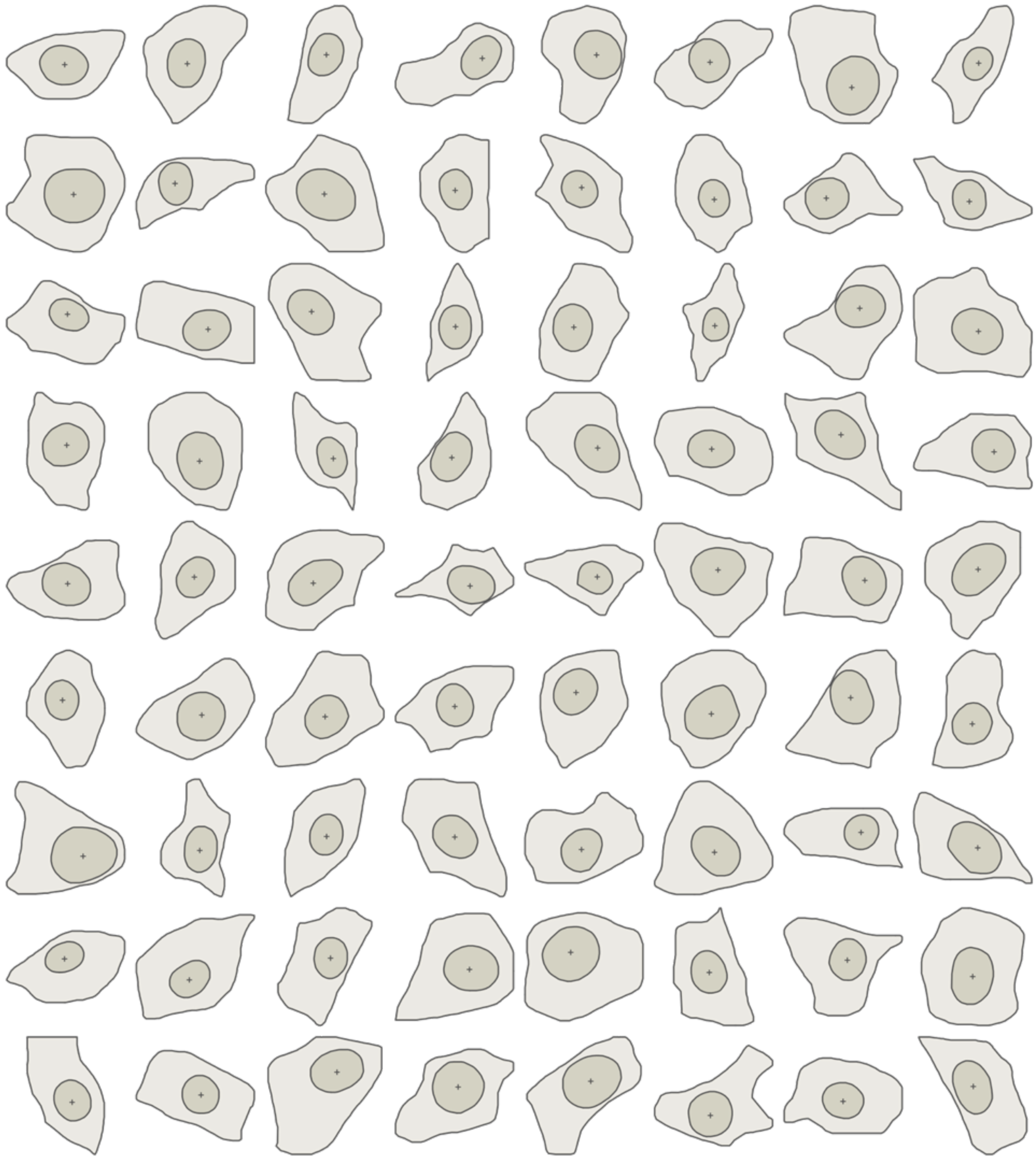

**Supplementary Figure 3. Cells and genes in the null baseline simulation.** Gene expression counts for five randomly selected genes are displayed for randomly selected cells in the baseline null simulation, where expression counts are randomly distributed within cells without any spatial patterns.

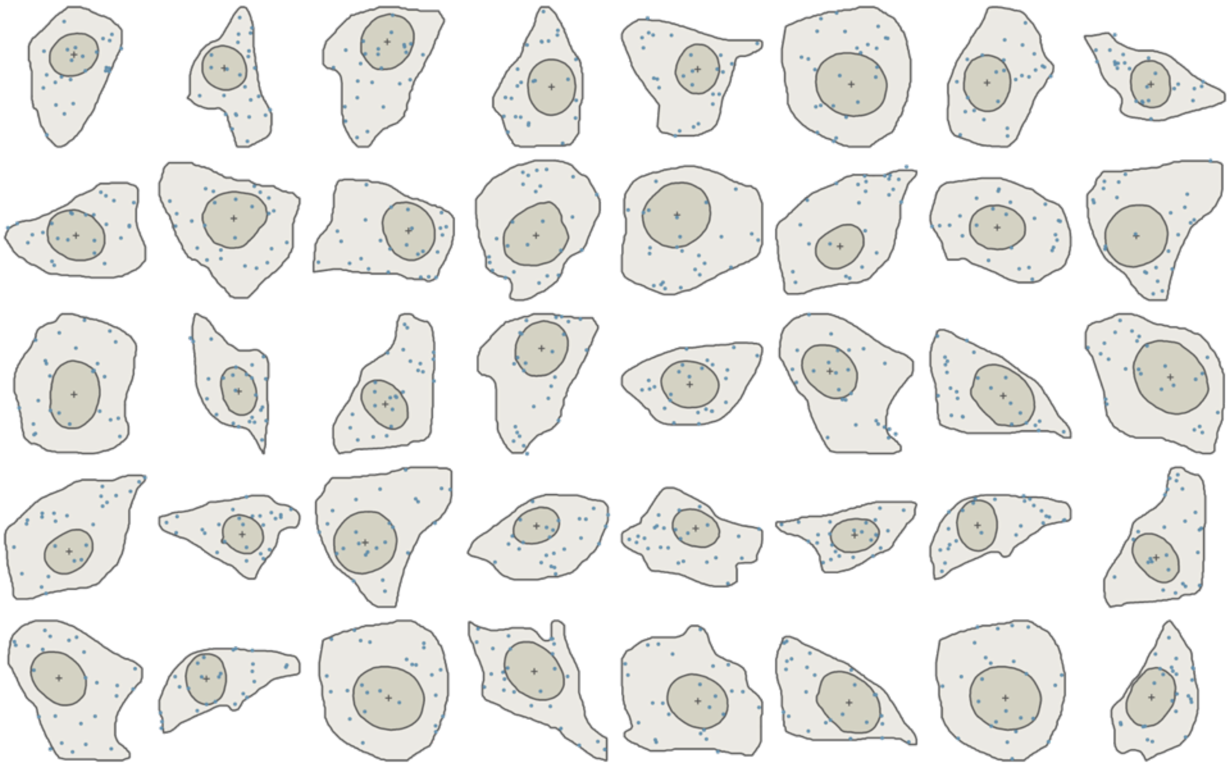

**Supplementary Figure 4. Cells and genes in the alternative baseline symmetric simulations, patterns 1-5.** Gene expression counts for five genes in randomly selected cells are shown for the alternative baseline symmetric simulation. Rows 1-5 correspond to Patterns 1-5, respectively. The subcellular expression intensity $\lambda(r)$ levels are visualized in gray color, where a darker color corresponds to a higher intensity.

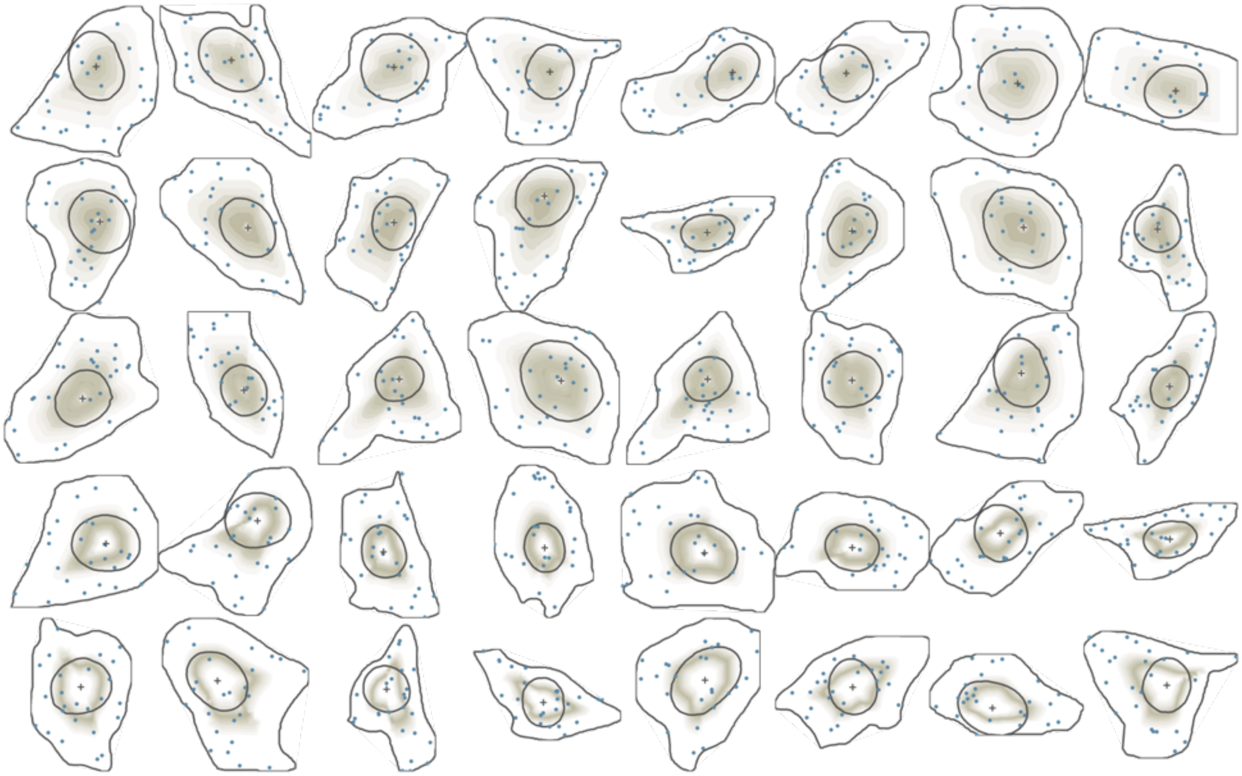

**Supplementary Figure 5. Cells and genes in the alternative baseline symmetric simulations, Patterns 6-11.** Gene expression counts for five genes in randomly selected cells are shown for the alternative baseline symmetric simulation. Rows 1-6 correspond to Patterns 6-11, respectively. The subcellular expression intensity λ(r) levels are visualized in gray color, where a darker color corresponds to a higher intensity.

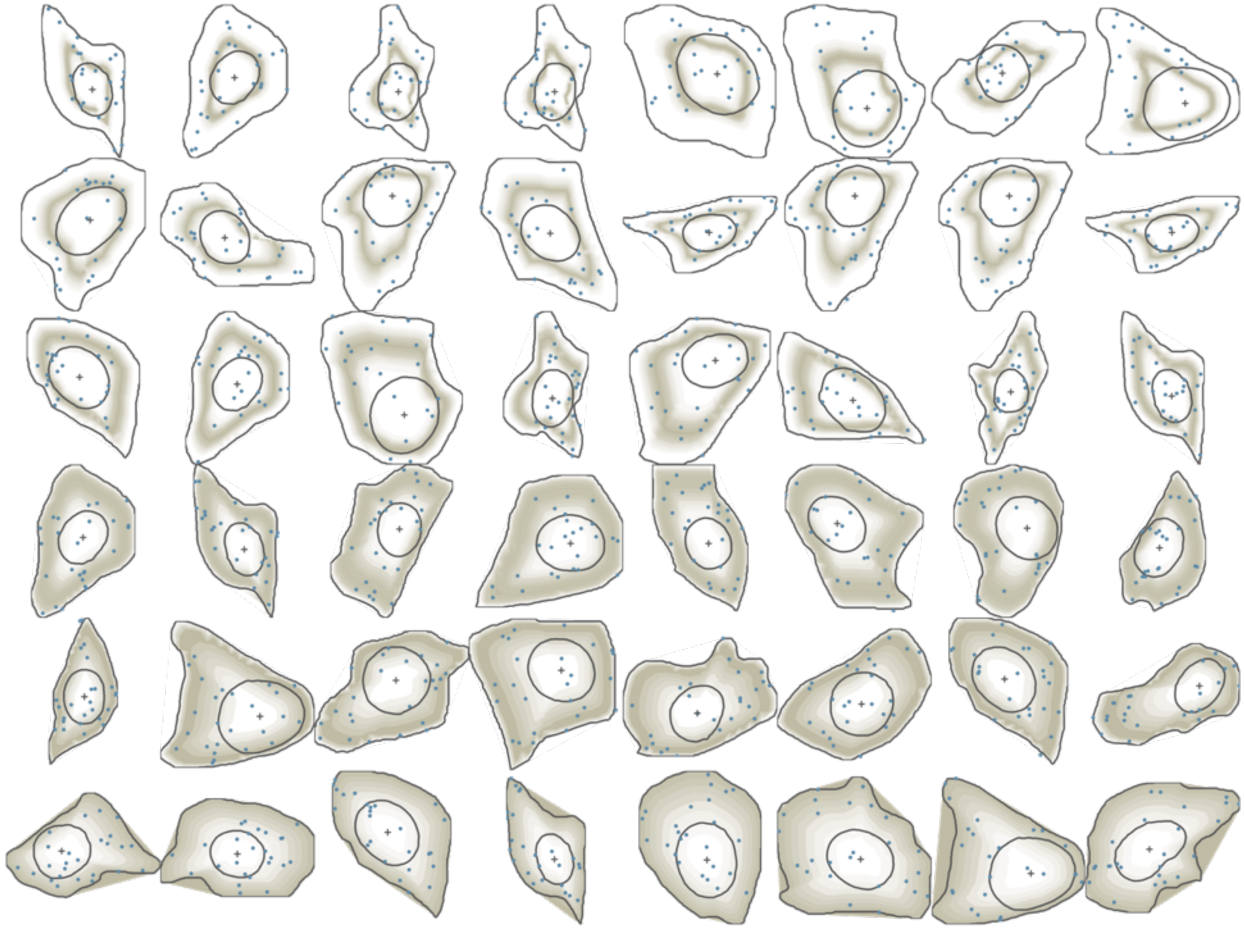

**Supplementary Figure 6. Cells and genes in the alternative asymmetric simulations.** Gene expression counts for one gene in randomly selected cells are shown for the alternative baseline asymmetric simulation across three settings: **a.** radial uniform pattern setting; **b.** radial-cyto pattern setting; **c.** punctate pattern setting. The subcellular expression intensity λ(r) levels are visualized in gray color, where a darker color corresponds to a higher intensity.

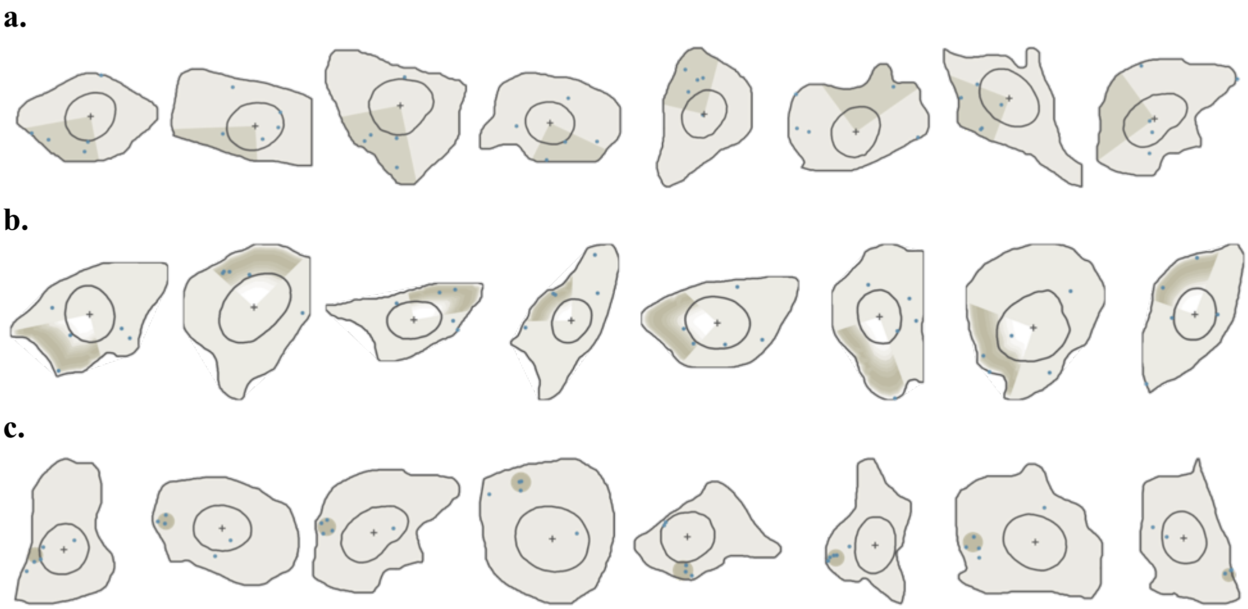

**Supplementary Figure 7. Quantile-quantile plots of the expected and observed -log10 P values in the null simulation with varying numbers of cells ($n$).** The gene expressions are randomly distributed spatially within the cells and the number of cells varies across $n$=10, 20, 50, 100, 200, 300, and 500. ELLA was compared to SPRAWL, Bento, and Wilcox methods.

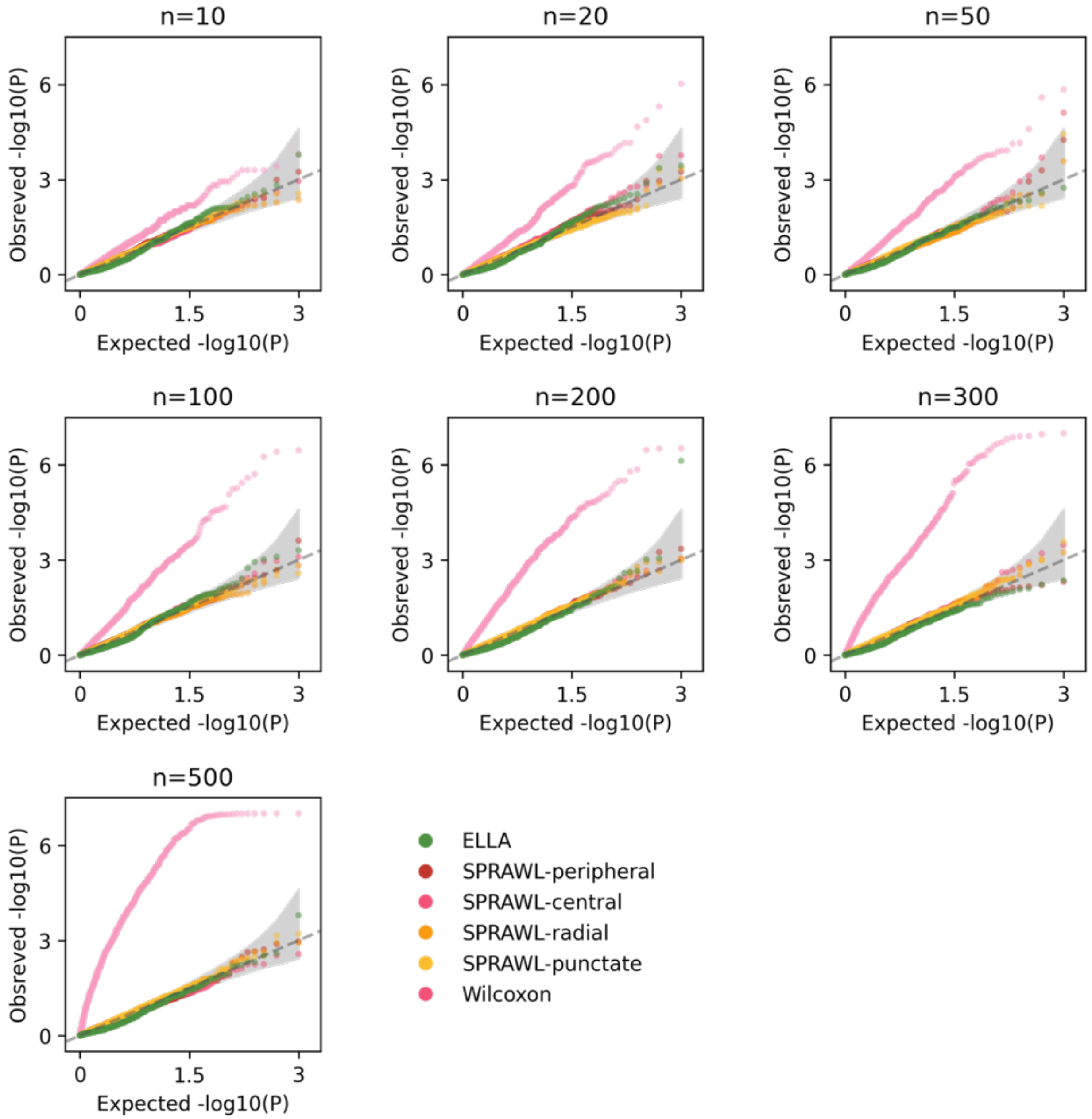

**Supplementary Figure 8. Quantile-quantile plots of the expected and observed -log10 P values in the null simulation with varying expression levels ($m$).** The gene expressions are randomly distributed spatially within the cells and the expression level varies across $m$=1, 2, 5, 10, 20, 50, and 100. ELLA was compared to SPRAWL, Bento, and Wilcox methods. SPRAWL radial and punctate metrics failed to produce any P values in the $m$=1 setting.

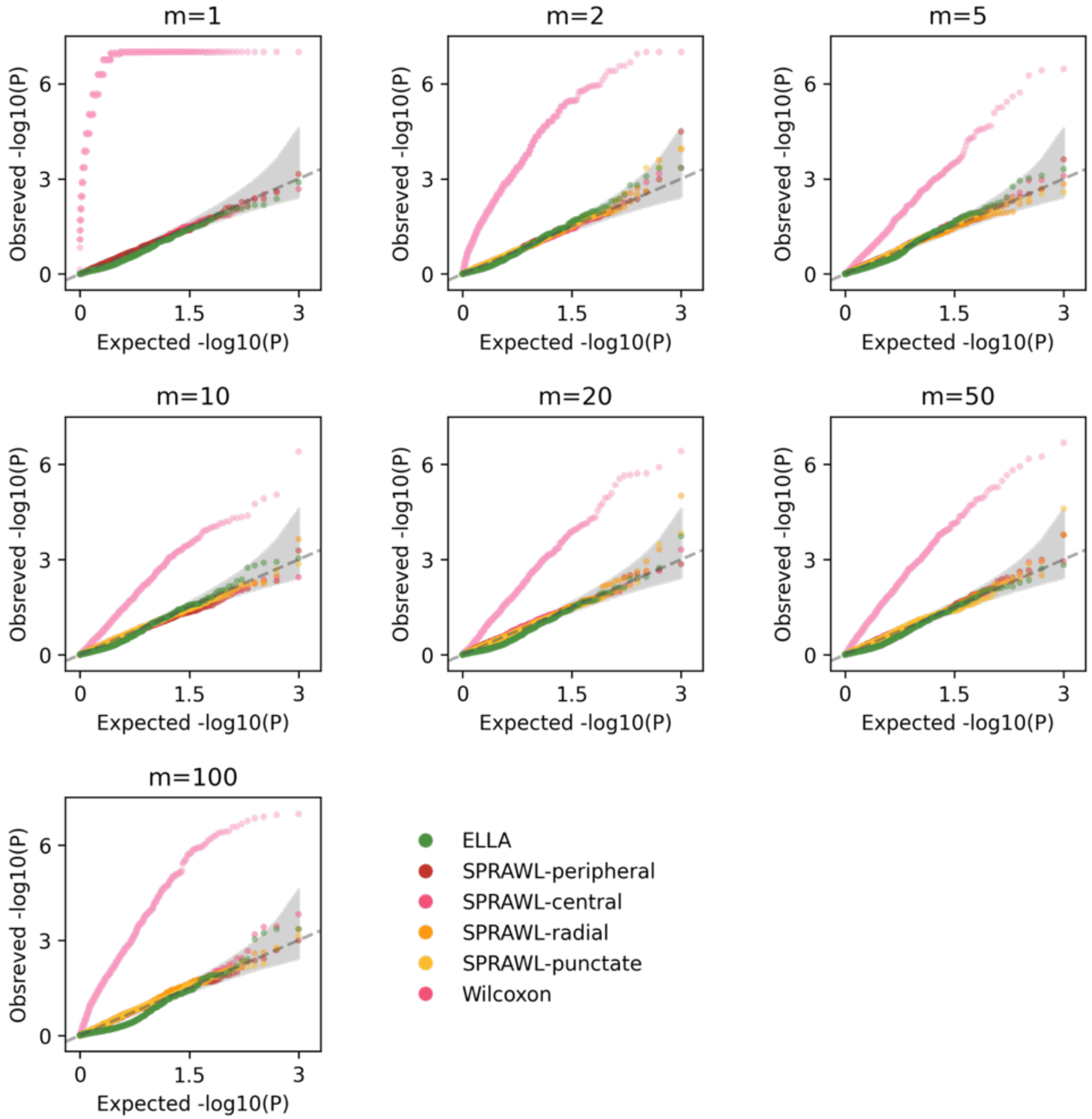

**Supplementary Figure 9. The heteroskedastic variances of the standardized nuclear and cytoplasmic expression counts used in the Wilcox method in the baseline null simulation.** Histogram of the standardized nuclear and cytoplasmic expression of 12 randomly selected genes in the baseline null simulation displaying the standardized nuclear expression counts have a smaller variance than the standardized cytoplasmic expression counts.

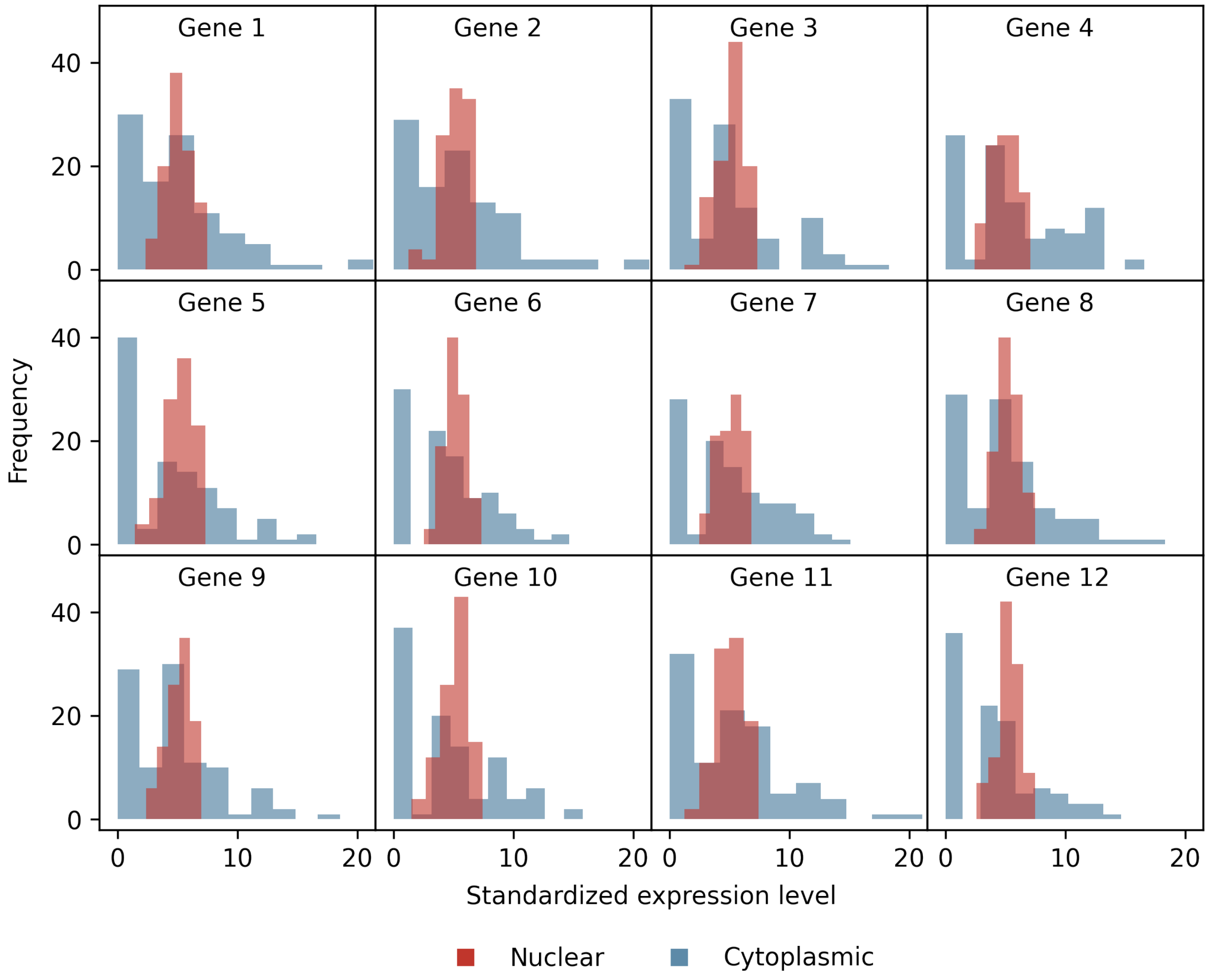

**Supplementary Figure 10. Alternative simulations with varying cell number ($n$), expression level ($m$), and pattern strength ($s$).** Scatter plots of the powers at 5% FDR of ELLA, SPRAWL, and Wilcox in the alternative simulation Pattern 1 with: **a.** varying number of cells $n$=10, 20, 50, 100, 200, 300, or 500; **b.** varying expression level $m$=1, 2, 5, 10, 20, 50, or 100; **c.** varying pattern strength $s$=0.1, 0.2, 0.3, 0.4, 0.5, 0.6, 0.7, 0.8, 0.9 or 1. The power of ELLA increases much more rapidly compared to the other compared methods.

**a.**

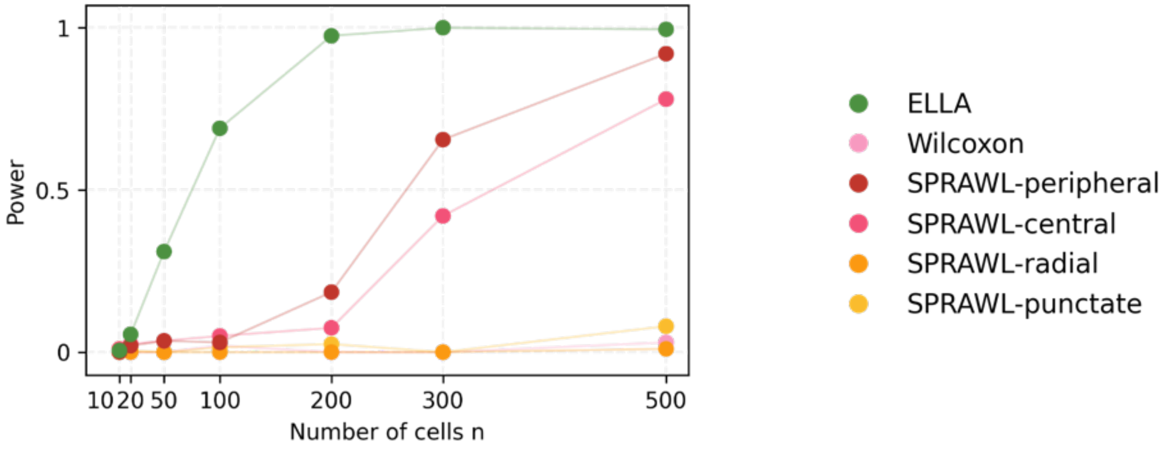

**b.**

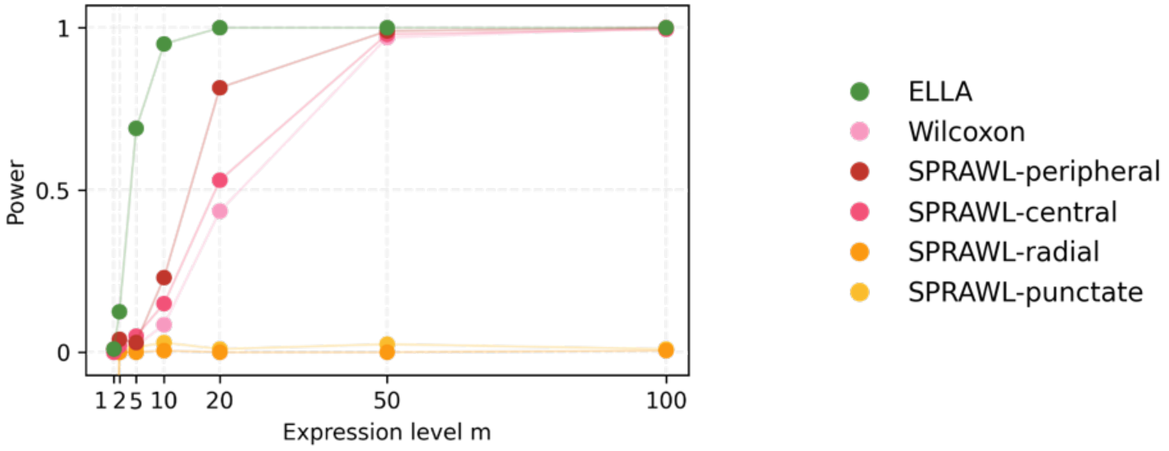

**c.**

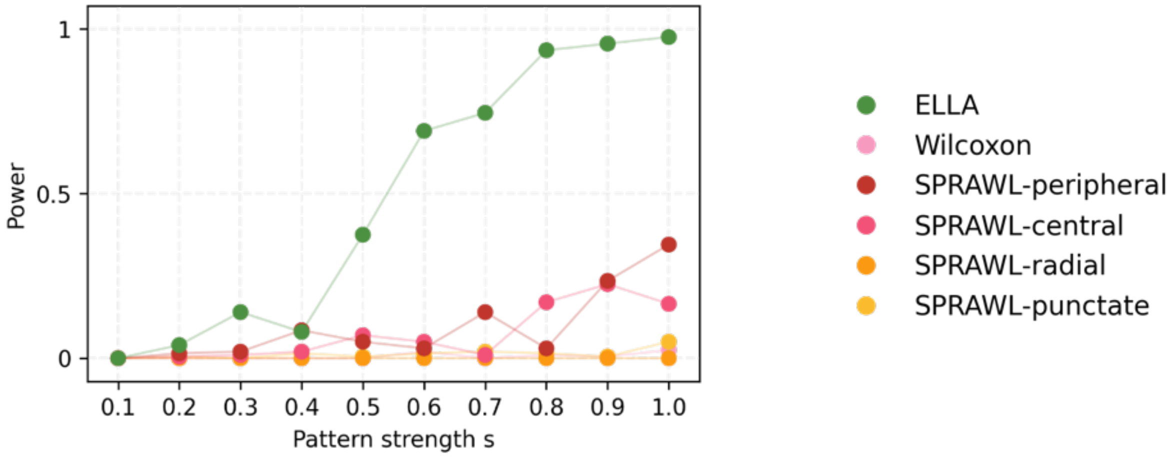

**Supplementary Figure 11. Expression intensity estimations of the eleven symmetric patterns in the baseline alternative simulations.** Plots display the estimated expression intensities across genes ($\hat{\lambda}(r)$; light green for significant genes, light grey for nonsignificant genes) overlayed with the true expression intensity (in dark green with dashed line) in each of the eleven baseline alternative symmetric simulation settings. **a.** Both the true and estimated subcellular expression intensities ($\lambda(r)$ and $\hat{\lambda}(r)$) are standardized by the areas under the curve. **b.** Both the true and estimated $\lambda(r)$ are standardized as $\left[\lambda(r) - \min_r \lambda(r)\right] / \left[\max_r \lambda(r) - \min_r \lambda(r)\right]$ to be between 0 and 1. The y-axis corresponds to the standardized intensity levels and the x-axis corresponds to the relative position (0-nuclear center, 1-cell boundary).

**a.**

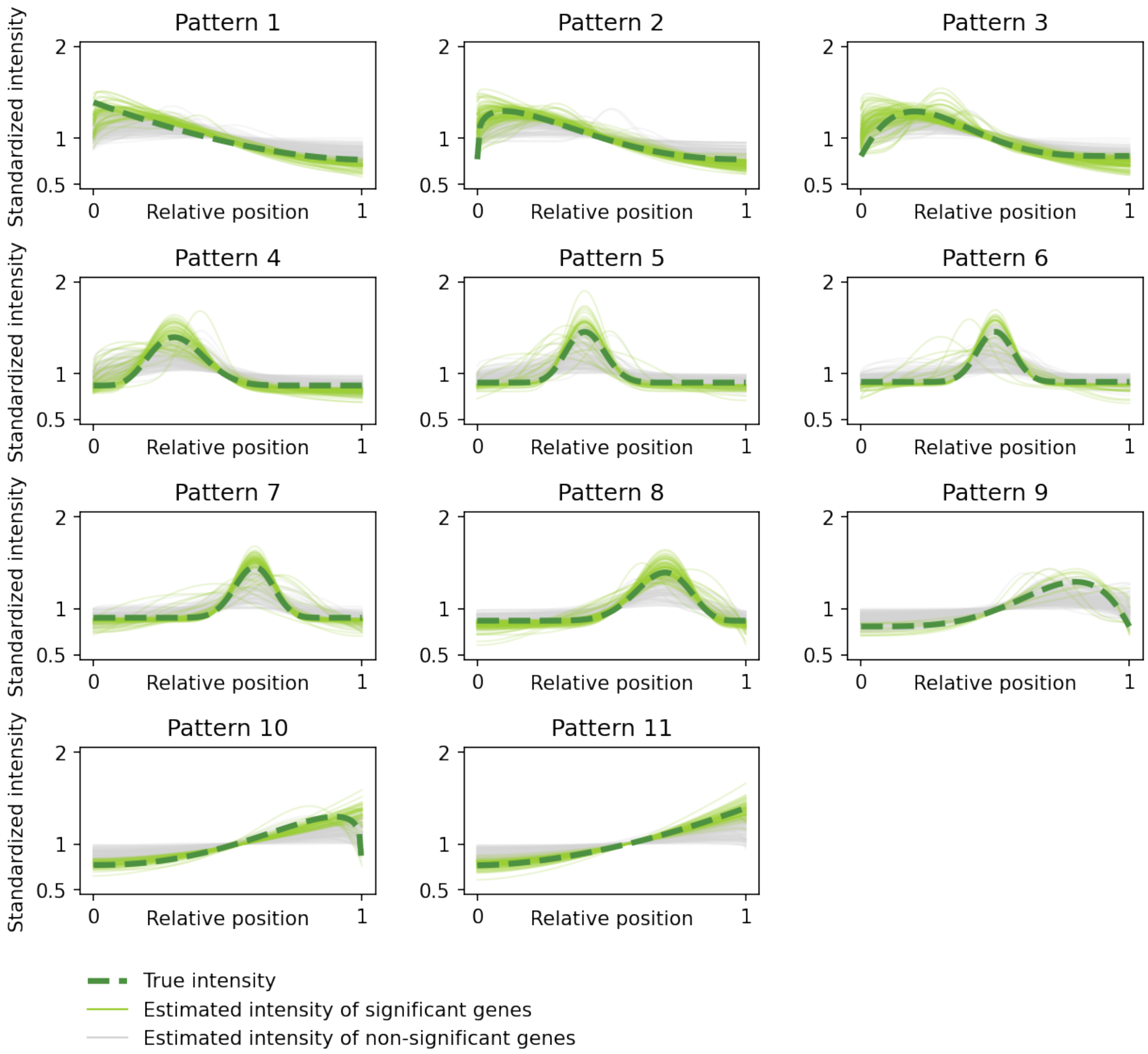

(Fig. S11 con'd)

**b.**

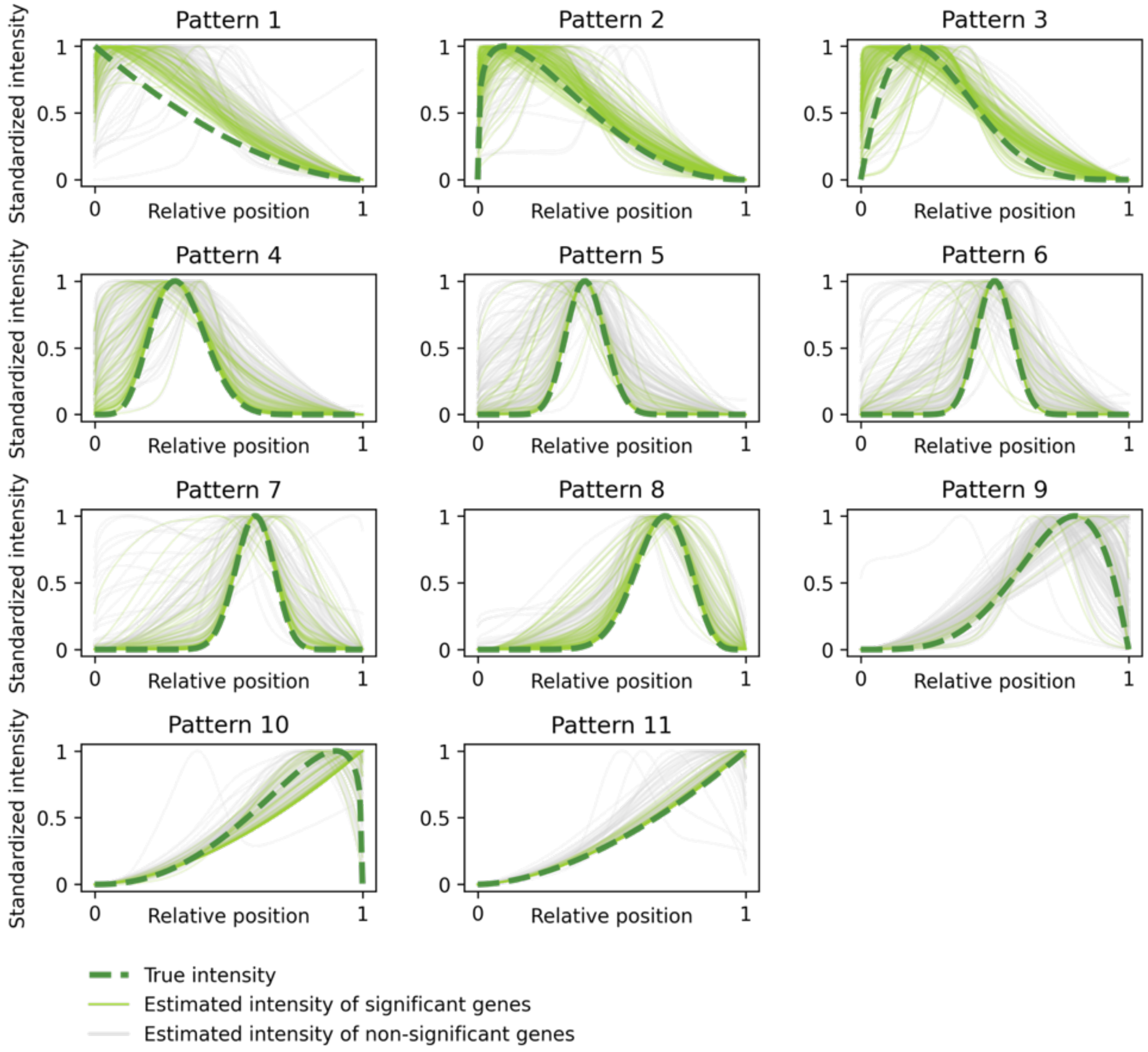

**Supplementary Figure 12. Expression intensity estimations of the three asymmetric patterns in the alternative simulations.** Plots display the estimated expression intensities across genes ($\hat{\lambda}(r)$; light green for significant genes, light grey for nonsignificant genes) overlayed with the true expression intensity (in dark green with dashed line) in the radial-unif, radial-cyto, and punctate-cyto pattern settings. Both the true and estimated subcellular expression intensities ($\lambda(r)$ and $\hat{\lambda}(r)$) are standardized as $\left[\lambda(r) - \min_r \lambda(r)\right] / \left[\max_r \lambda(r) - \min_r \lambda(r)\right]$ to be between 0 and 1 where the y-axis corresponding to the standardized intensity levels and the x-axis corresponding to the relative location (0-nuclear center, 1-cell boundary).

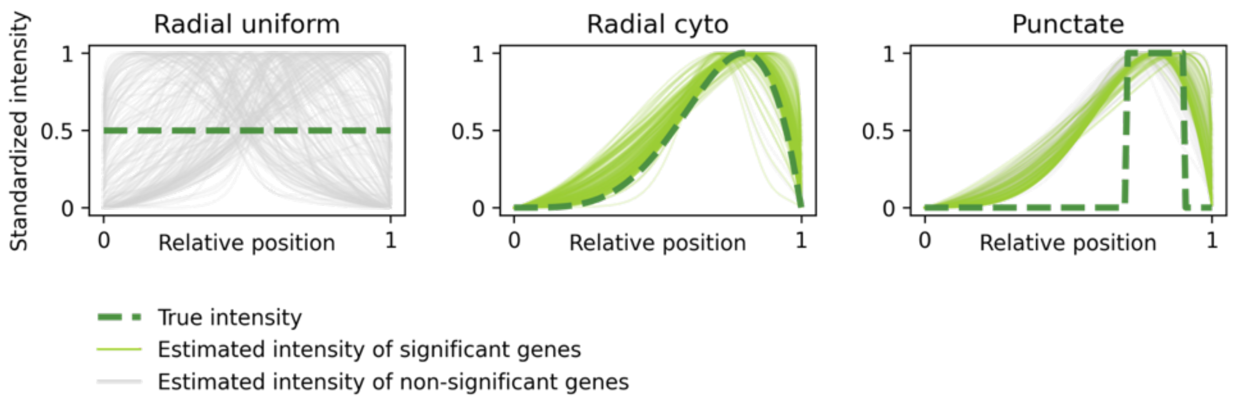

**Supplementary Figure 13. Expression pattern score estimations of the eleven symmetric patterns in the baseline alternative simulations.** Histograms display the estimated expression pattern scores across genes (light green for significant genes, light grey for nonsignificant genes) overlayed with the true pattern score (in dark green with dashed line) in each of the eleven baseline alternative symmetric simulation settings.

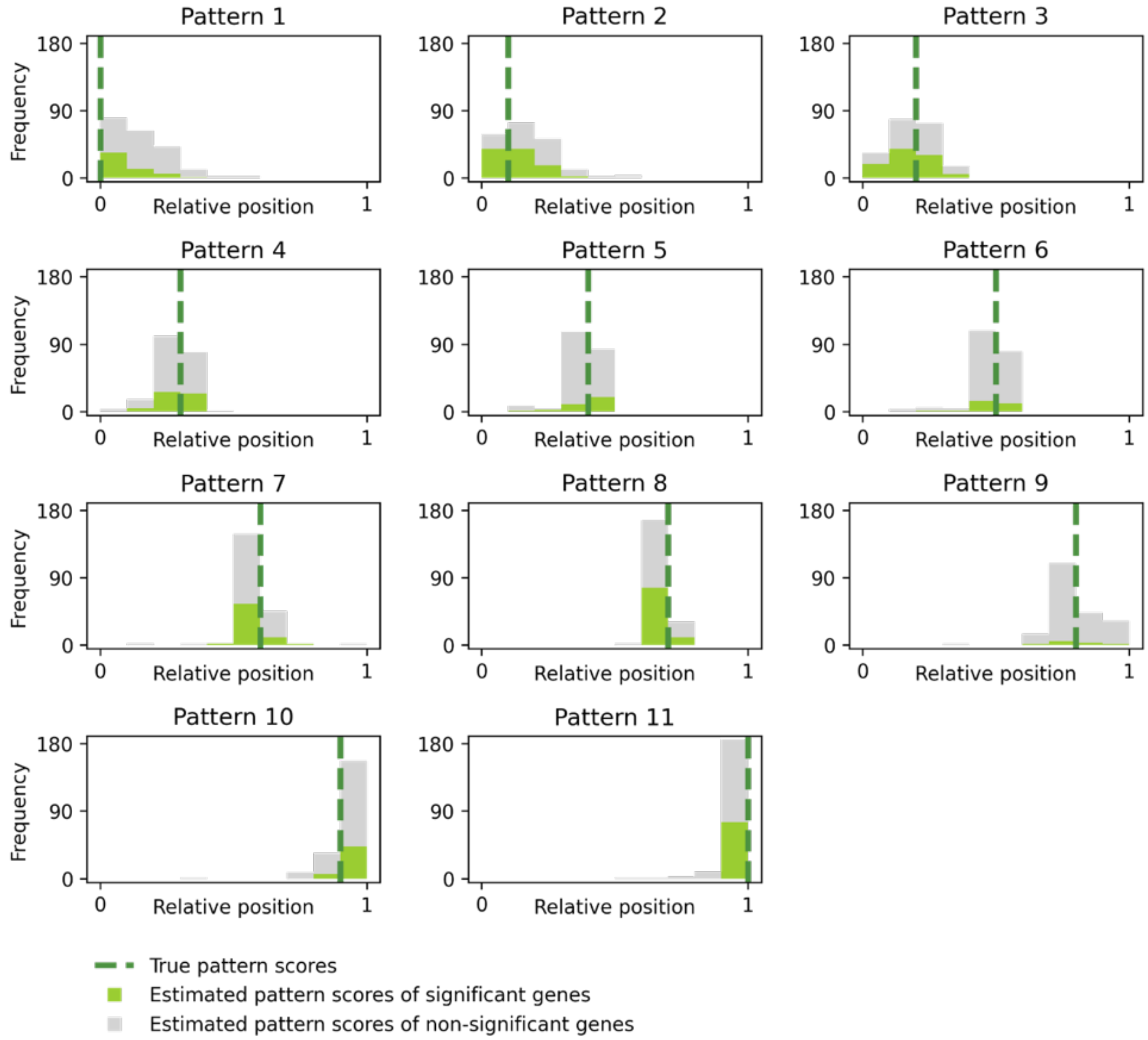

**Supplementary Figure 14. ELLA expression pattern score estimation of the three asymmetric patterns in the alternative simulations.** Histograms display the estimated expression pattern scores across genes (light green for significant genes, light grey for nonsignificant genes) overlayed with the true pattern score (in dark green with dashed line) in the radial-unif, radial-cyto, and punctate-cyto pattern settings.

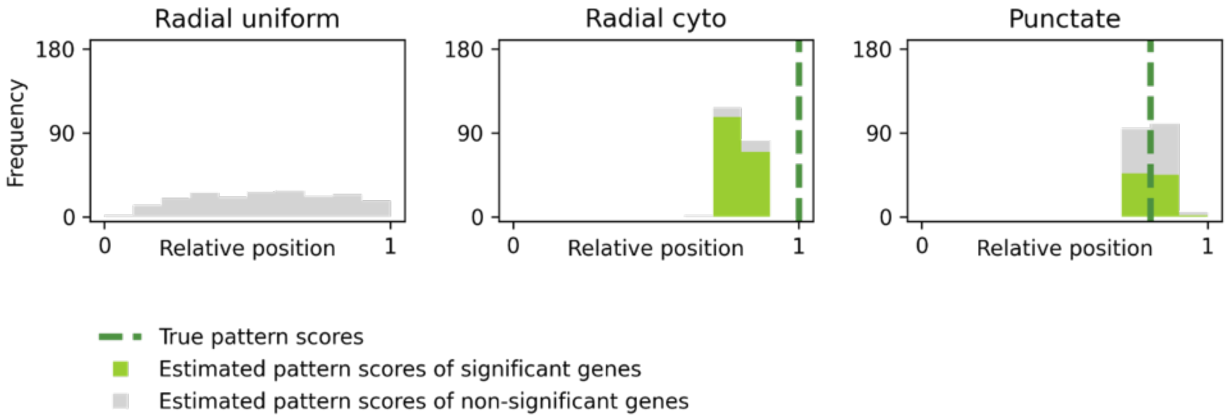

**Supplementary Figure 15. The setups of the additional simulations with one cell.** Additional simulations with only one cell were performed to compare ELLA with Bento. Five patterns (from column 1 to column 5) with expressions enriched in nucleus (2 patterns), nuclear edge (1), cytoplasm (1), and cellular boundary (1) under high expression level ($m$=30) and high pattern strength ($s$=9) were considered. **a.** Upper panels display the true expression intensity function $\lambda(r)$ across the five patterns. Middle panel displays the NHPP density function $\lambda^*(r)$ across patterns. Lower panel displays the probabilities of a gene expression falls in each bin of relative position $[0, 0.2)$, $[0.2, 0.4)$, $[0.4, 0.6)$, $[0.6, 0.8)$, $[0.8,1]$ across patterns. **b.** Gene expression counts for one gene in a randomly selected cell are shown across five settings.

**a.**

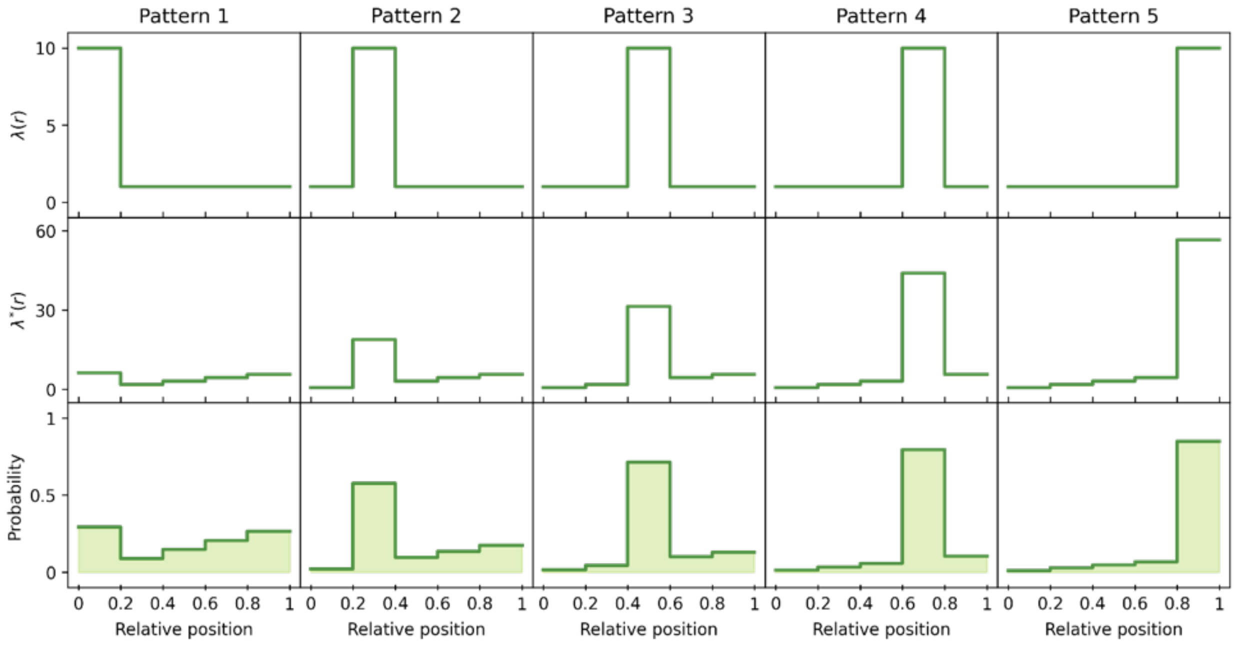

**b.**

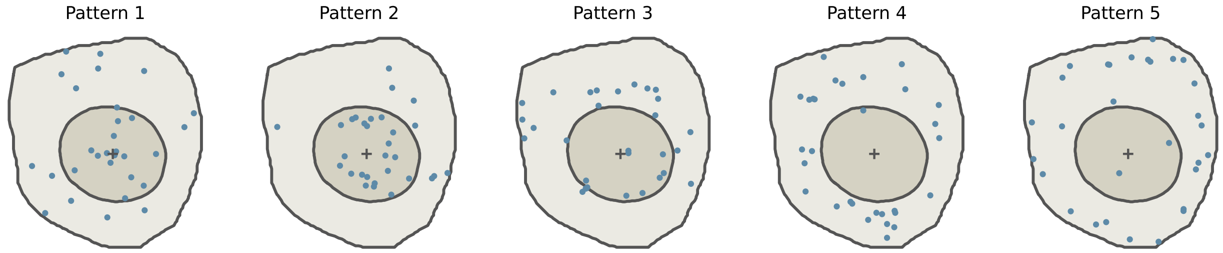

**Supplementary Figure 16. Expression intensity estimations of the five patterns in the additional simulations with one cell.** Plots display the estimated expression intensities across genes ($\hat{\lambda}(r)$; light green for significant genes, light grey for nonsignificant genes) overlayed with the true expression intensity (in dark green with dashed line) in the five patterns. Both the true and estimated subcellular expression intensities ( $\lambda(r)$ and $\hat{\lambda}(r)$ ) are standardized as $\left[\lambda(r) - \min_r \lambda(r)\right] / \left[\max_r \lambda(r) - \min_r \lambda(r)\right]$ to be between 0 and 1 where the y-axis corresponds to the standardized intensity levels and the x-axis corresponds to the relative location (0-nuclear center, 1-cell boundary).

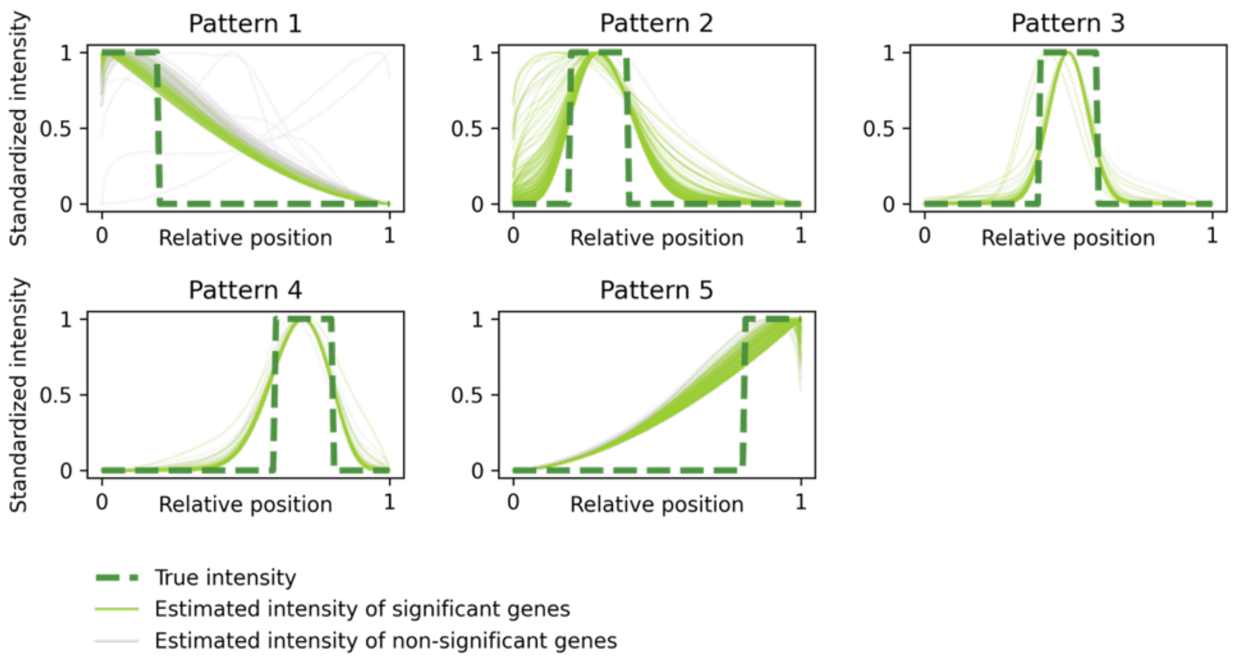

**Supplementary Figure 17. ELLA expression pattern score estimation estimations of the five patterns in the additional simulations with one cell.** Histograms display the estimated expression pattern scores across genes (light green for significant genes, light grey for nonsignificant genes) overlayed with the true pattern score (in dark green with dashed line) in the five patterns.

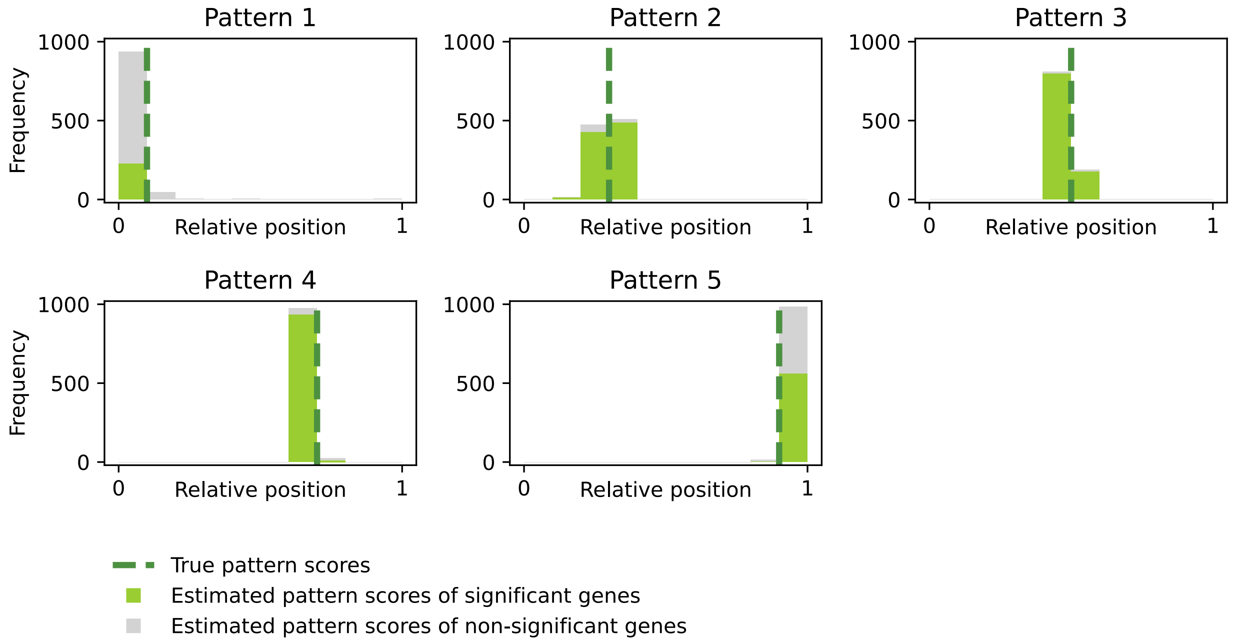

**Supplementary Figure 18. The concatenated 4X H&E images for the region of normal tiles and TD tiles in Seq-Scope.**

Region of normal tiles:

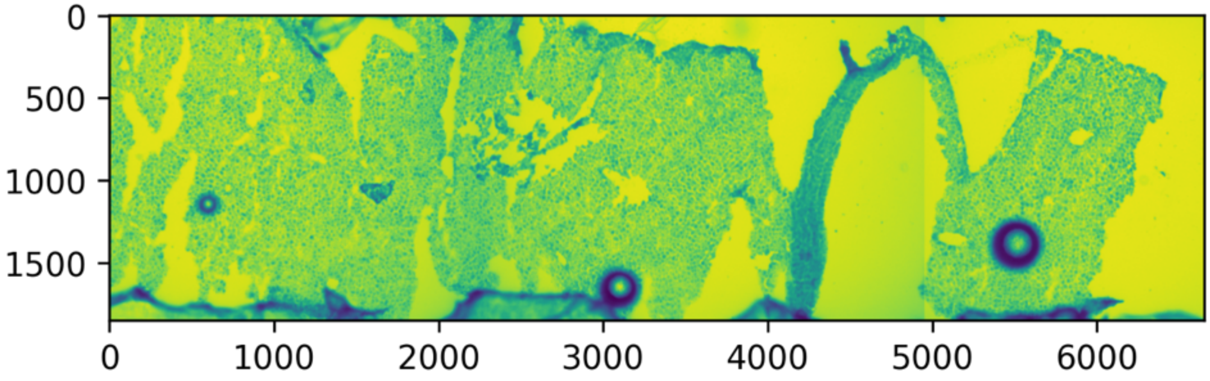

Region of TD tiles:

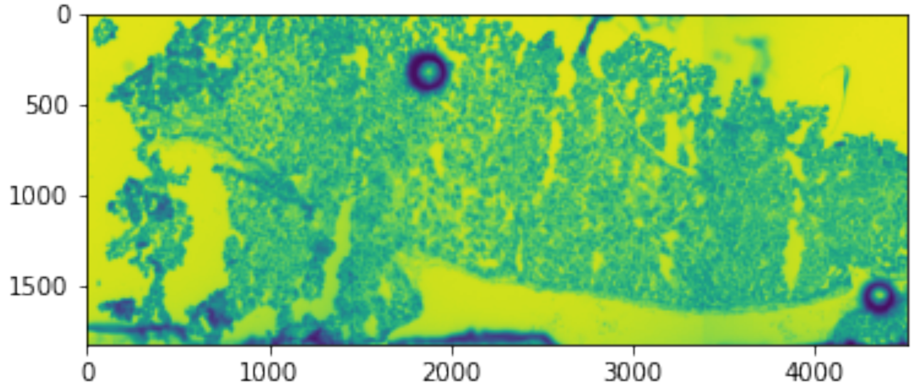

**Supplementary Figure 19. The concatenated 4X H&E images for the region of normal tiles and TD tiles after removing abnormal areas or subregions (yellow) with no tiles.**

Region of normal tiles:

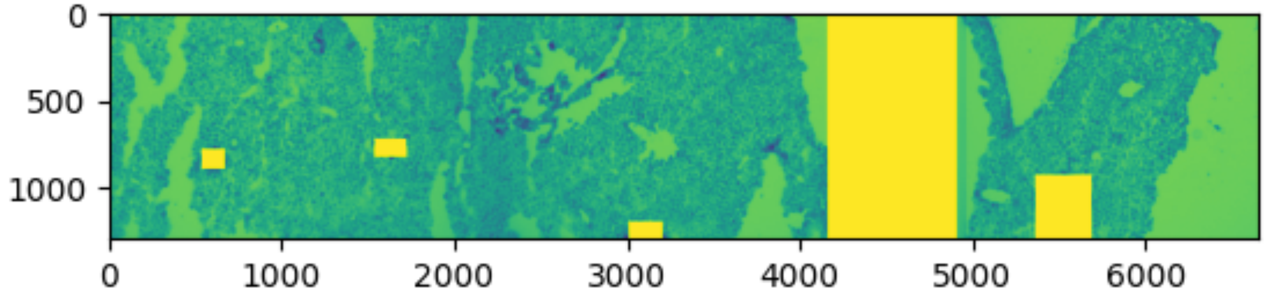

Region of TD tiles:

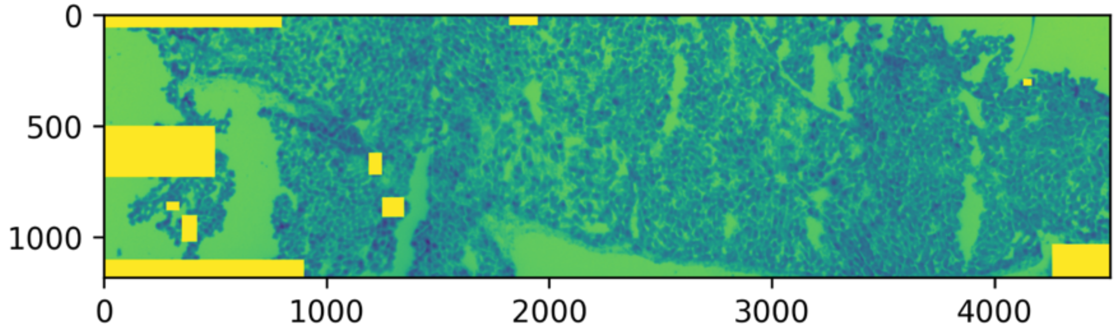

**Supplementary Figure 20. Cell segmentations produced by Cellpose based on raw normal 4X H&E images in Seq-Scope.** The cell segmentation boundaries (red) are overlayed on the H&E image (greyscale). Poor cell segmentation results were generated.

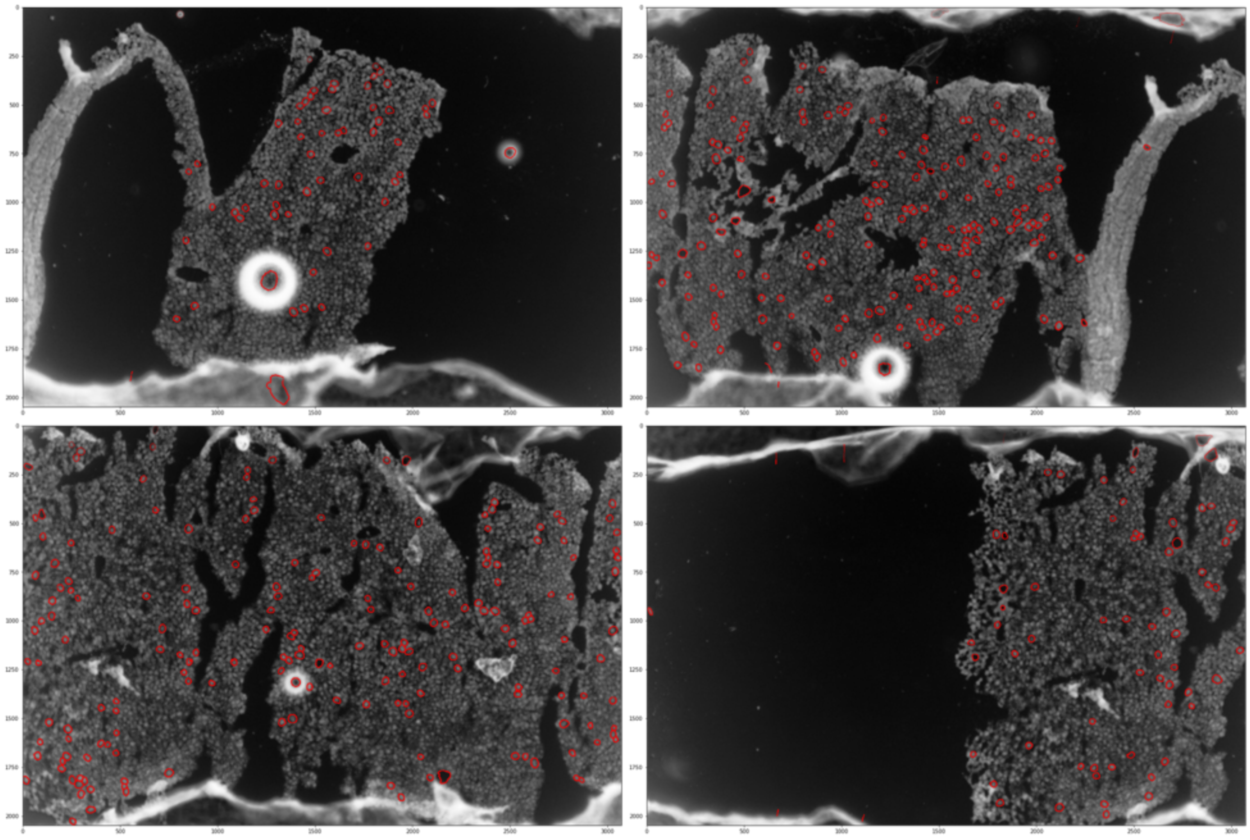

**Supplementary Figure 21. Cell segmentations produced by Cellpose based on raw TD 4X H&E images in Seq-Scope.** The cell segmentation boundaries (red) are overlayed on the H&E image (greyscale). Poor cell segmentation results were generated.

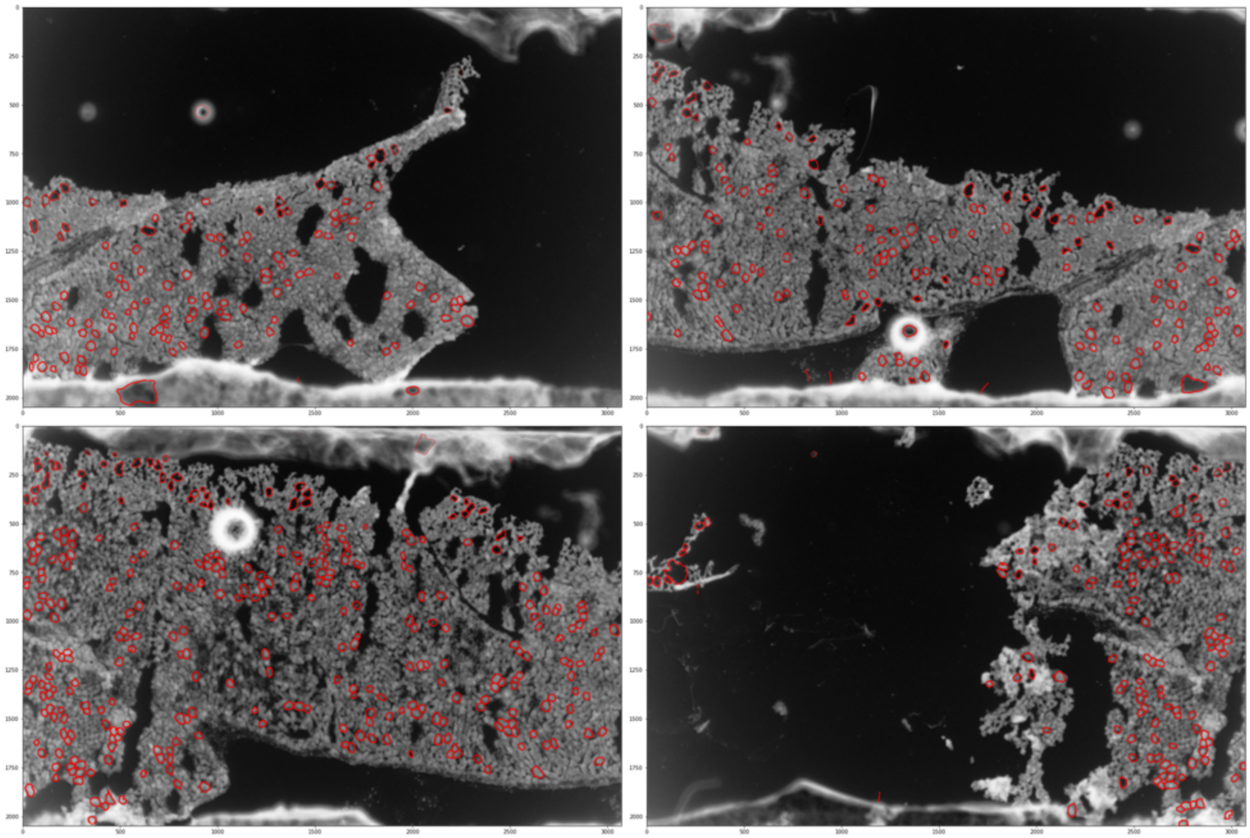

**Supplementary Figure 22. Cell segmentations produced by Cellpose based on the concatenated normal 4X H&E image removing abnormal areas or subregions with no tiles in Seq-Scope.** Upper panel with 4 images shows the outputs from Cellpose. Lower panel shows the cell segmentation boundaries (red) are overlayed on the H&E image (greyscale) produced by Cellpose.

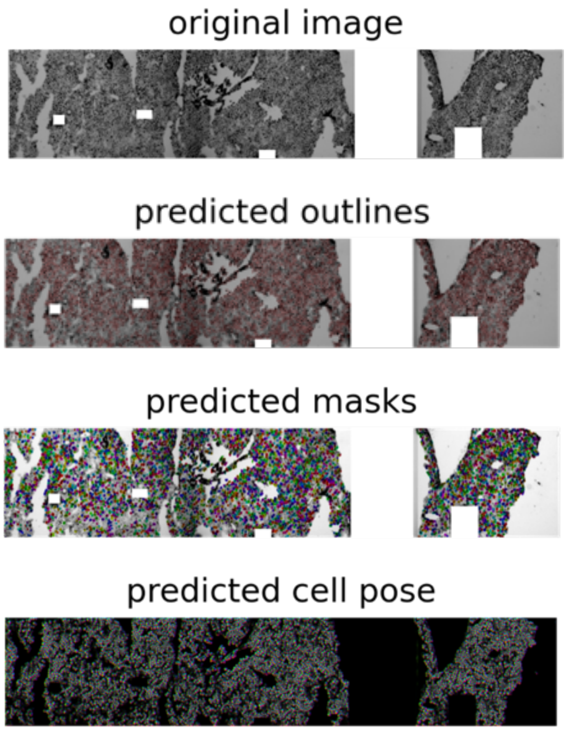

Cell segmentations:

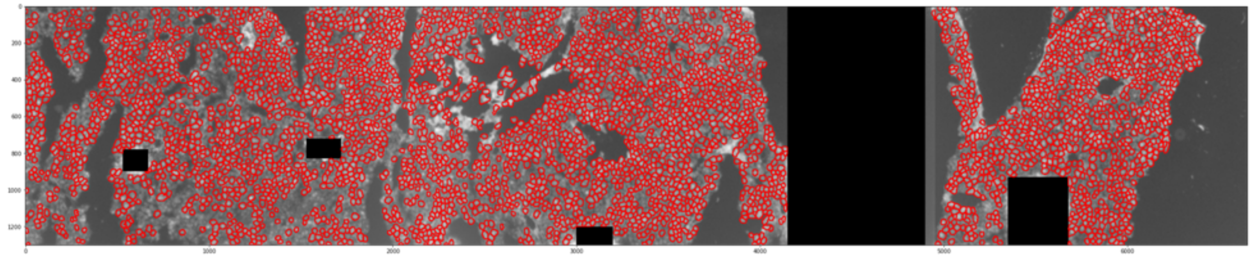

**Supplementary Figure 23. Cell segmentations produced by Cellpose based on the concatenated TD 4X H&E image removing abnormal areas or subregions with no tiles in Seq-Scope.** Upper panel with 4 images shows the outputs from Cellpose. Lower panel shows the cell segmentation boundaries (red) are overlayed on the H&E image (greyscale) produced by Cellpose.

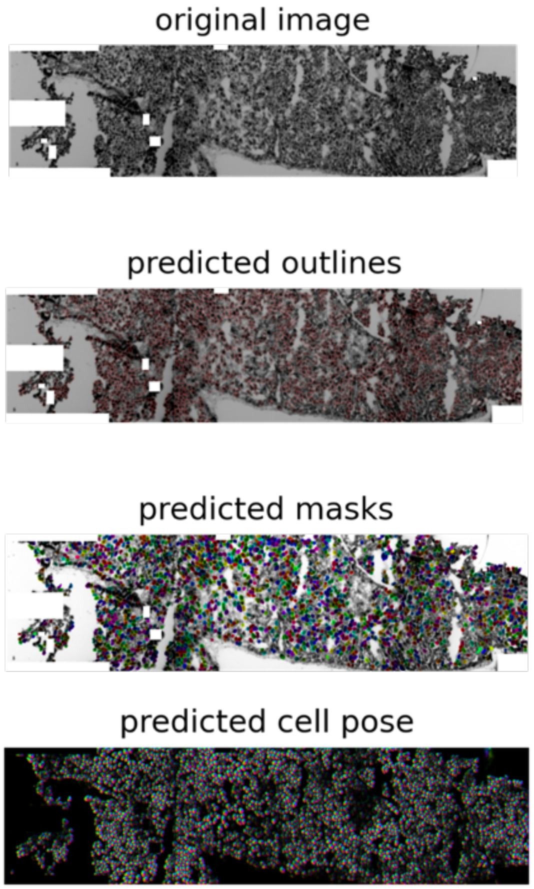

Cell segmentations:

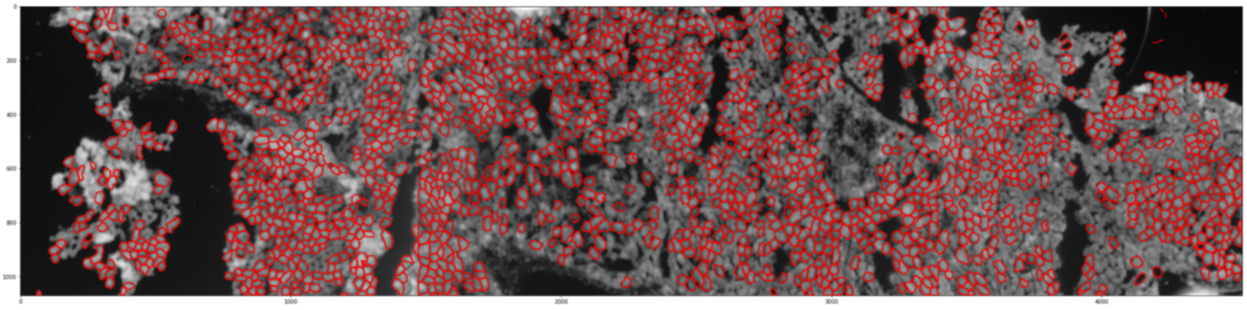

**Supplementary Figure 24. Total unspliced counts across tiles in Seq-Scope.** For each tile, the total unspliced expression counts of all genes across locations are plotted where a darker color corresponds to a higher count.

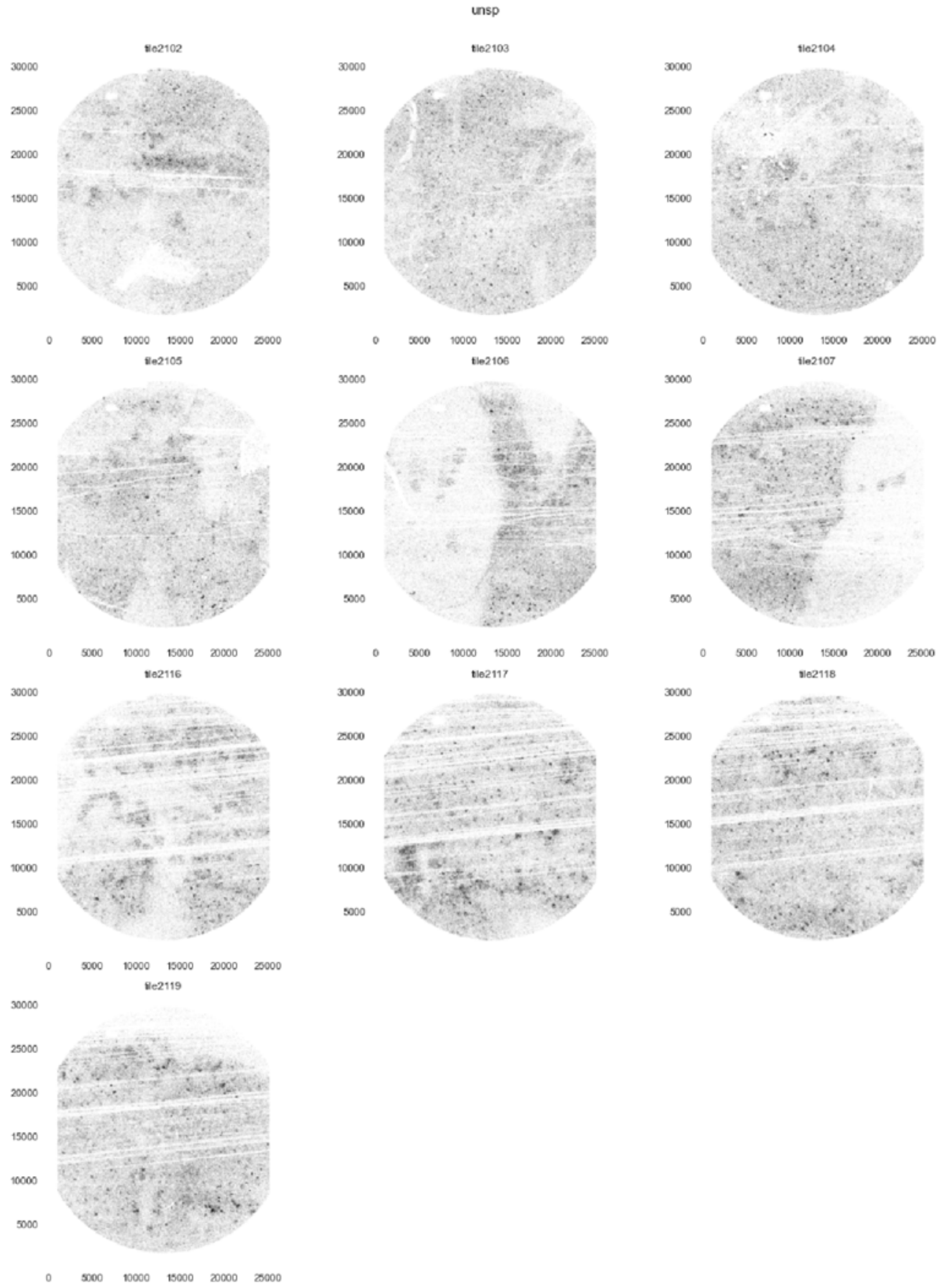

**Supplementary Figure 25. Total spliced counts across tiles in Seq-Scope.** For each tile, the total spliced expression counts of all genes across locations are plotted where a darker color corresponds to a higher count.

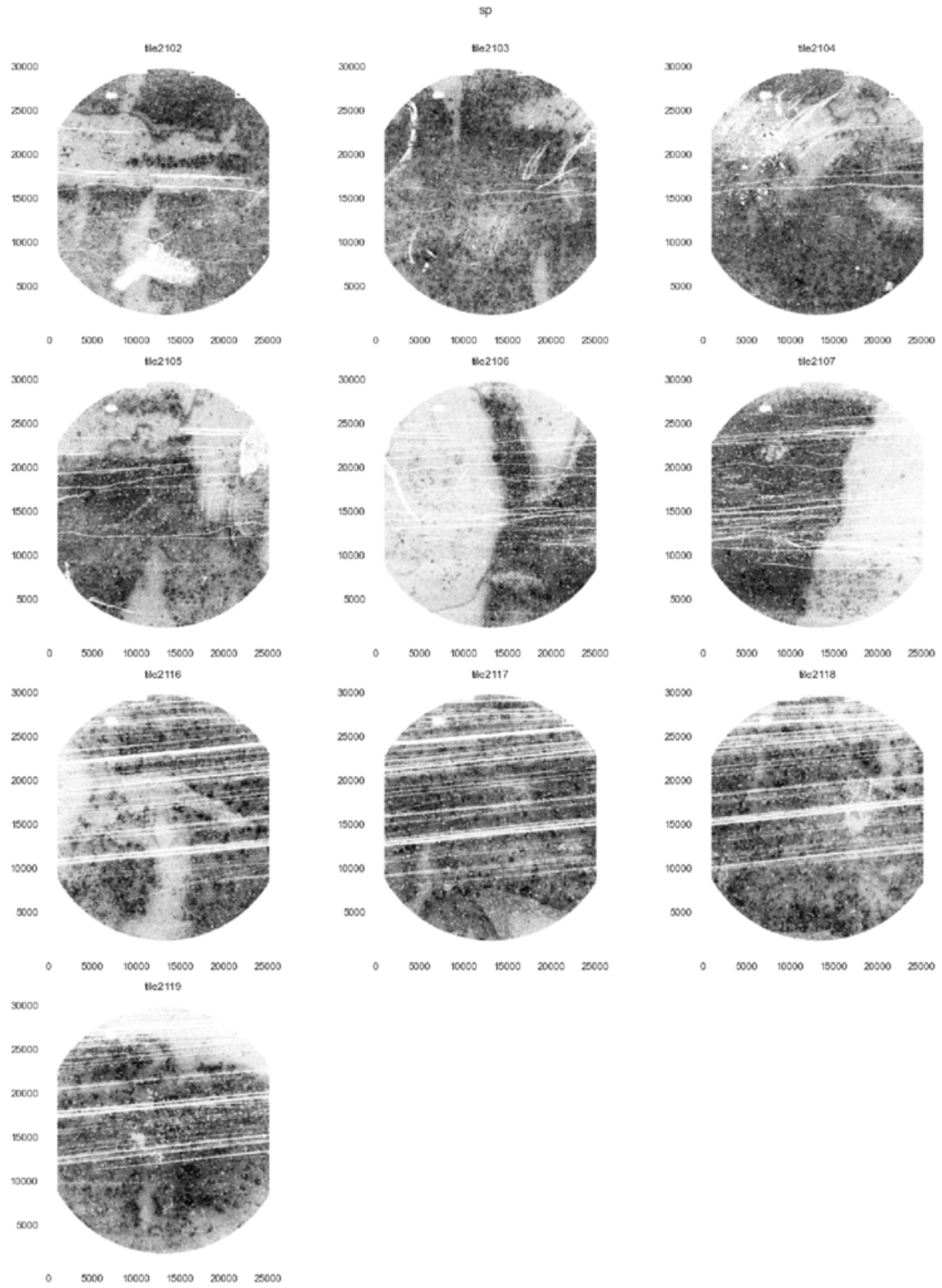

**Supplementary Figure 26. Overlay of the total unspliced counts (green) and total spliced counts (grey) across locations and tiles.**

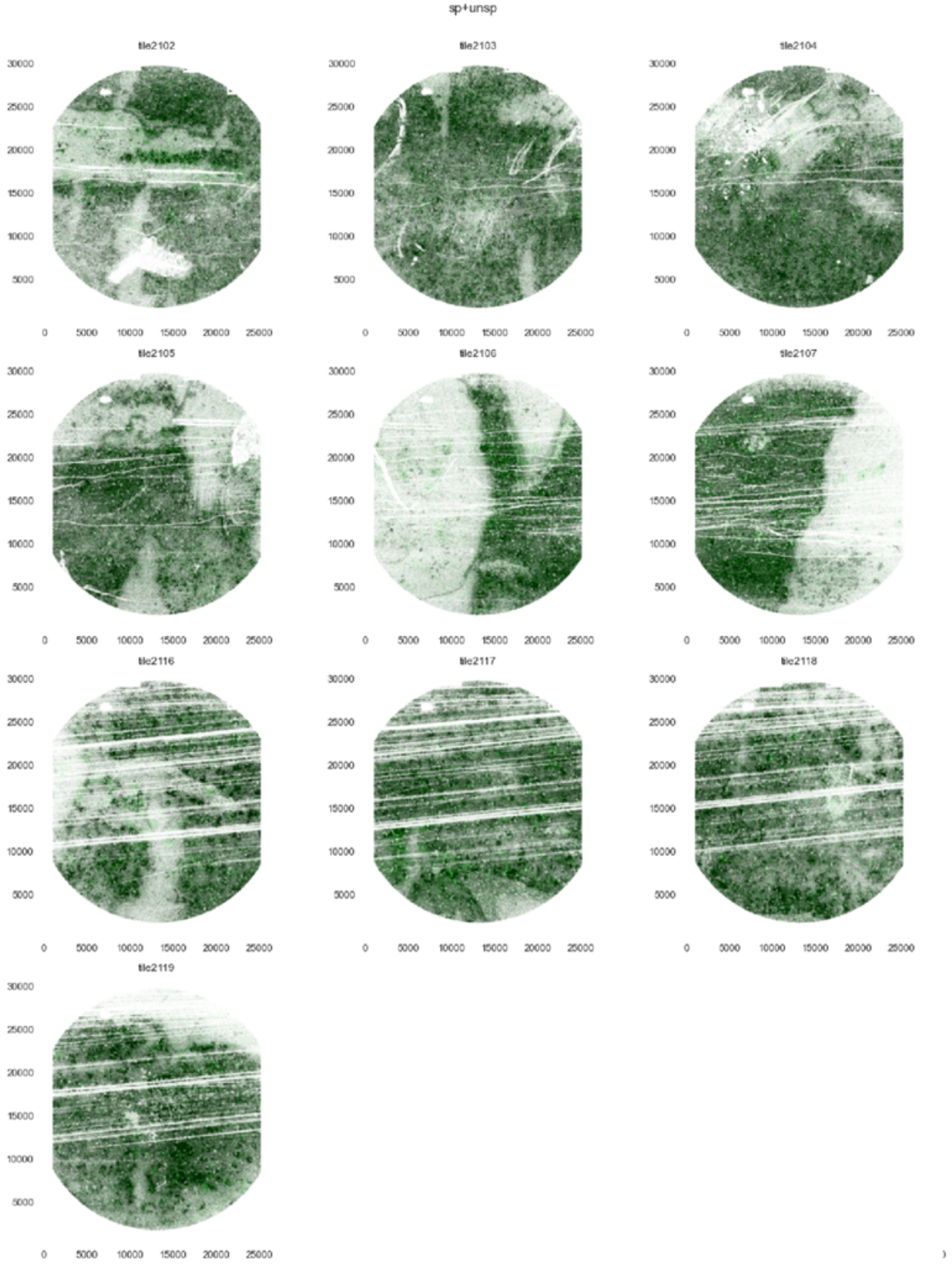

**Supplementary Figure 27. Aligned 4X H&E images for the normal tiles in Seq-Scope.** The 4X H&E images were manually aligned with the transcriptomics data. For each tile, the aligned H&E image (greyscale) is overlaid with the unspliced density (green) and the cell segmentation boundaries (red).

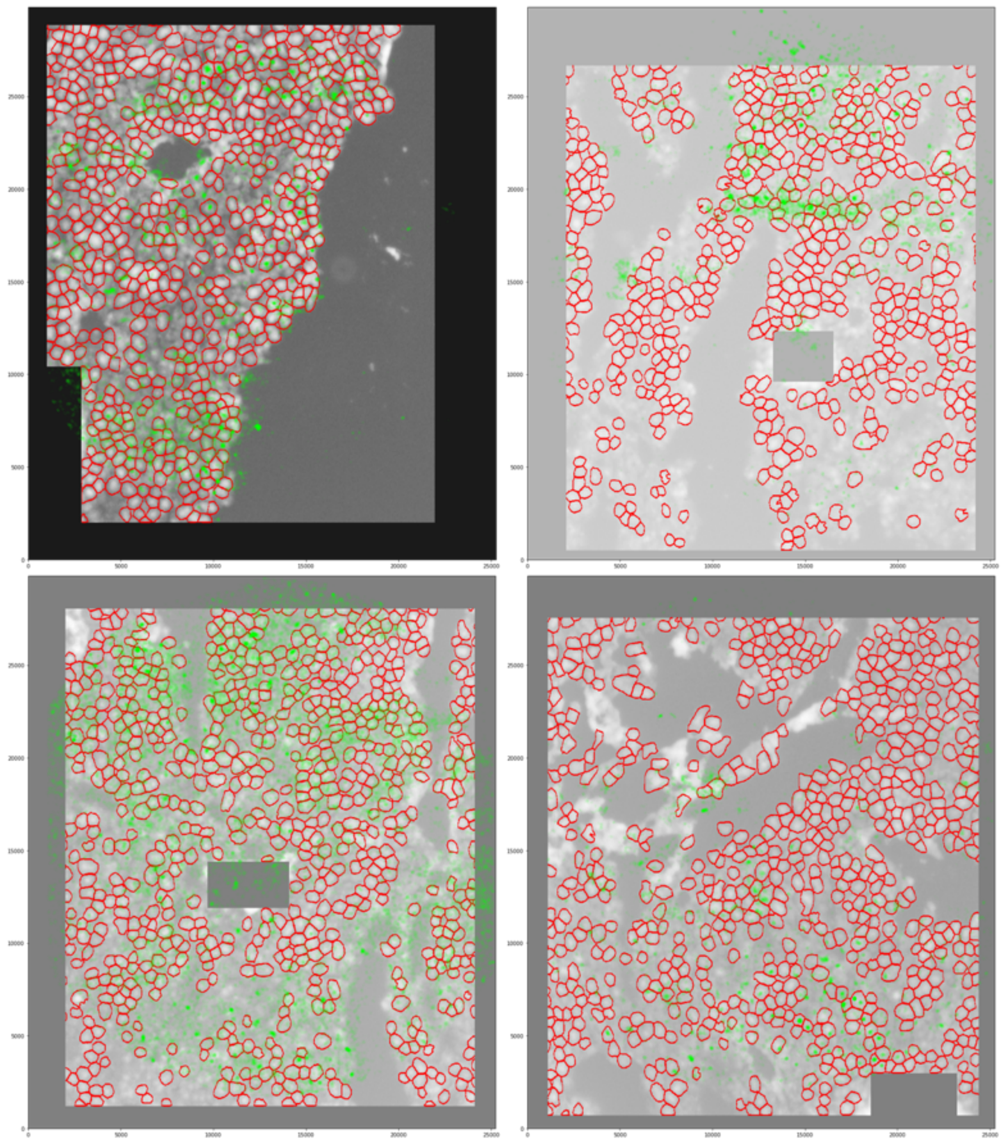

(Fig. S27 con'd)

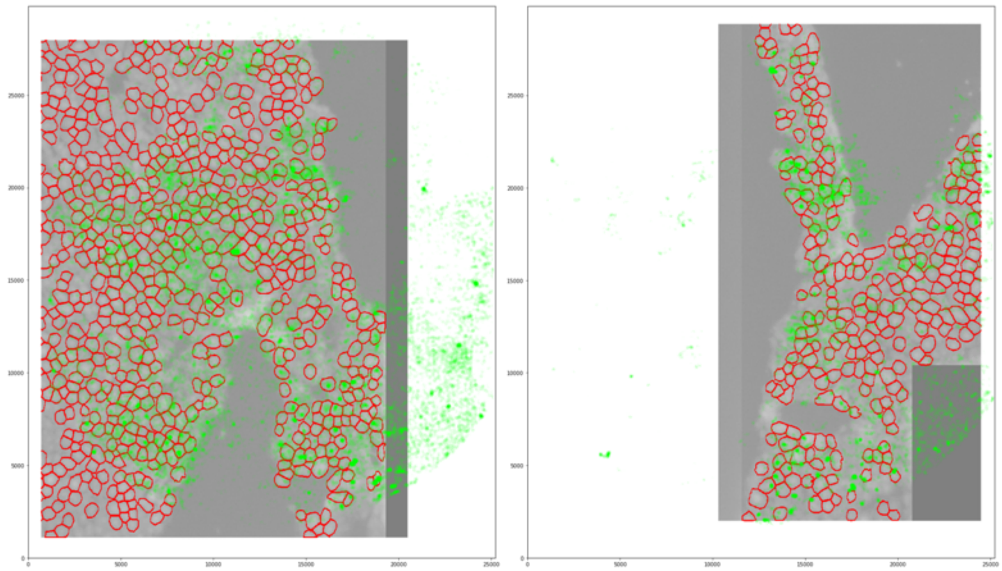

**Supplementary Figure 28. Aligned 4X H&E images for the TD tiles in Seq-Scope.** The 4X H&E images were manually aligned with the transcriptomics data. For each tile, the aligned H&E image (greyscale) is overlaid with the unspliced density (green) and the cell segmentation boundaries (red).

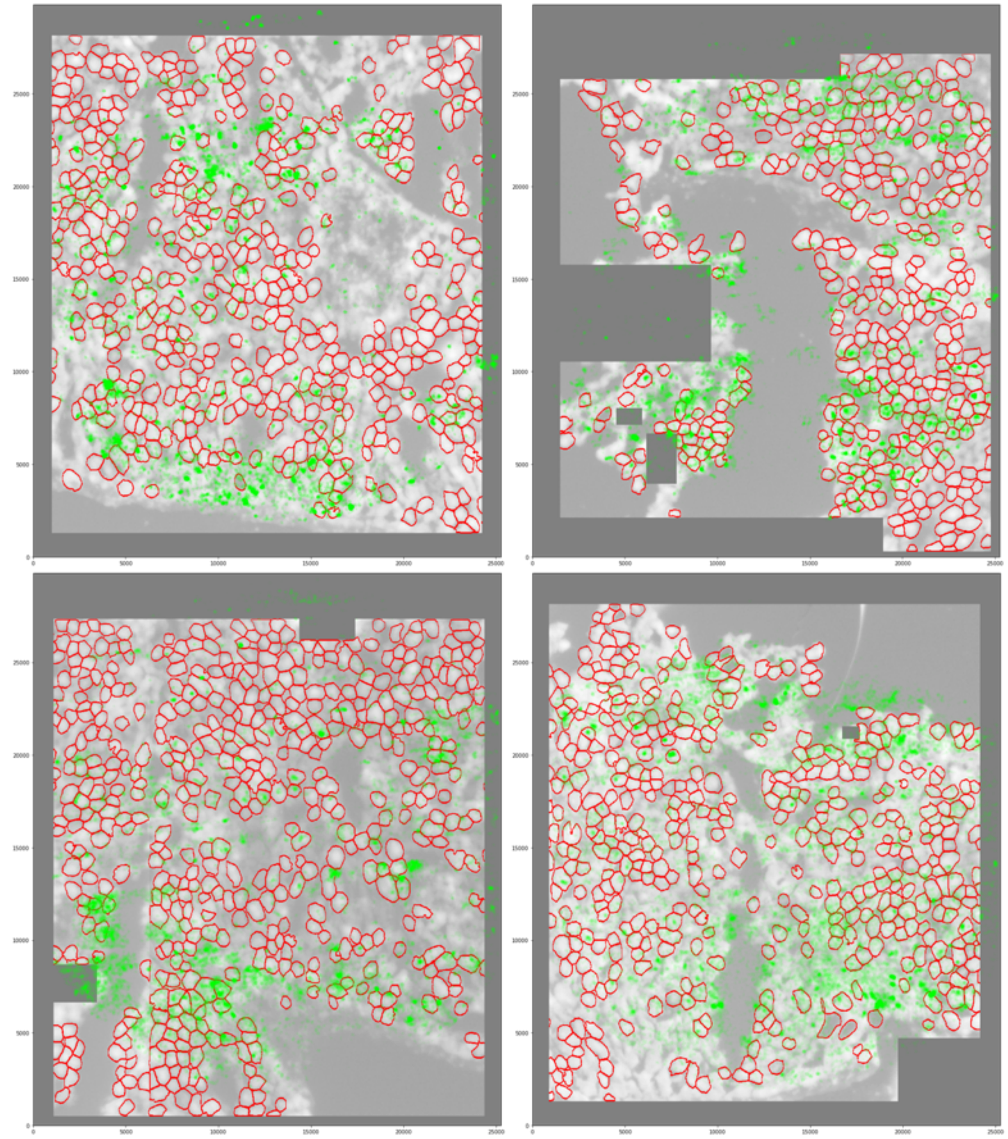

**Supplementary Figure 29. Cropped areas of cells across normal tiles in Seq-Scope.** Areas of cells were cropped from the tiles based on the cell segmentation boundaries transferred from the aligned H&E images to the tiles. The cell segmentation boundaries (red) are overlayed with: unspliced expression density (green), *Alb* expression (blue), MT genes expression (purple), nuclear gene expression (the 10 nuclear genes presented in Seq-Scope study, black), nuclear center (red crosses).

Tile 2102:

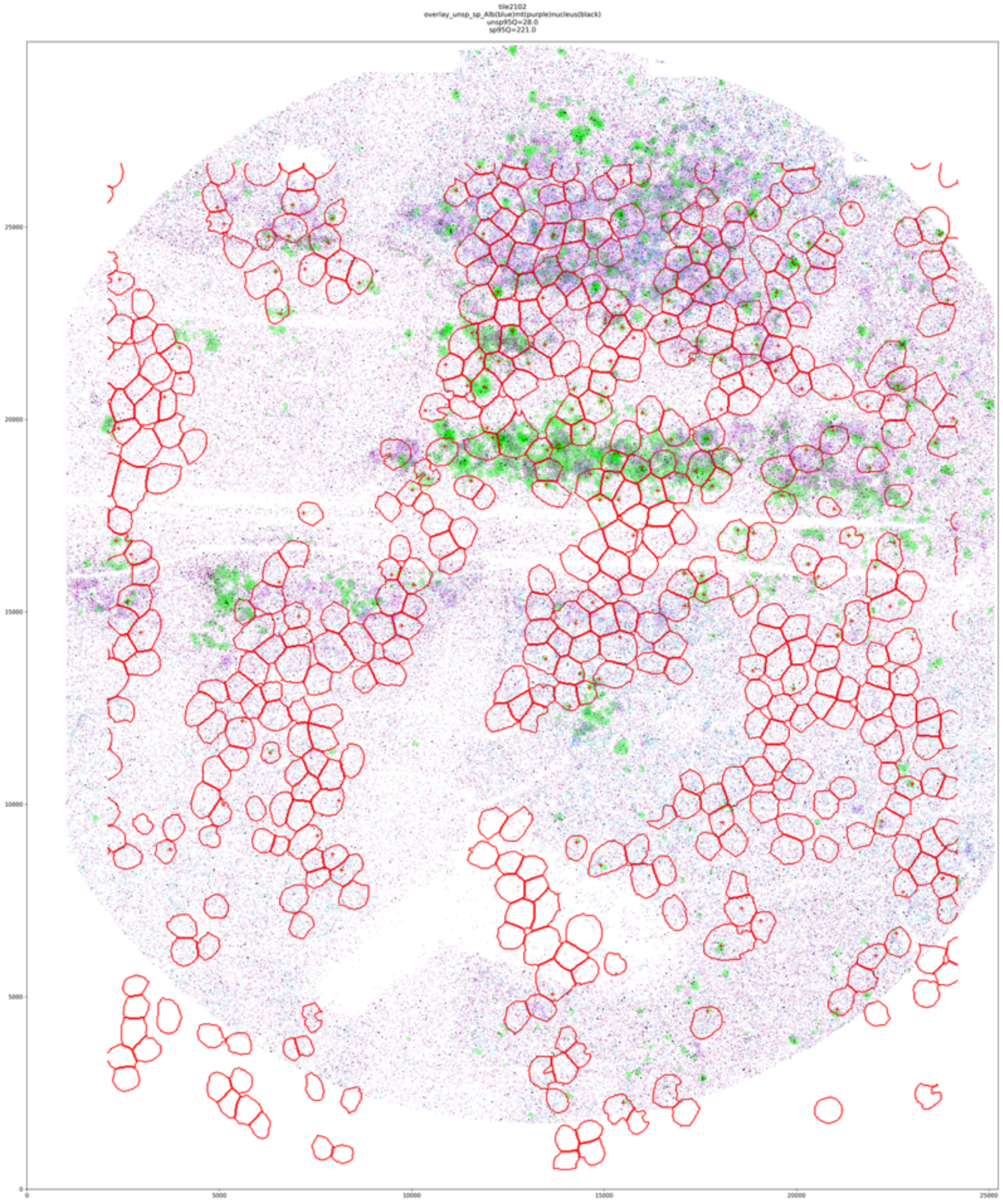

(Fig. S29 con'd)

Tile 2103:

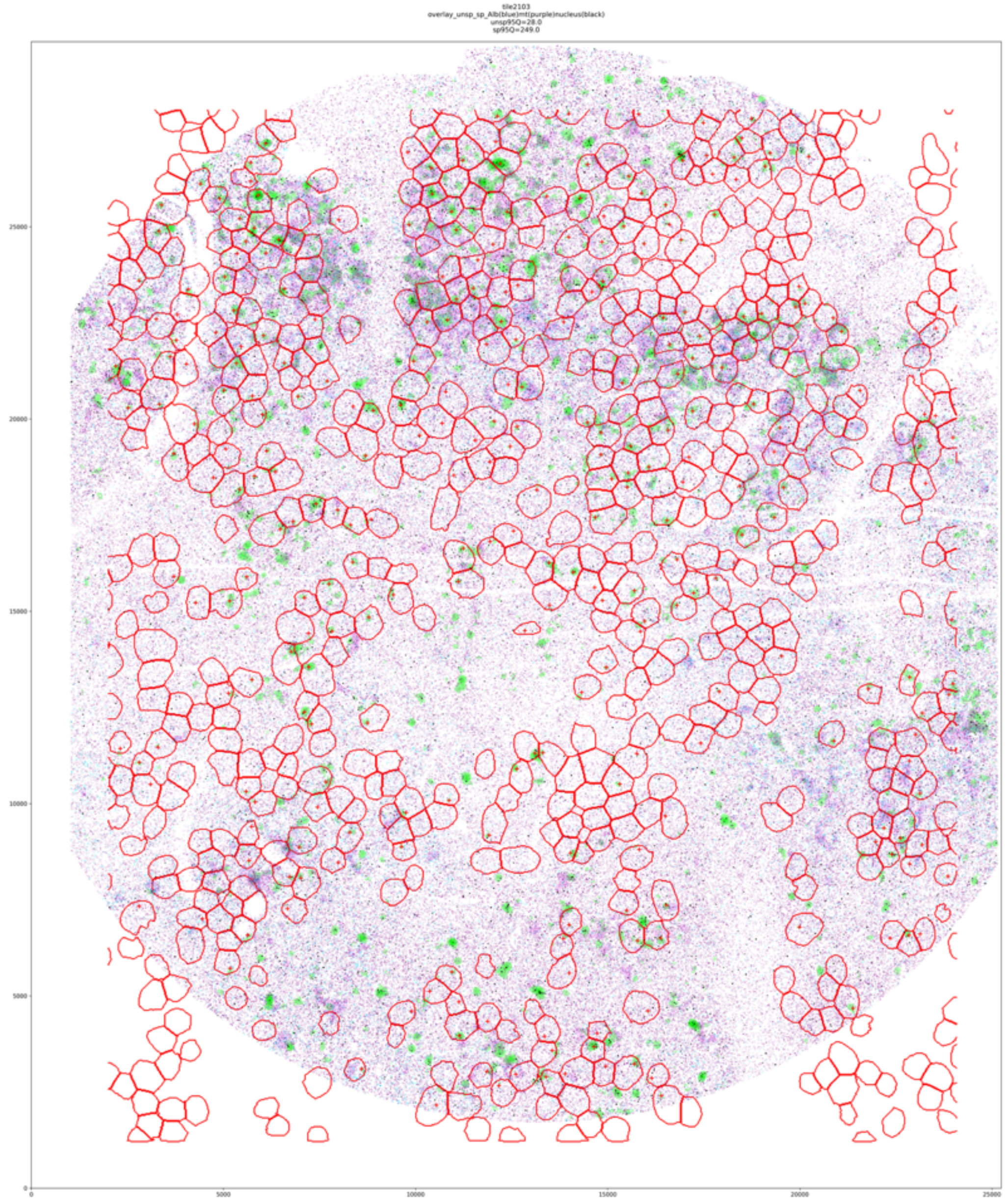

(Fig. 29 con'd)

Tile 2104:

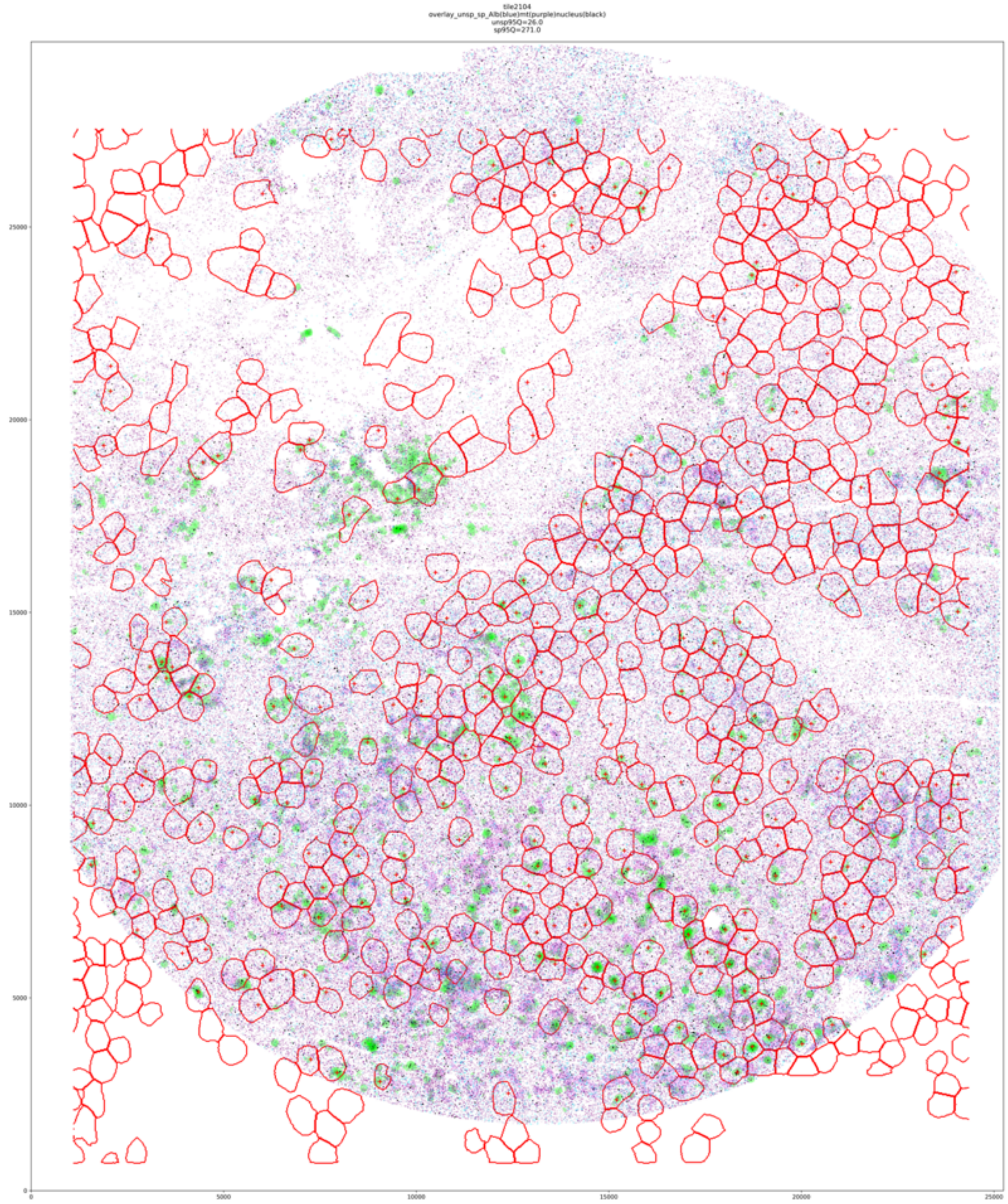

(Fig. S29 con’d)

Tile 2105:

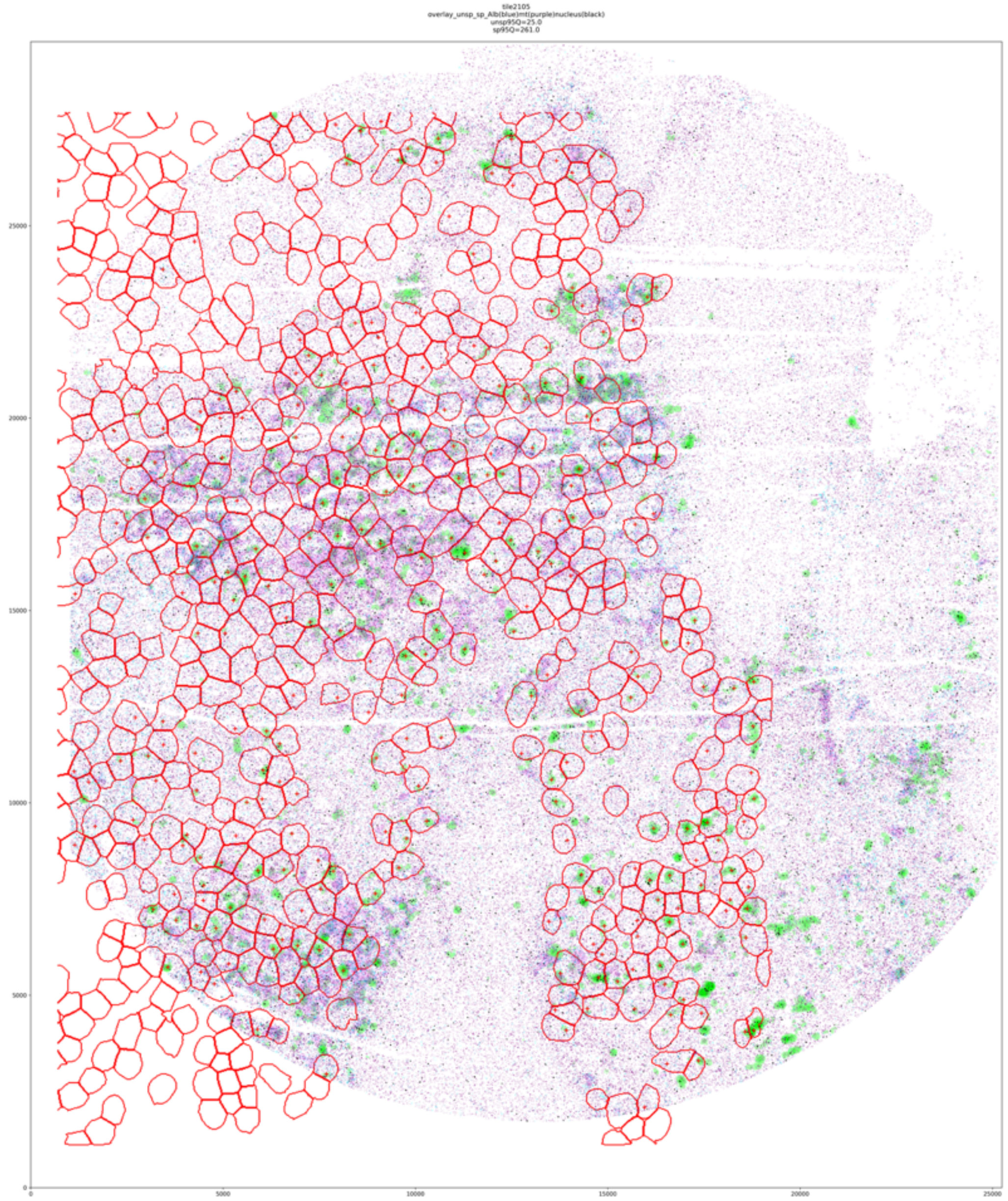

(Fig. S29 con’d)

Tile 2106:

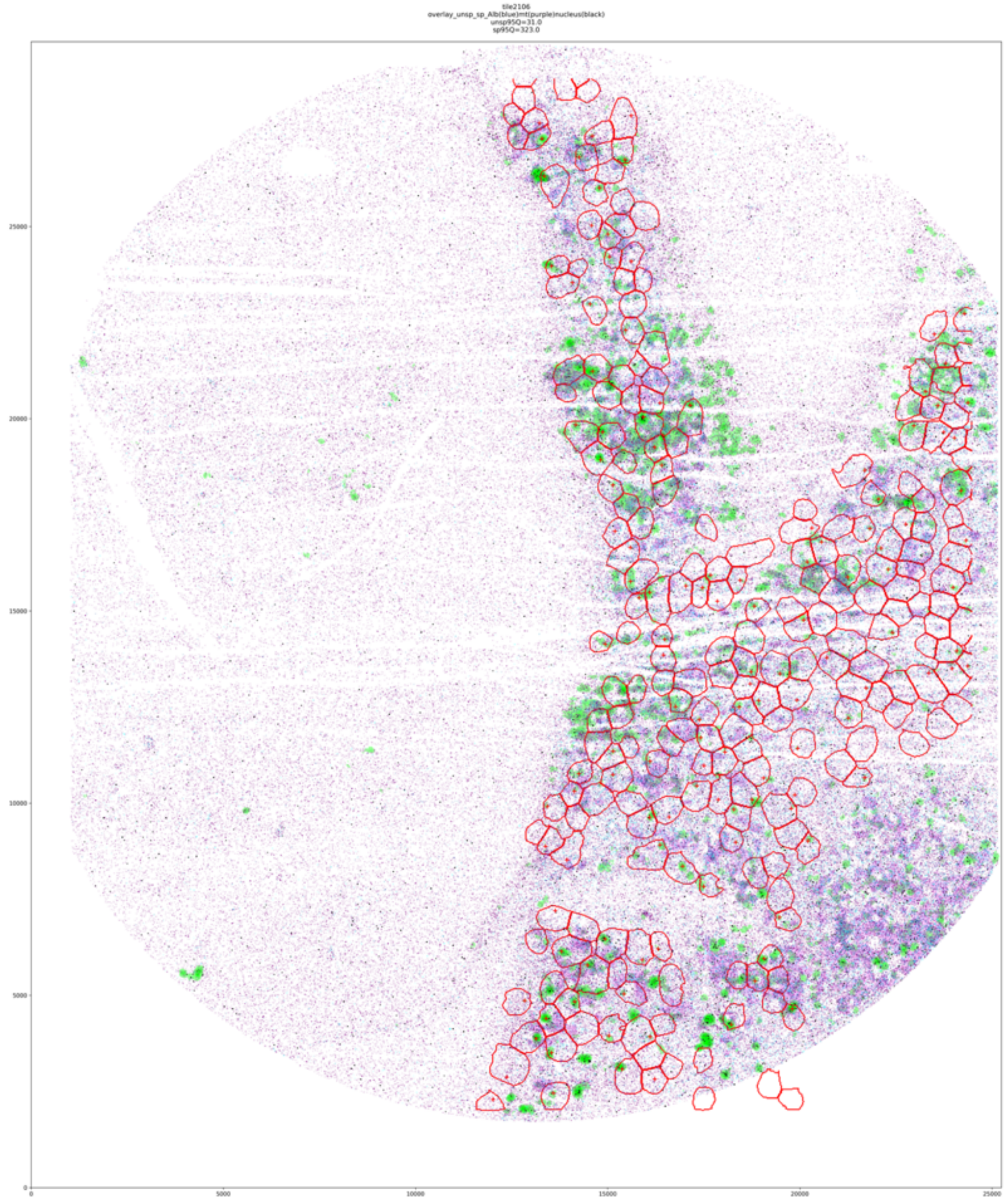

(Fig. S29 con’d)

Tile 2107:

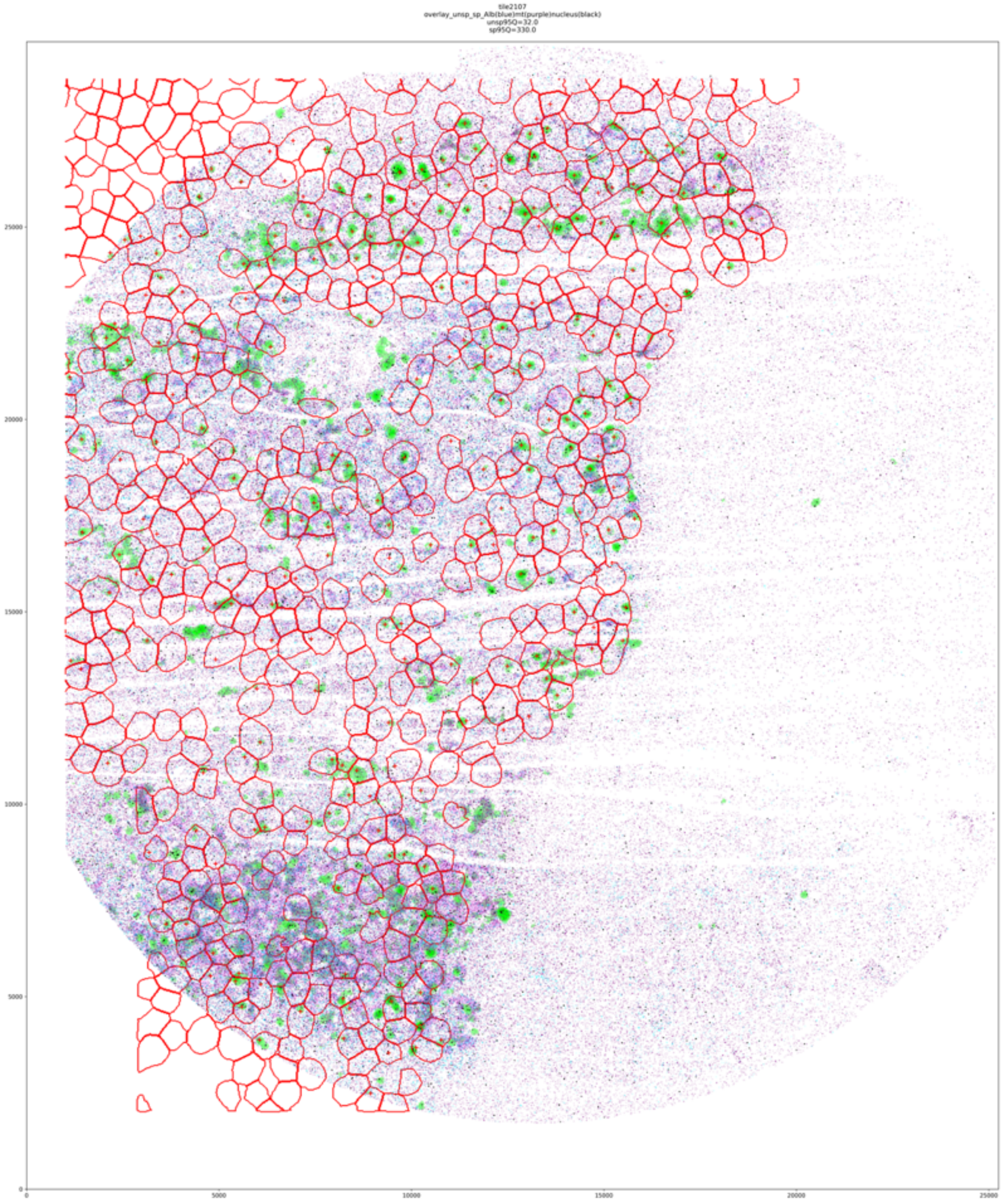

**Supplementary Figure 30. Cropped areas of cells across TD tiles in Seq-Scope.** Areas of cells were cropped from the tiles based on the cell segmentation boundaries transferred from the aligned H&E images to the tiles. The cell segmentation boundaries (red) are overlayed with: unspliced expression density (green), *Alb* expression (blue), MT genes expression (purple), nuclear gene expression (the 10 nuclear genes presented in Seq-Scope study, black), nuclear center (red crosses).

Tile 2116:

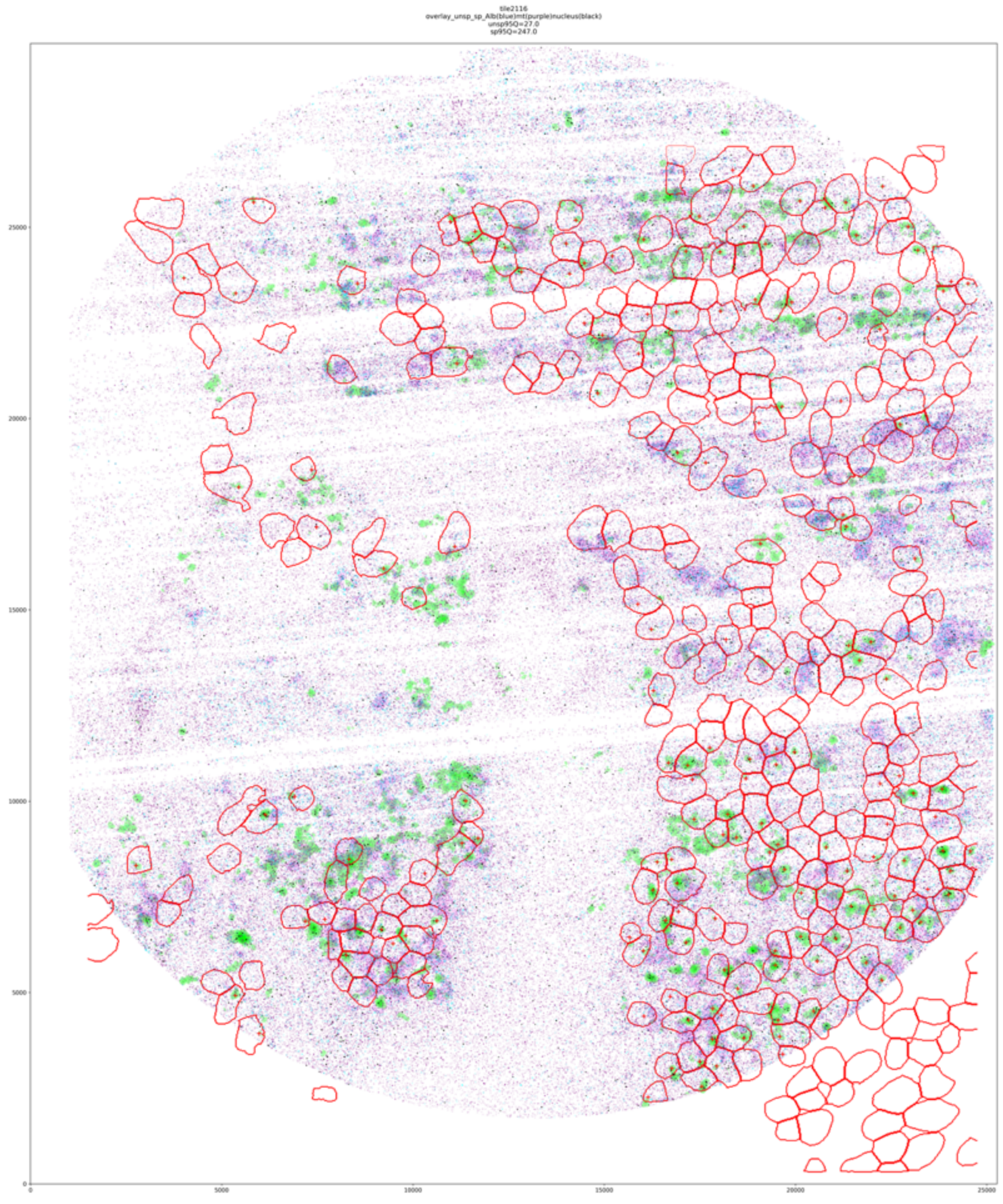

(Fig. S30 con’d)

Tile 2117:

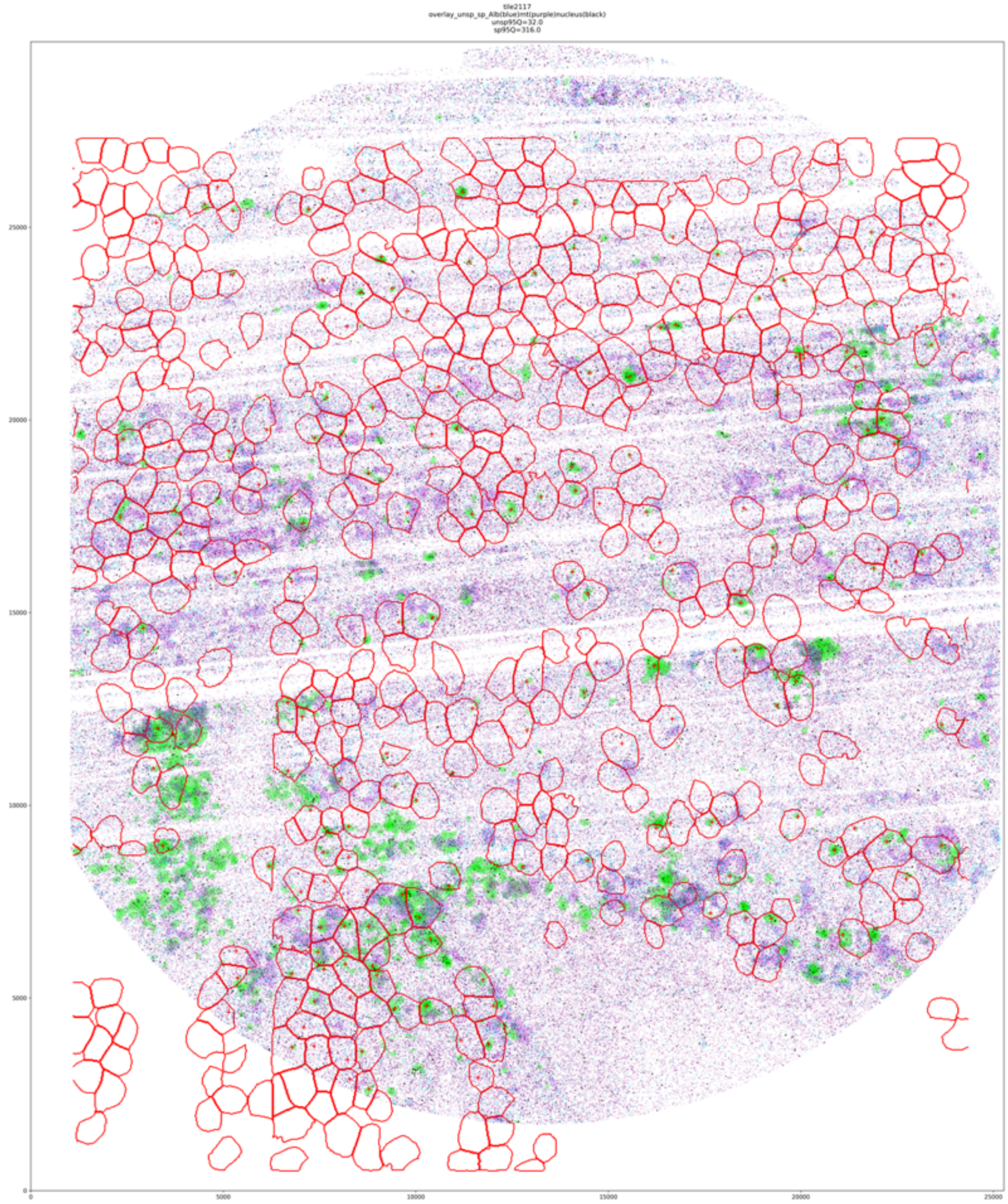

(Fig. S30 con’d)

Tile 2118:

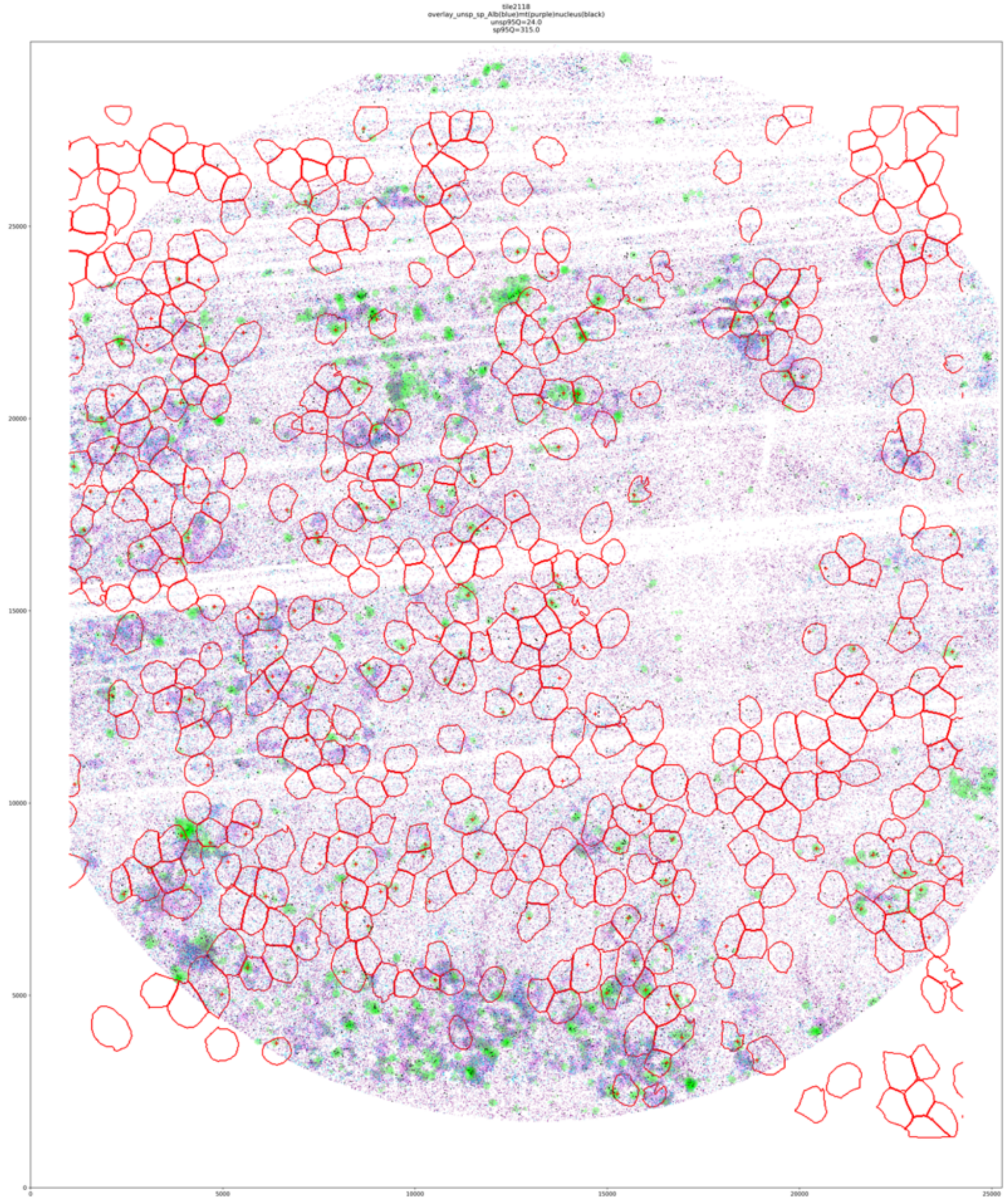

(Fig. S30 con’d)

Tile 2119:

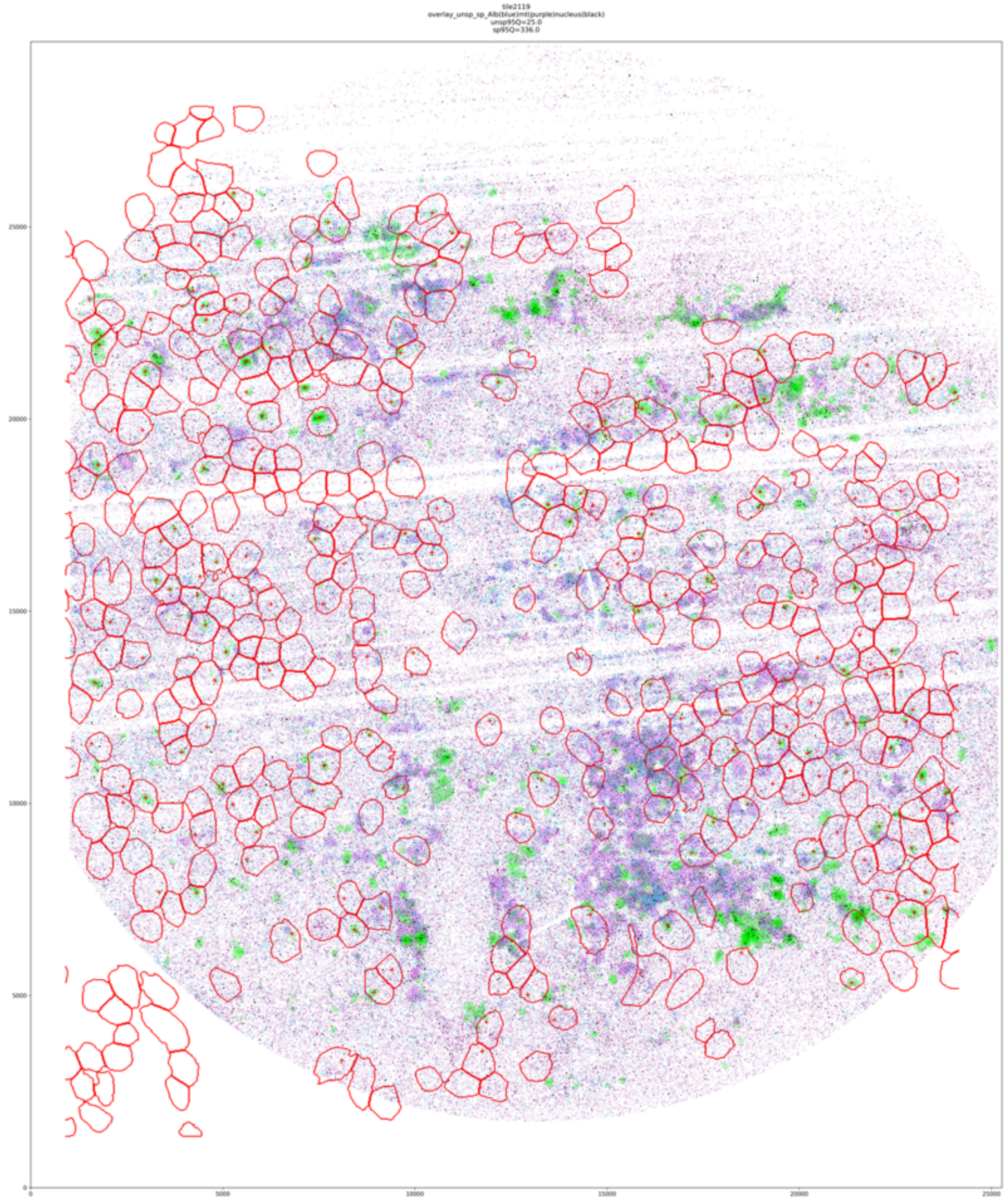

**Supplementary Figure 31. Number of genes and cells in the Seq-Scope mouse liver data. a.** Bar plot shows the number of genes analyzed by ELLA across cell types. **b.** Bar plot shows the number of cells available across cell types. **c.** Bar plot shows the number of genes detected by ELLA across cell types.

**a.**

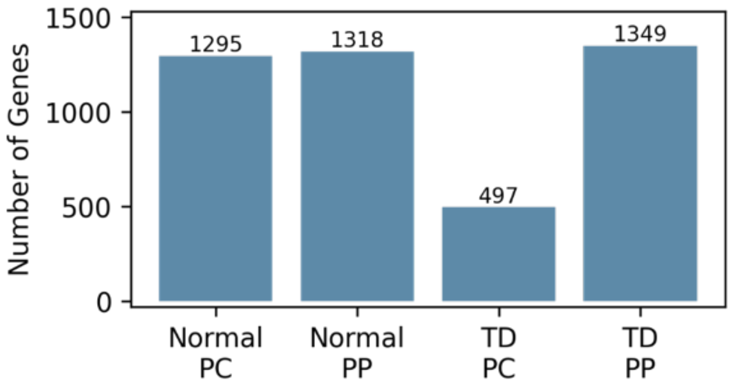

**b.**

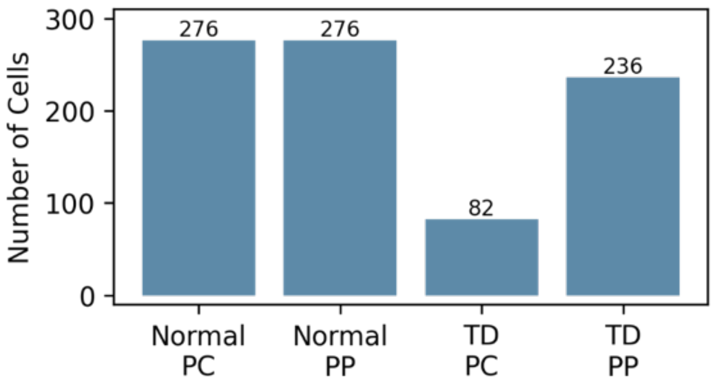

**c.**

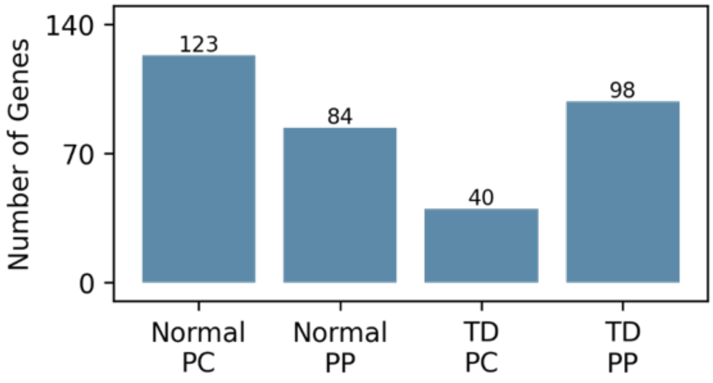

**Supplementary Figure 32. Cell type annotations across normal and TD tiles in Seq-Scope.** Nuclear centers of the cells with annotated cell types are plotted (PC red; PP green; NPC blue) across normal and TD tiles.

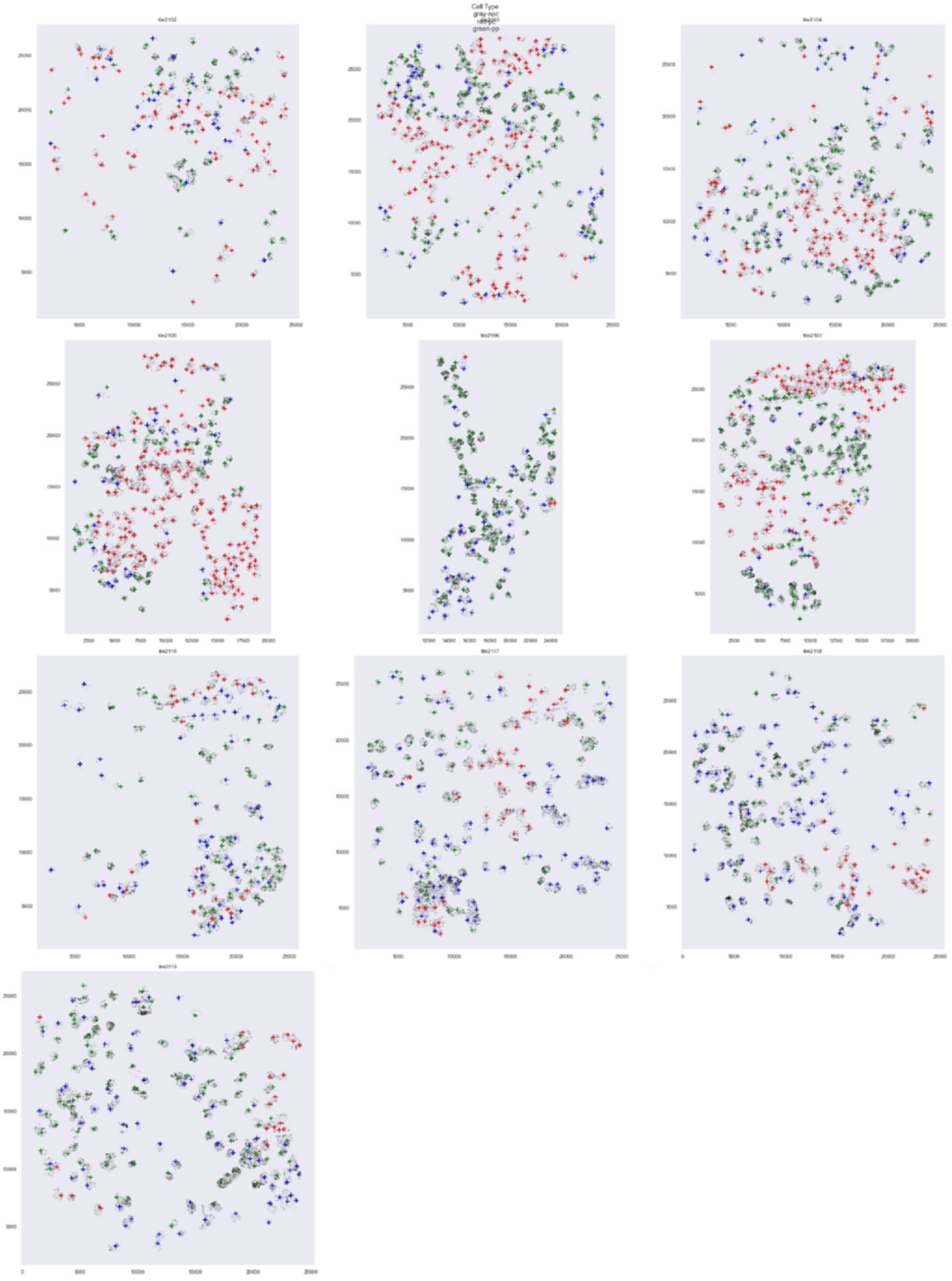

**Supplementary Figure 33. Estimated expression pattern for genes identified by ELLA in individual cell types in Seq-Scope. a.** The number and proportion of genes across five pattern clusters in each cell type. **b.** The estimated expression intensities for genes across clusters in each cell type. **c.** The estimated pattern score for genes across clusters in each cell type.

**a.**

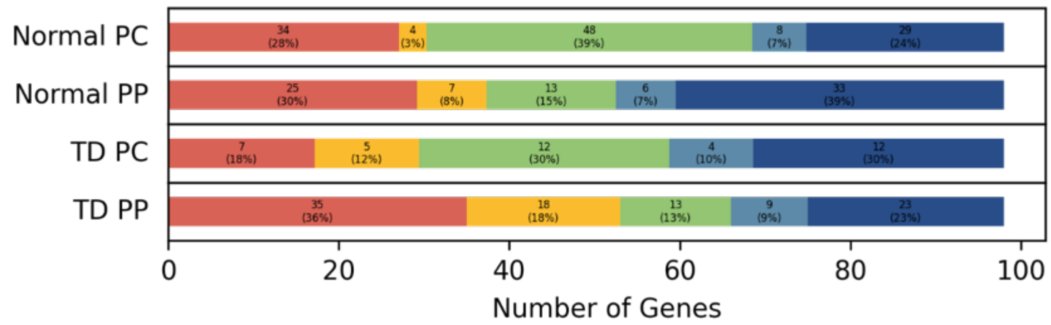

**b.**

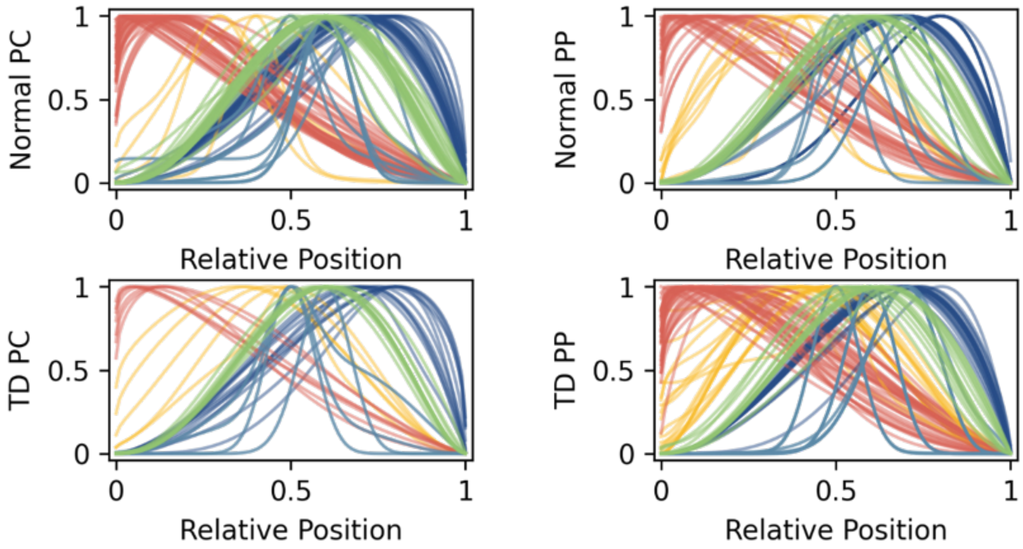

**c.**

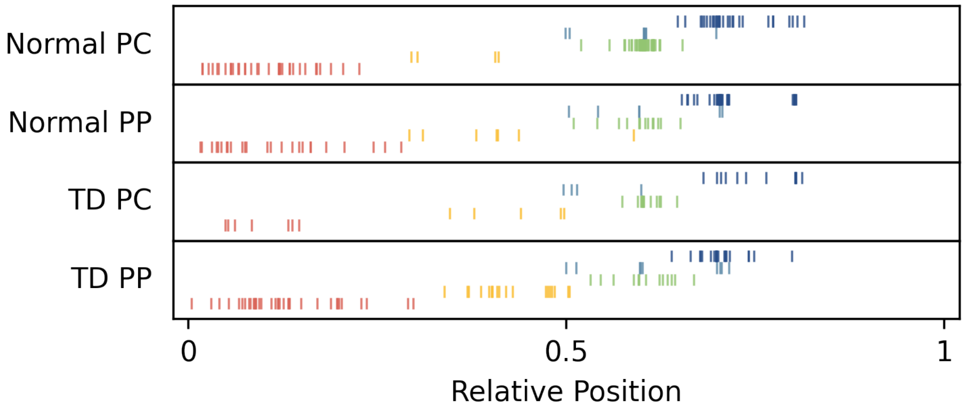

**Supplementary Figure 34. Estimated expression patterns for the 34 nuclear genes identified by ELLA in Seq-Scope normal PC cells.** The estimated standardized subcellular expression intensity functions are plotted.

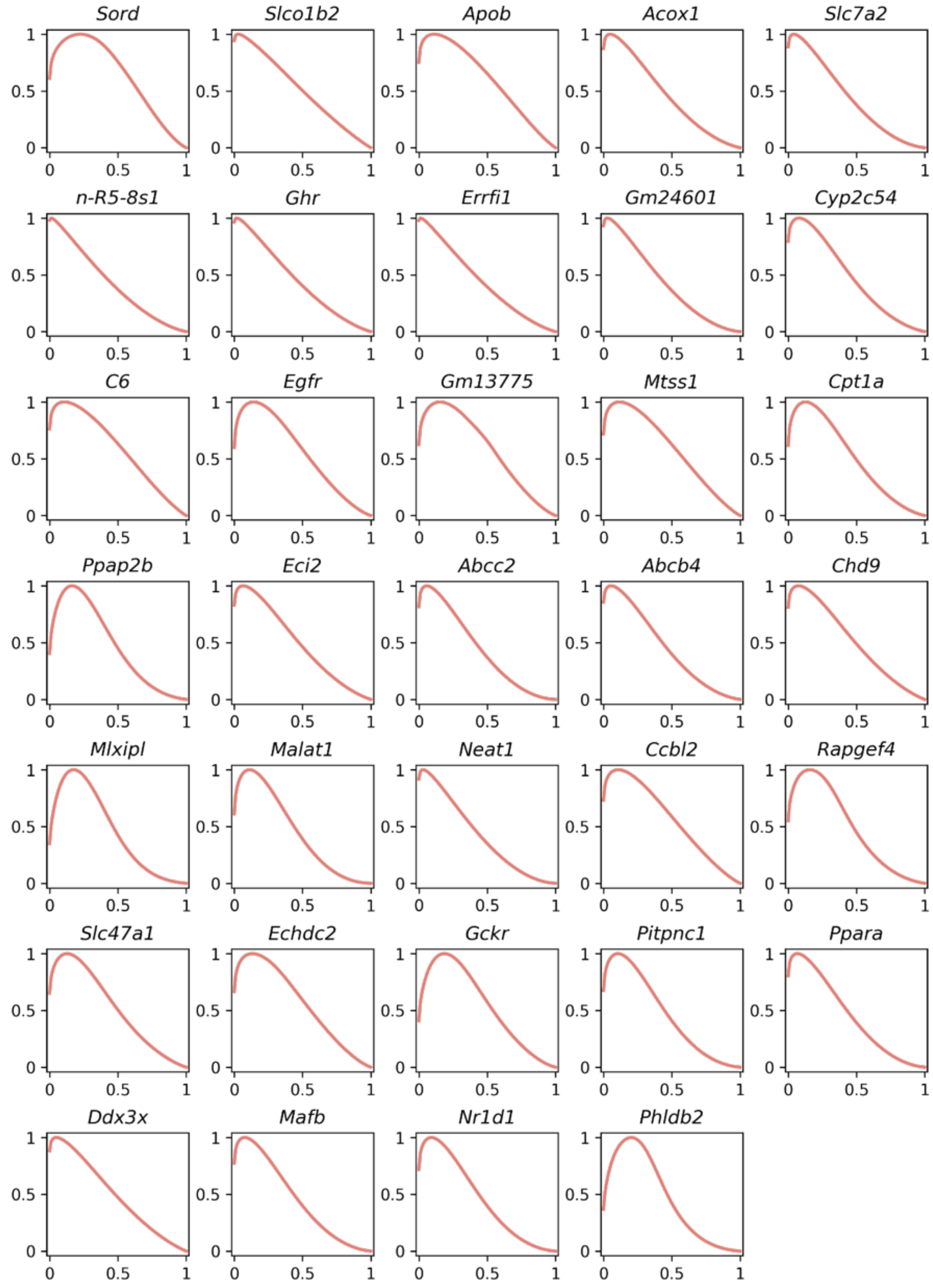

**Supplementary Figure 35. Estimated expression patterns for the mitochondrial genes and cell type marker genes in the Seq-Scope normal PC cell type. a.** The estimated standardized subcellular expression intensity functions of the 9 mitochondrial genes are plotted (cluster 3 genes in light green; cluster 5 genes in dark blue; nonsignificant genes in light grey). **b.** The estimated standardized subcellular expression intensity functions of the 3 cell type marker genes are plotted (cluster 3 genes in light green; cluster 5 genes in dark blue; nonsignificant genes in light grey).

**a.**

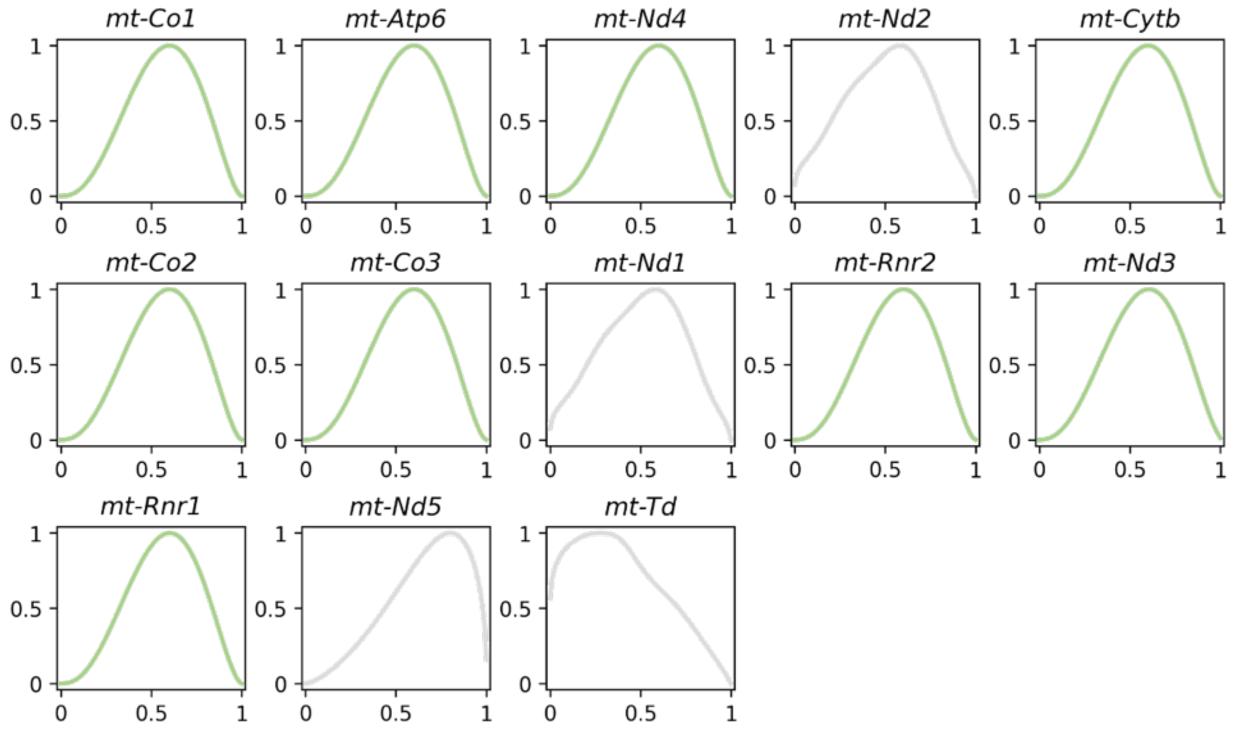

**b.**

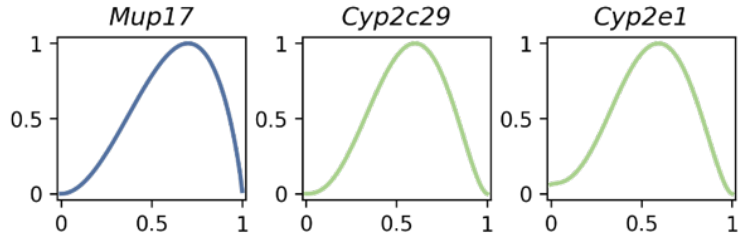

**Supplementary Figure 36. Nucleic acid staining image of slice E1S3 in Stereo-seq.** The full tissue region was split into eight subregions for Cellpose cell segmentation.

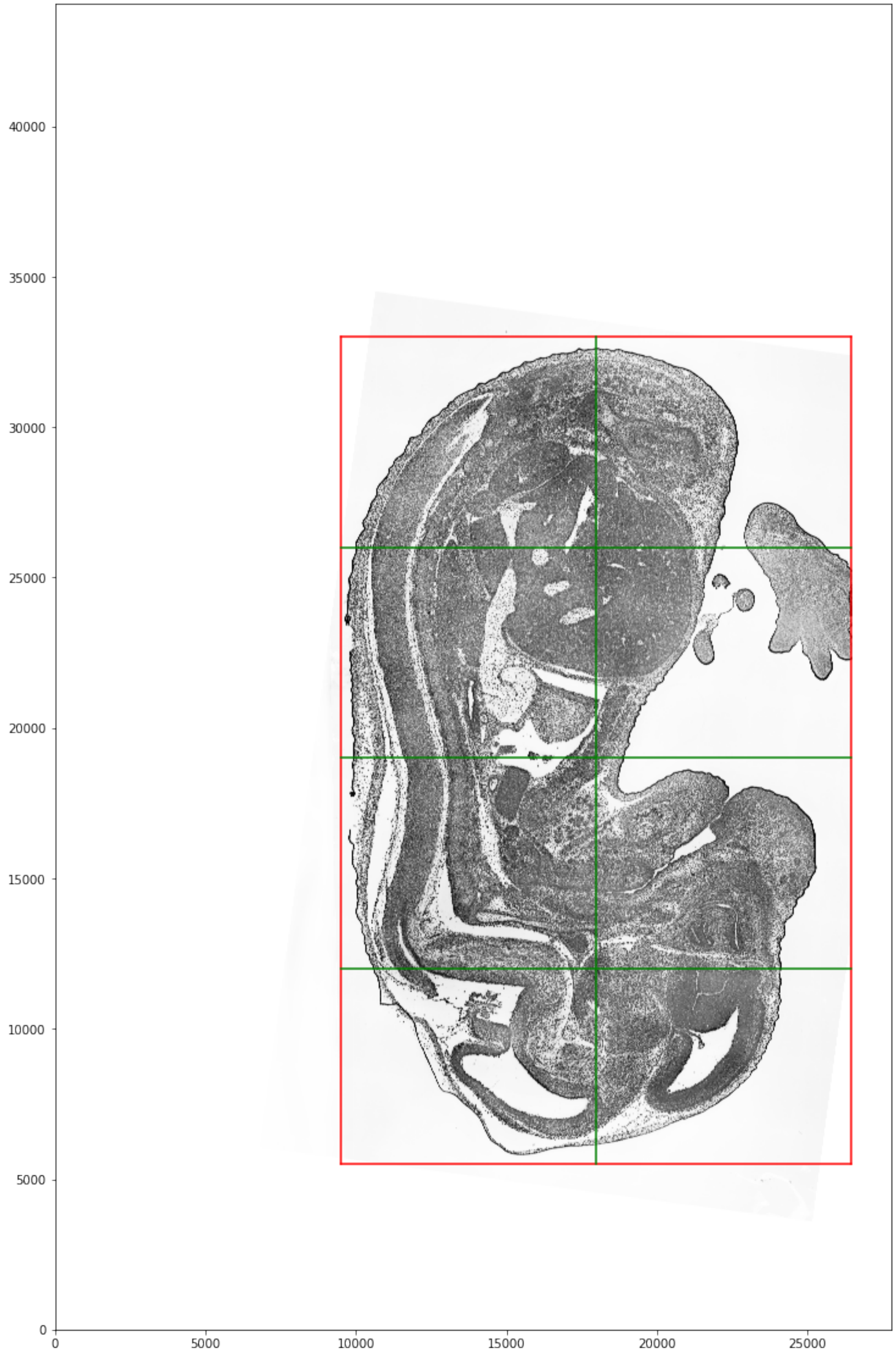

**Supplementary Figure 37. Cell segmentations from Cellpose in Stereo-seq.** Cell segmentation boundaries (blue) are overlayed with the nucleic acid staining image (greyscale), nuclear centers (red), and total gene expressions (pink) across subregions.

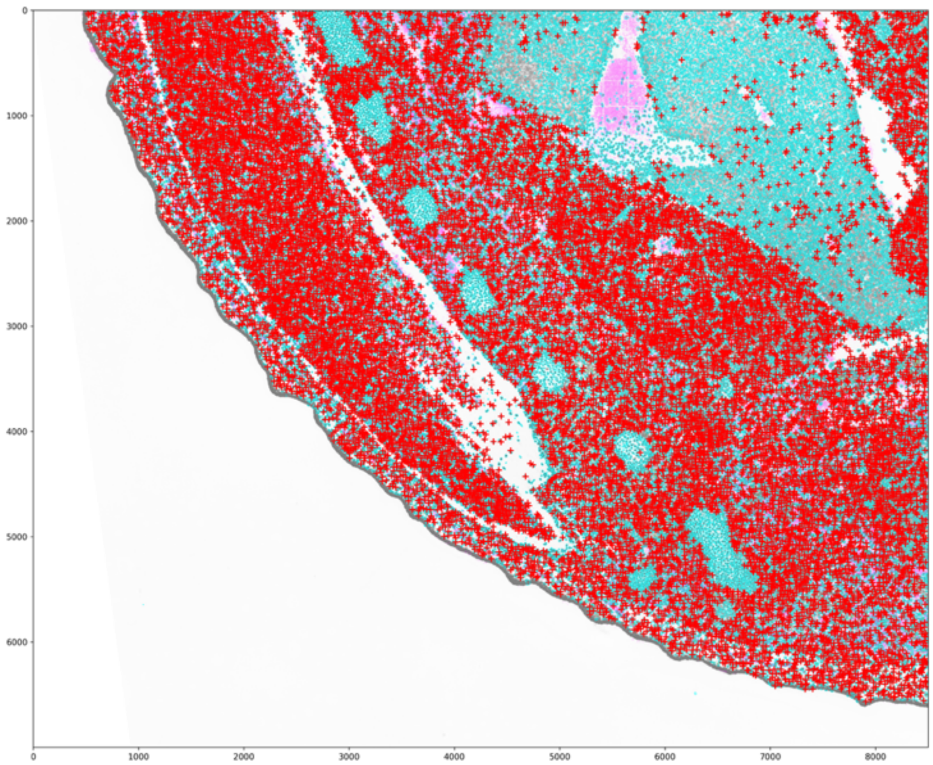

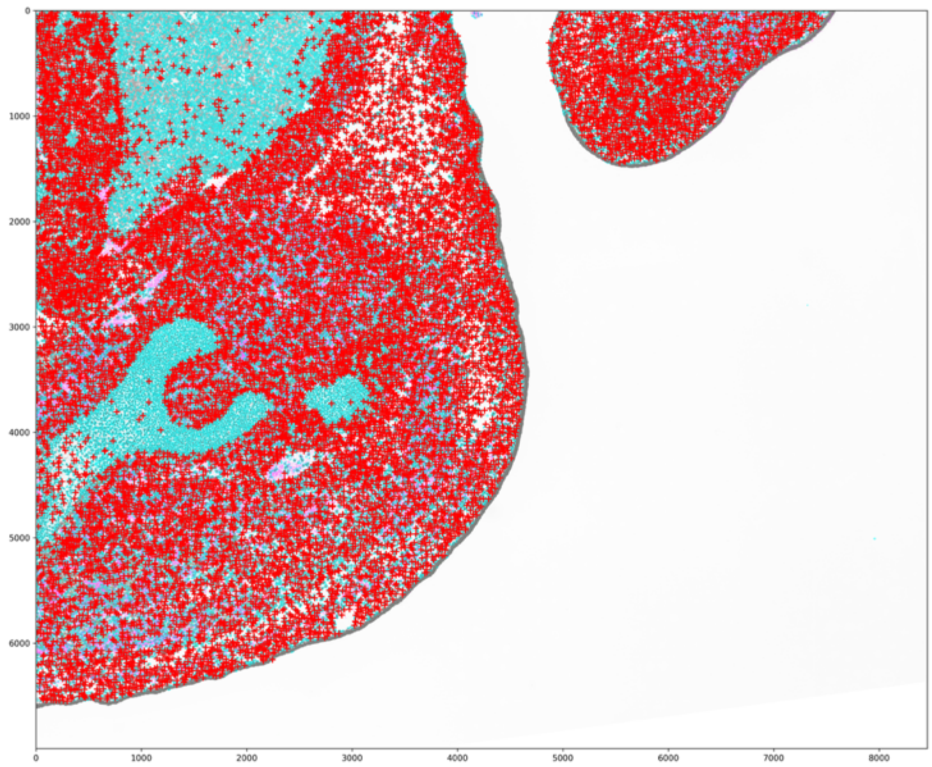

(Fig. S37 con'd)

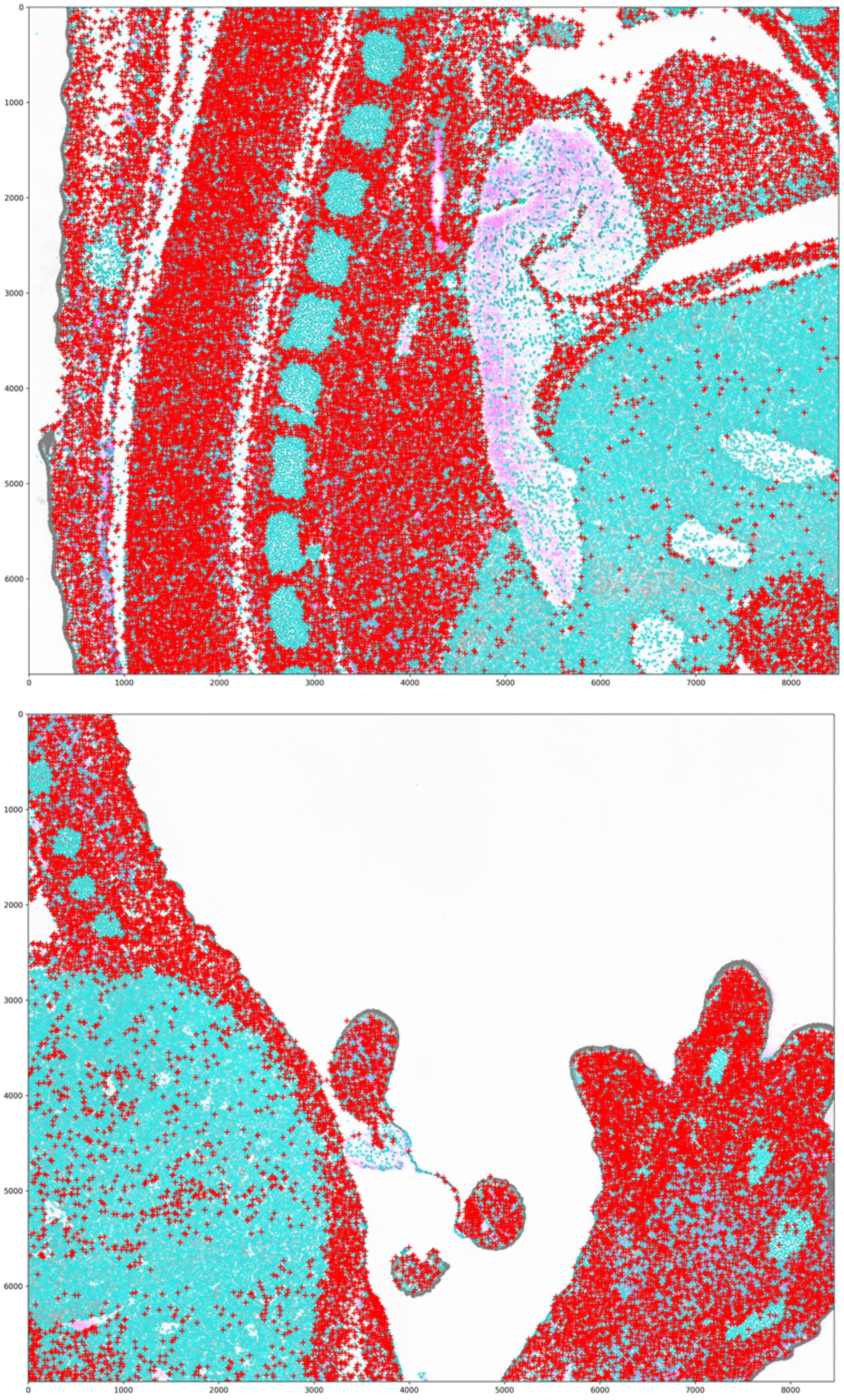
0
1000
2000
3000
4000
5000
6000
0
1000
2000
3000
4000
5000
6000
7000
8000
0
1000
2000
3000
4000
5000
6000
0
1000
2000
3000
4000
5000
6000
7000
8000

(Fig. S37 con'd)

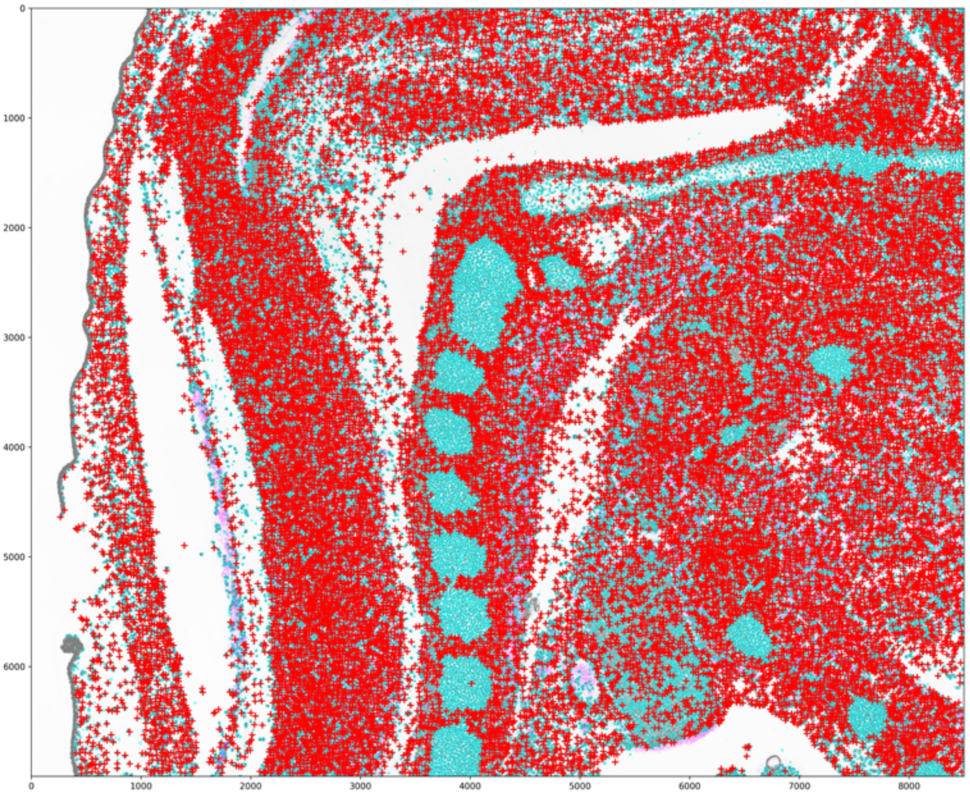
0
1000
2000
3000
4000
5000
6000
0
1000
2000
3000
4000
5000
6000
7000
8000

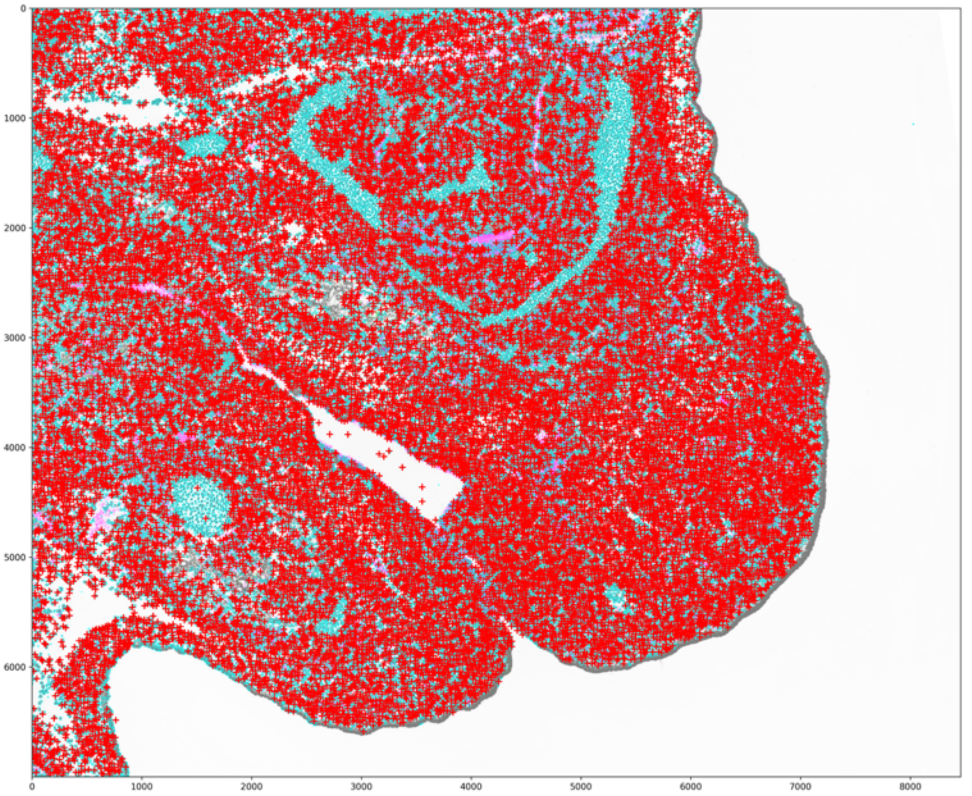
0
1000
2000
3000
4000
5000
6000
0
1000
2000
3000
4000
5000
6000
7000
8000

(Fig. S37 con’d)

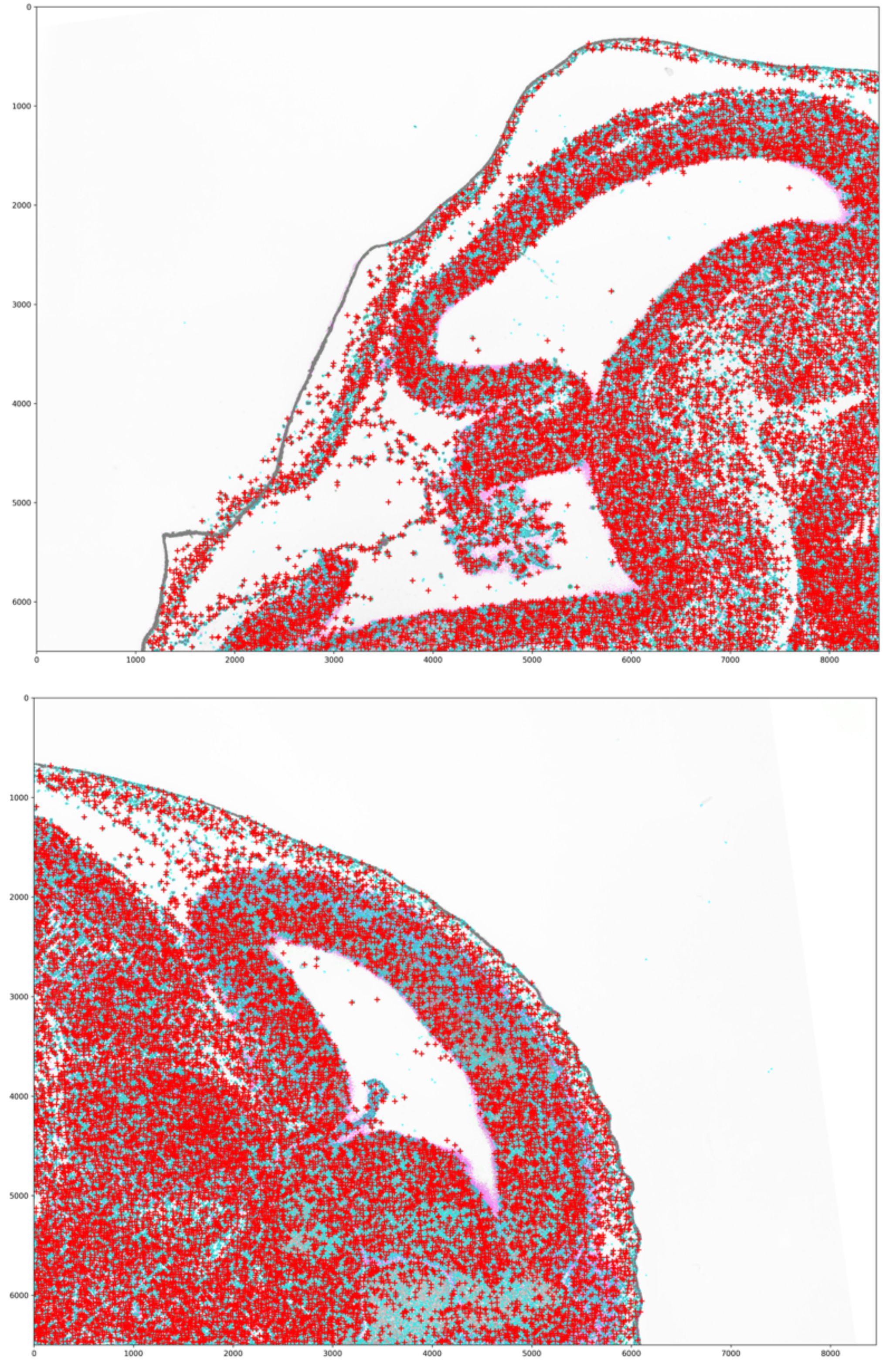
0
1000
2000
3000
4000
5000
6000
0
1000
2000
3000
4000
5000
6000
7000
8000
0
1000
2000
3000
4000
5000
6000
0
1000
2000
3000
4000
5000
6000
7000
8000

**Supplementary Figure 38. Cell type marker gene expression heatmap in Stereo-seq. a.** Heatmap with 100 randomly selected cells for the single cell (total) counts of the cell type marker genes of each of the 25 cell types; **b.** Heatmaps of the counts in (a) that are standardized across genes and cells.

**a.**

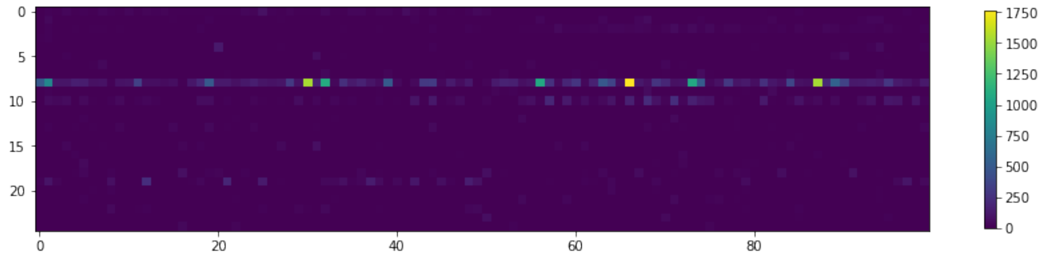

**b.**

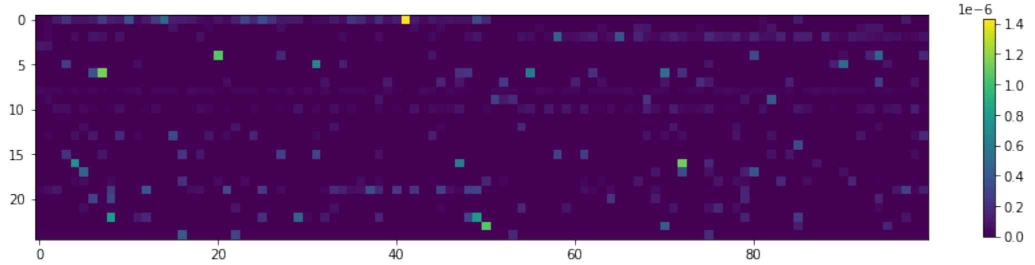

**Supplementary Figure 39. Number of genes and cells in the Stereo-seq mouse embryo data.** **a.** Bar plot shows the number of genes analyzed by ELLA across cell types. **b.** Bar plot shows the number of cells available across cell types. **c.** Bar plot shows the number of genes detected by ELLA across cell types.

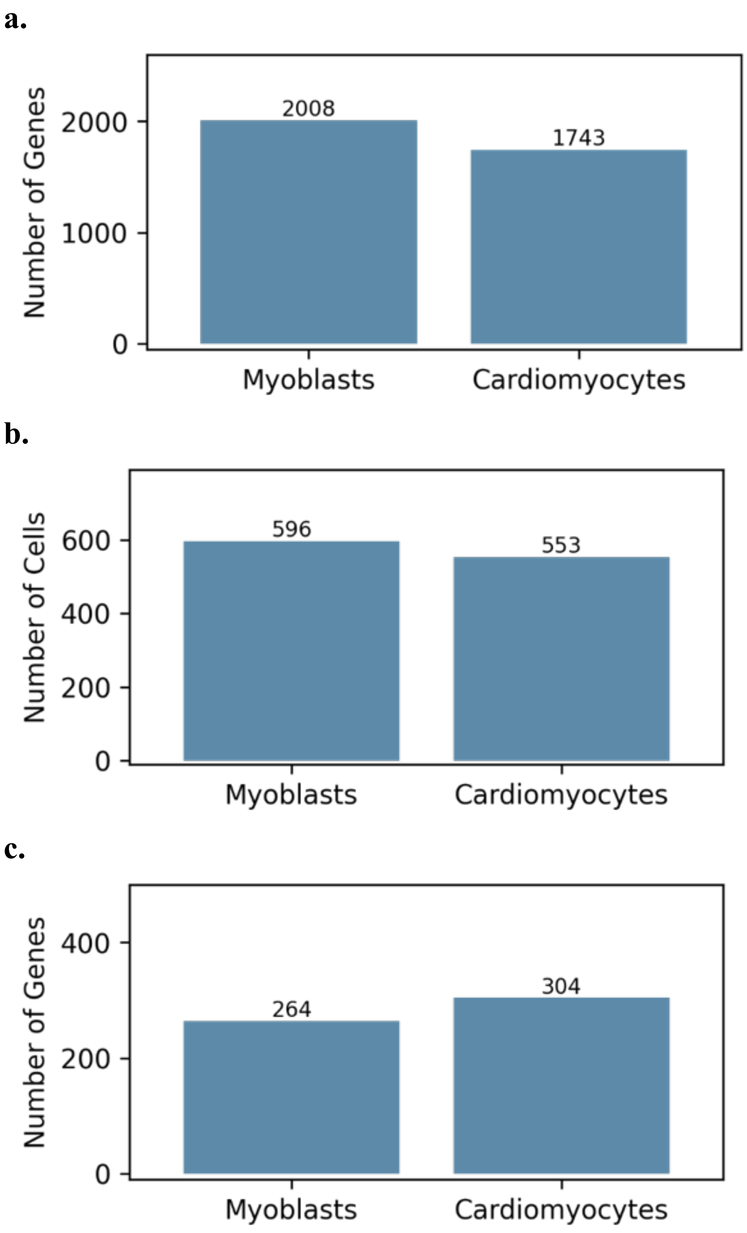

**Supplementary Figure 40. Histograms display the estimated ReadZS scores across 25 cell types in Stereo-seq.** The ReadZS scores of the positive strand genes are plotted in light green and the ReadZS scores of the negative strand genes are plotted in light blue.

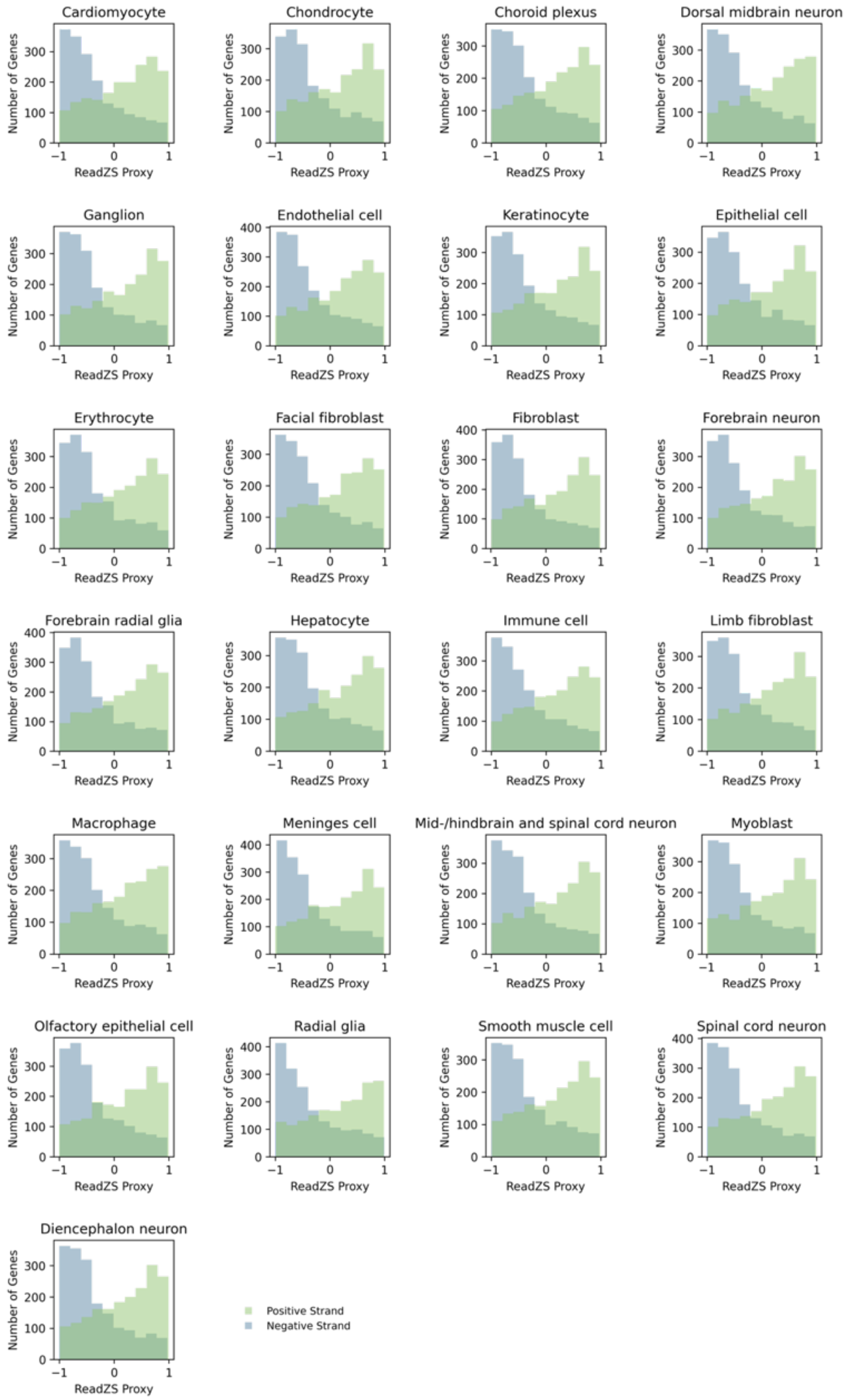

**Supplementary Figure 41. Boxplots show the estimated 3’ UTR lengths across ELLA pattern clusters of the 19 genes displaying significant variation across pattern clusters.** P values were obtained by examining the variation across pattern clusters based on Kruskal-Wallis H tests.

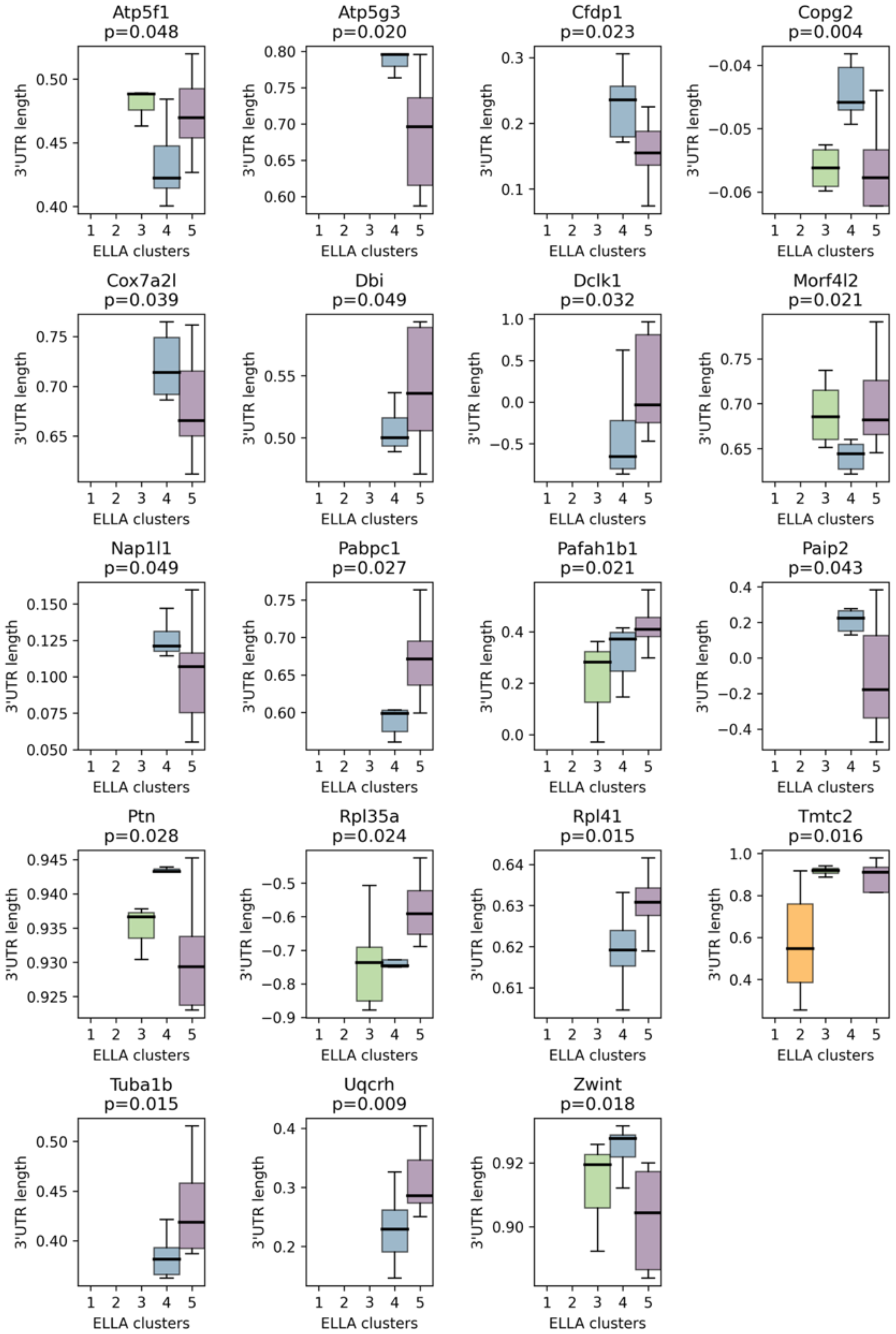

**Supplementary Figure 42. Scatter plots of estimated 3'UTR length and ELLA pattern strength for 21 genes of which 3'UTR length is correlated with ELLA pattern strength in Stereo-seq.** For each gene, a P value was obtained by examining the Pearson correlation between the estimated 3'UTR lengths (or the ReadZS scores) and the ELLA pattern strengths across cell types.

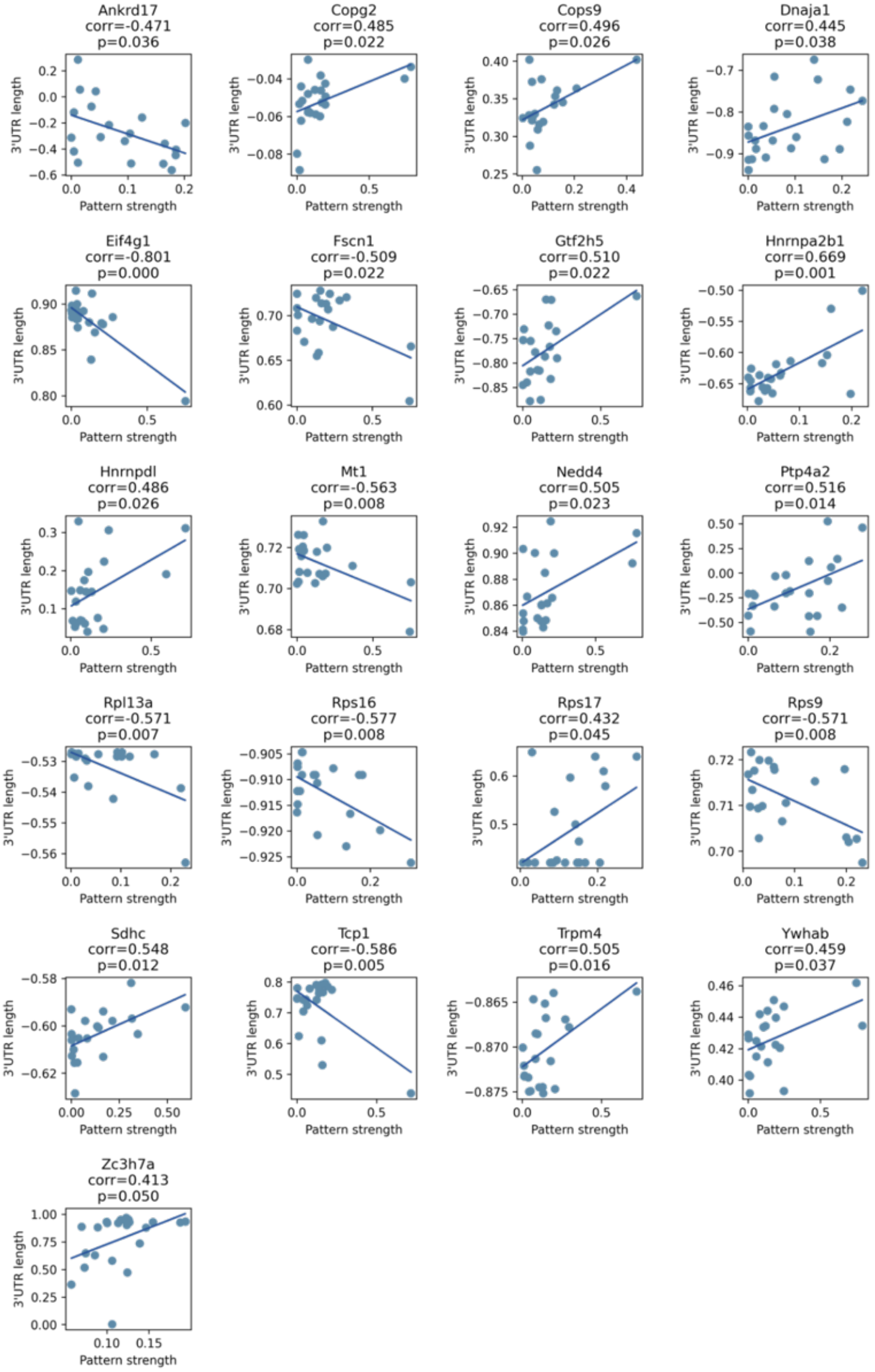

**Supplementary Figure 43. Scatter plots of estimated 3’UTR length and ELLA pattern score for 18 genes of which 3’UTR length is correlated with ELLA pattern score in Stereo-seq.** For each gene, a P value was obtained by examining the Pearson correlation between the estimated 3’UTR lengths (or the ReadZS scores) and the ELLA pattern scores across cell types.

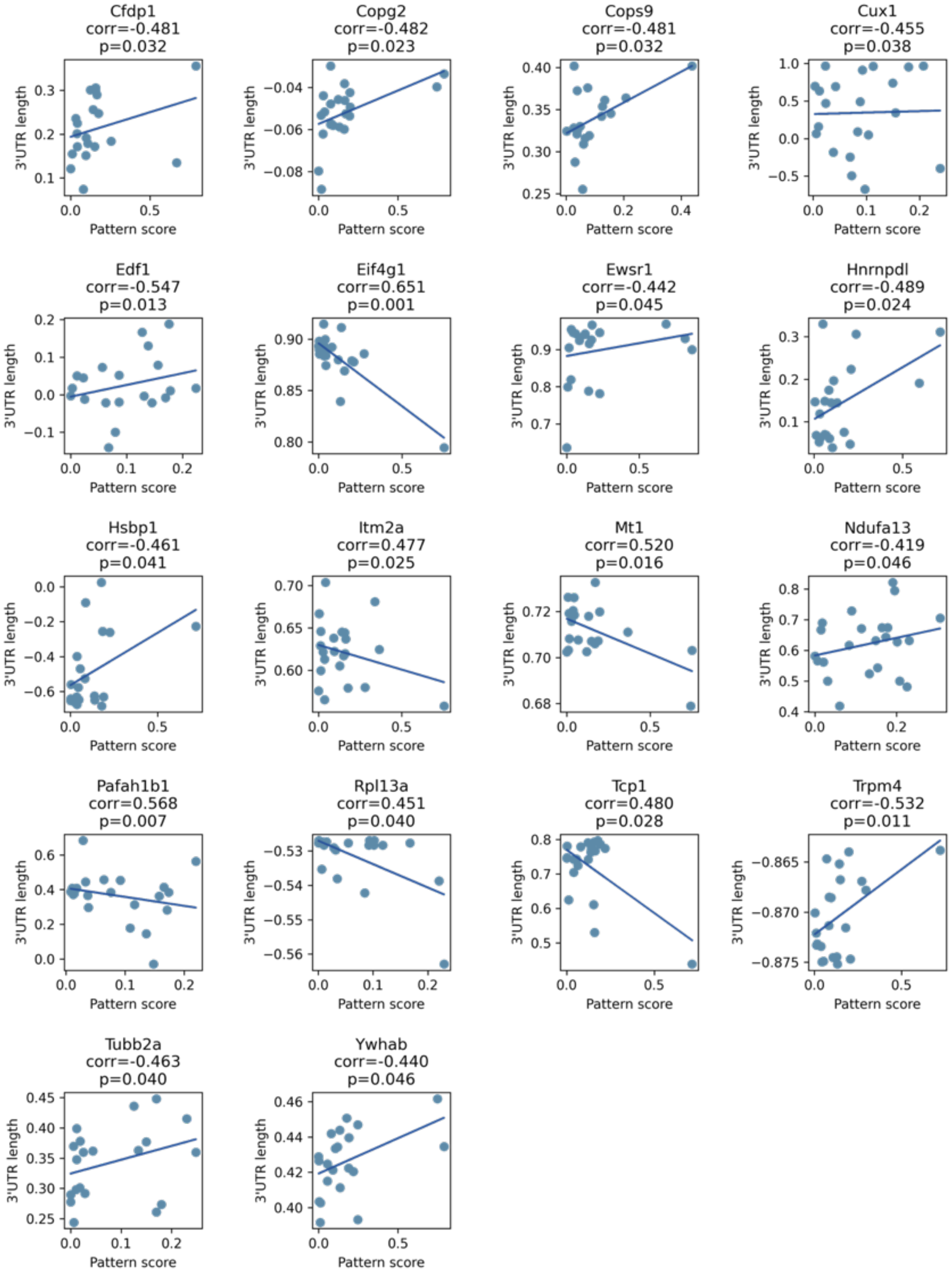

**Supplementary Figure 44. Estimated expression pattern for genes identified by ELLA in individual cell types in Stereo-seq. a.** The number and proportion of genes across five pattern clusters in each cell type. **b.** The estimated expression intensities for genes across clusters in each cell type. **c.** The estimated pattern score for genes across clusters in each cell type.

**a.**

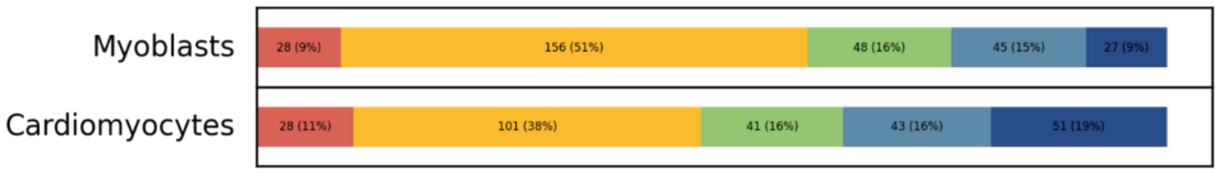

**b.**

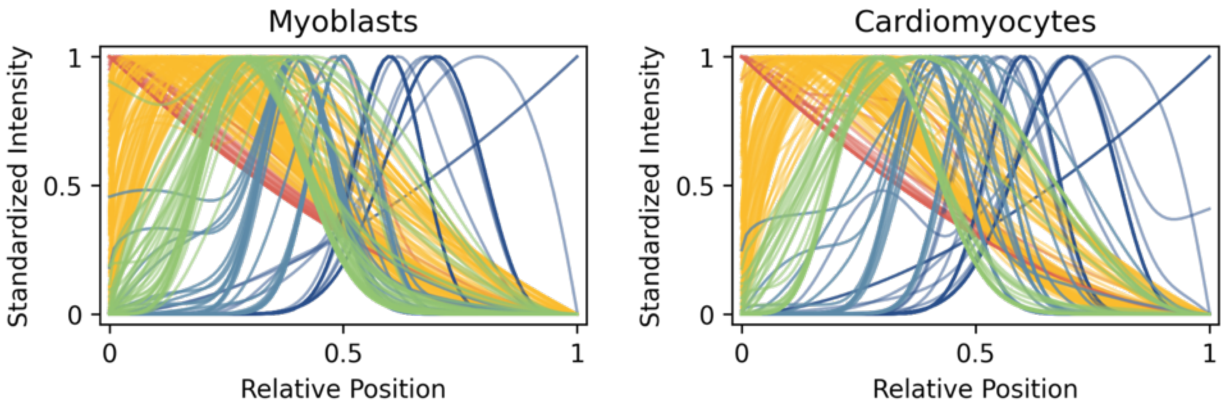

**c.**

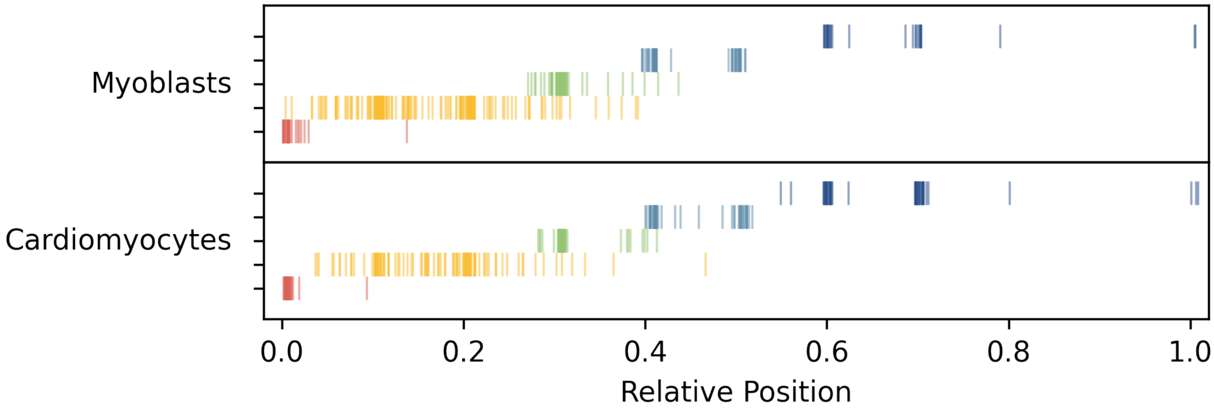

**Supplementary Figure 45. Venn plots of the numbers of commonly and uniquely detected genes in Myoblasts and Cardiomyocytes across the five pattern clusters in Stereo-seq.**

Cluster 1

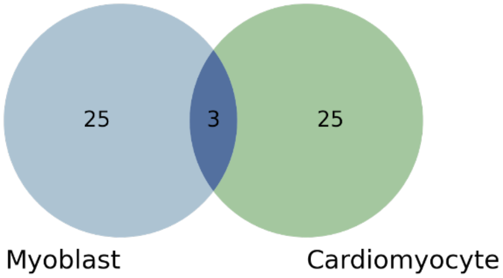

Cluster 2

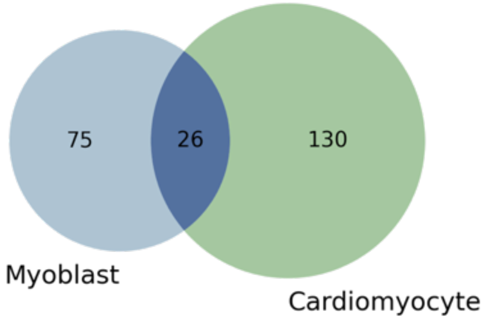

Cluster 3

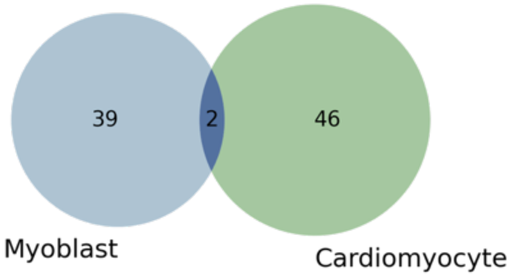

Cluster 4

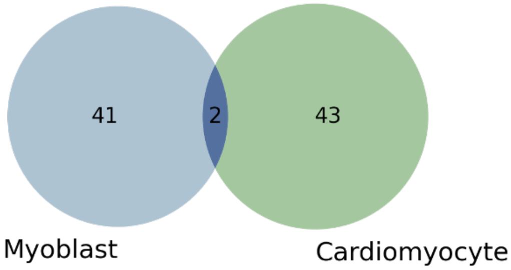

Cluster 5

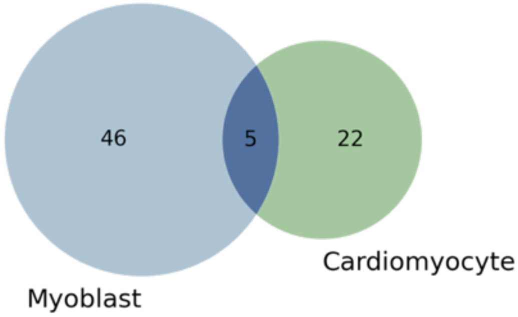

**Supplementary Figure 46. The estimated expression intensities of the commonly detected genes in Myoblasts (blue) and Cardiomyocytes (dashed green) across the five pattern clusters in Stereo-seq.** The estimated expression intensity of at most 5 genes in each cell type (solid blue in Myoblast, dashed green in Cardiomyocytes) are plotted per cluster (k=1-5).

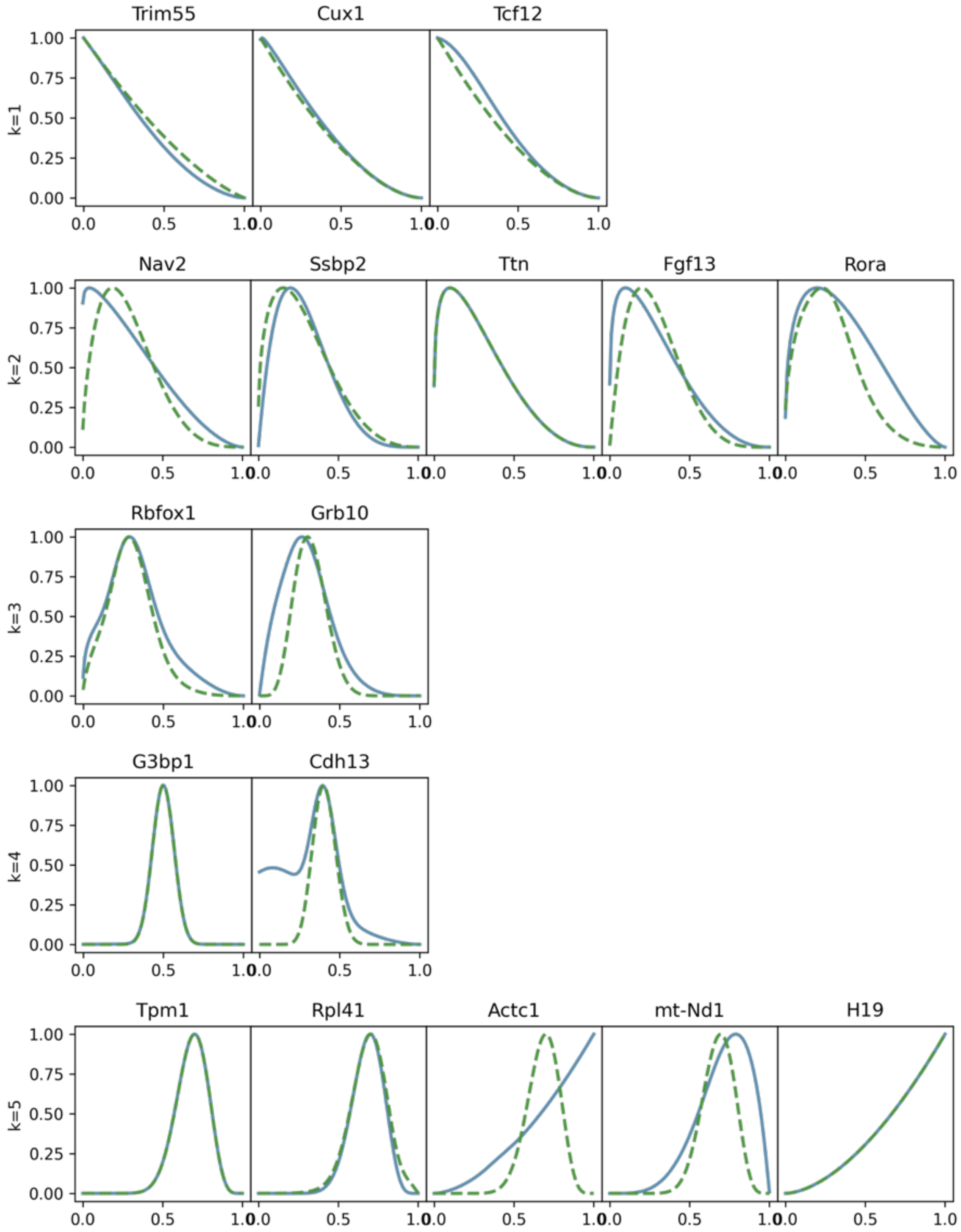

**Supplementary Figure 47. The commonly and uniquely detected transcription factors (TFs) in Myoblasts and Cardiomyocytes in Stereo-seq. a.** Venn plot shows the numbers of commonly and uniquely detected transcription factors in Myoblasts and Cardiomyocytes. **b-d.** Enriched gene sets (1% FDR) in GSEA analysis with GO Biological Processes for the commonly detected TFs, the Myoblast unique TFs, and the Cardiomyocyte unique TFs respectively.

**a.**

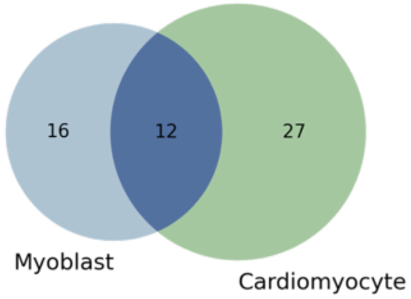

**b.**

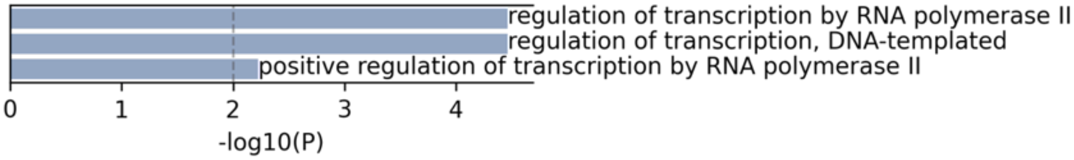

**c.**

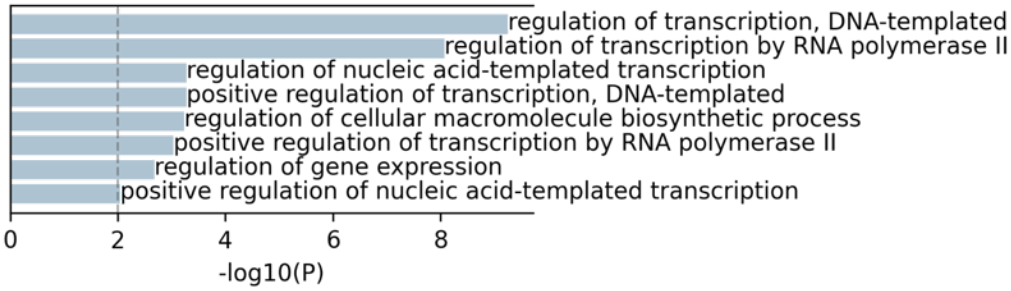

**d.**

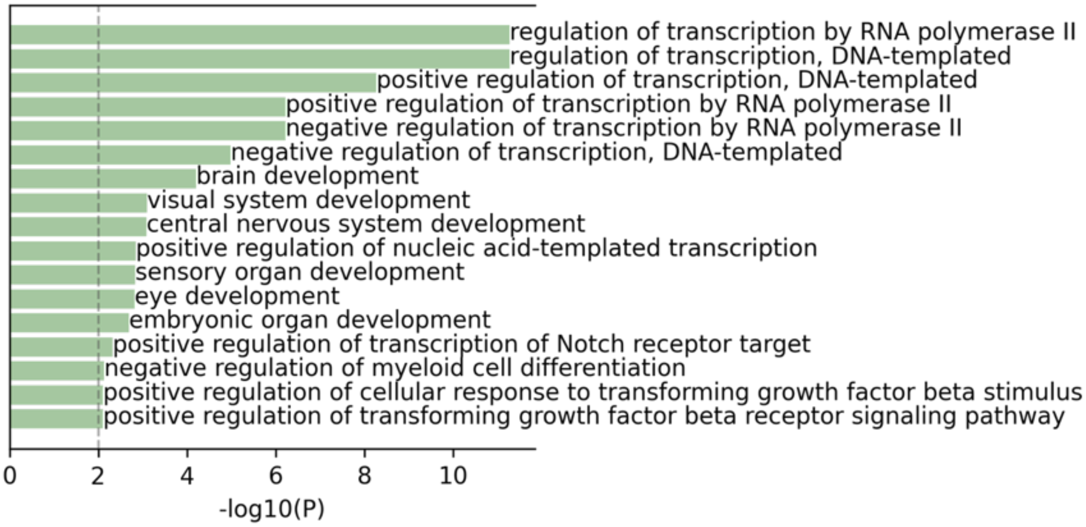

**Supplementary Figure 48. The commonly and uniquely detected long noncoding (lnc) genes in Myoblasts and Cardiomyocytes in Stereo-seq. a.** Venn plot shows the numbers of commonly and uniquely detected lnc genes in Myoblasts and Cardiomyocytes. b. The estimated expression intensity of the commonly detected seven genes in each cell type (solid blue in Myoblast, dashed green in Cardiomyocytes) are plotted.

**a.**

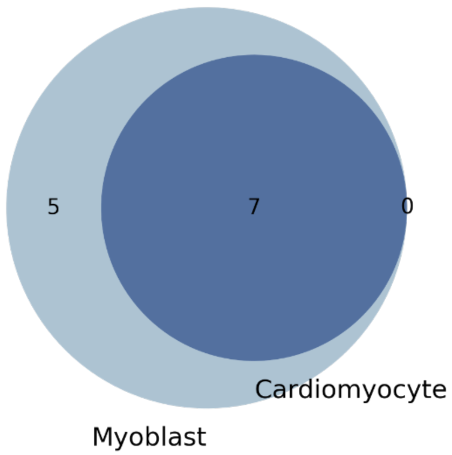

**b.**

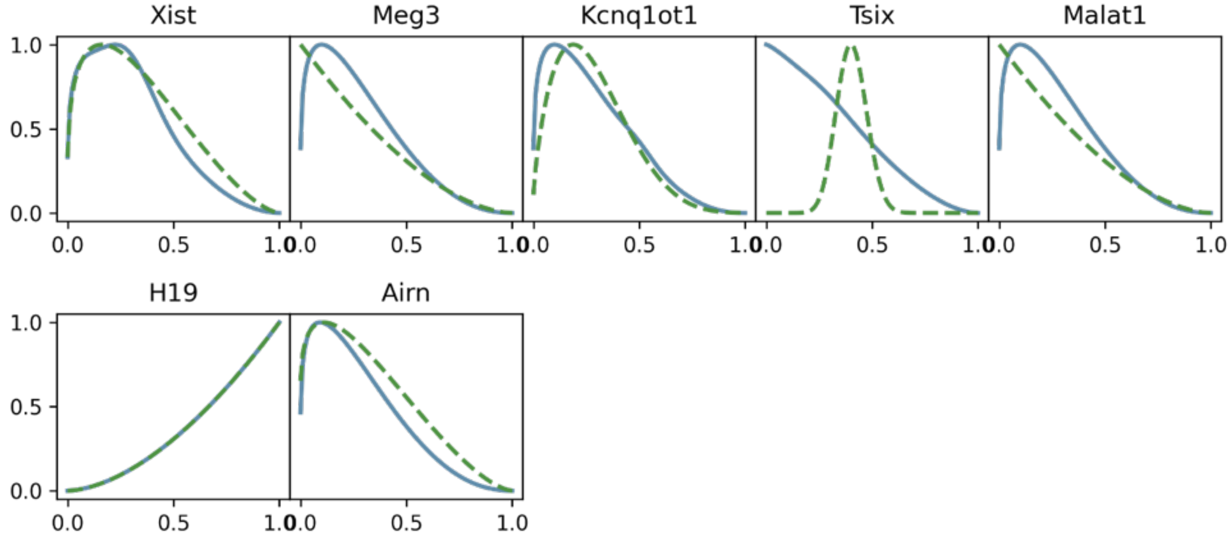

**Supplementary Figure 49. The commonly and uniquely detected mitochondrial genes in Myoblasts and Cardiomyocytes in Stereo-seq. a.** Venn plot shows the numbers of commonly and uniquely detected mitochondrial genes in Myoblasts and Cardiomyocytes. b. The estimated expression intensity of the commonly detected one gene (*mt-Nd1*) in each cell type (solid blue in Myoblast, dashed green in Cardiomyocytes) is plotted.

**a.**

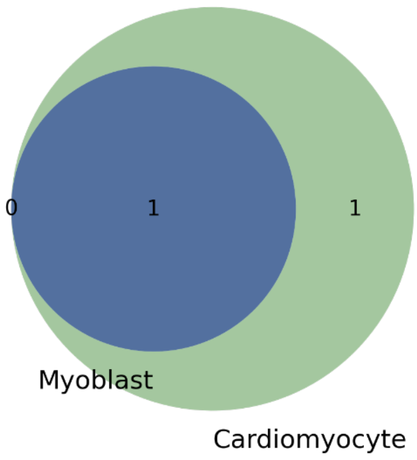

**b.**

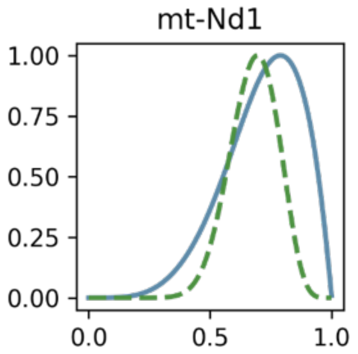

**Supplementary Figure 50. The estimated expression intensities of Myoblast and Cardiomyocyte marker genes in Stereo-seq across cell types.** The intensities of significant marker gene in the Myoblast cell type are plotted in (solid) blue, the intensities of significant marker gene in the Cardiomyocyte cell type are plotted in (dashed) green, the nonsignificant intensities are colored with (dashed) light grey. Two Cardiomyocyte cell type marker genes (*Acta1* and *Myh3*) are detected by ELLA as cytoplasmic localized (clusters 4) in the Cardiomyocyte cell type.

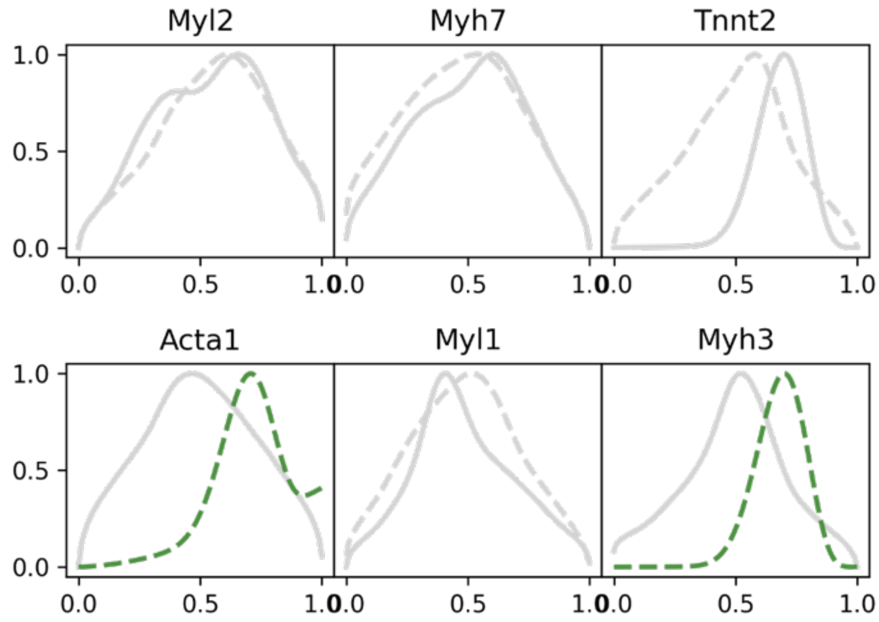

**Supplementary Figure 51. Preprocess of the seqFISH+ mouse embryonic fibroblast data. a.** Histogram of the average nucleus-cell ratio across cells. Dashed lines show the average nucleus-cell ratio and the cutoff ratios corresponding to two standard deviations from the average. **b.** Demo gene expressions (green dots) in randomly selected cells overlayed with nuclear segmentations (grey), cell segmentations (grey), and nuclear centers (red crosses).

**a.**

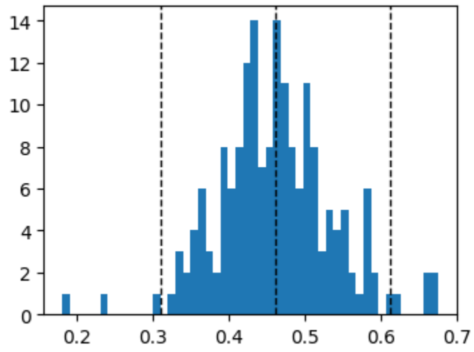

**b.**

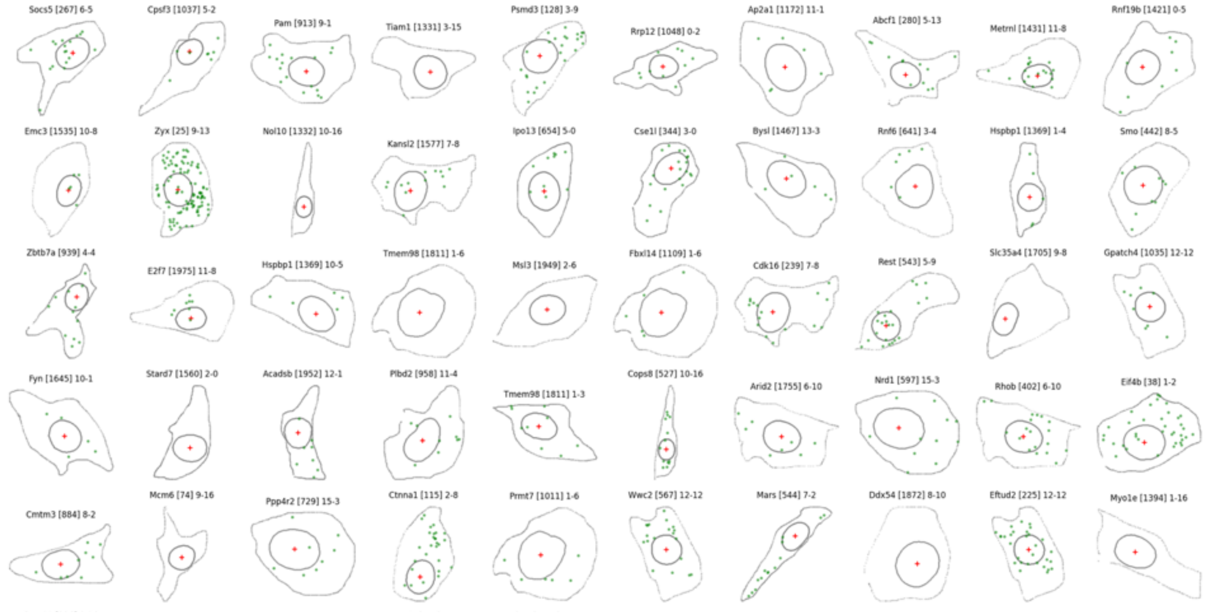

**Supplementary Figure 52. Stacked bar plot shows the pattern clusters identified by ELLA for three lists of localized genes in the seqFISH+ study.** The ELLA detected pattern clusters for the 20 nuclear and perinuclear genes, 20 cytoplasm genes, and 20 protrusion genes (1 out of 20 was not included in the ELLA analysis).

SeqFISH+ reported patterns:

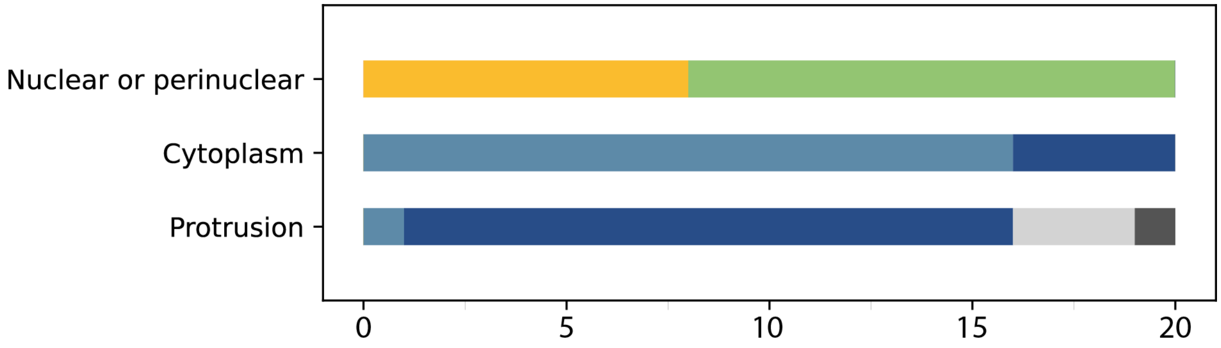

ELLA detected patterns:

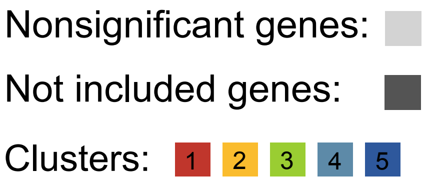

**Supplementary Figure 53. Real cells used for comparing Bento and ELLA.** Cell boundary and nuclear boundary are displayed for the 20 embryonic fibroblast cells from the seqFISH+ data that are used to compare Bento and ELLA. Both ELLA and Bento were applied to analyze one cell at a time.

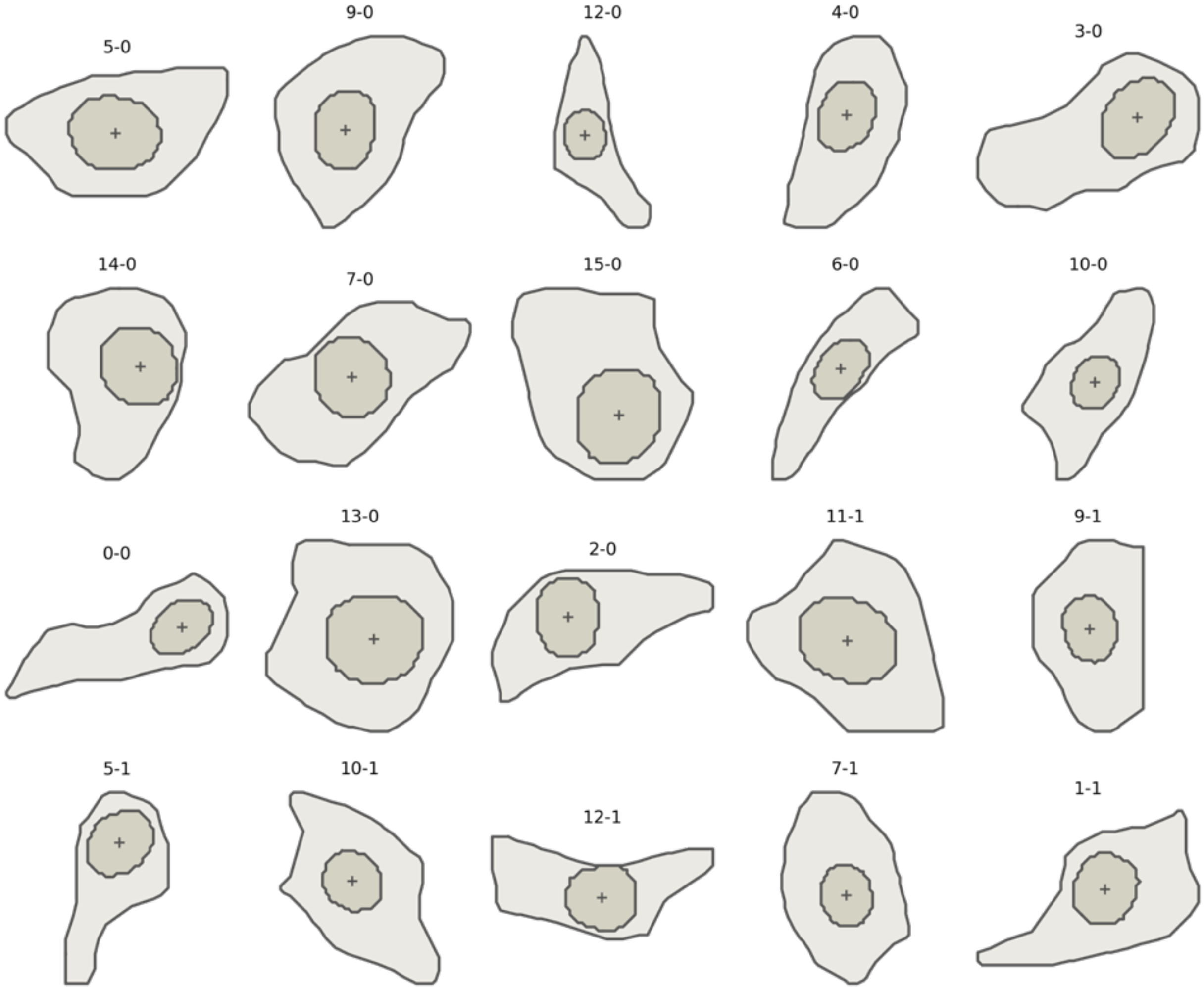

**Supplementary Figure 54. The expression patterns identified by Bento and ELLA in the seqFISH+ cell by cell analysis. a.** Pie chat shows the proportion of genes across different patterns assigned by Bento. **b.** Pie chat shows the proportion of genes across five pattern clusters identified by ELLA. Within each pattern cluster, the proportion of gene achieves statistical significance (5% FDR) is highlighted with a corresponding dark color. **c.** The estimated expression intensities for all genes across five clusters estimated by ELLA.

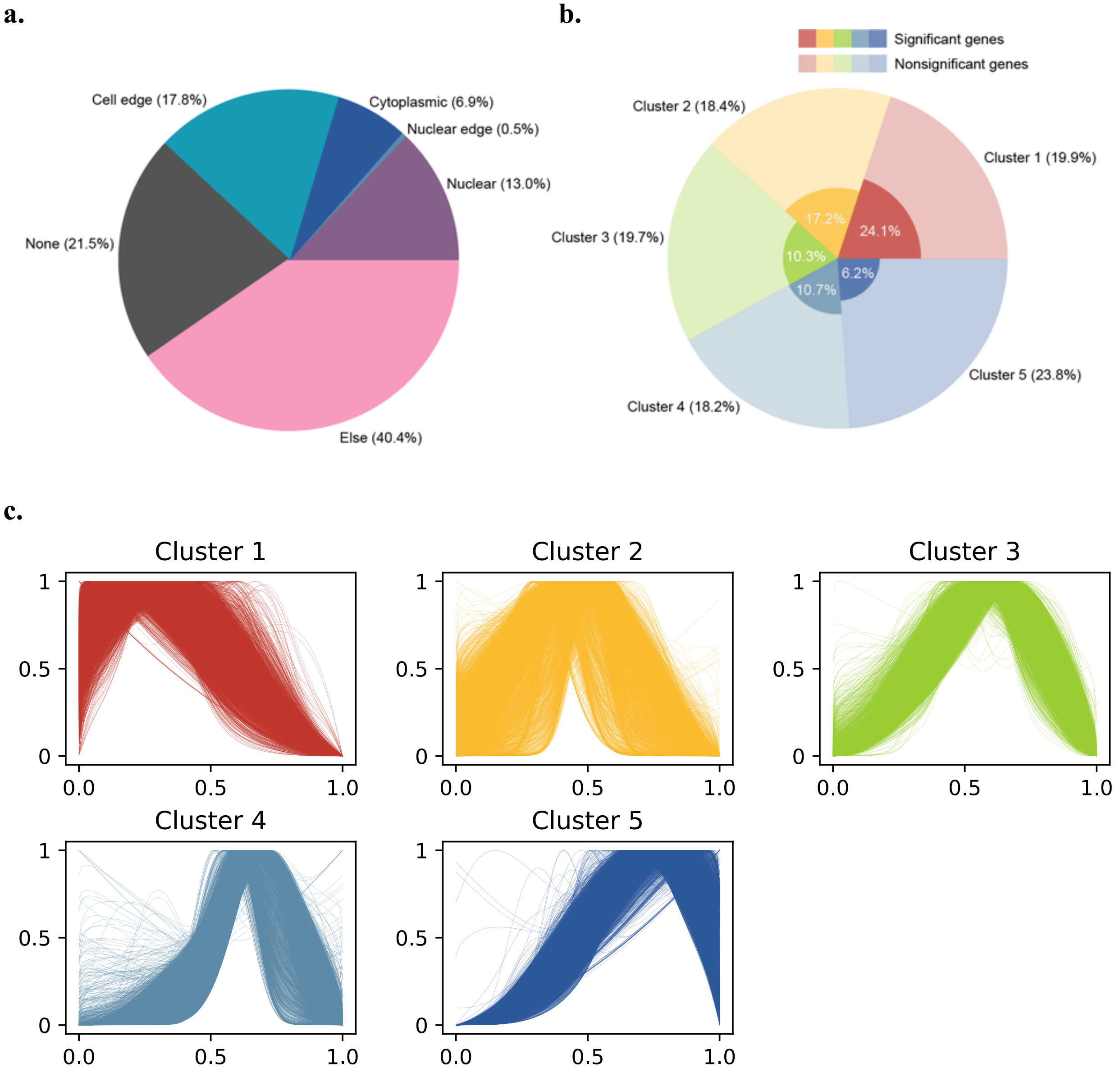

**Supplementary Figure 55. Sankey plot compares expression patterns identified by Bento and ELLA in the seqFISH+ cell by cell analysis.** Left bars show the patterns of genes assigned by Bento. Bar length is proportional to the number of genes of each pattern. Right bars show the pattern clusters identified by ELLA. For example, ELLA-1 refers to cluster 1 nonsignificant genes, ELLA-sig 1 refers to cluster 1 significant genes based on 5% FDR. A band connecting a left bar (a Bento category) and a right bar (an ELLA category) represents the number of genes assigned to a certain pattern by Bento (on the left) that was identified by ELLA as a certain pattern (on the right). Width of bands is proportional to the number of genes.

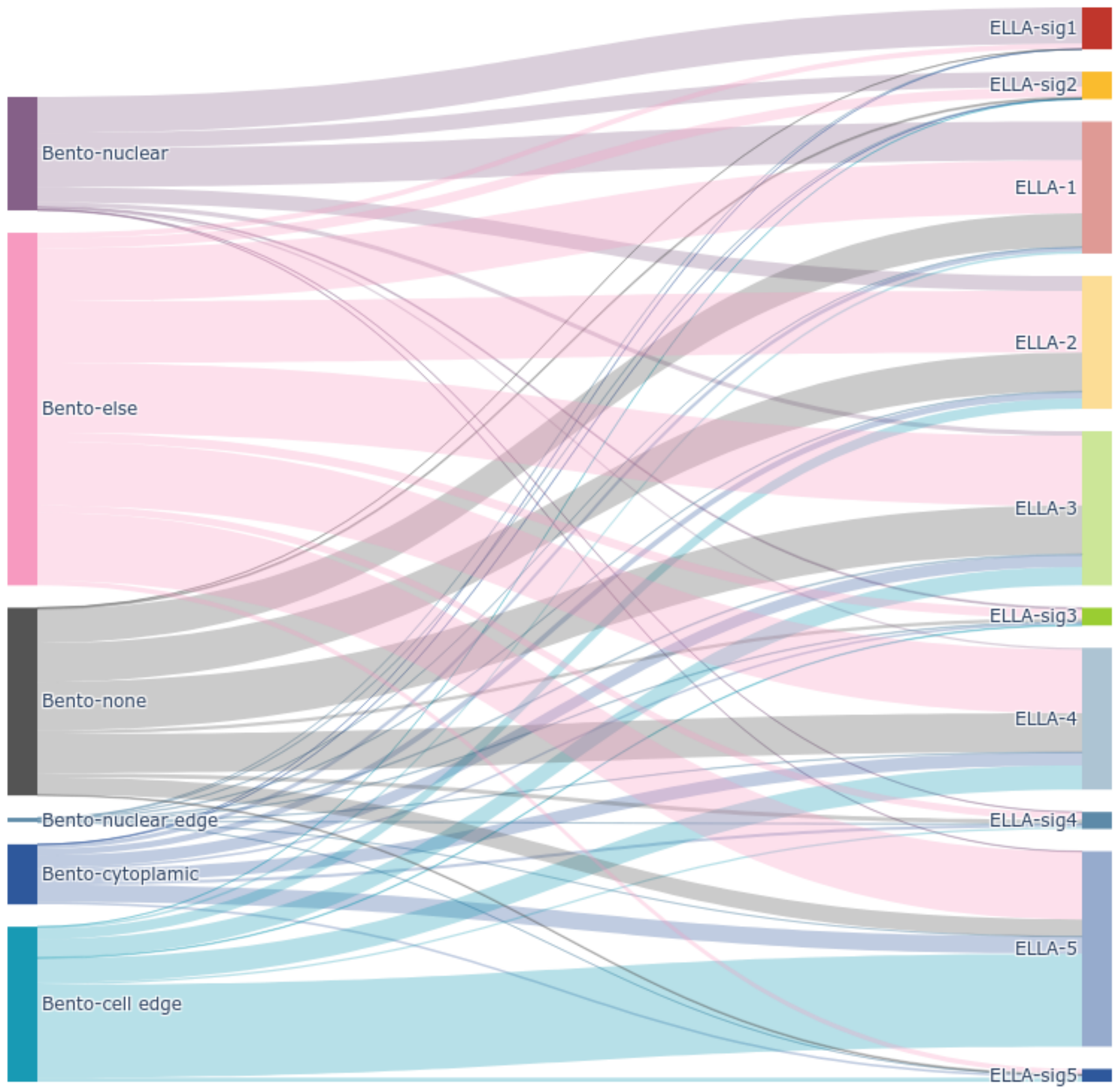

**Supplementary Figure 56. Preprocess of the MERFISH adult mouse brain data.** In a subregion, the total gene expressions (grey dots) are overlayed with cell segmentations across five z stacks (varying colors), and cell centroids (red crosses).

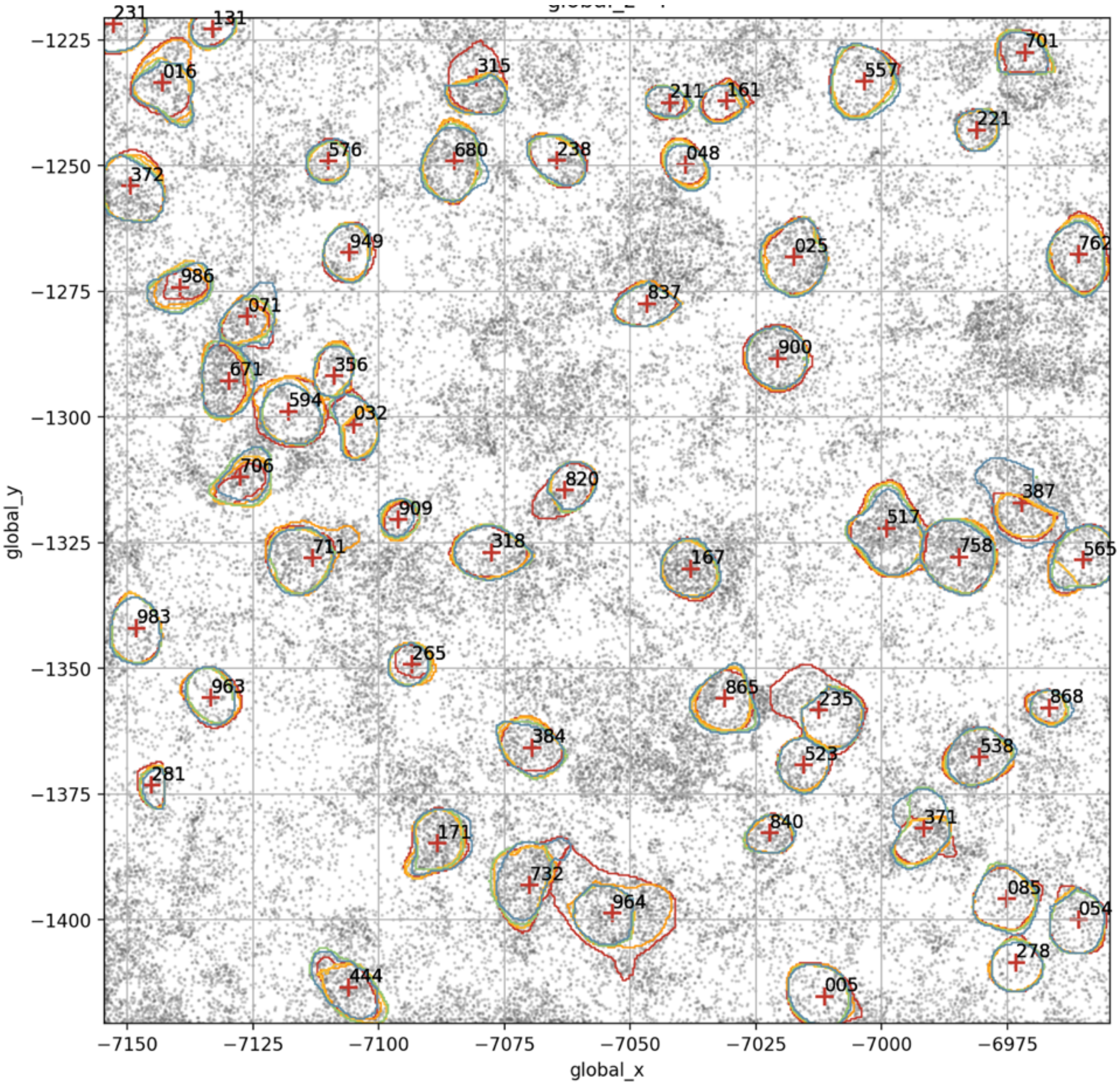

**Supplementary Figure 57. Number of genes and cells in the MERFISH adult mouse brain data. a.** Bar plot shows the number of genes analyzed by ELLA across cell types. **b.** Bar plot shows the number of cells available across cell types. **c.** Bar plot shows the number of genes detected by ELLA across cell types.

**a.**

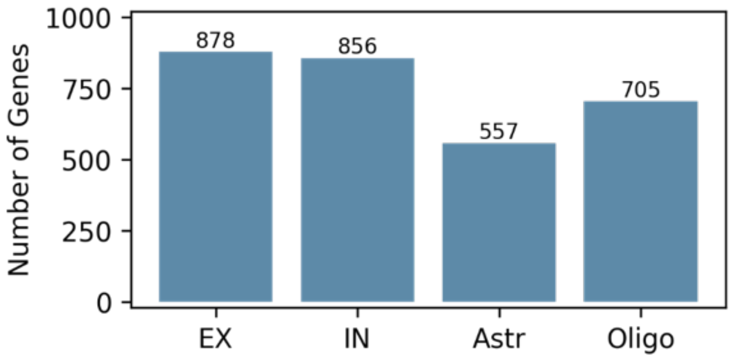

**b.**

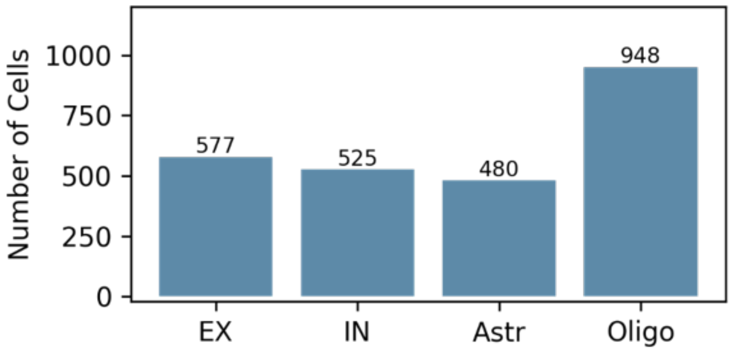

**c.**

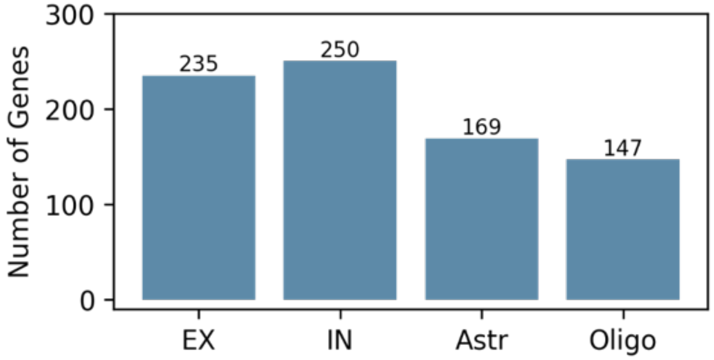

**Supplementary Figure 58. Sankey plots compare expression patterns identified by SPRAWL and ELLA in the MERFISH adult mouse brain data.** Left bars show the proportion of significant genes (purple) and nonsignificant genes (light grey) detected by a SPRAWL metric. Right bars show proportions of gene across pattern clusters (colored) identified by ELLA and the proportion of nonsignificant genes (light grey). Bar length is proportional to the number of genes. A band connecting a left bar (a SPRAWL category) and a right bar (an ELLA category) represents the number of genes assigned to a certain category by SPRAWL (on the left) that is identified by ELLA as a certain category (on the right). Width of bands is proportional to the number of genes. **a.** SPRAWL-peripheral metric and ELLA, **b.** SPRAWL-central metric and ELLA, **c.** SPRAWL-radial metric and ELLA, **d.** SPRAWL-punctate metric and ELLA.

**a.**

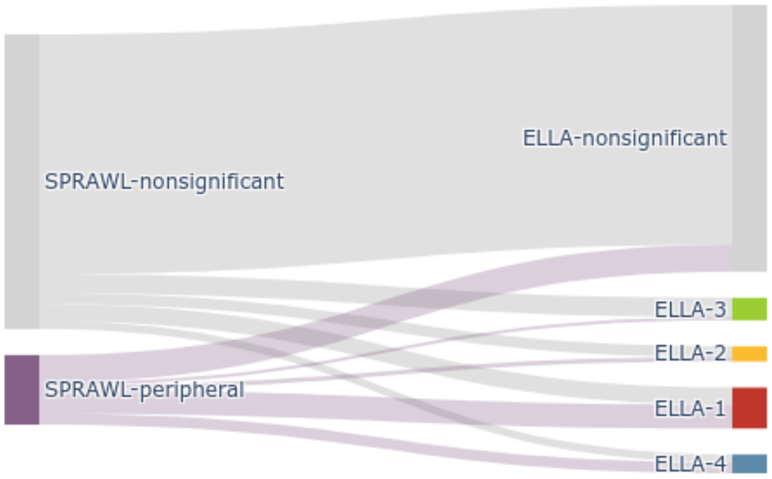

**b.**

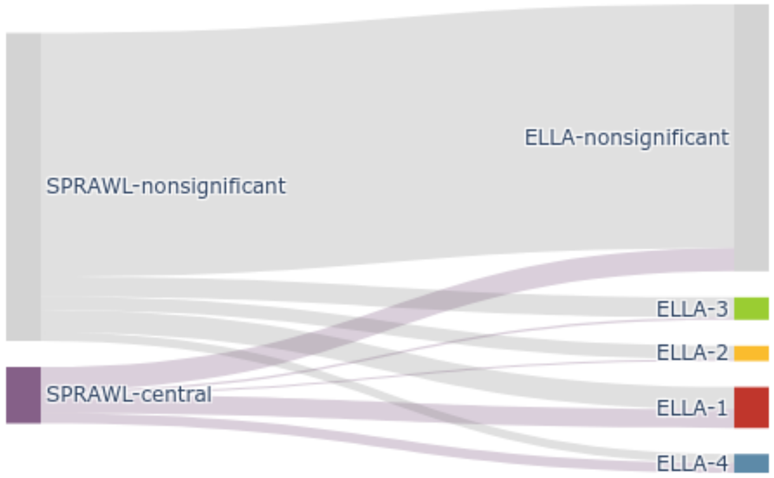

(Fig. S58 con'd)

**c.**

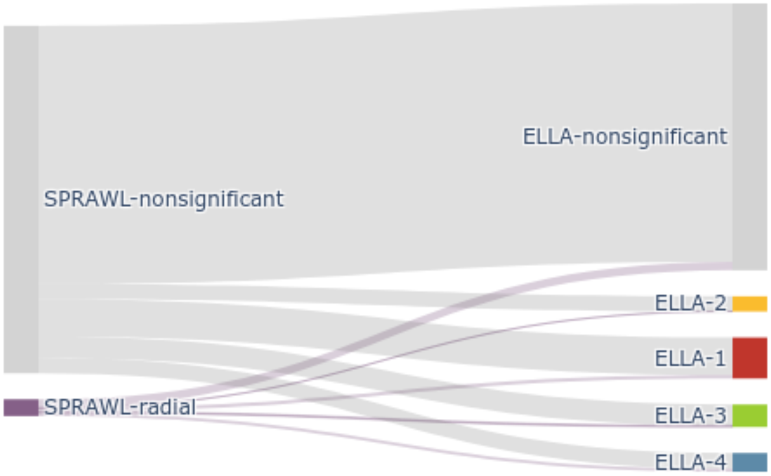

**d.**

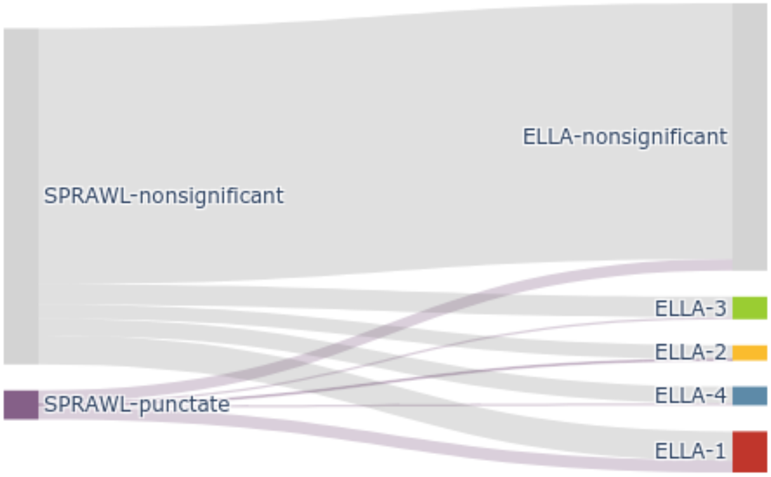

**Supplementary Figure 59. GSEA for genes in pattern clusters 1-2 in the MERFISH adult mouse brain data.** Stem plots show the -log10 P values of the top 10 enriched gene sets in GSEA analysis for genes in pattern cluster 1 and 2 respectively. Included gene sets: GO Biological Process (2021), GO Cellular Component (2021), GO Molecular Function (2021), and KEGG (2019).

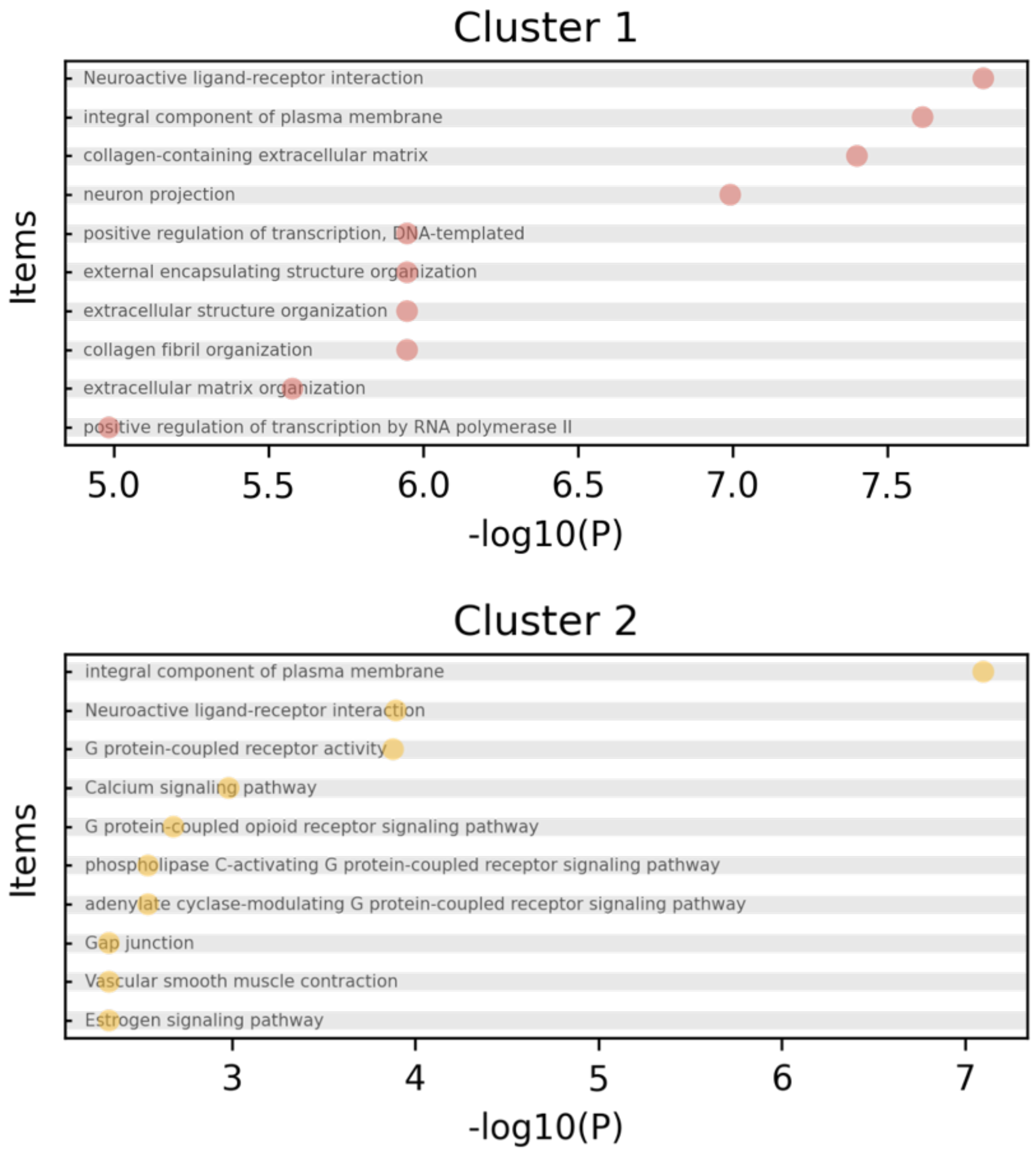

**Supplementary Figure 60. Estimated expression pattern for genes identified by ELLA in individual cell types in the MERFISH adult mouse brain data. a.** The number and proportion of genes across five pattern clusters in each cell type. **b.** The estimated expression intensities for genes across clusters in each cell type. **c.** The estimated pattern score for genes across clusters in each cell type.

**a.**

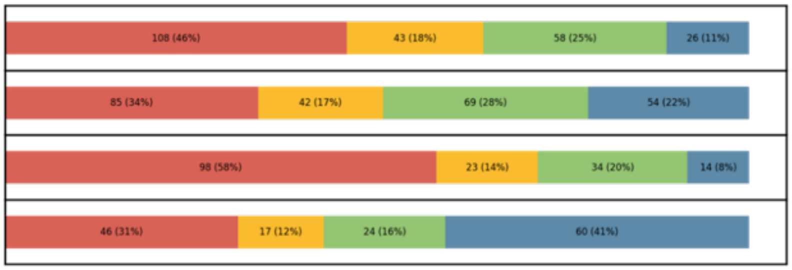

**b.**

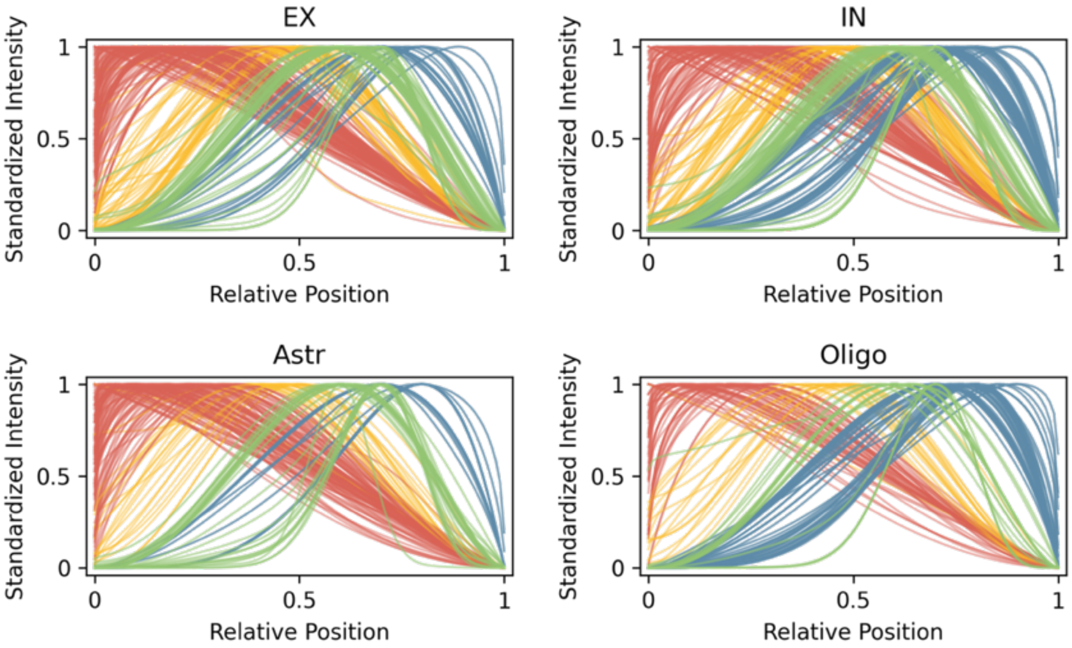

**c.**

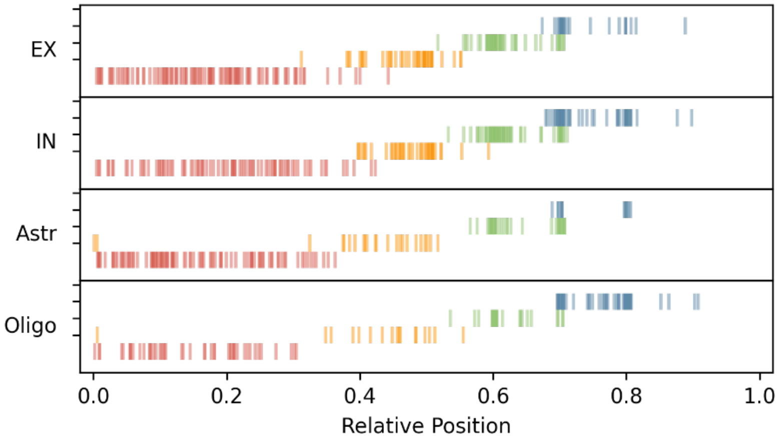

**Supplementary Figure 61. Venn plots of the numbers of commonly and uniquely detected genes in EX and IN cells across four pattern clusters in the MERFISH adult mouse brain data.**

Cluster 1

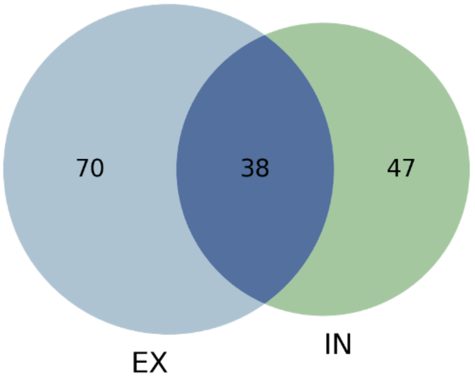

Cluster 2

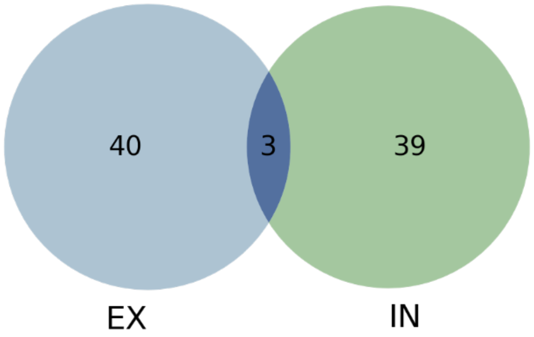

Cluster 3

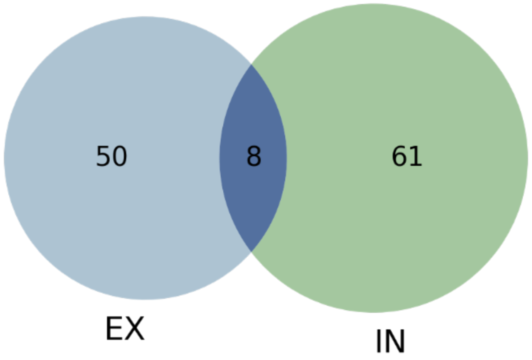

Cluster 4

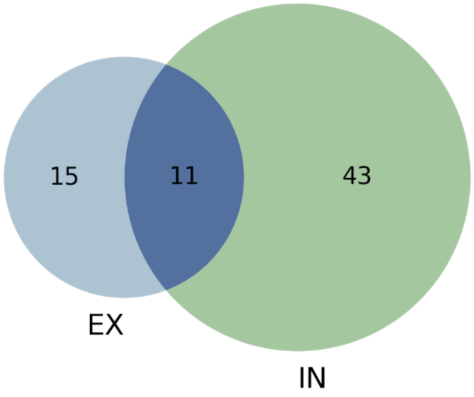

**Supplementary Figure 62. The estimated expression intensities of the commonly detected genes in EX (blue) and IN (dashed green) cells across four pattern clusters in the MERFISH adult mouse brain data.** The estimated expression intensities of at most 5 genes in each cell type are plotted per cluster (k=1-5).

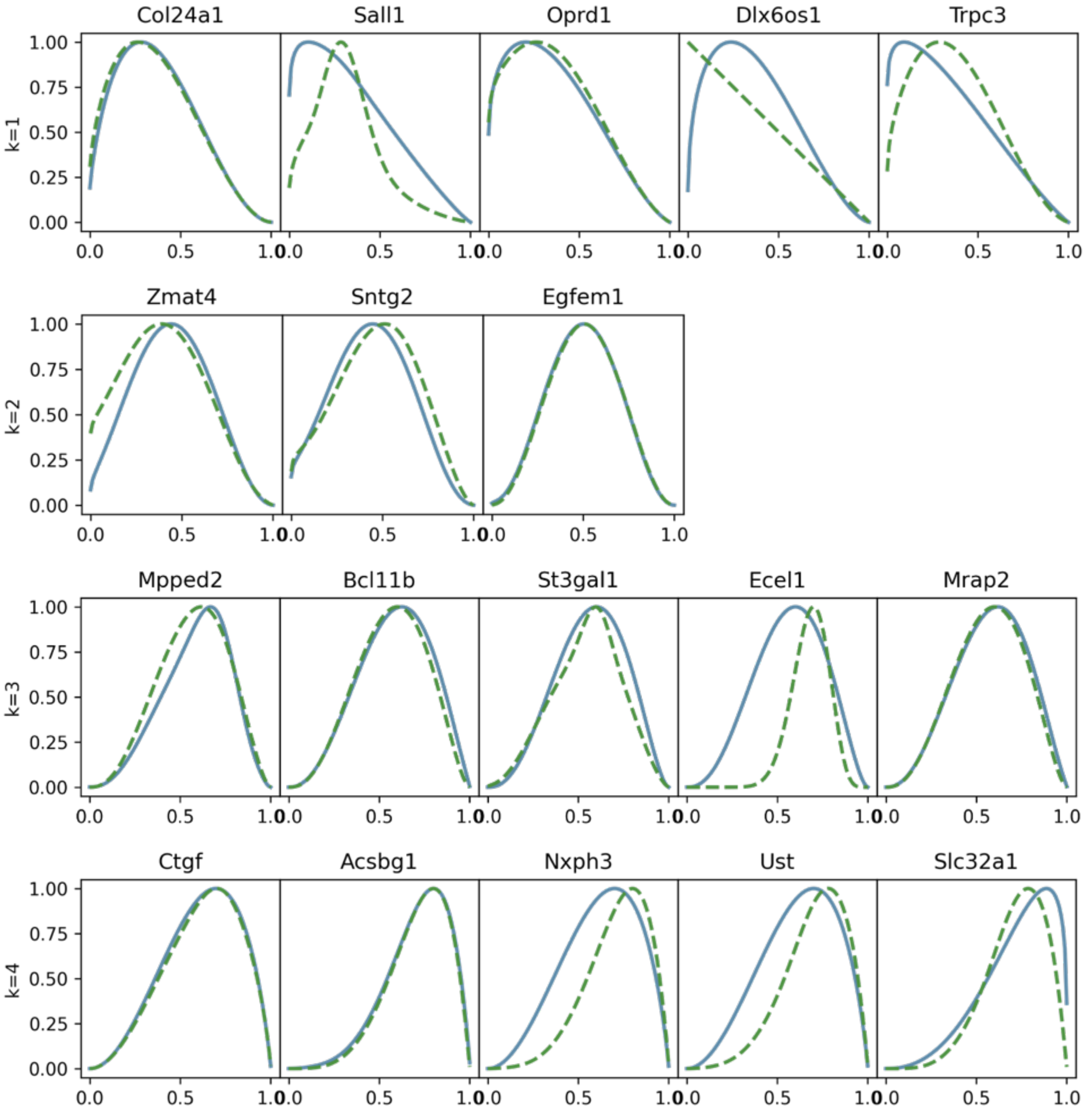

**Supplementary Figure 63. The commonly and uniquely detected transcription factors (TFs) in EX and IN cells in the MERFISH adult mouse brain data. a.** Venn plot shows the numbers of commonly and uniquely detected transcription factors in EX and IN cells. **b.** The estimated expression intensity of the 20 (out of 126) commonly detected TFs across cell types (solid blue in EX, dashed green in IN) are plotted.

**a.**

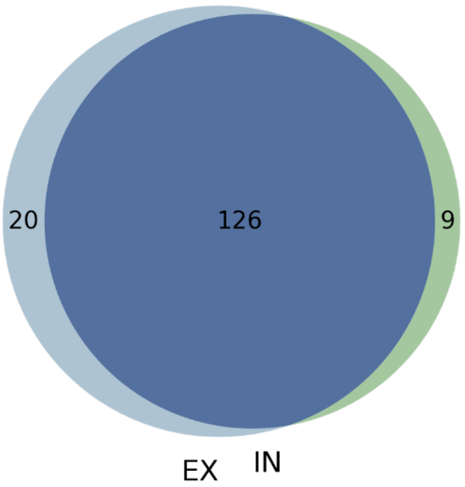

**b.**

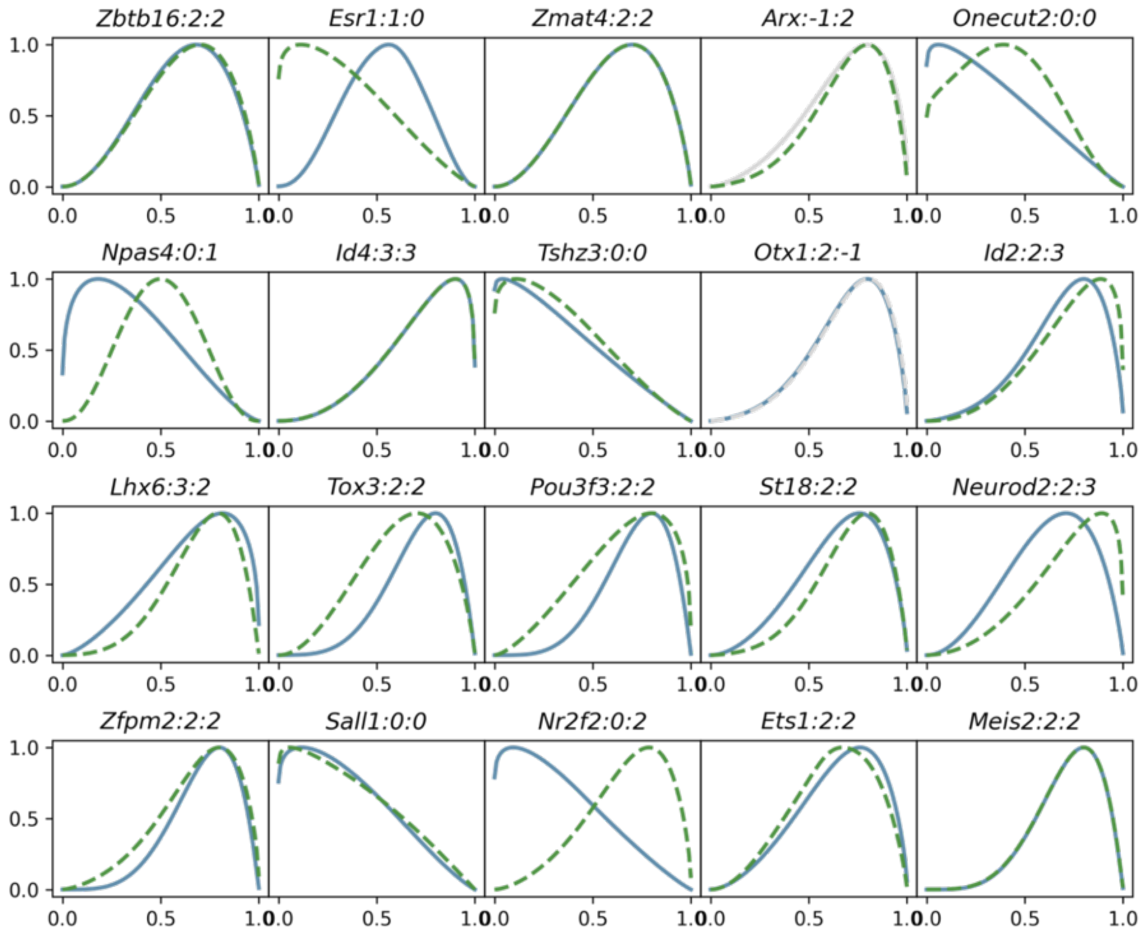

**Supplementary Figure 64. GSEA for the commonly detected transcription factors in EX and IN cells in the MERFISH adult mouse brain data.** Plot shows the -log10 P values of the top 50 enriched gene sets in the GSEA with GO Biological Processes.

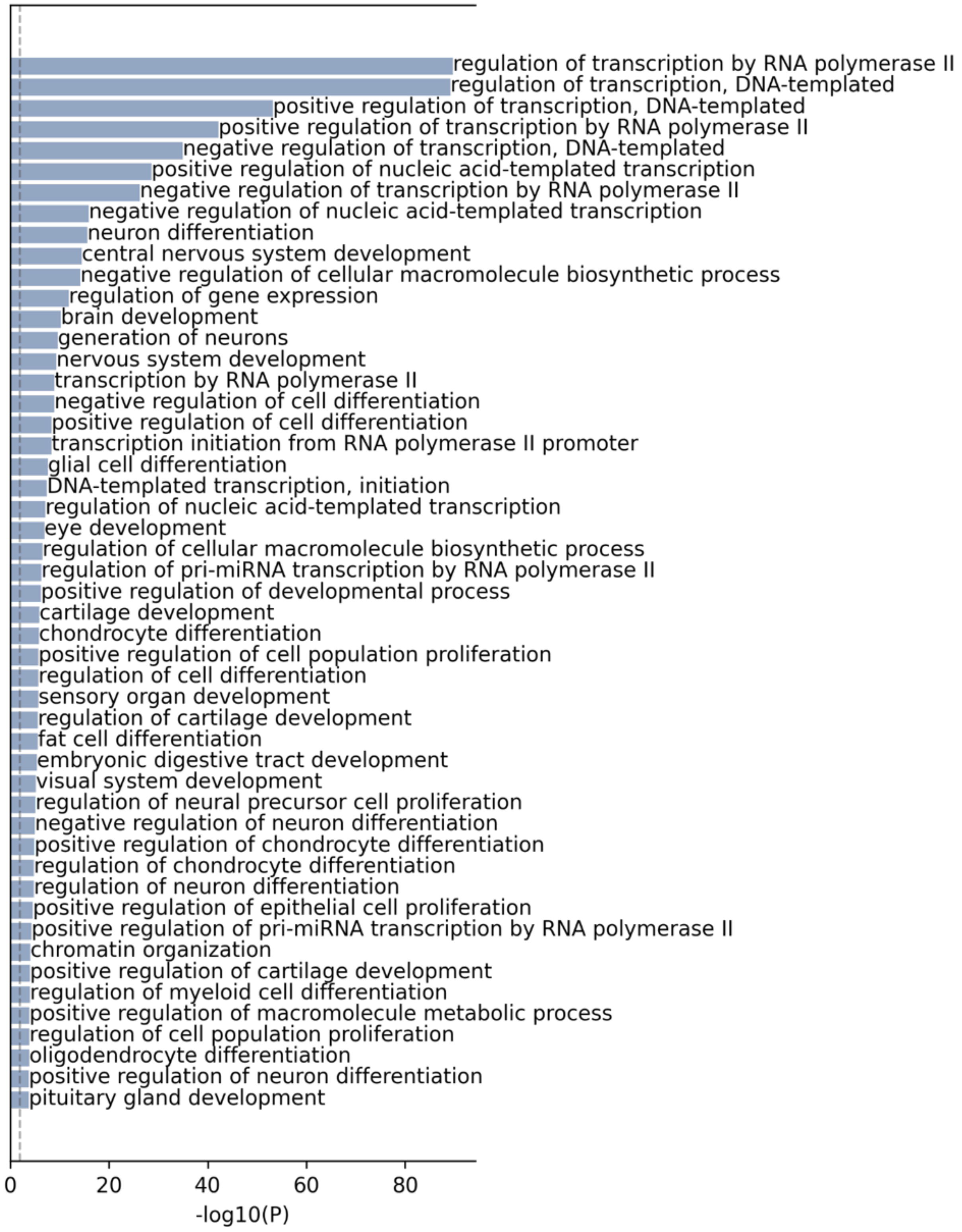

**Supplementary Figure 65. GSEA for the uniquely detected transcription factors (TFs) in EX and IN cells in the MERFISH adult mouse brain data. a.** Plot shows the -log10 P values of enriched gene sets for the TFs uniquely identified in the EX with GO Biological Processes based on 5% FDR. **b.** Plot shows the -log10 P values of enriched gene sets for the TFs uniquely identified in the IN with GO Biological Processes based on 5% FDR.

**a.**

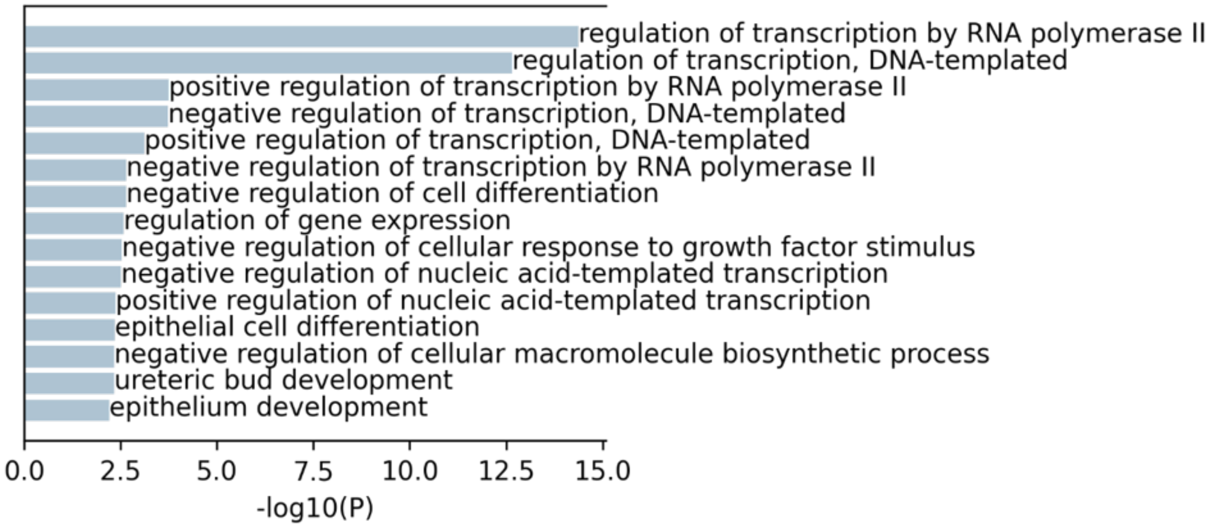

**b.**

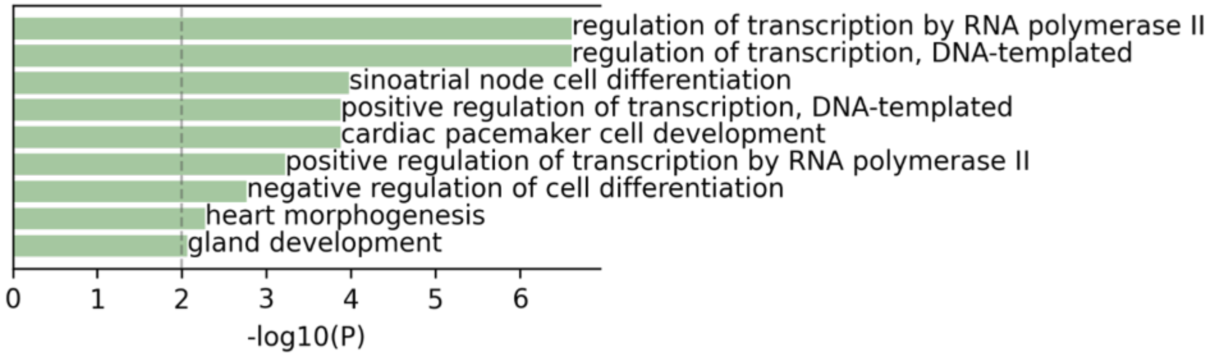

**Supplementary Figure 66. The commonly and uniquely detected long noncoding (lnc) genes in EX and IN cells in the MERFISH adult mouse brain data. a.** Venn plot shows the numbers of commonly and uniquely detected lnc genes in EX and IN. b. The estimated expression intensities of the commonly detected four genes in each cell type (solid blue in EX, dashed green in IN) are plotted.

**a.**

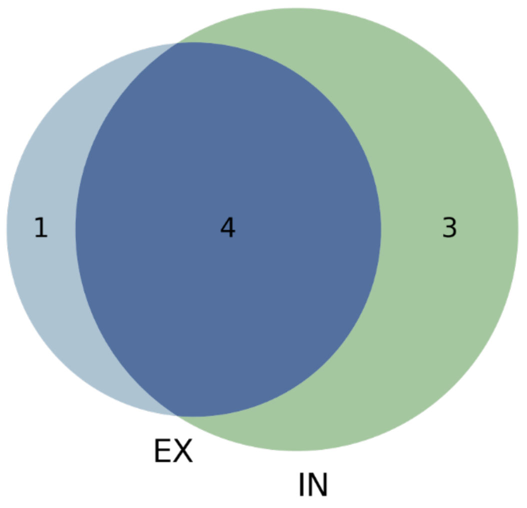

**b.**

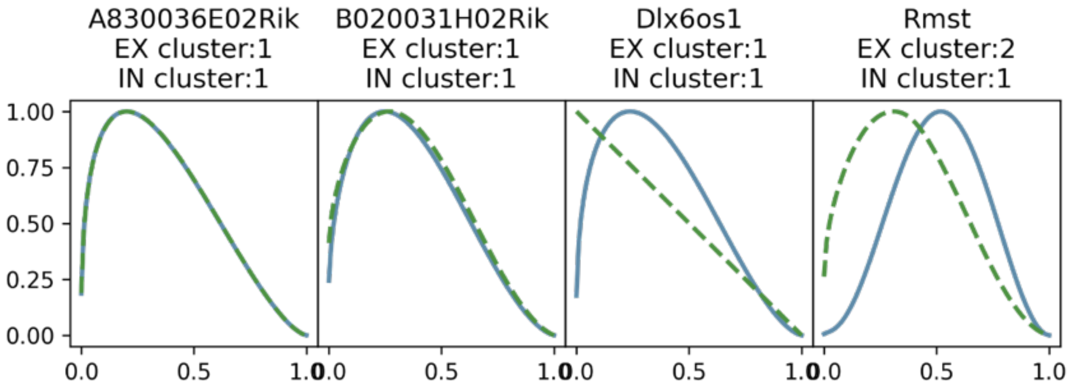

**Supplementary Figure 67. The estimated expression intensities of EX and IN marker genes in the MERFISH adult mouse brain data.** The intensities of significant marker genes in the EX cell type are plotted in (solid) blue, the intensities of significant marker genes in the IN cell type are plotted in (dashed) green, and the nonsignificant intensities are colored with (dashed) light grey.

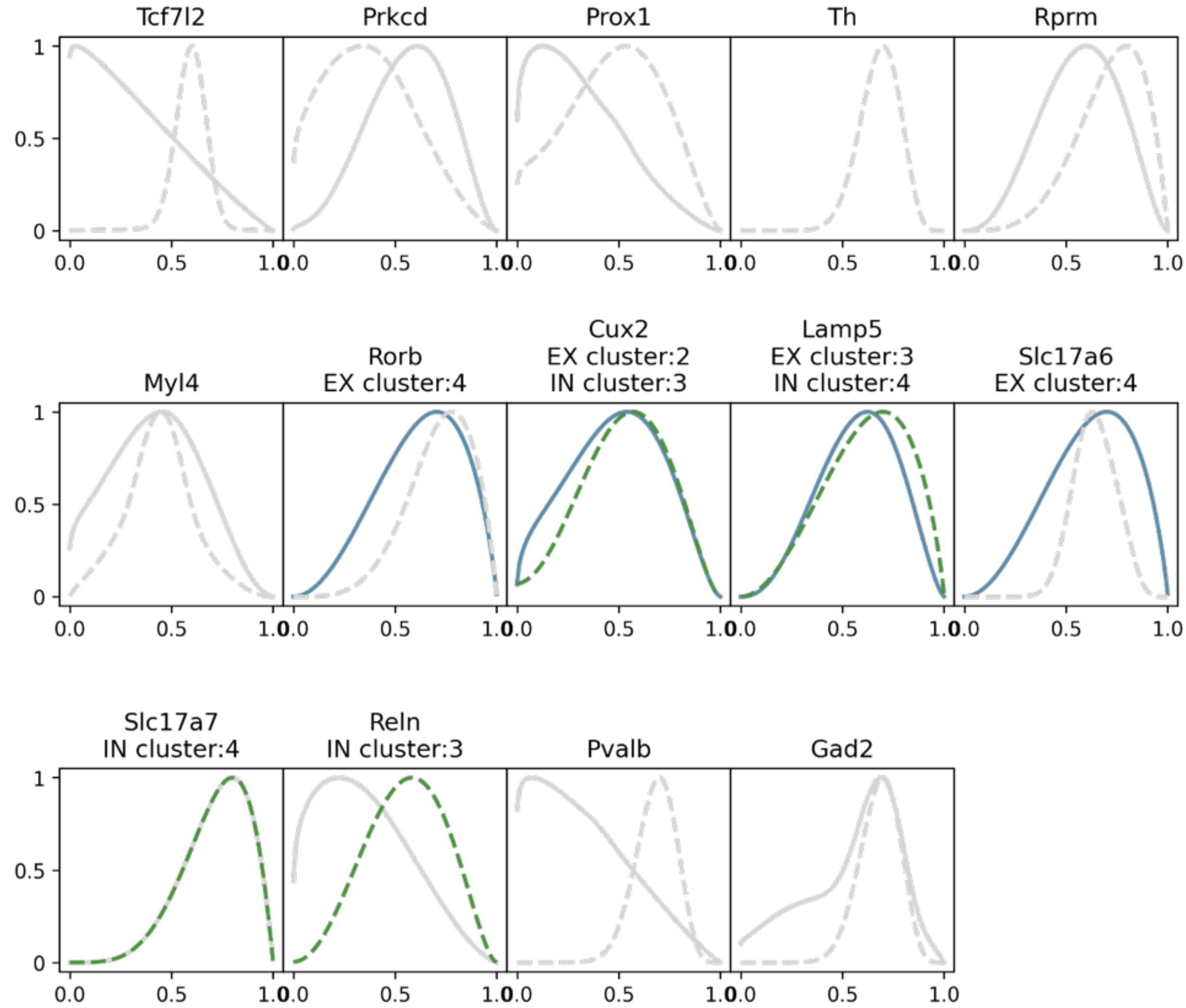

**Supplementary Figure 68. ELLA cell registration and ELLA default kernel functions $\varphi(r)$.** **a.** We use one embryonic fibroblast cell (cell 3-14) in the seqFISH+ data to illustrate the original cell (left) with the expression of one gene (*Aamp*) and the corresponding registered cell (right) with the expression of the same gene. For the transcript of focus (red dot), the relative position is computed as d1/(d1+d2). **b.** Multiple cells in seqFISH+ demonstrating the subcellular spatially gene expression prior to and after cell registration. **c.** The 22 default Beta kernel functions in ELLA. Each kernel function is standardized by its min and max value as $[\varphi(r) - \min \varphi(r)]/[\max \varphi(r) - \min \varphi(r)]$ for plotting. The first row shows the first set of 11 kernel functions with strong patterns and the max intensity values are taken at relative position=0, 0.1, …, 1 respectively; and the second row shows the second set of 11 kernel functions with less strong patterns and the max intensity values are taken at relative position=0, 0.1, …, 1 respectively.

**a.**

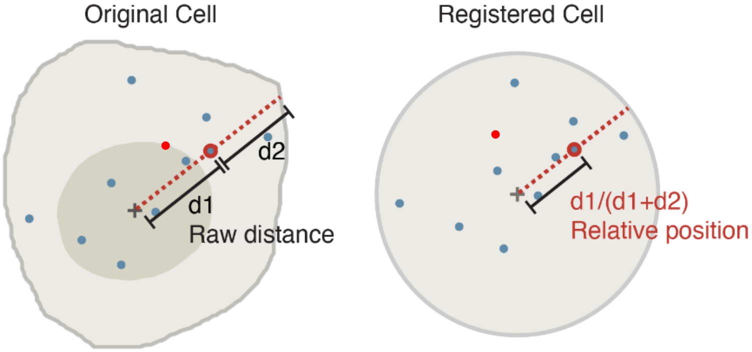

**b.**

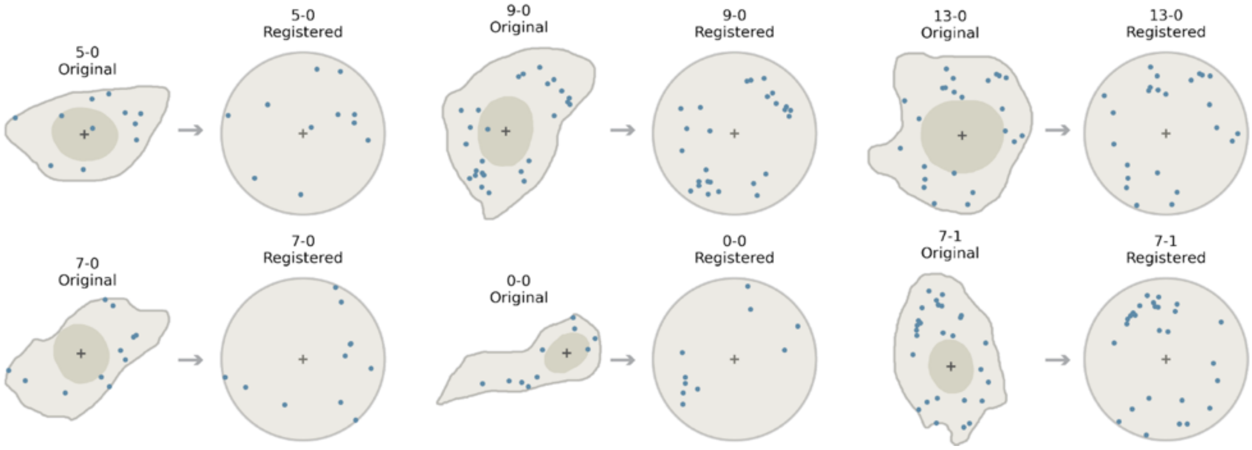

**c.**

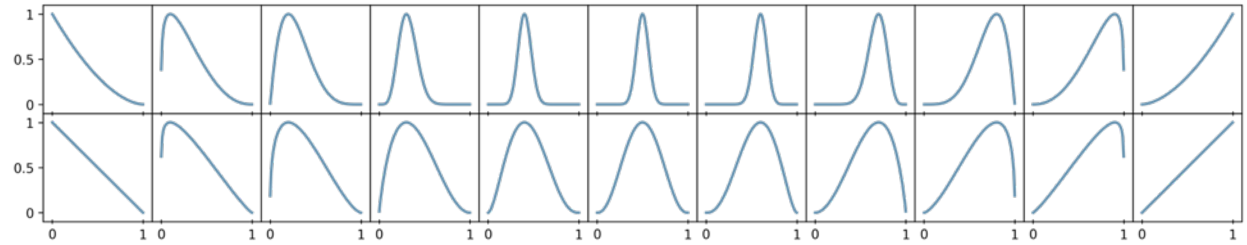

**Supplementary Figure 69. Cell segmentations produced by Cellpose based on the normal 10X H&E images in Seq-Scope.** Cell segmentation boundaries (red) are overlayed on the H&E image (greyscale) produced by Cellpose.

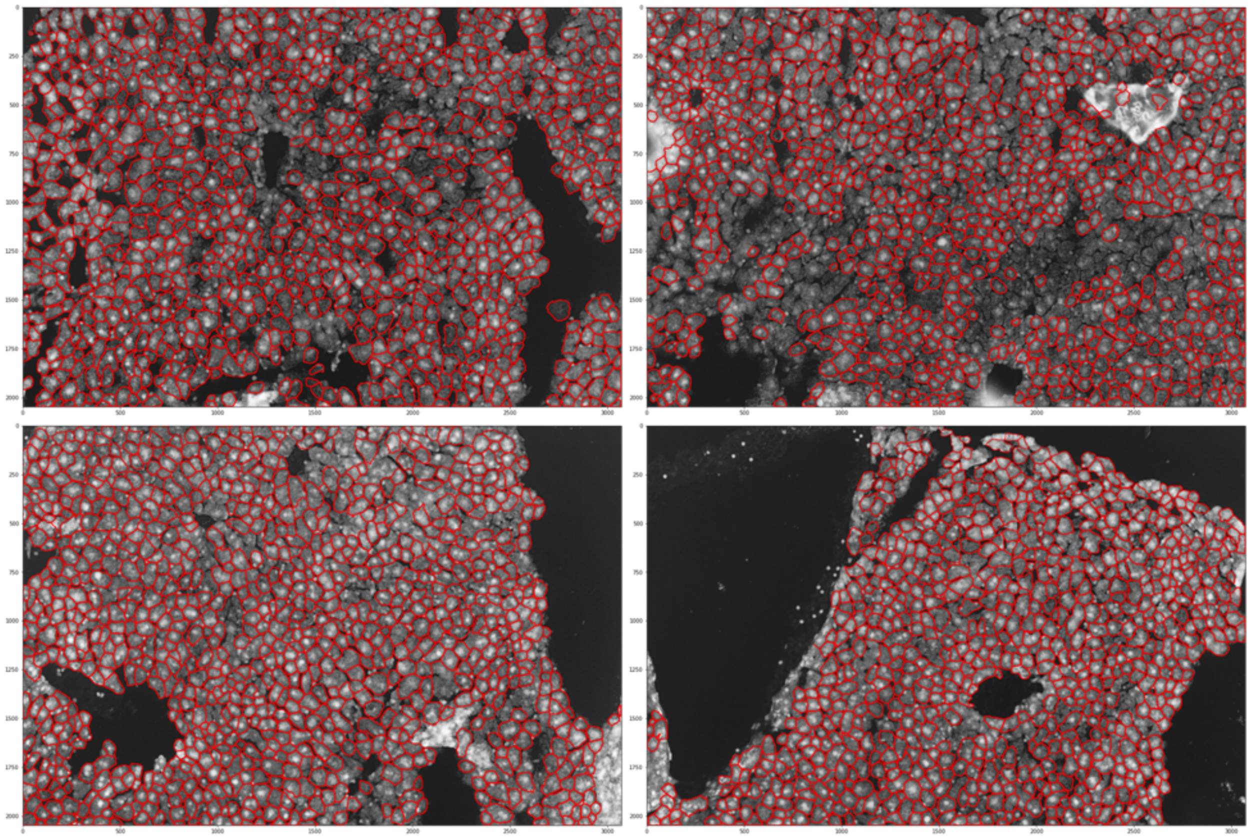

**Supplementary Figure 70. Cell segmentations produced by Cellpose based on the TD 10X H&E images in Seq-Scope.** Cell segmentation boundaries (red) are overlayed on the H&E image (greyscale) produced by Cellpose.

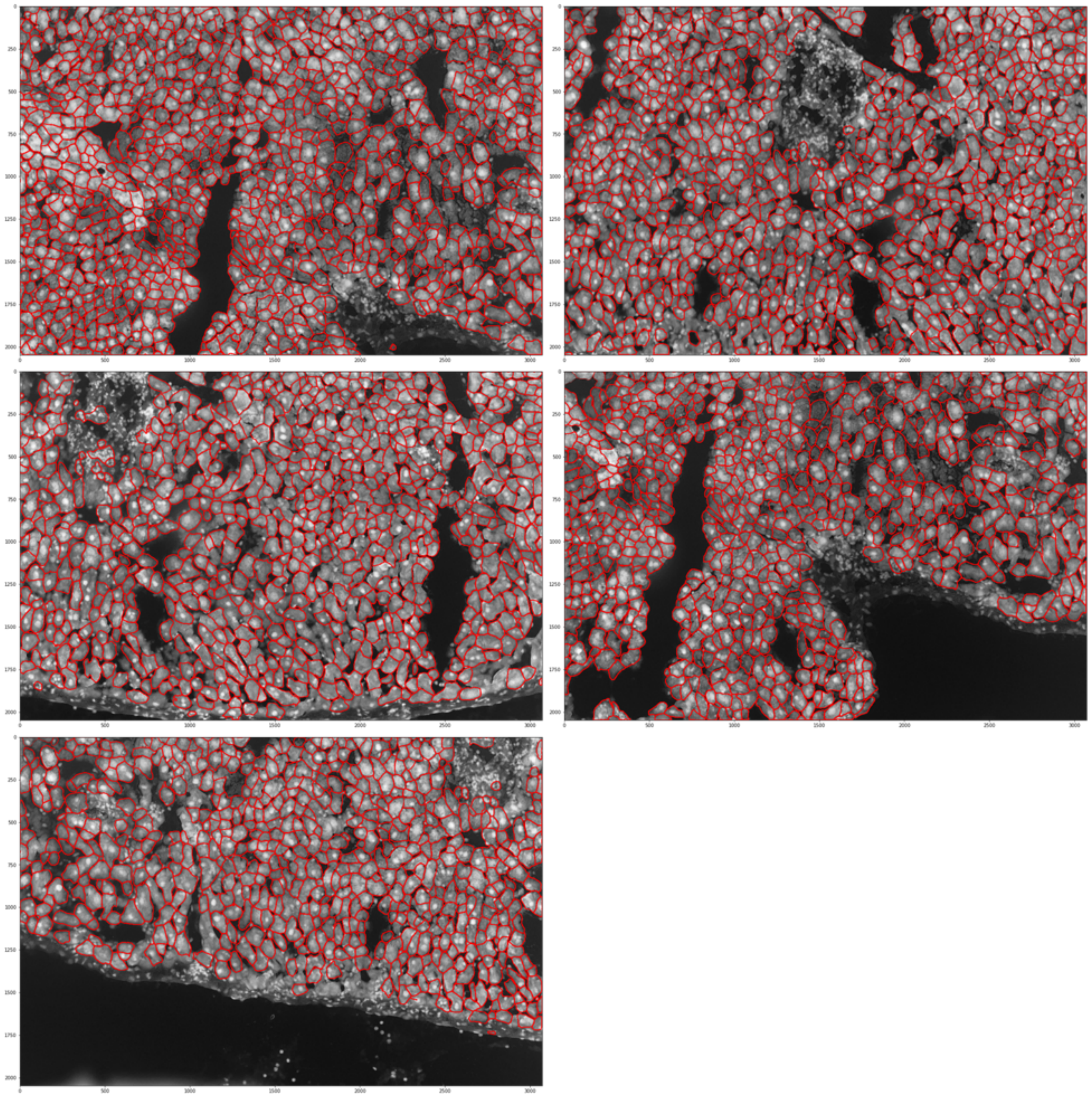

**Supplementary Figure 71. Aligned 10X H&E images for the normal tiles in Seq-Scope.** The 10X H&E images were manually aligned with the transcriptomics data. For each tile, the aligned H&E image (greyscale) is overlaid with the unspliced density (green) and the cell segmentation boundaries (red).

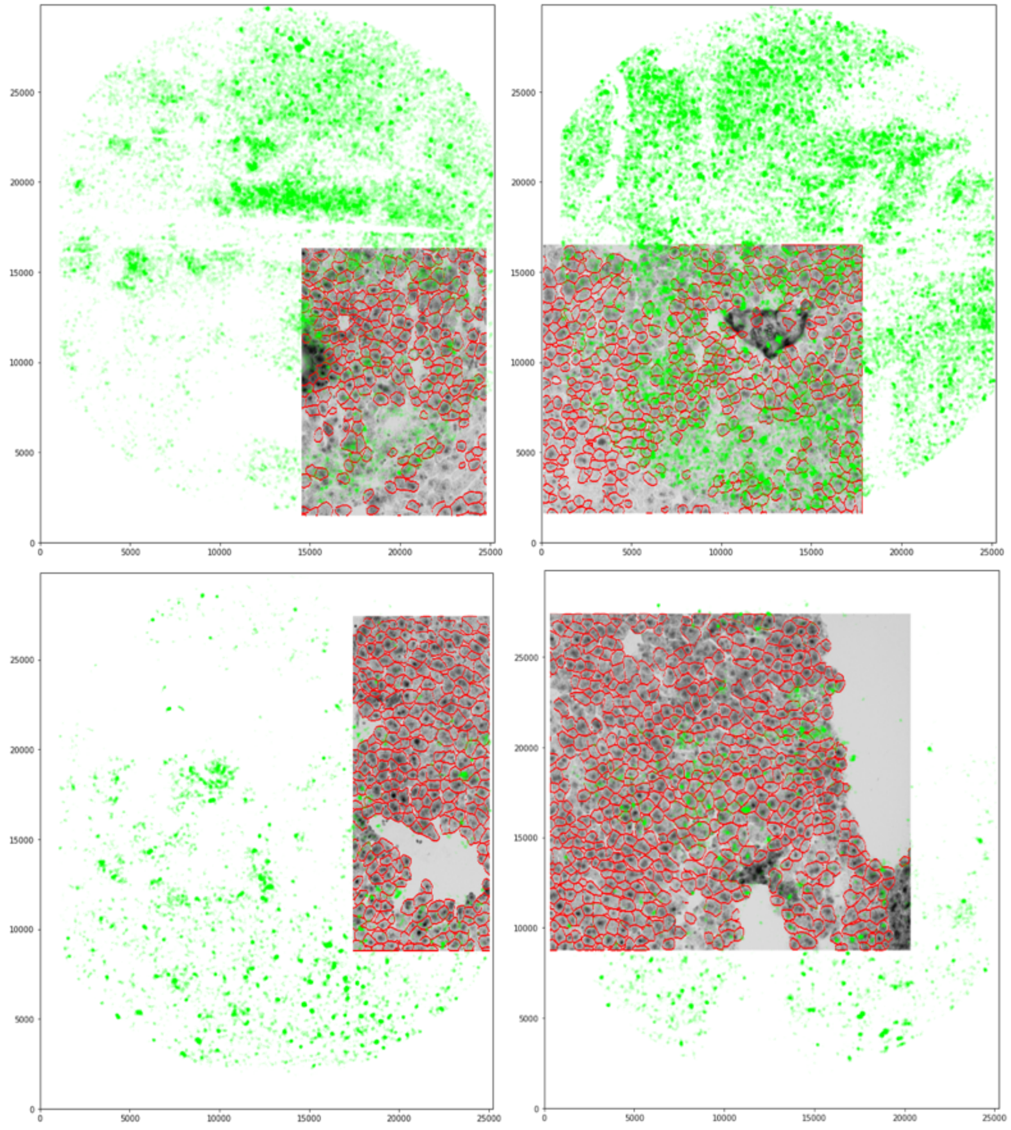

**Supplementary Figure 72. Aligned 10X H&E images for the TD tiles in Seq-Scope.** The 10X H&E images were manually aligned with the transcriptomics data. For each tile, the aligned H&E image (greyscale) is overlaid with the unspliced density (green) and the cell segmentation boundaries (red).

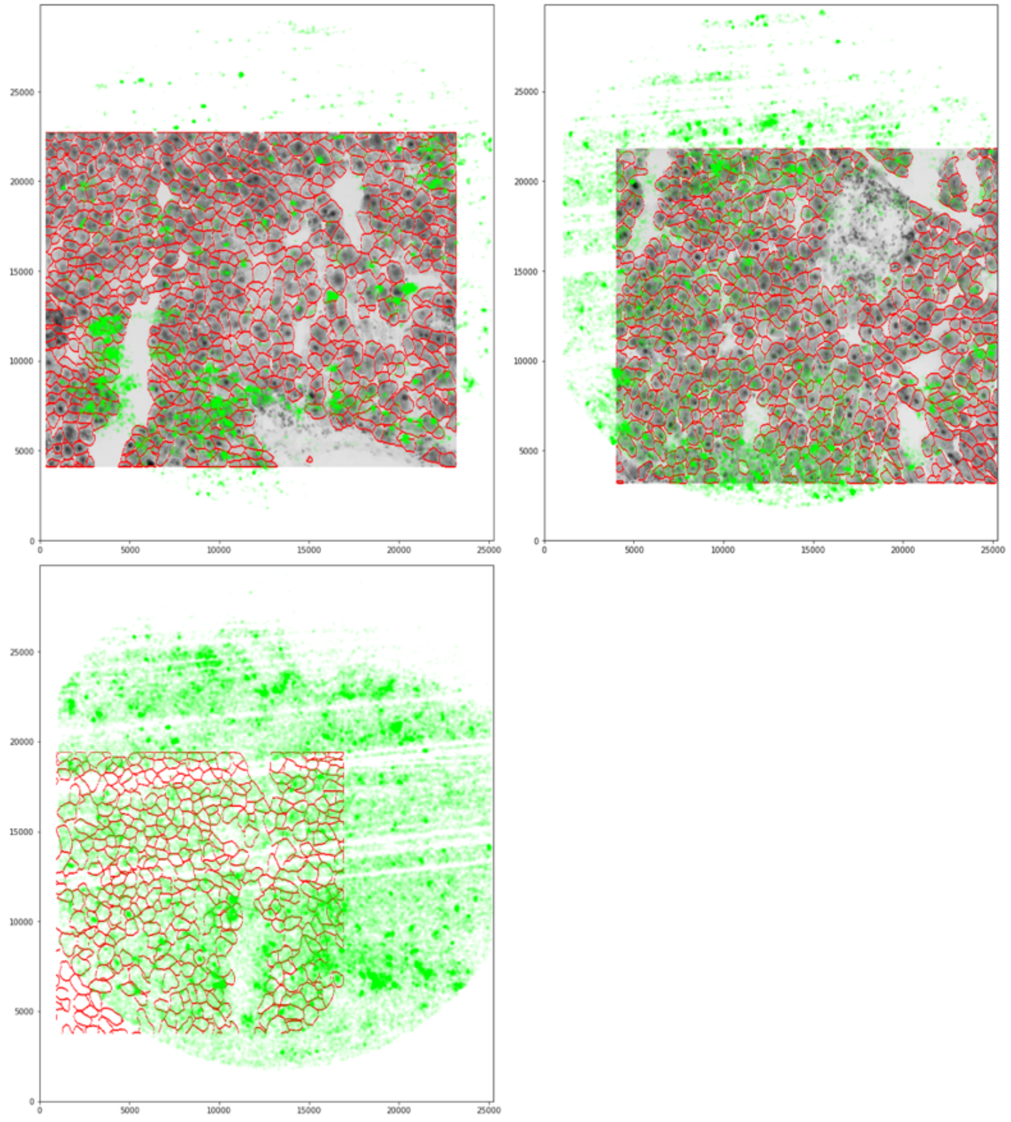

**Supplementary Figure 73. MERFISH human O2-US cell line data. a.** Cell boundary, nuclear boundary, and the expression of one randomly selected gene are displayed for randomly selected O2-US cells. **b.** Histogram of the average nucleus-cell ratio across cells. Dashed lines show the average nucleus-cell ratio and the cutoff ratios corresponding to two standard deviations from the average.

**a.**

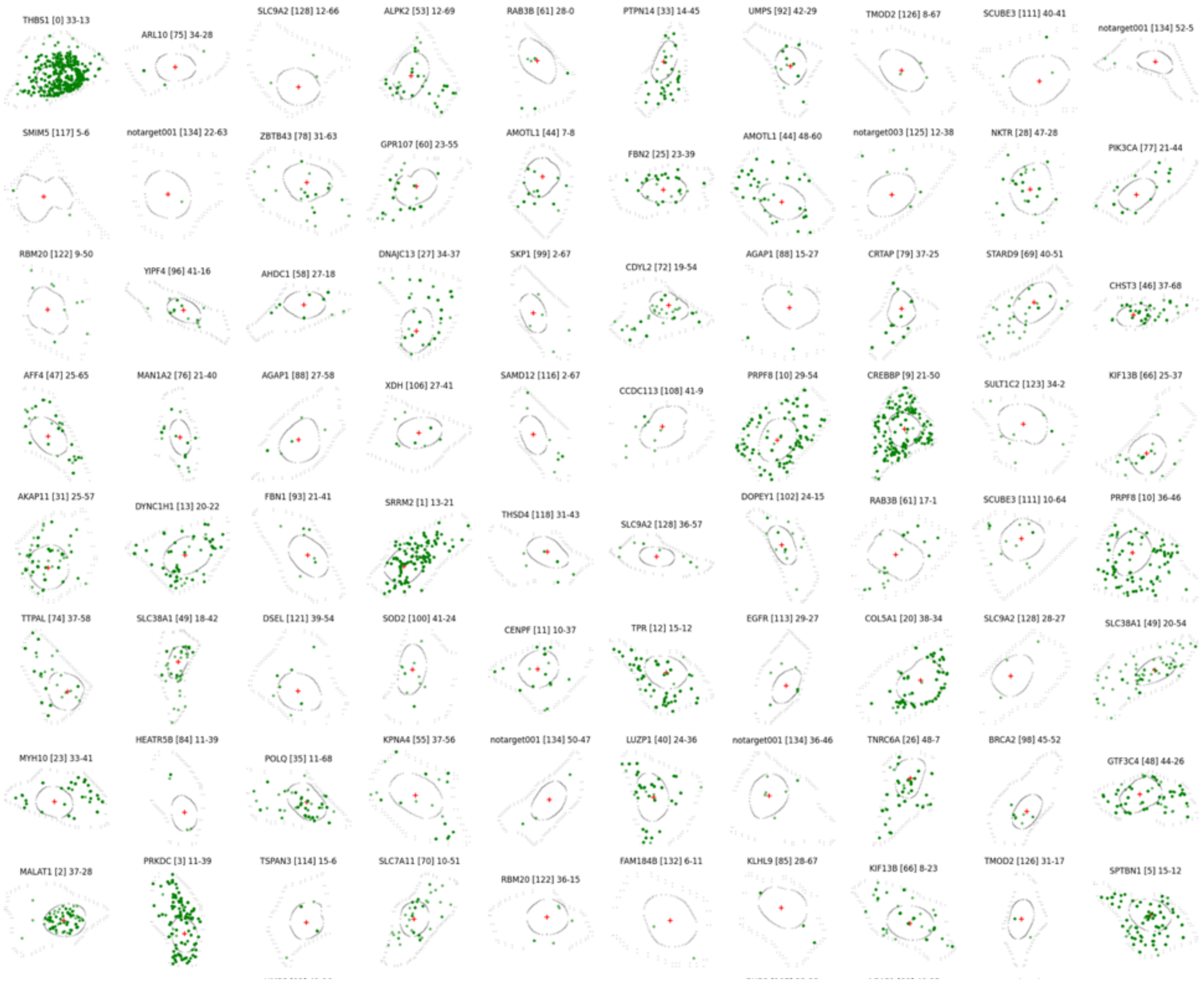

**b.**

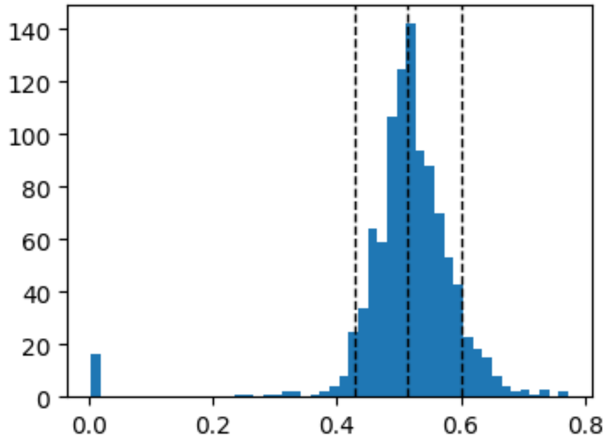

**Supplementary Figure 74. ELLA estimates of the MERFISH O2-US cell line data. a.** Estimated spatial expression pattern for genes in each of the seven gene pattern clusters identified by ELLA. Upper panel shows the number and proportion of genes across seven pattern clusters. Middle panel displays the estimated expression intensities for genes across clusters. Lower panel displays the estimated pattern score for genes across clusters. **b.** Example genes and cells for the seven pattern clusters. Upper panel shows the estimated expression intensities for seven genes, one from each pattern cluster. Lower panel displays the expressions of the corresponding genes within one selected cell, overlayed with cell boundary and nuclear center (cross).

**a.**

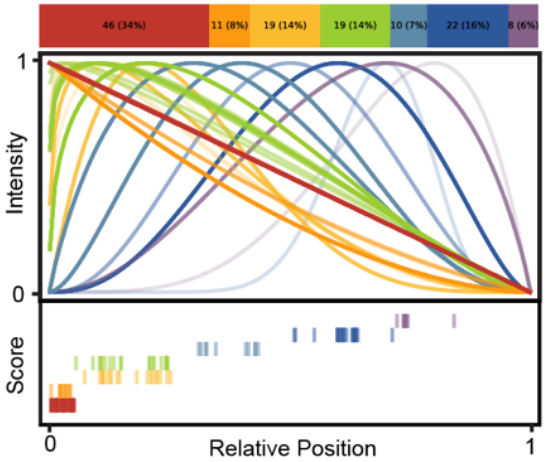

**b.**

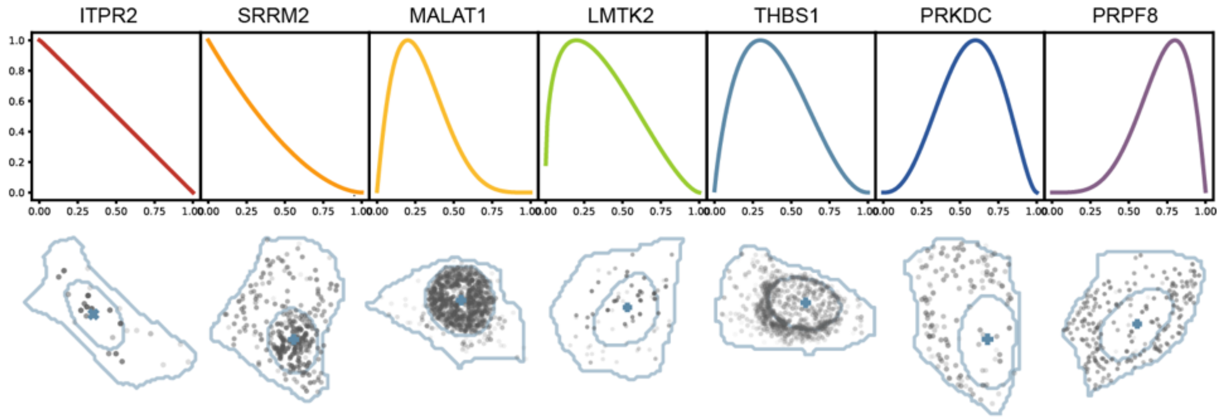

**Supplementary Figure 75. Cell by cell analysis on normal tile 2105 in Seq-Scope. a.** ELLA estimation for gene *Alb* across cells. The *Alb* expression (blue dots) is overlayed with the 10X H&E image (greyscale), cell centers (crosses), cell segmentations (red), and the best fit kernels are marked on each cell. **b.** In each subplot and gene, bar plot shows number of cells with significant spatial gene expression P<0.001 (green), number of cells with significant spatial gene expression P<0.05 and >0.001 (blue), and number of cells with nonsignificant spatial gene expression P>0.05 (grey), across 9 kernels.

**a.**

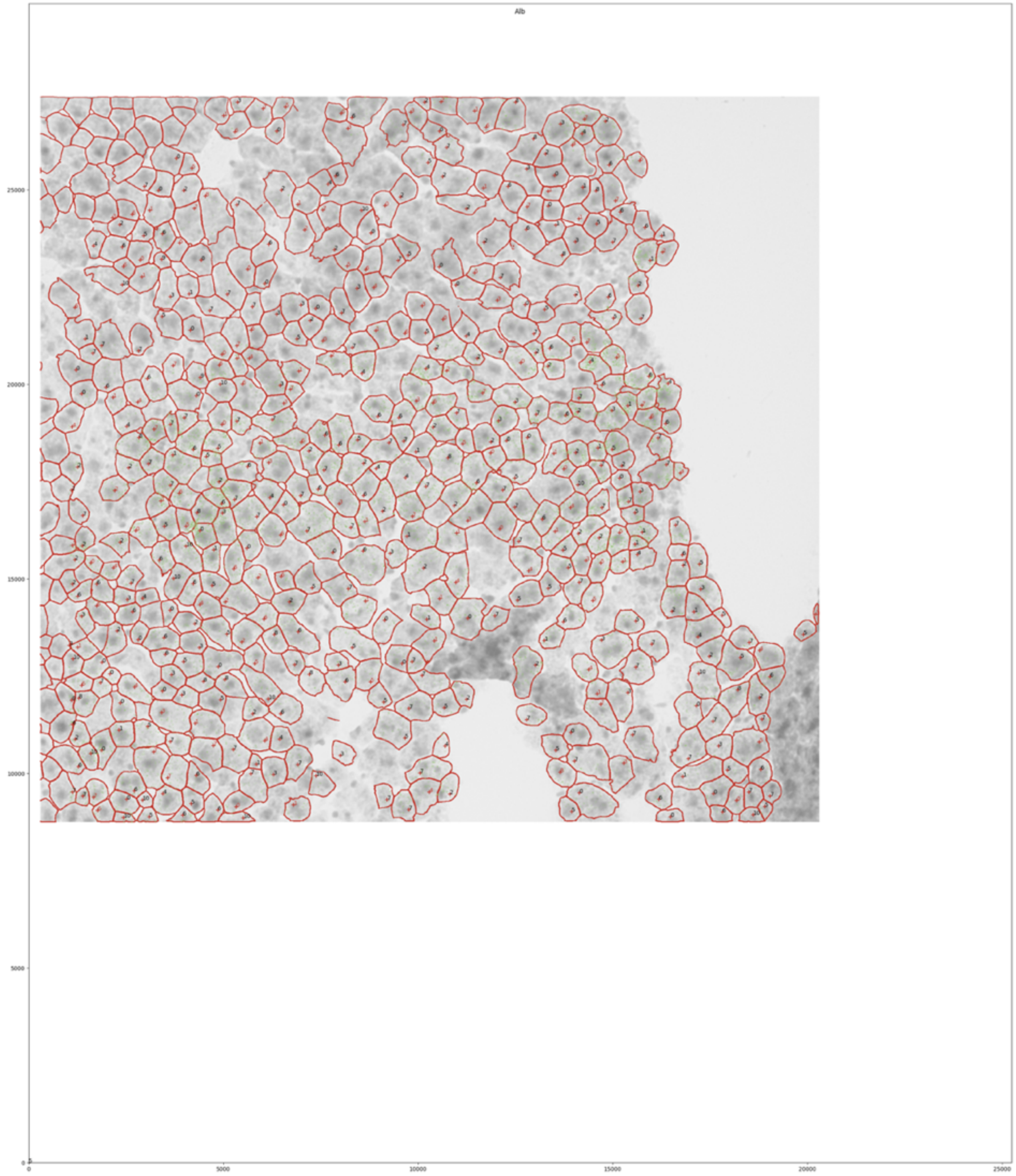

(Fig. S75 con'd)

**b.**

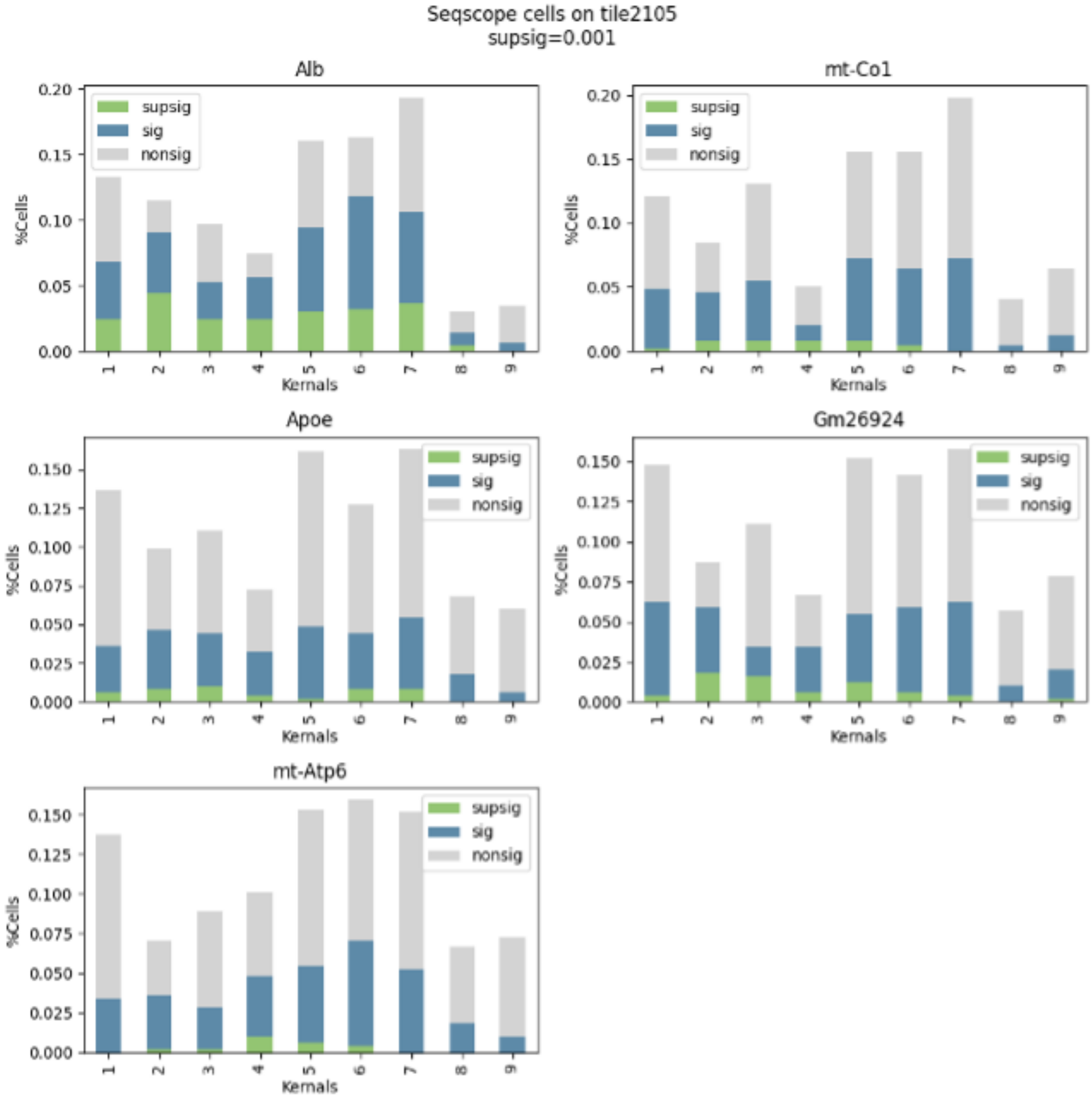

**Supplementary Figure 76. Cell by cell analysis on normal tile 2107 in Seq-Scope. a.** ELLA estimation for gene *Alb* across cells. The *Alb* expression (blue dots) is overlayed with the 10X H&E image (greyscale), cell centers (crosses), cell segmentations (red), and the best fit kernels are marked on each cell. **b.** In each subplot and gene, bar plot shows number of cells with significant spatial gene expression P<0.001 (green), number of cells with significant spatial gene expression P<0.05 and >0.001 (blue), and number of cells with nonsignificant spatial gene expression P>0.05 (grey), across 9 kernels.

**a.**

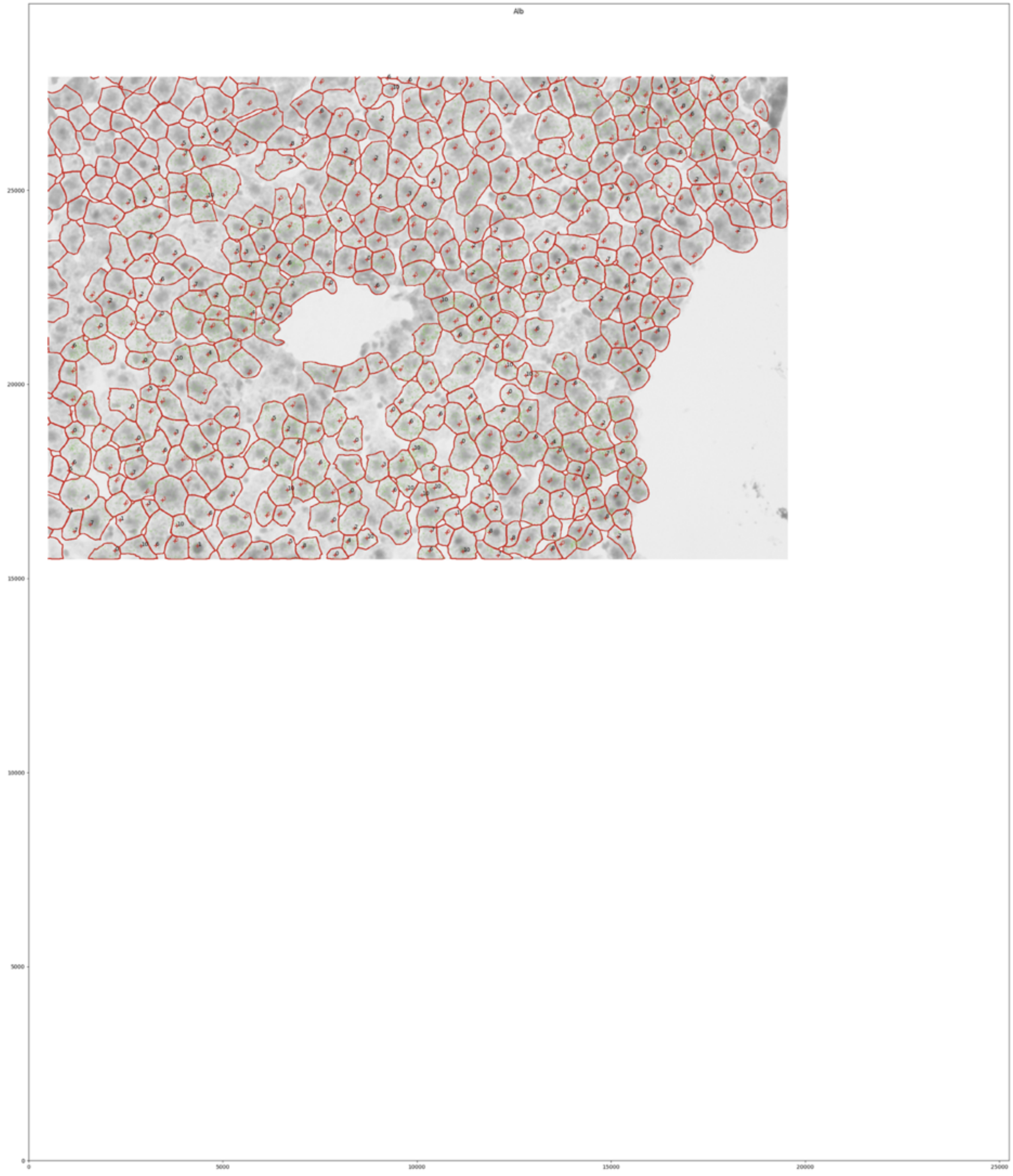

(Fig. S76 con'd)

**b.**

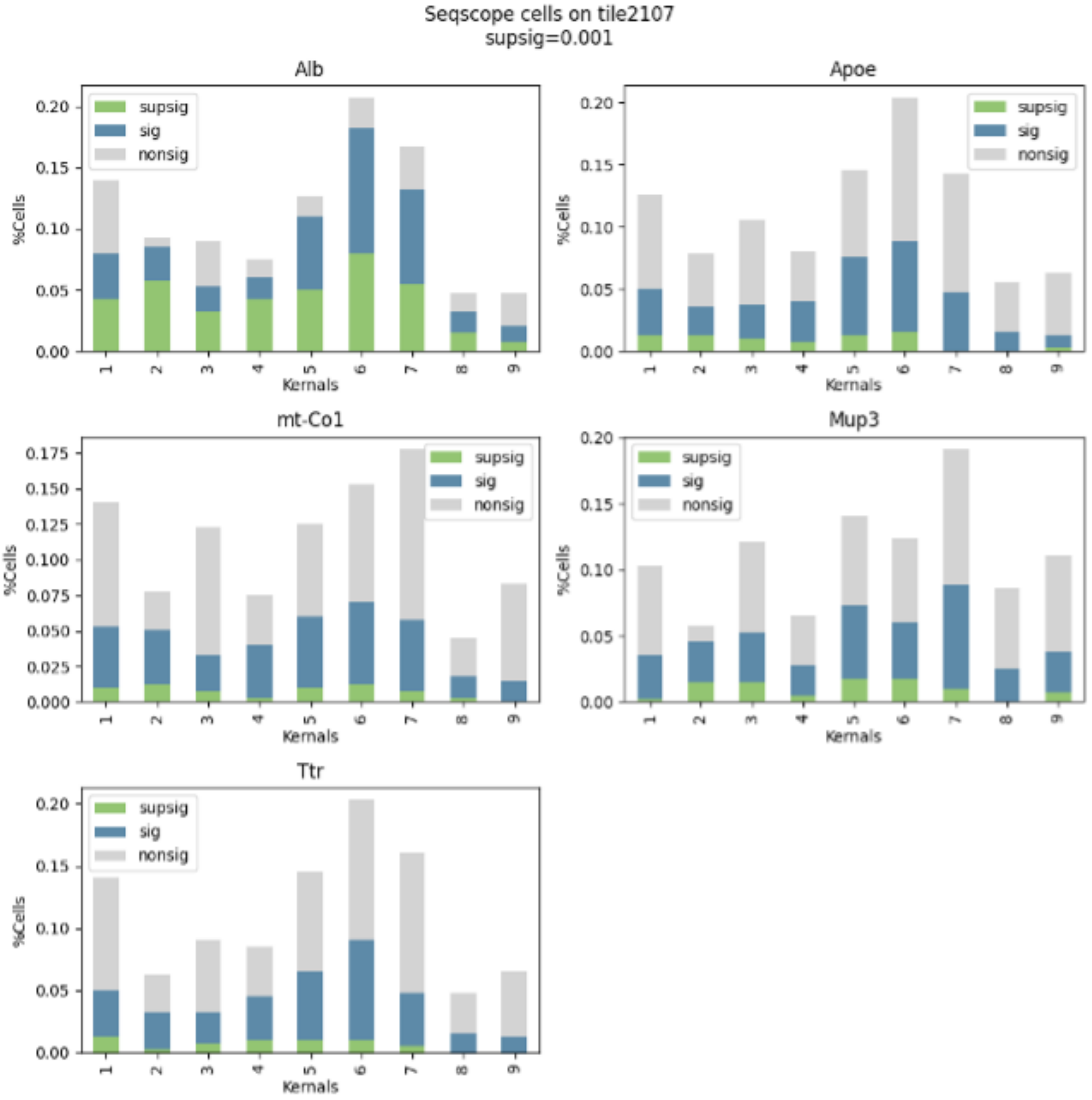

**Supplementary Figure 77. Cell by cell analysis on TD tile 2117 in Seq-Scope. a.** ELLA estimation for gene *Alb* across cells. The *Alb* expression (blue dots) is overlayed with the 10X H&E image (greyscale), cell centers (crosses), cell segmentations (red), and the best fit kernels are marked on each cell. **b.** In each subplot and gene, bar plot shows number of cells with significant spatial gene expression P<0.001 (green), number of cells with significant spatial gene expression P<0.05 and >0.001 (blue), and number of cells with nonsignificant spatial gene expression P>0.05 (grey), across 9 kernels.

**a.**

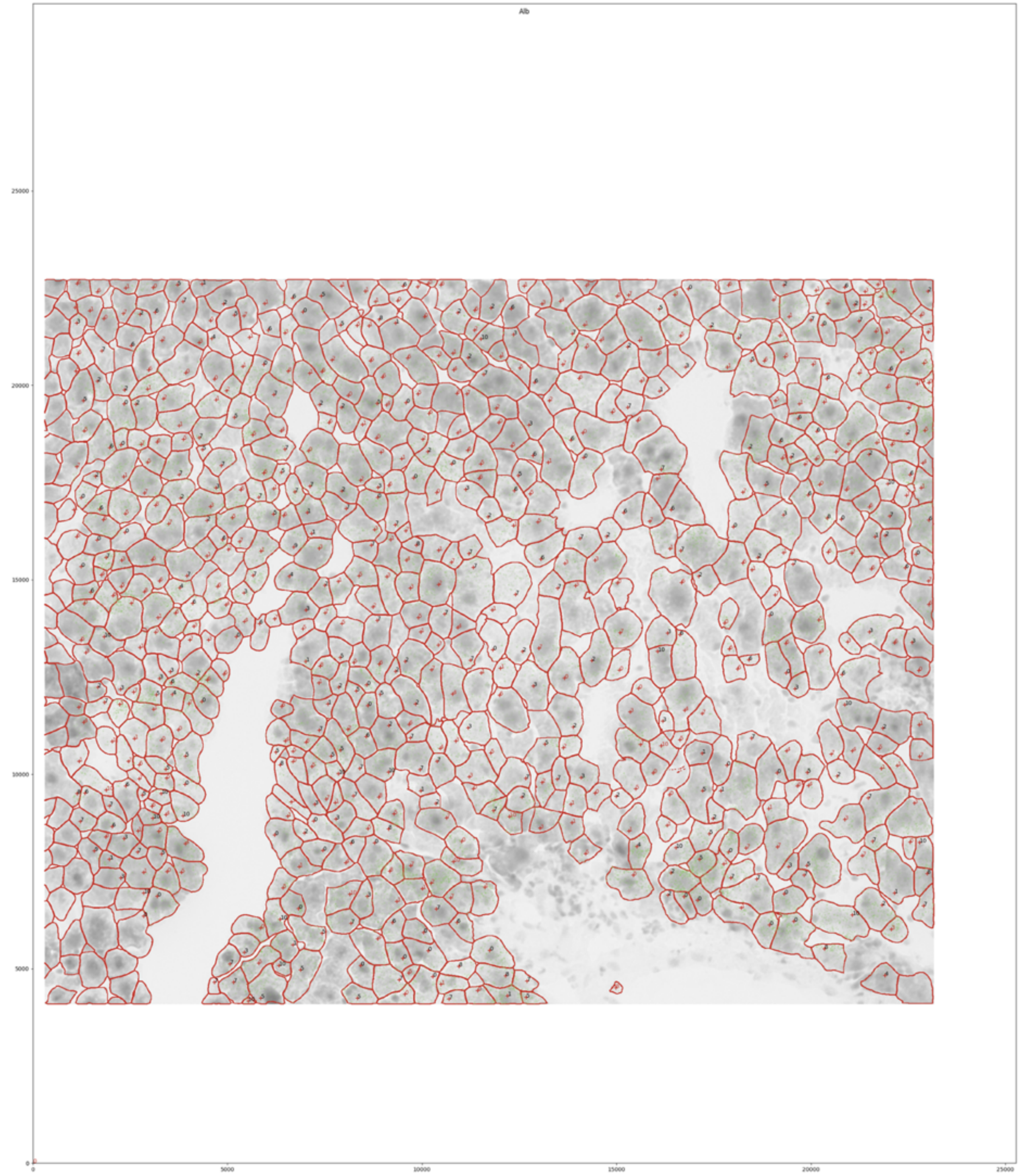

(Fig. S77 con'd)

**b.**

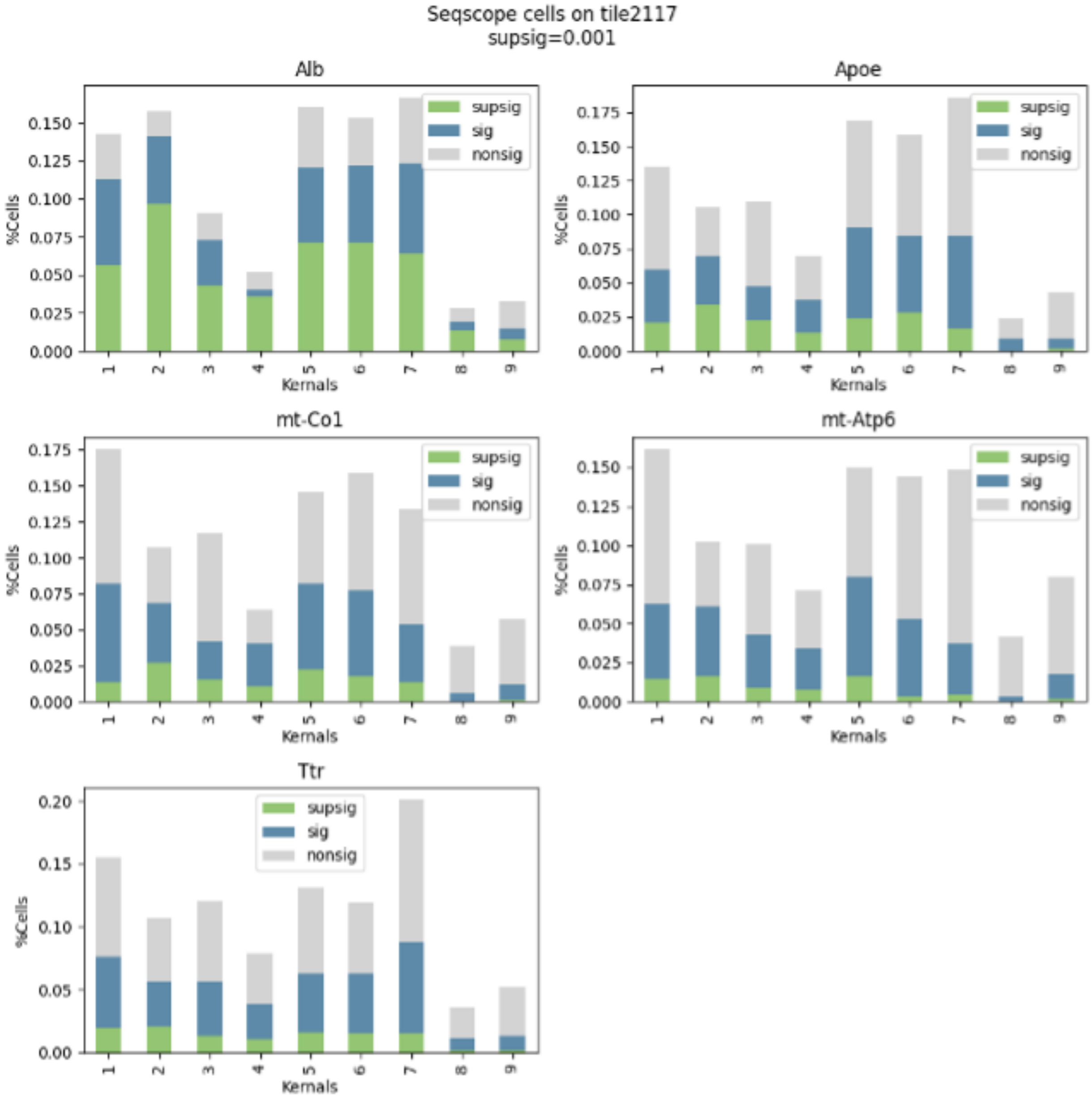

**Supplementary Figure 78. Cell by cell analysis on TD tile 2118 in Seq-Scope. a.** ELLA estimation for gene *Alb* across cells. The *Alb* expression (blue dots) is overlayed with the 10X H&E image (greyscale), cell centers (crosses), cell segmentations (red), and the best fit kernels are marked on each cell. **b.** In each subplot and gene, bar plot shows number of cells with significant spatial gene expression P<0.001 (green), number of cells with significant spatial gene expression P<0.05 and >0.001 (blue), and number of cells with nonsignificant spatial gene expression P>0.05 (grey), across 9 kernels.

**a.**

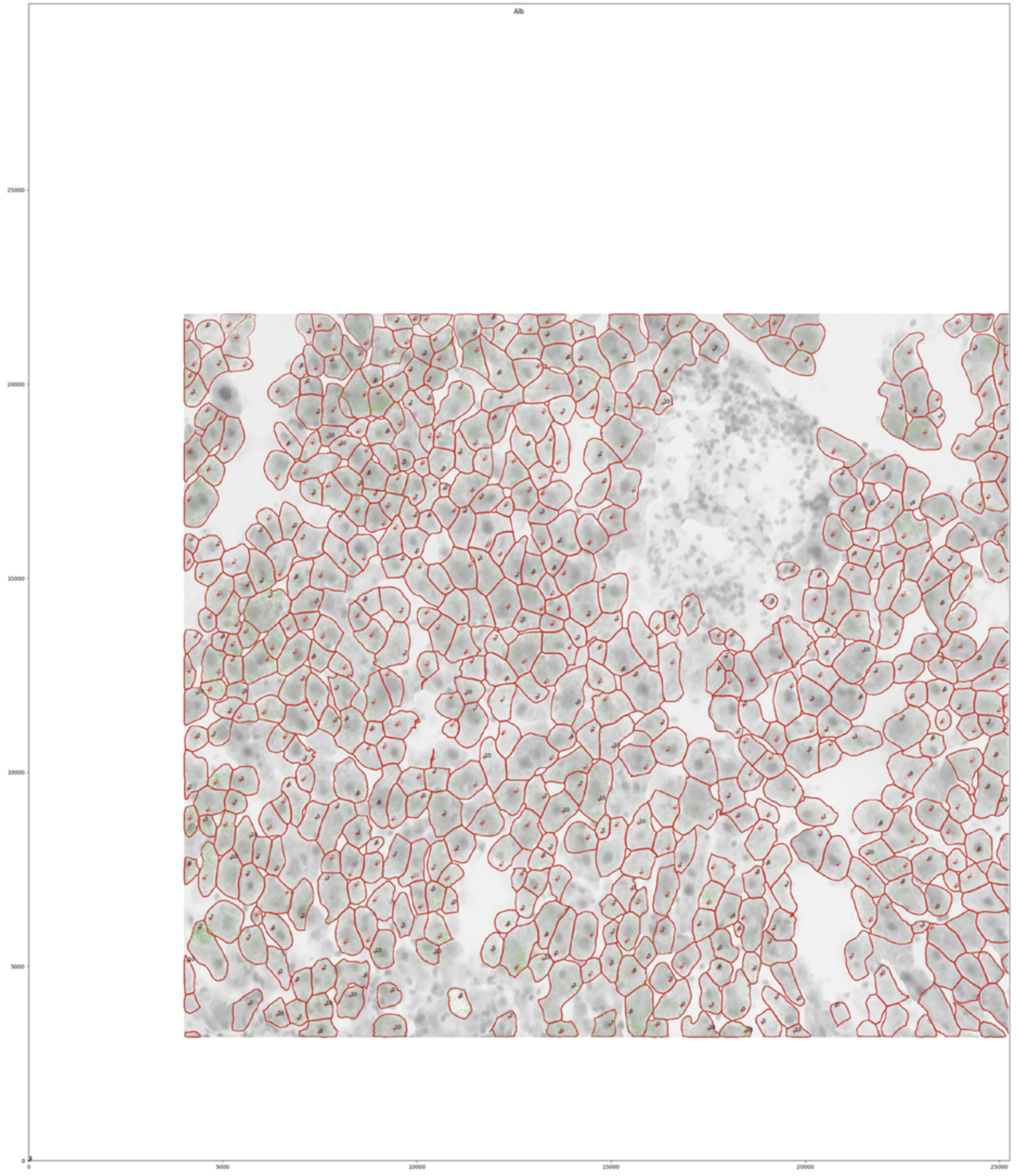

(Fig. S78 con'd)

**b.**

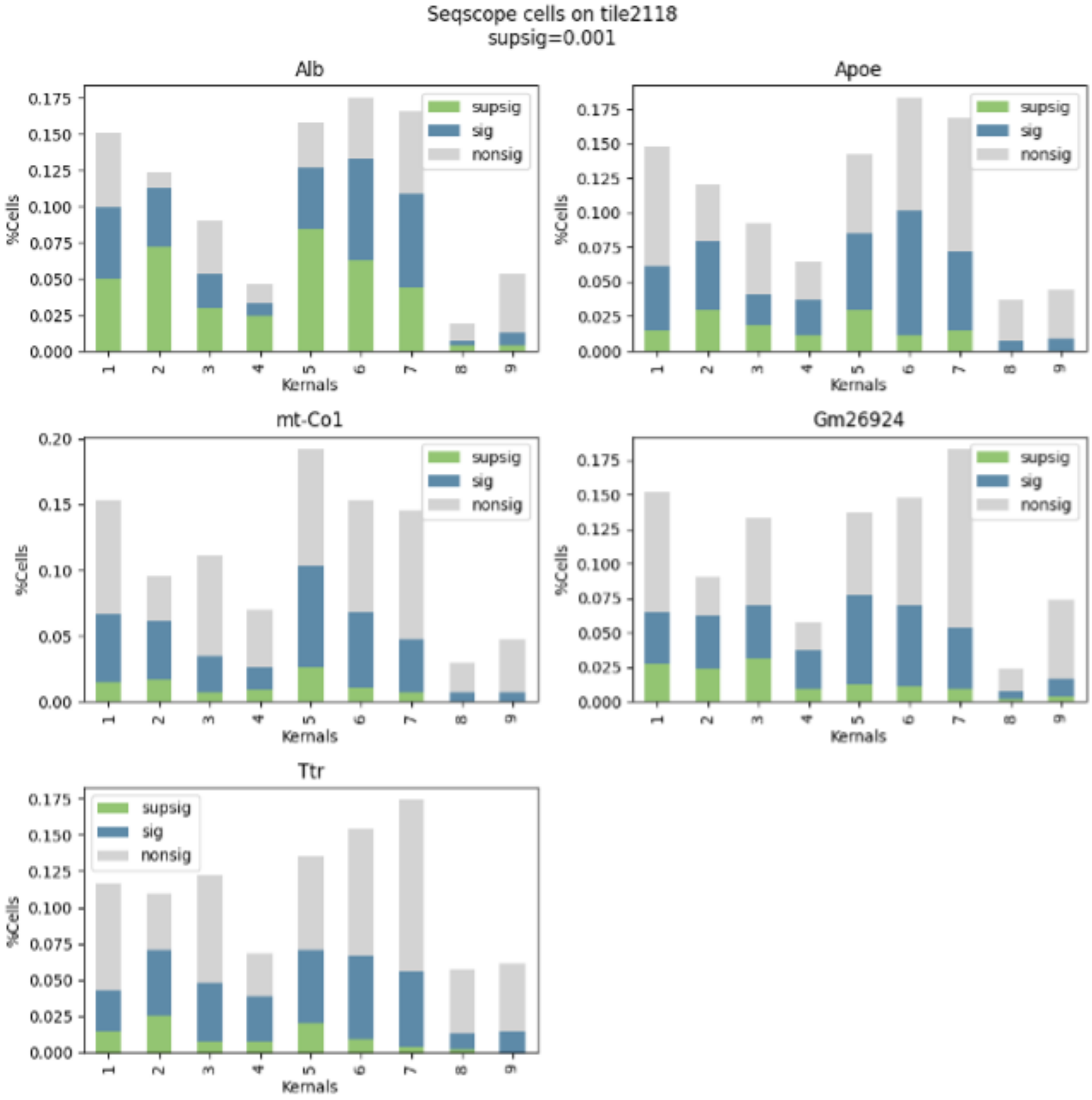

### Supplementary Tables

**Supplementary Table 1. Compatibility and required inputs for ELLA, SPRAWL, Bento, and Wilcox methods.**

| **Compatible with:** | **ELLA** | **SPRAWL** | **Bento** | **Wilcox** |
|---|---|---|---|---|
| One cell | ☑ (green) | ☒ (red) | ☑ (green) | ☒ (red) |
| Multiple cells | ☑ (green) | ☑ (green) | ☒ (red) | ☑ (green) |
| Imaging data | ☑ (green) | ☑ (green) | ☑ (green) | ☒ (yellow) |
| Sequencing data | ☑ (green) | ☒ (red) | ☒ (red) | ☒ (yellow) |
| P values | ☑ (green) | ☒ (yellow) | ☒ (red) | ☒ (yellow) |
| Number of patterns | Various | 4 | 5 | 1 |

☑ (green) : A method is compatible with a certain scenario.
☒ (red) : A method is not compatible with a certain scenario.
☒ (yellow) : A method is only compatible with a certain scenario under certain conditions.

| **Required inputs:** | **ELLA** | **SPRAWL** | **Bento** | **Wilcox** |
|---|---|---|---|---|
| Nuclear center | ☑ (green) | ☒ (red) | ☒ (red) | ☒ (red) |
| Nuclear boundary | ☒ (red) | ☒ (red) | ☑ (green) | ☑ (green) |
| Cell centroid | ☒ (red) | ☑ (grey) | ☒ (red) | ☒ (red) |
| Cell boundary | ☑ (green) | ☑ (grey) | ☑ (green) | ☑ (green) |

☑ (green) : An input is required for a certain method.
☑ (grey) : SPRAWL does not explicitly require cell centroid and boundary as inputs but infer them from the spatial expression data.
☒ (red) : An input is not required for a certain method.

**Supplementary Table 2. The heteroskedastic variances of the standardized nuclear and cytoplasmic expression counts used in the Wilcox method in the null simulations.** The average variances (across genes) of the standardized nuclear expression and the standardized cytoplasmic expression across null simulation settings. The standardized nuclear expression counts have smaller variances than the standardized cytoplasmic expression counts across settings.

| n | Nuclear Var | Cytoplasmic Var | m | Nuclear Var | Cytoplasmic Var |
|---|---|---|---|---|---|
| 10 | 4.36 | 1.16 | 1 | 1.96 | 0.52 |
| 20 | 4.37 | 1.16 | 2 | 2.69 | 0.76 |
| 50 | 4.37 | 1.16 | 5 | 4.29 | 1.19 |
| 100 | 4.29 | 1.19 | 10 | 6.05 | 1.68 |
| 200 | 4.35 | 1.16 | 20 | 8.68 | 2.33 |
| 300 | 4.36 | 1.16 | 50 | 13.77 | 3.69 |
| 500 | 4.27 | 1.19 | 100 | 18.98 | 5.38 |

**Supplementary Table 3. Accuracy of the estimated expression intensity in the baseline alternative simulations.** The average KL divergence between the standardized estimated intensities $\hat{\lambda}(r)$ and the true intensity across genes of the eleven symmetric baseline simulation patterns and the three asymmetric simulation patterns.

| Symmetric patterns | | Asymmetric patterns | |
|---|---|---|---|
| Pattern | KL divergence | Pattern | KL divergence |
| 1 | 0.04 | Radial uniform | 0.21 |
| 2 | 0.03 | Radial cytoplasmic | 0.03 |
| 3 | 0.06 | Punctate | 0.62 |
| 4 | 0.22 | | |
| 5 | 0.30 | | |
| 6 | 0.30 | | |
| 7 | 0.22 | | |
| 8 | 0.09 | | |
| 9 | 0.06 | | |
| 10 | 0.02 | | |
| 11 | 0.00 | | |
| Average: | 0.12 | Average: | 0.29 |

**Supplementary Table 4. Accuracy of the estimated pattern scores in the baseline alternative simulations.** The accuracy of the estimated pattern scores in the eleven symmetric baseline simulation patterns and the three asymmetric simulation patterns. An estimated pattern score of a gene is counted as an accurate estimate if its deviation from the true pattern score is no more than 0.1.

| **Symmetric patterns** | | **Asymmetric patterns** | |
|---|---|---|---|
| Pattern | Accuracy | Pattern | Accuracy |
| 1 | 40% | Radial uniform | NA |
| 2 | 66% | Radial cytoplasmic | 99% |
| 3 | 76% | Punctate | 98% |
| 4 | 90% | | |
| 5 | 95% | | |
| 6 | 94% | | |
| 7 | 97% | | |
| 8 | 99% | | |
| 9 | 76% | | |
| 10 | 96% | | |
| 11 | 93% | | |

**Supplementary Table 5. Accuracy of the estimated expression patterns and pattern scores in the additional simulations with one cell.** The average KL divergence between the standardized estimated intensities $\hat{\lambda}(r)$ and the true intensity across genes of the five simulation patterns. And the accuracy of the estimated pattern scores of the five simulation patterns. An estimated pattern score (of a gene) is counted as an accurate estimate if its deviation from the true pattern score is no more than 0.1.

| Pattern | KL divergence | Accuracy |
|---|---|---|
| 1 | 0.60 | 98% |
| 2 | 0.36 | 99% |
| 3 | 0.13 | 100% |
| 4 | 0.30 | 100% |
| 5 | 0.56 | 100% |

**Supplementary Table 6. The 34 nuclear genes (cluster 1) in normal PC cells identified by ELLA in Seq-Scope.**

| Gene ID | Gene type | Gene ID | Gene type |
|---|---|---|---|
| *Sord* | Protein coding | *Abcc2* | Protein coding |
| *Slco1b2* | Protein coding | *Abcb4* | Protein coding |
| *Apob* | Protein coding | *Chd9* | Protein coding |
| *Acox1* | Protein coding | *Mlxipl* | Protein coding |
| *Slc7a2* | Protein coding | *Malat1* | Lnc RNA |
| *n-R5-8s1* | rRNA | *Neat1* | Lnc RNA |
| *Ghr* | Protein coding | *Ccbl2* | Protein coding |
| *Errfi1* | Protein coding | *Rapgef4* | Protein coding |
| *Gm24601* | Protein coding | *Slc47a1* | Protein coding |
| *Cyp2c54* | Protein coding | *Echdc2* | Protein coding |
| *C6* | Protein coding | *Gckr* | Protein coding |
| *Egfr* | Protein coding | *Pitpnc1* | Protein coding |
| *Gm13775* | Lnc RNA | *Ppara* | Protein coding |
| *Mtss1* | Protein coding | *Ddx3x* | Protein coding |
| *Cpt1a* | Protein coding | *Mafb* | Protein coding |
| *Ppap2b* | Protein coding | *Nr1d1* | Protein coding |
| *Eci2* | Protein coding | *Phldb2* | Protein coding |

**Supplementary Table 7. Pattern clusters of the mitochondrial and cell type marker genes in normal PC cells from ELLA in Seq-Scope.**

| Normal PC mitochondrial genes | | Normal PC cell type marker genes | |
|---|---|---|---|
| Gene ID | Cluster | Gene ID | Cluster |
| *mt-Co1* | 3 | *Mup17* | 5 |
| *mt-Atp6* | 3 | *Cyp2c29* | 3 |
| *mt-Nd4* | 3 | *Cyp2e1* | 3 |
| *mt-Cytb* | 3 | | |
| *mt-Co2* | 3 | | |
| *mt-Co3* | 3 | | |
| *mt-Rnr2* | 3 | | |
| *mt-Nd3* | 3 | | |
| *mt-Rnr1* | 3 | | |

**Supplementary Table 8. Shape parameters of the 22 default Beta kernel functions in ELLA.** The 22 kernel functions include two sets (each contains 11) of Beta pdfs where Set 1 corresponds to strong patterns and Set 2 corresponds to less strong patterns (Fig. S41).

| Set 1 | | | Set 2 | | |
|---|---|---|---|---|---|
| Mode | $a_0$ | $b_0$ | Mode | $a_0$ | $b_0$ |
| 0.0 | 1 | 2.71 | 0.0 | 1 | 2 |
| 0.1 | 1.26 | 3.34 | 0.1 | 1.13 | 2.19 |
| 0.2 | 2.05 | 5.19 | 0.2 | 1.38 | 2.52 |
| 0.3 | 6.99 | 14.98 | 0.3 | 1.88 | 3.06 |
| 0.4 | 19.41 | 28.62 | 0.4 | 2.73 | 3.60 |
| 0.5 | 28.50 | 28.50 | 0.5 | 3.50 | 3.50 |
| 0.6 | 28.62 | 19.41 | 0.6 | 3.60 | 2.73 |
| 0.7 | 14.98 | 6.99 | 0.7 | 3.06 | 1.88 |
| 0.8 | 5.19 | 2.05 | 0.8 | 2.52 | 1.38 |
| 0.9 | 3.34 | 1.26 | 0.9 | 2.19 | 1.13 |
| 1.0 | 2.71 | 1 | 1.0 | 2 | 1 |

**Supplementary Table 9. A summary of all the simulation settings.**

**Null simulations**

| | | |
|---|---|---|
| Baseline setting | n=100, m=5 | 1 setting |
| Varying n | n=10, 20, 50, 200, 300, 500 | 6 settings |
| Varying m | m=1, 2, 10, 20, 50, 100 | 6 settings |
| | | **Total: 13 settings** |

**Alternative simulations, across 11 symmetric patterns**

| | | | | |
|---|---|---|---|---|
| Baseline setting | n=100, m=5, s=0.6 | $\alpha$=max $\varphi(r)$ | $\beta$=s | 1 setting |
| Varying n | n=10, 20, 50, 200, 300, 500 | $\alpha$=max $\varphi(r)$ | $\beta$=s | 6 settings |
| Varying m | m=1, 2, 10, 20, 50, 100 | $\alpha$=max $\varphi(r)$ | $\beta$=s | 6 settings |
| Varying s | s=0.1, 0.2, …, 1.0 | $\alpha$=max $\varphi(r)$ | $\beta$=s | 9 settings |
| | | | | **Total: 22 settings** |

**Alternative simulations, other pattern scenarios**

| | |
|---|---|
| Radial uniform pattern | 1 setting |
| Radial cytoplasmic pattern | 1 setting |
| Punctate pattern | 1 setting |
| | **Total: 3 settings** |

**Additional alternative simulations with one cell**

| | | |
|---|---|---|
| n=1, m=30, s=9 | Pattern 1 | 1 setting |
| | Pattern 2 | 1 setting |
| | Pattern 3 | 1 setting |
| | Pattern 4 | 1 setting |
| | Pattern 5 | 1 setting |
| | | **Total: 5 settings** |

**Supplementary Table 10. Cell type marker genes in Seq-Scope.**

| Cell type | Marker genes |
|---|---|
| **PC** | *Glul, Oat, Cyp2a5, Mup9, Mup17, Cyp2c29, Cyp2e1* |
| **PP** | *Mup20, Alb, Cyp2f2, Serpina1e, Ass1, Hamp, Mup11* |
| **HSC-N** | *Ecm1, Dcn, Sod3, Prelp* |
| **HSC-A** | *Col3a1, Col1a1, Col1a2, Acta2* |
| **M$\varphi$-Kupffer** | *Clec4f, Cd5l, Marco, C1qc* |
| **M$\varphi$-Inflamed** | *Cd74, H2-Aa, H2-Ab1, H2-Eb1* |
| **Hep-Injured** | *Saa1, Saa2, Hp, Lrg1* |
| **HPC** | *Spp1, Mmp7, Clu, Epcam* |

**Supplementary Table 11. Mouse embryo cell type marker genes in Stereo-seq.**

| Cell type | Marker genes |
|---|---|
| **Cardiomyocyte** | *Myl2, Myh7, Tnnt2* |
| **Chondrocyte** | *Col2a1, Col9a1, Col11a, Runx2* |
| **Choroid plexus** | *Ttr, Enpp2, Igfbp2* |
| **Dorsal midbrain neuron** | *Tfap2b, Lhx9, Zic1* |
| **Ganglion** | *Nefl, Nefm, Sncg* |
| **Endothelial cell** | *Pecam1, Kdr, Ptprm* |
| **Keratinocyte** | *Krt4, Krtdap, Krt10* |
| **Epithelial cell** | *Krt19, Epcam, Krt8, Krt5, Adh1, Foxp2* |
| **Erythrocyte** | *Hba-a2, Hba-a1, Hbb-bs* |
| **Facial fibroblast** | *Trps1, Pax3, Wnt5a* |
| **Fibroblast** | *Col1a2, Col3a1, Dcn* |
| **Forebrain neuron** | *Neurod6, Tbr1, Neurod2* |
| **Forebrain radial glia** | *Fabp7, Sox2, Pou3f3* |
| **Hepatocyte** | *Afp, Alb* |
| **Immune cell** | *S100a8, S100a9* |
| **Limb fibroblast** | *Mecom, Gas2, Ebf1* |
| **Macrophage** | *Mrc1, C1qc, Csf1r* |
| **Meninges cell** | *Ptgds, Trpm3, Ranbp3l* |
| **Mid-/hindbrain and spinal cord neuron** | *Rtn1, Nnat, Stmn2* |
| **Myoblast** | *Acta1, Myl1, Myh3* |
| **Olfactory epithelial cell** | *Gstm1, Ebf2, Fstl5* |
| **Radial glia** | *Fabp7, Sox2, Slc1a3* |
| **Smooth muscle cell** | *Acta2, Myh11, Tagln* |
| **Spinal cord neuron** | *Npy, Cck, Lingo2* |
| **Diencephalon neuron** | *Tcf712, Ntng1, Tenm2* |

**Supplementary Table 12. Adult mouse brain cell type marker genes in Stereo-seq.**

| Cell type | Marker genes |
|---|---|
| EX | *Rprm*, *Myl4*, *Rorb*, *Cux2*, *Lamp5*, *Slc17a7* |
| IN | *Reln*, *Pvalb*, *Gad2* |
| Astr | *Sox9*, *Aqp4*, *Gfap*, *Mfge8*, *Agt* |
| Oligo | *Sox10* |
| Others | *Cldn5*, *Acta2*, *Tcf7l2*, *Prkcd*, *Prox1*, *Th*, *Slc17a6* |

**Supplementary Table 13. All mitochondria genes measured in Seq-Scope and a list of 10 nuclear genes presented in the Seq-Scope study.**

| | |
|---|---|
| **Mitochondria genes** | *mt-Tf, mt-Rnr1, mt-Tv, mt-Rnr2, mt-Tl1, mt-Nd1, mt-Ti, mt-Tq, mt-Tm, mt-Nd2, mt-Tw, mt-Ta, mt-Tn, mt-Tc, mt-Ty, mt-Co1, mt-Ts1, mt-Td, mt-Co2, mt-Tk, mt-Atp8, mt-Atp6, mt-Co3, mt-Tg, mt-Nd3, mt-Tr, mt-Nd4l, mt-Nd4, mt-Th, mt-Ts2, mt-Tl2, mt-Nd5, mt-Nd6, mt-Te, mt-Cytb, mt-Tt, mt-Tp* |
| **10 nuclear genes presented in the Seq-Scope study** | *Neat1, Malat1, Mlxipl, n-R5-8s1, Gm24601, Echdc2, D4Wsu53e, Aspg, Mafb, Sema4g* |

**Supplementary Table 14. Three lists of localized genes in the seqFISH+ study.**

| | |
|---|---|
| **20 nuclear or nuclear edge genes** | *Col1a1, Fn1, Fbln2, Col6a2, Bgn, Nid1, Lox, P4hb, Aebp1, Emp1, Col5a1, Sdc4, Postn, Col3a1, Pdia6, Col5a2, Itgb1, Calu, Pdia3, Cyr61* |
| **20 cytoplasmic genes** | *Ddb1, Myh9, Actn1, Tagln2, Kpnb1, Hnrnpf, Ppp1ca, Hnrnpl, Pcbp1, Tagln, Fscn1, Psat1, Cald1, Snd1, Uba1, Hnrnpm, Cap1, Ssrp1, Ugdh, Caprin1* |
| **20 protrusion genes** | *Cyb5r3, Sh3pxd2a, Ddr2, Net1, Trak2, Kif1c, Kctd10, Dynll2, Arhgap11a, Gxylt1, H6pd, Gdf11, Dync1li2, Palld, Ppfia1, Naa50, Ptgfr, Zeb1, Arhgap32, Scd1* |

### Supplementary Notes

#### 1. 3'UTR analysis in the Stereo-seq data

We used a modified version of ReadZS [3] provided by SPRAWL [4], which we refer to as ReadZS-SPRAWL, to measure 3' UTR length in the Stereo-seq data. Specifically, we applied ELLA to 25 cell types on 90,974 cells on E1S3, with 195 to 2,892 genes measured per cell type (Methods). We focused the analysis on 435 genes that are detected by ELLA in no less than 20 out of the 25 cell types. For each selected gene, we applied ReadZS-SPRAWL to quantify its 3' UTR length with ReadZS score across physical locations on the slice and extracted its median ReadZS score for each cell type. For each gene in turn, in the cell types where the gene is detected by ELLA, we applied the Kruskal-Wallis H test on the 3'UTR lengths across patterns 1-5, whereas we skipped the patterns where less than three lengths were available. In the analysis, we detected 19 genes that have significantly varying 3'UTR lengths across subcellular localization patterns. We computed the Pearson correlation between 3'UTR length and ELLA pattern strength across available cell types, yielding 21 genes with significant correlation. We also computed the Pearson correlation between 3'UTR length and ELLA pattern scores across available cell types, yielding 18 genes with significant correlation.

#### 2. Derivation of the estimation weights in ELLA

Here, we provide details for the derivation of the estimation weights for the estimated intensity functions in ELLA through Bayesian model averaging. Specifically, let $\Lambda_l$ denote the space of functions that contains all potential $\lambda_l(r)$ in the form of linear transformed $\varphi_l(r)$ i.e., $\lambda_l(r) = \alpha_l + \beta_l\varphi_l(r)$ with arbitrary scalars $\alpha_l$ and $\beta_l$. Then the weights are in the form of

$$w_l = \frac{\sum_{\lambda_l(r)\in\Lambda_l} L(D|\lambda_l(r), M_l)P(\lambda_l(r)|M_l)}{\sum_{j=1}^{k}\left[\sum_{\lambda_j(r)\in\Lambda_j} L(D|\lambda_j(r), M_j)P(\lambda_j(r)|M_j)\right]}.$$

We approximate the summation $\sum_{\lambda_l(r)\in\Lambda_l}\cdot$ using $\max_{\lambda_l(r)\in\Lambda_l}\cdot$, as the summation is dominated by the largest summand when the sample size is large. We let $P\big(\lambda_j(r)\big|M_j\big), j = 1,\ldots,k$ all equal, corresponding to equal prior probability. Therefore,

$$w_l = \frac{\max_{\lambda_l(r)\in\Lambda_l} L(D|\lambda_l(r), M_l)}{\sum_{j=1}^{k}\left[\max_{\lambda_j(r)\in\Lambda_j} L(D|\lambda_j(r), M_j)\right]},$$

where $\max_{\lambda_l(r)\in\Lambda_l} L(D|\lambda_l(r), M_l)$ is the maximized likelihood value based on kernel function $\varphi_l(r)$. With the weights, we can obtain $\hat{\lambda}(r)$ as the final estimated subcellular spatial expression intensity function for the gene of focus in a given cell type.

#### 3. Seq-Scope data preprocess

##### 3.1 Unspliced and spliced expression

We followed the instructions in the Seq-Scope study to extract the unspliced and spliced gene expression counts using STtools based on STARsolo's velocyto (details in the Seq-Scope github page: https://github.com/jyxi7676/STtools). We merged the unspliced/spliced counts for all the genes at each location across tiles.

##### 3.2 H&E image segmentation

We obtained cell boundary segmentations from the 4X H&E image provided by Seq-Scope using Cellpose [5]. To do so, we manually concatenated all images for the normal and TD tiles separately, removed abnormal regions (defined as regions with extremely high or low intensities, such as bubbles and edges) as well as regions with no tiles in the concatenated images, and applied Cellpose to the two concatenated images (one for normal tissue and one for TD tissue).

##### 3.3 Alignment of the H&E image with the spatial transcriptomics data

We aligned the H&E images with the spatial transcriptomics data and transferred the cell segmentation information obtained from the H&E images to the transcriptomics coordinate space. To do so, we first transformed the H&E images using affine transformation, where we obtained an initial transformation using the template matching algorithm in python based on tile 2107 and fine-tuned the parameters for the other tiles based on that of tile 2107. The intensity of the H&E images allows us to visualize cellular morphology on the tissue. On the other hand, we obtained the total unspliced counts in the spatial transcriptomics data, which allows us to label the cell nuclei. Afterwards, we manually aligned the H&E images with the spatial gene expression data based on the cell morphology in the H&E images and the cellular nucleus in spatial transcriptomics.

After alignment, we transferred the cell segmentations to the spatial transcriptomics coordinate space. We retained gene expression counts that are assigned either to a specific cell or to the background. For each cell in turn, we allocated its nuclear center to the location with the maximum total unspliced expression intensity (kernel size=201) within the region that is 200 units away from the boundary. We removed cells whose i) unspliced intensity at the detected center was below the 95% quantile across locations of the tile or ii) or spliced intensity at the detected center was above the 95% quantile across locations of the tile.

##### 3.4 Cell type annotation

We obtained lists of cell type marker genes from the Seq-Scope study ([6], Tab. S10). We computed for each cell its total counts of marker genes and obtained cell type annotations (details in the Methods). The annotated cells display spatial patterns that are largely agreed with the Seq-Scope study. We removed cells with extreme widths or heights (details in the Methods).

#### 4. Stereo-seq data preprocess

##### 4.1 Unspliced and spliced expression

We used STARsolo velocyto to obtain the unspliced and splice expression matrix of the E1S3 slice. The STARsolo arguments were chosen based on SAW (https://github.com/STOmics/SAW) and the description in the Stereo-seq manuscript [7].

##### 4.2 Nucleic acid staining image segmentation

We examined the cell segmentations provided by the Stereo-seq study and found many merged cells incorrectly identified as single cells. Therefore, we applied Cellpose to the nucleic acid staining image to obtain cell segmentations. We split the full E1S3 nucleic acid staining image (2.58 GB) into eight subregions and applied Cellpose to each subregion in parallel to reduce memory consumption and reduce computational time.

##### 4.3 Assign spatial gene expression counts to cells

The nucleic acid staining image and the transcriptomics data have already been aligned. Therefore, we transferred the cell segmentations to the transcriptomics data (the bin1 data provided by Stereo-seq) directly to obtain cells in each of the eight subregions in a similar way as the Seq-Scope preprocessing steps described earlier. We assigned the nuclear centers to the location with the maximum unspliced intensity. We filtered out cells with a center that is less than 2 units from the cell boundary. We filtered out nuclear centers with unspliced intensity less than 90% quantile or spliced intensity greater than 90% quantile across all locations.

##### 4.4 Cell type annotation

We carried out cell type annotation on cells across the eight subregions in a similar way as described in the Seq-Scope preprocessing steps. The annotation was based on the cell type marker genes of 25 cell types provided by Stereo-seq (Tab. S11). We excluded cells with extreme widths or heights and cells whose nuclear center is too close to its cell boundary (details in the Methods).

#### 5. MERFISH human osteosarcoma data analysis

We analyzed a human U2-OS osteosarcoma cell line data generated by MERFISH [1], which contains 130 genes measured on 887 U2-OS cells.

##### 5.1 Preprocessing

The MERFISH U2-OS cell line data was generated in Bento (details in [1]). Nucleus and cell segmentations are provided by Bento in the form of a set of points with x and y coordinates on the nuclear boundary and a set of points with x and y coordinates on the cell boundary (Fig. S73a). We included all 130 genes in the analysis. For each cell in turn, we first obtained its nuclear center as the geometric center of all the nuclear boundary points. We computed its average nuclear radius ($r_n$) as the mean distance between the nuclear center and all nuclear boundary points. We computed its average cell radius ($r_c$) as the mean distance between the nuclear center and all cell boundary points. We further computed its nucleus-cell ratio as $r_n/r_c$. We excluded cells whose nucleus-cell ratio is beyond one standard deviation from the average ratio across all cells, yielding 887 cells for analysis (Fig. S73b). The estimated expression patterns, pattern clusters, pattern scores, sample genes and cells are shown in Fig. S74.

##### 5.2 ELLA analysis

We applied ELLA to analyze the preprocessed data (details in Methods). At an FDR of 5%, all genes are identified by ELLA to display subcellular expression patterns in the human U2-OS cells, including five transcription factors. The subcellular expression of the detected genes can be clustered into seven distinct expression pattern clusters (Fig. S74a): 95 genes (70%) display one of the four nuclear expression patterns (clusters 1-4), 10 (7%) genes display a nuclear edge expression pattern (cluster 5), and 30 genes (22%) display one of the two cytoplasmic expression patterns near the cellular membrane (clusters 6-7). Example cells from the seven clusters are shown in Fig. S74b. Specifically, the only long noncoding gene measured in this dataset, *MALAT1*, is identified to be nuclear localized (pattern 3). Three transcription factors measured in this dataset are detected to be nuclear localized (pattern 1), including the histone methyltransferase *PRDM2*, the BTB/POZ zinc-finger family gene *ZBTB43* involved in chromatin remodeling, transcriptional regulation, and the control of cell proliferation, as well as *AHDC1* which is known for its role in DNA binding. One transcription factor *ZNF592* is detected to be cytoplasmic localized (pattern 6). *ZNF592* encodes a member of the zinc finger protein family characterized by their ability to bind DNA, RNA, or protein via distinctive zinc finger motifs.

#### 6. Seq-Scope cell by cell analysis

Besides the cell by cell analysis in the seqFISH+ data, we also explored cell by cell analysis with high-quality cells in the Seq-Scope data.

##### 6.1 Preprocessing

Here, we applied the same data preprocessing steps as in the multiple-cell analysis to high-resolution 10X H&E images to obtain the corresponding cell segmentations. We again obtained nuclear centers based on unspliced expression intensity, registered cells with gene expressions to the unit circle, and extracted the relative positions of mRNAs as described before.

##### 6.2 Cell by cell analysis

We applied ELLA to analyze one cell at a time on two normal tissue tiles 2105, 2107 and two TD tissue tiles 2117, 2118, with 2105 as the main example. We included the top 20 highly expressed genes across all the four tiles for analysis. For ease of computation and simplicity, we used a customized set of 9 Beta kernel functions including Beta(1,4), Beta(1,2), Beta(2,5), Beta(2,3), Beta(2,2), Beta(3,2), Beta(5,2), Beta(2,1), and Beta(4,1), with a mode centering at 0, 0, 0.2, 0.33, 0.5, 0.67, 0.8, 1, 1 respectively. We visualized the *Alb* gene expression on tile 2105 (Fig. S75a) where the best fit kernel labels in each cell were marked out. A portion of cells achieved statistical significance with various kernels due to the potential heterogeneity of its subcellular localization across cells (Fig. S75b). The results of the remaining three tiles are shown in Fig. S76-78. *Alb* appears to be more likely nuclear localized on the TD tiles than the normal tiles.
